# Naturally occurring SARS-CoV-2 gene deletions close to the spike S1/S2 cleavage site in the viral quasispecies of COVID19 patients

**DOI:** 10.1101/2020.06.03.129585

**Authors:** Cristina Andres, Damir Garcia-Cehic, Josep Gregori, Maria Piñana, Francisco Rodriguez-Frias, Mercedes Guerrero-Murillo, Juliana Esperalba, Ariadna Rando, Lidia Goterris, Maria Gema Codina, Susanna Quer, Maria Carmen Martín, Magda Campins, Ricard Ferrer, Benito Almirante, Juan Ignacio Esteban, Tomás Pumarola, Andrés Antón, Josep Quer

## Abstract

The SARS-CoV-2 spike (S) protein, the viral mediator for binding and entry into the host cell, has sparked great interest as a target for vaccine development and treatments with neutralizing antibodies. Initial data suggest that the virus has low mutation rates, but its large genome could facilitate recombination, insertions, and deletions, as has been described in other coronaviruses. Here, we deep-sequenced the complete SARS-CoV-2 *S* gene from 18 patients (10 with mild and 8 with severe COVID-19), and found that the virus accumulates deletions upstream and very close to the S1/S2 cleavage site, generating a frameshift with appearance of a stop codon. These deletions were found in a small percentage of the viral quasispecies (2.2%) in samples from all the mild and only half the severe COVID-19 patients. Our results suggest that the virus may generate free S1 protein released to the circulation. We propose that natural selection has favored a “Don’t burn down the house” strategy, in which free S1 protein may compete with viral particles for the ACE2 receptor, thus reducing the severity of the infection and tissue damage without losing transmission capability.

## INTRODUCTION

RNA viruses replicate using their own RNA-dependent RNA polymerase (RdRp), which lacks proofreading mechanisms and is prone to mutate at high rates (10^−3^ - 10^−5^ substitutions/nucleotide/replication cycle), lending the virus a quasispecies structure(*1*, *2*). Previous studies with severe acute respiratory syndrome coronavirus (SARS-CoV) and mouse hepatitis viruses have reported moderate mutation rates of 9.06×10^−7^ and 2.5×10^−6^ subs/site/cycle respectively, below the expected range for RNA viruses(*3*). This is consistent with a role for non-structural protein (nsp) 14 in RNA proofreading or repair functions because of its 3’-5’ exonuclease (ExoN) activity. Nonetheless, the large size of the CoV RNA genome increases the probability that deletions will be generated and recombination events will take place, which could facilitate adaptation to new host environments, as occurs with jumping between species^1, 2^. One naturally occurring deletion on 29 nucleotides in the open reading frame (ORF) 8 of SARS-CoV after human-to-human transmission was found to be associated with attenuation of replication(*4*).

The low mutation rate, high human-to-human transmissibility (R_0_=2.2)(*5*), and absence of human pre-existing immunity against SARS-CoV-2 could explain its rapid spread through the human population, with very high sequence identity (99.9%) between isolates recovered all over the world (sequence published in the repository sequence data banks, GISAID and GenBank). The high pathogenicity of the virus, the severity of COVID-19 and the lack of an effective antiviral treatment or vaccine has pushed the scientific community worldwide to develop, in record time, a solution for this pandemic(*6*).

Among the SARS-CoV-2 structural proteins, including spike (S), envelope (E), and membrane (M) constituting the viral coat, and the nucleocapsid (N) protein that packages the viral genome, the S glycoprotein is the most promising as a therapeutic and vaccine target. The S protein is encoded by the *S* gene, and following trimerization, it composes the spikes of the characteristic viral particle crown (corona). The S protein is essential for SARS-CoV-2 to infect a host cell(*7*), by recognizing and binding to the human cell receptor, angiotensin-converting enzyme 2 (ACE2)(*8*), and possibly (with lower affinity) to other receptors, such as CD209L (L-SIGN), also used by SARS-CoV(*9*) and dipeptidyl peptidase 4 (DPP4), used by MERS(*10*).

The *S* gene has 3,822 nucleotides with 1,273 amino acids (GenBank reference sequence MN908947.3). It has five essential domains: the receptor-binding domain (RBD), O-linked glycan residues flanking a polybasic S1/S2 cleavage site, fusion peptide (FP), heptad repeats HR1 and HR2, and a transmembrane domain (TM). The S1 RBD includes 6 amino acid positions that show high affinity for the human ACE2 receptor, which is widely distributed, but mainly present in alveolar type 2 (AT2) cells of the lungs(*11*). Once the virus is attached to the host cell receptor cleavage occurs between subunits S1 and S2, and subunit S2 drives the viral and cellular membranes to fuse(*12*). Thus, S1 recognizes and binds to the human cell receptor ACE2, whereas S2 directly facilitates entry into the host cell. Both functions are crucial for infection, and therein lies the interest of S as a target for the development of vaccines and antiviral agents.

Because of the importance of the S protein, we carried out a deep-sequencing study of the *S* gene in upper respiratory tract samples from 18 patients with mild or severe SARS-CoV-2 disease. Of particular note, hot-spot deletion sites were found in minority mutants located upstream and very close to the S1/S2 and S2’ cleavage sites, suggesting that these genomes code for a truncated S protein. The variants were significantly more prevalent in patients with mild than those with severe disease. Thus, their effect on the protein could constitute a favorable regulatory mechanism emerging in the viral quasispecies to modulate the pathological effect of the infection. Discussion is provided on the implications this observation may have in the biology of SARS-CoV-2.

## PATIENTS AND METHODS

### PATIENTS

Upper respiratory tract specimens (naso/oropharyngeal swabs or nasopharyngeal aspirates) from individuals consulting in the emergency room were collected for SARS-CoV-2 testing in the Department of Microbiology at Hospital Universitari Vall d’Hebron (HUVH), Barcelona (Spain). Samples from 18 patients with no previous comorbidities other than COVID-19 were included in the study. As defined by CDC criteria (https://www.cdc.gov/coronavirus/2019-ncov/hcp/clinical-guidance-management-patients.html), 10 patients had a mild clinical presentation of COVID-19 (absence of viral pneumonia and hypoxia, no hospitalization requirement, able to manage their illness at home), whereas 8 patients had severe disease (ICU requirement for supportive management of complications of severe COVID-19, eg, pneumonia, hypoxemic respiratory failure, sepsis, cardiomyopathy and arrhythmia, acute kidney injury, and other complications). All patients, both those with mild and severe disease, had a favorable outcome with resolution of the infection.

### METHODS

#### Detection of SARS-CoV-2

The diagnosis of COVID-19 was performed by two tests, an in-house PCR assay using the primer/probe set from the CDC 2019-nCoV Real-Time RT-PCR Diagnostic Panel (Qiagen, Germany) and a commercial real-time RT-PCR assay, the Allplex 2019-nCoV Assay (Seegene, Korea).

#### SARS-CoV-2 sequencing

The eighteen respiratory specimens were inactivated by mixing 140 μL of sample with 560 μL of AVL buffer (Qiagen, Hilden, Germany). Extraction of nucleic acids was then performed using the QIAmp Viral RNA Mini Kit (Qiagen, Hilden, Germany) following the manufacturers’ instructions but without the RNA carrier, obtaining a final elution of 30 μL.

The complete *S* gene was amplified using a double PCR. The first RT-PCR step consisted in amplifying 2 large fragments, 3314 base pairs (bp) and 3591bp in length, respectively. The 5’ end of primer 1 and 3’ end of primer 2 were designed to be outside the *S* region to ensure that we were amplifying SARS-CoV-2 genomic RNA, and not subgenomic RNA. (Table S13).

The SuperScript III One-Step RT-PCR System with Platinum Taq HiFi DNA Polymerase (Invitrogen; Carlsbad, CA, USA) was used for the RT-PCR. Reverse transcription was done at 50°C for 30 min, followed by a retrotranscriptase inactivation step at 94°C for 2 min. Next, 30 cycles of PCR amplification were performed as follows: denaturation at 94°C for 15 sec, annealing at 54°C for 30 sec, and elongation at 68°C for 5 min. After the last cycle, amplification ended with a final elongation step at 68°C for 5 min.

The second round of amplification (nested) was done using overlapping internal primer pairs to amplify fragments 470bp to 313bp in length. The FastStart High-Fidelity PCR System dNTPack (Sigma, St. Louis, MO, CA) was used for this purpose, as follows: activation at 94°C for 4 min, followed by 30 cycles with denaturation at 94°C for 30 sec, annealing at 55°C for 30 sec, and elongation at 72°C for 40 sec, ending with a single elongation step at 72°C for 7 min.

PCR products were purified using the QIAquick Gel Extraction Kit (Qiagen, Hilden, Germany) with QG buffer, following the manufacturers’ instructions, and eluted DNA was quantified by fluorometry using the QUBIT dsDNA BR Assay Kit (ThermoFisher, MA, USA). For each patient, PCR products were normalized to 1.5 ng/μL, pooled in a single tube, and purified using KAPA Pure Beads (KapaBiosystems, Roche, Pleasanton, CA, USA) to ensure that no short DNA fragments were present in the library. Library preparation was done using the KAPA Hyper Prep Kit (Roche Applied Science, Pleasanton, CA, USA) and each pool was individually indexed using the SeqCap Adapter Kit A/B (Nimblegen, Roche, Pleasanton, CA USA). After library enrichment and a second clean-up with KAPA Pure Beads, the pools were quantified again using the QUBIT dsDNA BR Assay Kit and quality-tested using the 4150 TapeStation System (Agilent). All pools underwent a final normalization to 4 nM, and 10 μL of each pool was added to the final library tube. The final library was qPCR-quantified using the KAPA Library Quantification Kit (KapaBiosystems, Roche, Pleasanton, CA USA) in a LightCycler 480 system (Roche) to obtain the precise concentration of indexed DNA. PhiX internal DNA control (*PhiX V3*, Illumina, San Diego, CA) was added to the final dilution. The library was loaded in a MiSeq Reagent Kit 600V3 cartridge (Illumina, San Diego, CA) and sequenced using the MiSeq platform (Illumina, San Diego, CA).

#### Bioinformatics analysis. InDel study

The sequence analysis aimed to obtain high-quality haplotypes fully covering the amplicons. The pipeline comprises the following steps:

1. Amplicons were reconstructed from the corresponding R1 and R2 paired ends using FLASH(*13*) and setting a minimum of 20 overlapping bases and a maximum of 10% mismatches. Low-quality reads that did not match meet the requirements were discarded.
2. Next, all reads with more than 5% of bases below a Phred score of Q30 were filtered out.
3. The reads were demultiplexed by matching primers, allowing a maximum of three mismatches, and the primers were trimmed at both read ends. Identical reads were collapsed to haplotypes with the corresponding frequencies as read counts. A fasta file was generated with each pool/primer/strand combination. The reverse haplotypes were reverse complemented.
4. Raw forward and reverse haplotypes were multiple aligned with **MU**ltiple **S**equence **C**omparison by **L**og-**E**xpectation (Muscle)(*14*), then separated into strands, and haplotypes common to both strands at abundances ≥0.1% were identified. Low-abundance haplotypes (<0.1%) and those unique to one strand were discarded. The haplotypes common to both strands, with frequencies not below 0.1% are called consensus haplotypes, and are the basis of subsequent computations.

The amino acid alignments were computed as follows:

1. Gaps were removed and haplotypes translated to amino acids.
2. The translated stops generated were identified, and haplotypes were trimmed after the stop.
3. Resulting amino acid haplotypes were realigned with Muscle (EMBL-EBI https://www.ebi.ac.uk/Tools/msa/muscle/).

All computations were made in the R language and platform(*15*), developing in-house scripts using Biostrings(*16*) and Ape(*17*) packages.

## RESULTS

Eighteen coronavirus disease-19 (COVID-19) patients (10 mild, 8 severe) were included in the study. In total, 48,746,647 reads, ranging from 81,202 to 597,558 reads per amplicon (median 171,478), were analyzed from upper respiratory tract samples (Table 1), using 13 overlapping amplicons covering the complete S protein. Thus, we studied 3,749,742 complete S genes, with a mean of 208,319 per patient. Results related to amplicon positions, coverage, percentage of the master sequence, gap incidence per patient and per amplicon, and premature stop codons are available as Supplementary Tables S1-S10.

**Table 1.** Patients’ characteristics with mild and severe COVID-19. *This patient was still at the ICU at the time this manuscript was submitted. #P16 had clinical symptoms according to a severe disease, but he was not physically hospitalized into the ICU.

| Sample Id<br>(P=patient) | Real-time PCR<br>cycle<br>threshold (Ct)<br>value | COVID-19<br>classifica<br>tion | Sample Type | Sex<br>(F=female;<br>M=male) | Age<br>(years) | Days at<br>Intensive<br>Care Unit<br>(ICU) |
| --- | --- | --- | --- | --- | --- | --- |
| P01 | 19,00 | mild | nasopharyngeal aspirate | F | 34 | no admission |
| P02 | 25,00 | mild | nasopharyngeal aspirate | F | 54 | no admission |
| P03 | 16,40 | mild | nasopharyngeal/oropharyngeal swab | F | 42 | no admission |
| P04 | 23,10 | mild | nasopharyngeal/oropharyngeal swab | F | 25 | no admission |
| P05 | 25,98 | mild | nasopharyngeal/oropharyngeal swab | M | 52 | no admission |
| P06 | 21,45 | mild | nasopharyngeal/oropharyngeal swab | F | 42 | no admission |
| P07 | 25,94 | mild | nasopharyngeal/oropharyngeal swab | F | 25 | no admission |
| P14 | 23,71 | mild | nasopharyngeal aspirate | F | 26 | no admission |
| P15 | 27,32 | mild | nasopharyngeal/oropharyngeal swab | M | 41 | no admission |
| P18 | 15,50 | mild | nasopharyngeal/oropharyngeal swab | M | 74 | no admission |
| P08 | No data | severe | nasopharyngeal/oropharyngeal swab | F | 51 | 4 |
| P09 | 25,36 | severe | nasopharyngeal/oropharyngeal swab | M | 49 | 3 |
| P10 | 21,23 | severe | nasopharyngeal/oropharyngeal swab | F | 47 | 16 |
| P11 | 36,01 | severe | nasopharyngeal/oropharyngeal swab | M | 45 | 27 |
| P12 | 31,04 | severe | nasopharyngeal/oropharyngeal swab | M | 51 | 23 |
| P13 | 22,94 | severe | nasopharyngeal aspirate | F | 44 | >29* |
| P16 | 34,35 | severe | nasopharyngeal/oropharyngeal swab | F | 45 | # |
| P17 | 30,77 | severe | nasopharyngeal/oropharyngeal swab | M | 49 | 10 |

Deletions were not randomly accumulated along the S gene, but instead, were found at specific regions (Figure 1, Figures S1-S27). Deletions coded as delta (Δ1-Δ18) ranged from 1 to 42 nucleotides lost (Table 2). In some cases, the sequence recovered the correct reading frame, in others, the frameshift caused the appearance of a premature stop codon very close to the deletion site, whereas in still others, a new amino acid segment appeared.

**Figure 1.**
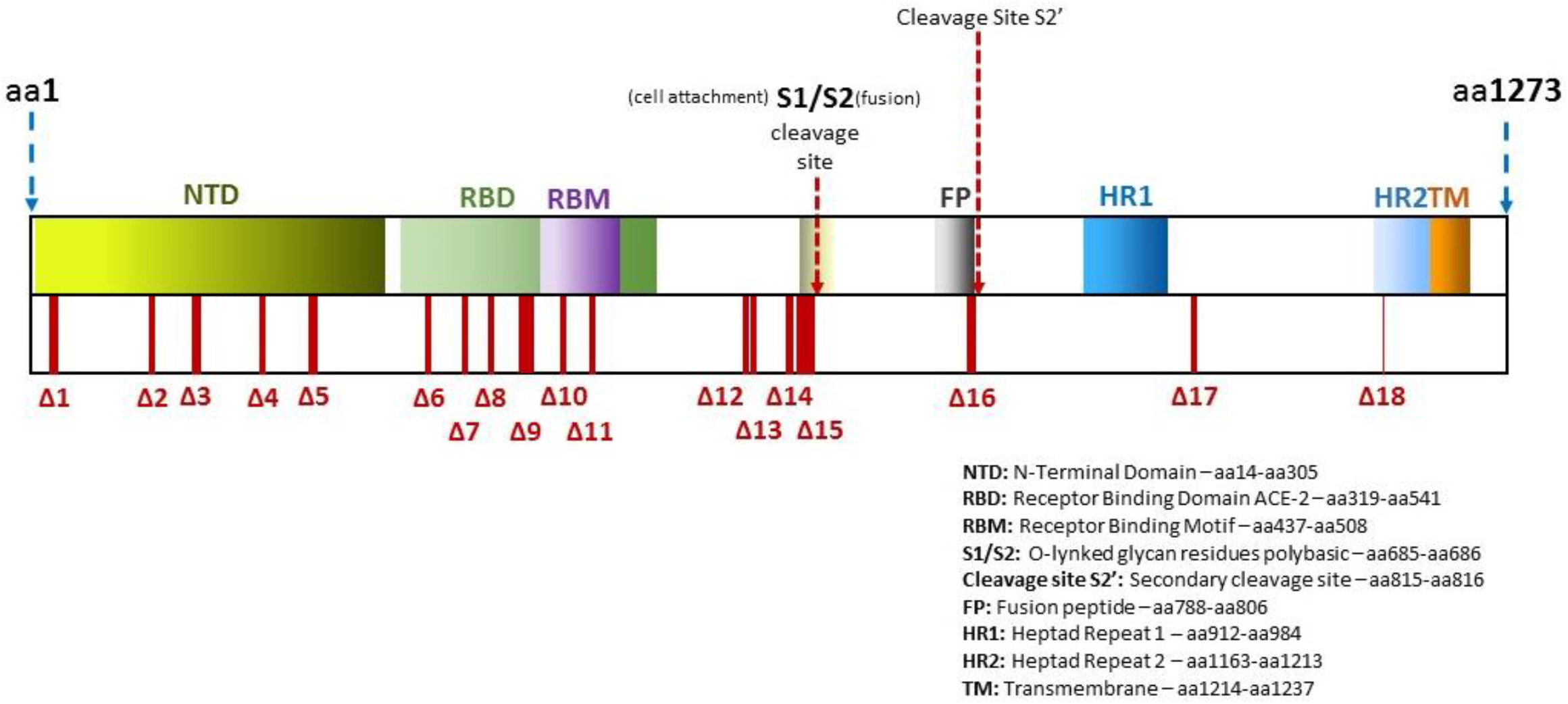
Schematic location of deletions found along the Spike gene and protein(*28*).

**Table 2.**
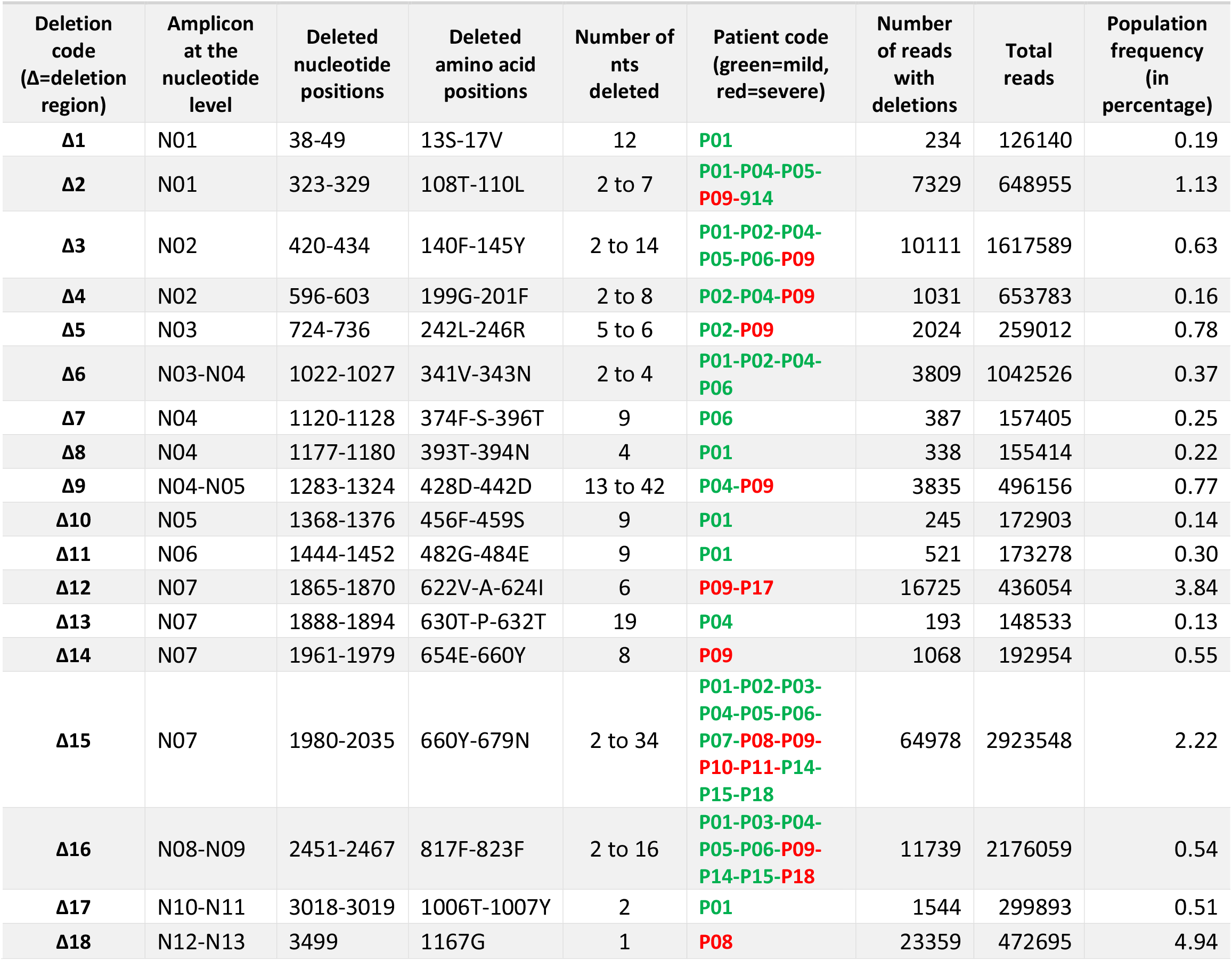
List of deletions found along the spike gene

| Deletion code<br>(Δ=deletion region) | Amplicon at the nucleotide level | Deleted nucleotide positions | Deleted amino acid positions | Number of nts deleted | Patient code (green=mild, red=severe) | Number of reads with deletions | Total reads | Population frequency (in percentage) |
| --- | --- | --- | --- | --- | --- | --- | --- | --- |
| Δ1 | N01 | 38-49 | 13S-17V | 12 | P01 | 234 | 126140 | 0.19 |
| Δ2 | N01 | 323-329 | 108T-110L | 2 to 7 | P01-P04-P05-P09-914 | 7329 | 648955 | 1.13 |
| Δ3 | N02 | 420-434 | 140F-145Y | 2 to 14 | P01-P02-P04-P05-P06-P09 | 10111 | 1617589 | 0.63 |
| Δ4 | N02 | 596-603 | 199G-201F | 2 to 8 | P02-P04-P09 | 1031 | 653783 | 0.16 |
| Δ5 | N03 | 724-736 | 242L-246R | 5 to 6 | P02-P09 | 2024 | 259012 | 0.78 |
| Δ6 | N03-N04 | 1022-1027 | 341V-343N | 2 to 4 | P01-P02-P04-P06 | 3809 | 1042526 | 0.37 |
| Δ7 | N04 | 1120-1128 | 374F-S-396T | 9 | P06 | 387 | 157405 | 0.25 |
| Δ8 | N04 | 1177-1180 | 393T-394N | 4 | P01 | 338 | 155414 | 0.22 |
| Δ9 | N04-N05 | 1283-1324 | 428D-442D | 13 to 42 | P04-P09 | 3835 | 496156 | 0.77 |
| Δ10 | N05 | 1368-1376 | 456F-459S | 9 | P01 | 245 | 172903 | 0.14 |
| Δ11 | N06 | 1444-1452 | 482G-484E | 9 | P01 | 521 | 173278 | 0.30 |
| Δ12 | N07 | 1865-1870 | 622V-A-624I | 6 | P09-P17 | 16725 | 436054 | 3.84 |
| Δ13 | N07 | 1888-1894 | 630T-P-632T | 19 | P04 | 193 | 148533 | 0.13 |
| Δ14 | N07 | 1961-1979 | 654E-660Y | 8 | P09 | 1068 | 192954 | 0.55 |
| Δ15 | N07 | 1980-2035 | 660Y-679N | 2 to 34 | P01-P02-P03-P04-P05-P06-P07-P08-P09-P10-P11-P14-P15-P18 | 64978 | 2923548 | 2.22 |
| Δ16 | N08-N09 | 2451-2467 | 817F-823F | 2 to 16 | P01-P03-P04-P05-P06-P09-P14-P15-P18 | 11739 | 2176059 | 0.54 |
| Δ17 | N10-N11 | 3018-3019 | 1006T-1007Y | 2 | P01 | 1544 | 299893 | 0.51 |
| Δ18 | N12-N13 | 3499 | 1167G | 1 | P08 | 23359 | 472695 | 4.94 |

Deletions were found in all amplicons, but they were mainly observed at frequencies <1% (Table 2). Most deletions in amplicons N04, N05, N06, N10, N11, N12, and N13, were found in only 1 or 2 patients, whereas deletions in amplicons N01, N02 N08 and N09, ranging from 2 to 16 nucleotides, were observed in 4 to 9 patients. A deletion of 6 nucleotides in amplicon N07 (nt 1865-1870), generating a stop codon, was present at a frequency of 3.84% of the quasispecies in samples from patients P09 and P17. The largest deletion, involving 42 nucleotides (nt 1283-1324) and found in N05 of patient P09, resulted in a loss of 14 amino acids, but the reading frame recovered.

A striking result was the accumulation of deletions (“hot-spot”) in amplicon N07, between nucleotides 1980-2035 (aa Y660-N679) in 14/18 (78%) patients, which included 100% of the patients with mild disease (P01-P07, P14, P15 and P18), and only half of those with severe disease (P08, P09, P10 and P11). In this particular hot-spot, deletions Δ2 to Δ34 were produced (Figure 2, Table 2). Among the severe patients, P12, P13, and P16 had no deletions in the N07 amplicon, and P17 showed a deletion outside this hot-spot location (Table S11). Viral variants carrying these deletions were significantly more frequent in mild than severe COVID-19 patients (Fisher test: odds-ratio: 95% confidence interval 0.0 - 0.9605; p=0.02288) (Table S12).

**Figure 2.**
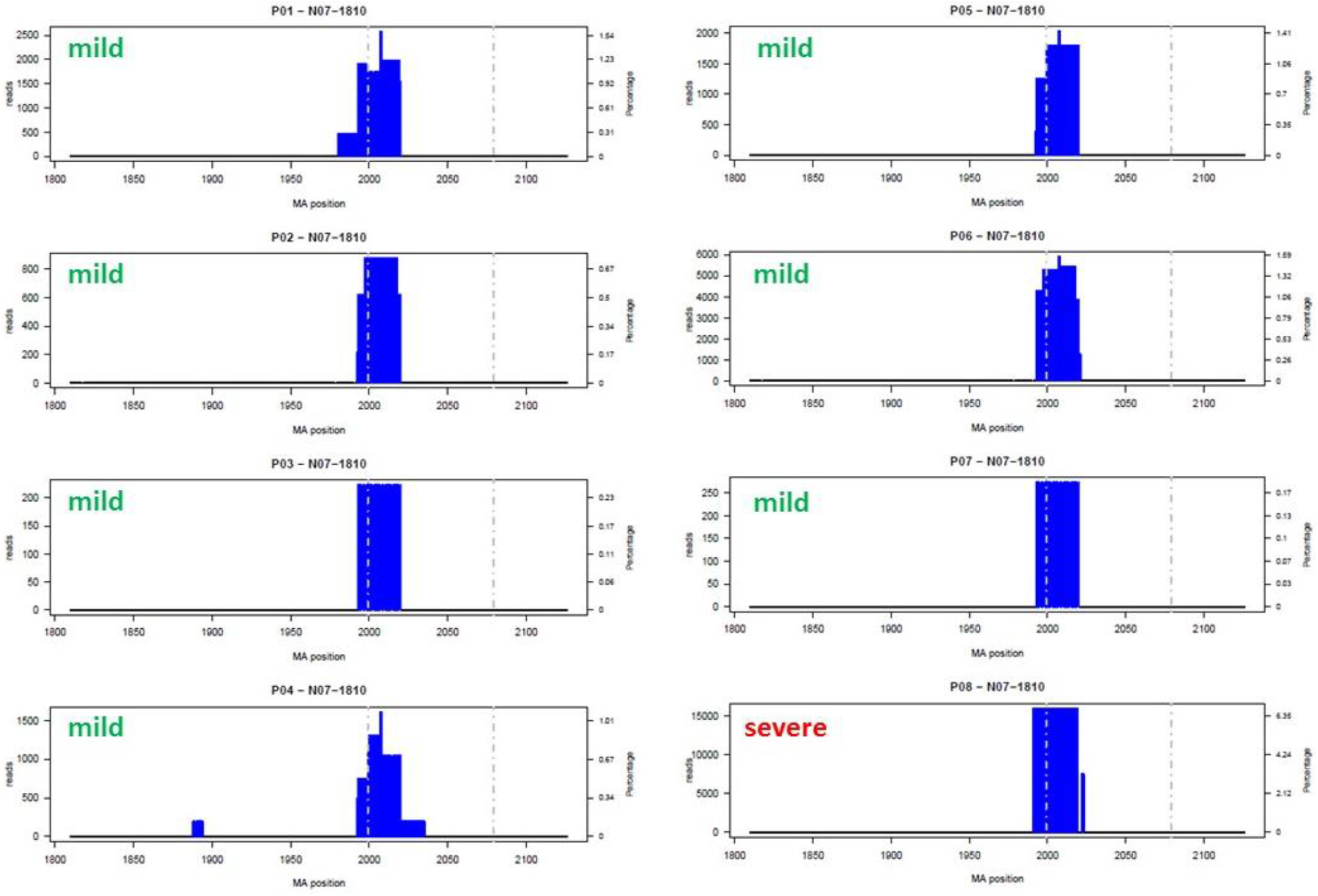
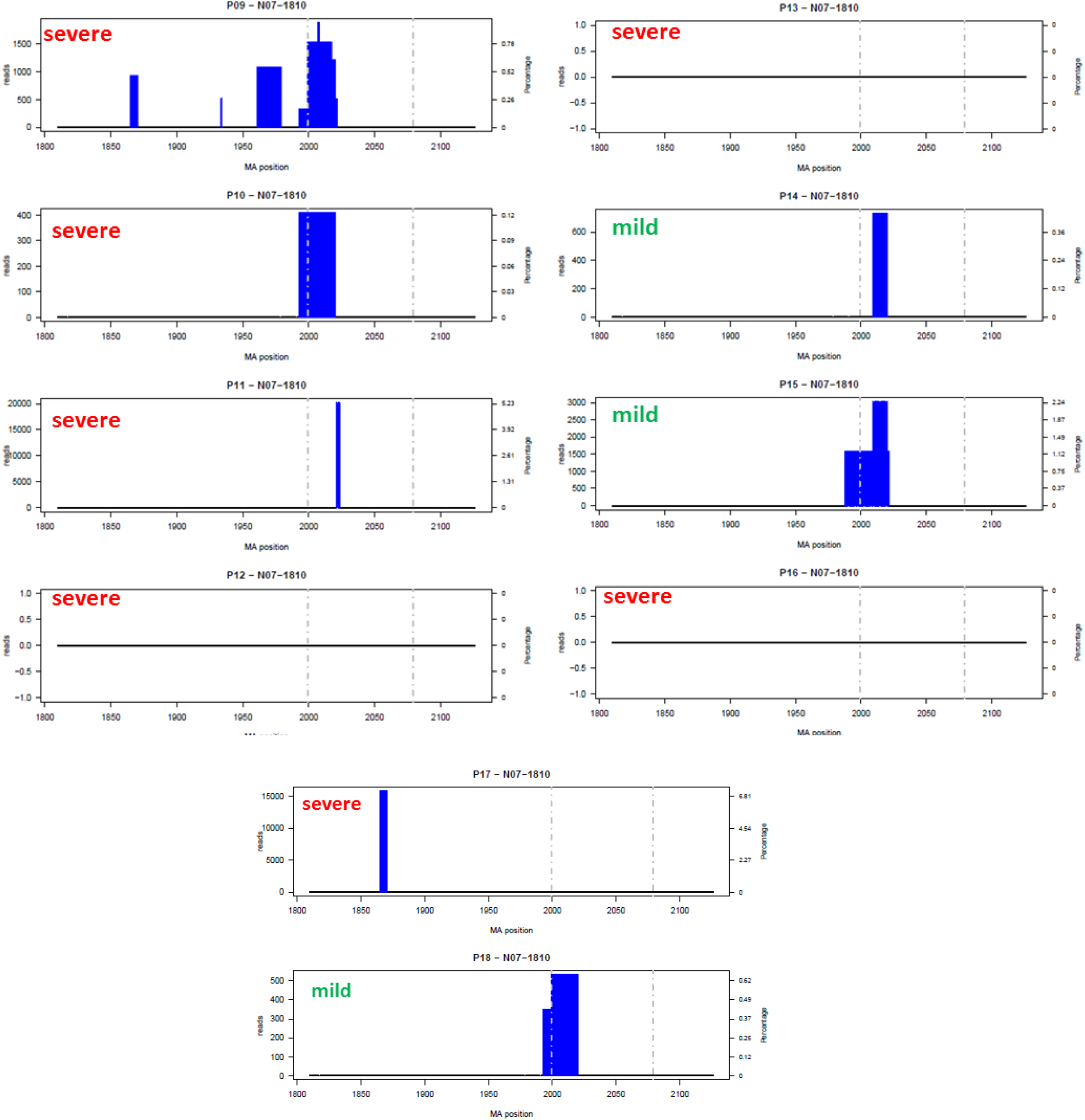
Bar plot of deletions in amplicon N07 in the 18 patients (P01-P18) at the nucleotide level. The x axis provides the multiple alignment (MA) nucleotide positions and the amplitude of the deletions by subregions, and the y axis shows the frequency of the deletion (percentage) on the right and the number of reads on the left. As no insertions were observed the MA positions correspond to S gene positions. Dashed lines indicates S1/S2 (left) and S2’ (right) cleavage sites. Bar plots for the 18 patients and by amplicons are provided in supplementary material (Figures S1 to S14 for nucleotides and S15 to S27 for amino acids).

Among the total of 43, Δ15 deletions in amplicon N07, a premature stop codon appeared immediately after the deletion site in five cases (11.6%) and the reading frame recovered after losing 4, 5, or 7 amino acids in six cases. However, a frameshift that changed the reading frame and caused the appearance of a premature stop codon several amino acids later was generated in most of the deletions 32/43 (74.4%), and in consequence the S1/S2 cleavage site and the polybasic domain (PRRAR/S) disappeared. In 39 of the 43 (90.7%) N07 deletions, a TATA box-like motif (nt 2,007-2,010) was lost. In this particular region, the deletion was characterized by a similar 3’ cutting edge (Table S11). An interesting result at the amino acid level was that regardless of the starting point of the deletion (nt 654, 663, 664, 665, 666, 667 or 671), in 9 of the mild patients (all except P14) and in 2 severe ones (P09 and P10), the frameshift caused by the deletion generated a new peptide motif, RLRLILLGGHVV*, with a stop codon (*) at the end (Table S11).

In 9 patients, a second deletion hot-spot was found deleting a number of nucleotides (from 2 to 16) between positions 2451 and 2467 (aa 817F-823F), coinciding with the secondary S cleavage site (S2’). The hot spot was located between nt2431 (811K) and nt2454 (818I), just after the exact S2’ cleavage site (KPSKR/SFI) (Table 2).

## DISCUSSION

Here, we describe the naturally occurring deletions in the SARS-CoV-2 S gene in a set of patients with mild or severe COVID 19. The deletions mainly clustered in two hot-spot regions, one (Δ15, affecting aa660-aa679) located upstream but very close to the S1/S2 cleavage site (aa 685/686) and the second (Δ16 affecting aa817-823) situated just upstream to the secondary cleavage site S2’ (aa 815/816). These two deletions were found in most of the patient samples studied, and notably, the Δ15 deletion was present in 100% of patients with mild infection and in half of those with severe disease, three fourth of the studied patients (Table 2). This finding suggests that the deletions are not sporadic events even though these deletions were seen in a relatively small percentage of the viral quasispecies (2.2% for Δ15; and 0.54% for Δ16). The mutants could be interpreted as a strategy that natural selection has favored during the SARS-CoV-2 infectious life cycle to facilitate extensive spread of the infection, as is discussed below.

This study involved deep-sequencing of the complete SARS-CoV-2 spike gene using 13 overlapping amplicons in laboratory confirmed samples for SARS-CoV-2 in 18 patients. In studies with other SARS-CoV viruses, several subgenomic RNAs were reported to be generated during the cell cycle(*6*, *18*). To exclusively study the genomic viral RNA of SARS-CoV-2, RT-PCR was performed using two large PCR products in which the 5’ end of primer pair 1 and the 3’ end of primer pair 2 were designed to be outside the spike region (5’ end in ORF1ab and 3’ end in ORF3a) (Table S13, Figure S28). Taking into consideration that CoV have 3’-5’ ExoN activity (nsp14 protein), consistent with a proofreading mechanism to correct mutations during replication, the deep-sequencing analysis accepted mutants present at a low frequency of ≥0.1%. Because the possibility of PCR artefacts, deep-sequencing point mutations, or deletion of single nucleotides generated mainly at homopolymeric sites, we did not include single deletions unless they were found in different patients and in overlapping amplicons at higher frequencies (>1%). No insertions were found.

Entry of the viral genome into the cell depends on recognition and binding of the surface subunit S1 to the ACE2 human receptor(*11*), whereas the S2 subunit is responsible for fixing the S protein to the viral membrane surface. After binding to the ACE2 cell receptor, the S protein is primed by the serine-protease, TMPRSS2, which leads to S protein cleavage at S1/S2 and S2’(*8*). After cleavage, S1 remains attached to ACE2, while subunit S2 anchors the viral and cellular membranes, inducing fusion and viral entry. The Δ15 deletion (Table 2) mainly causes a frameshift that generates an in-frame stop codon. The presence of this new stop codon would result in translation of a truncated S, which would consist of an almost complete S1 subunit, and total absence of the S2 subunit responsible for anchoring S to the lipid membrane of the viral particle. The absence of the S2 anchor peptide suggests that S1 could be produced as a “free” protein (free S1). As S1 is located on the exposed outside of SARS-CoV-2 in the crown structures, it could have hydrophilic domains and be a soluble peptide with potential for release outside the infected cell, in the lower respiratory tract and even to plasma. (Figure 3). These free soluble proteins, which are not a part of the viral cycle or components of the viral particles have also been observed in other viral infections. For example, a huge amount of “empty” subviral genomic particles, consisting of viral envelope proteins (HBsAg), are often found in plasma of patients with hepatitis B virus (HBV) infection. These empty particles are produced and secreted during HBV infection, and have an immunomodulatory role(*19*). In addition, soluble HBV e antigen (HBeAg), which is not a component of the viral particles and shares immunoactive epitopes with the HBV core antigen (HBcAg viral capsid component), is detected during HBV infection and has an immunomodulator role(*20*).

**Figure 3.**
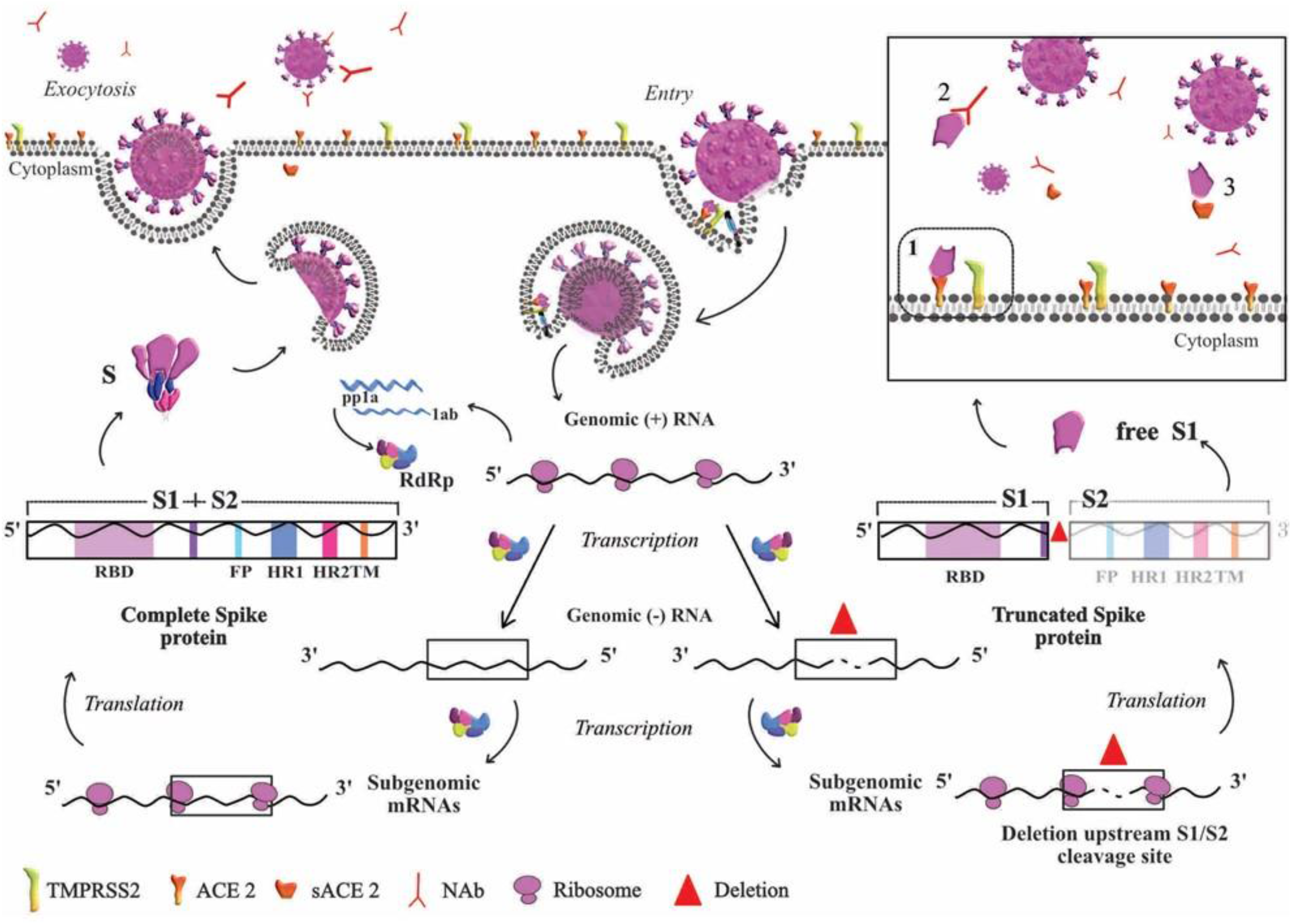
Based on the life cycle of SARS-CoV, this diagram represents the hypothesis derived from our results. Entry of the virus in the host cell is shown at the top right of the diagram. At the transcription step, two scenarios are depicted: to the left, the viral particle resulting from normal S protein, and to the right the viral particle resulting from truncated S. In normal conditions, once the nucleoprotein is freed into the cytoplasm ss+RNA is translated into the non-structural proteins required for transcription. ss+RNA is transcribed into ss-RNA and later into genomic ss+RNA which is encapsidated (left side of the figure). Once the complete viral particle has been formed, it is secreted from the cell by exocytosis. The right side of the figure depicts the situation when a deletion occurs in the S gene during transcription of the complete genome and before subgenomic mRNAs are generated to produce the structural proteins. Translation of a deleted subgenomic spike mRNA would lead to a truncated S protein composed of the S1 domain without S2, which could be shed outside the cell as free S1. The box depicts possible destinations of free S1, which could bind to 1) the ACE2 cell receptor, 2) S1-specific neutralizing antibodies, or 3) free ACE2 receptor. The red triangle indicates the deletion in genomic RNA. Abbreviations: ACE2, angiotensin converting enzyme 2; mRNA, messenger RNA; NAb; neutralizing antibodies; pp1a, polyprotein 1a; RdRp, RNA-dependent RNA polymerase; S, spike; S1, subunit S1 at the N-terminal domain of the S protein, which includes receptor binding domain (RBD); S2, subunit S2 located at the C-terminal domain of S protein, which includes fusion peptide (FP), heptad repeat (HR) domain 1 and 2, and the transmembrane domain (TM); ss, single stranded; ss+RNA, single-stranded positive sense RNA; TMPRS22, human serine protease TMPRSS2

Human respiratory syncytial virus (HRSV) is another respiratory virus with the ability to produce pre-anchored proteins. The attachment protein (G) of HRSV is an anchored protein whose main function is viral attachment to the host’s cell membrane through a still unknown receptor(*21*). As in many other viruses, this protein has several functions, and in this case, because of the existence of a second start codon, a soluble form of G protein lacking the anchor is produced, and this is shed to the extracellular medium(*22*) in abundant quantities by infected cells. The function of soluble, free G is to inhibit toll-like receptors, thereby modulating the host’s immune response. Free G also binds to the host’s neutralizing antibodies, which are mainly directed to this protein. In this way, neutralization of circulating virions is reduced, favoring viral infection(*23*).

The free S1 binding subunit of SARS-CoV-2 without its membrane anchor S2 could have similar functions (Figure 3). One putative action of secreted free S1 protein might be to attach to the human ACE2 cell receptor, thereby competing with complete viral particles to re-infect or newly infect respiratory tract cells, resulting in less severe disease. This could be interpreted as an effect of natural selection to attenuate the infection and facilitate its persistence with minimal damage, increasing the human-to-human transmission into the community. This strategy, which we have dubbed “Don’t burn down the house” is supported by the finding that these minor variants carrying these deletions were statistically more frequent in patients with mild that severe COVID19. This self-modulating viral strategy has also been seen in hepatitis delta virus (HDV) infection, where one viral antigen (short HDV antigen SHDAg) enhances HDV replication, while a second antigen (large HDV antigen LHDAg), produced after a stop codon edition (TAG to TGG) by cellular adenosine deaminase, acts as a negative regulator of replication(*24*).

The fact that the truncated S protein was present in only a low percentage of the entire viral quasispecies suggests that natural selection may have designed a favorable equilibrium in which a limited number of deleted virions are generated to balance virus production with infection of new cells during disease progression. A likely reason for maintaining a minority population of genomes with deletions able to produce free S1 protein would be to infect a host while causing minimal damage, which would greatly facilitate transmission of the virus within the population. It would be of interest to determine whether the percentage of these viral mutants changes during disease progression and whether there are differences between the quasispecies isolated from upper and lower respiratory tract specimens. As a consequence of the frameshift a new peptide motif, RLRLILLGGHVV* appeared in several sequences with a deletion that started in different nucleotide points. Additional work is needed to determine whether acquisition of this peptide motif has biological consequences.

Two other putative consequences of the *S* mutants might be that free S1 protein could bind with S-specific antibodies, acting as a decoy and weakening the immune response, or to circulating ACE2, released from the cell membrane to plasma(*25*, *26*), with cardiovascular effects. However, as the deletions were mainly found in patients with mild disease and considering the zoonotic origin of the virus (animal immune and cardiac systems differ from human ones) and the short time that the virus has been evolving in the human population, we believe that the most likely reason for maintaining a minor population of mutant genomes able to produce free S1 protein would be to cause an infection with limited damage in the host, thus facilitating transmission and persistence of the virus in the population. The observation of mutation hot spots in the S gene opens the door to further work on a number of potentially related aspects.

In a recent study (27), Lau et al reported the results of plaque purification of Vero-E6 cultured SARS-CoV-2 genomes obtained from nasopharyngeal aspirate of a COVID-19 patient. The authors found deletions of 10 to 15 nucleotides at the S1/S2 junction. In a further experiment, infection of hamsters with virus containing these variants led to attenuated viral disease. These findings strongly support our hypothesis that deletions close to the S1/S2 cleavage site may be a phenomenon favoured by natural selection to enhance spread of the SARS-CoV-2. The authors failed to detect these deletions in this and other clinical specimens, but this may be attributable to the relatively low sequencing throughput of the Sanger technique used. In the cell culture experiments, the lack of genomes with deletions that generate a premature stop codon in the S gene can be easily explained, as the truncated S gene would produce non-infectious particles.

To conclude, in-depth sequencing of the SARS-CoV-2 *S* gene in 18 patients with COVID-19 enabled identification of a naturally occurring deletion very close to the S1/S2 cleavage site. Our results indicate that the mutant S would have a large impact on the S protein, and suggest that the virus could produce free S1, which may have implications regarding the candidacy of S protein as a target for vaccination and antiviral treatment strategies. The deletions were significantly more prevalent in patients with mild than in those with severe disease, supporting the notion that they are a strategy of natural selection to decrease the injury caused after onset of the infection. In this “Don’t burn down the house” strategy, the ability of the virus to bind with ACE2 receptor and spread to others would be unchanged; thus its propensity for transmission would be enhanced by a mildly affected host. To prove this hypothesis, it is essential to further investigate whether the truncated S protein (free S1) is present in respiratory tract specimens and in plasma.

## Acknowledgements

This work was partially supported by the Direcció General de Recerca i Innovació en Salut (DGRIS) Catalan Health Ministry Generalitat de Catalunya through Vall d’Hebron Research Institute (VHIR), European Development Regional Fund (ERDF) “A way to achieve Europe”, by Spanish Network for Research in Infectious Diseases (REIPI RD16/0016/0003) and Centro para el Desarrollo Tecnológico Industrial (CDTI) from the Spanish Ministry of Economy and Business, grant number IDI-20200297.

The authors thank Celine Cavallo for English language support.

## Author Contributions

CA, DG-C, MP and MCM have significantly contributed to designing the experimental work and performed the technical work involving RNA extraction, amplification, and deep sequencing JG and MG-M developed the software used in the study

FRF, JE, AR, LG, BA, RF and MGC collected samples and actively participated in discussions and corrections related to the draft

SQ designed the graphics and actively participated in the draft discussion and corrections.

MC, LG, BA, RF, JIE, and TP participated in the analysis and interpretation of the data, and substantially revised the draft

AA and JQ conceived the work, designed the primers, led the data analysis, wrote the draft, and led the discussion.

## Ethics Committee approval

The work has been approved by Vall d’Hebron University Hospital Ethical Committee reference number PR(AG)259-2020.

## REFERENCES

1. J. J. Holland, Replication error, quasispecies populations and extreme evolution rates of RNA viruses (Oxford University Press, Oxford. UK, 1993).

2. E. Domingo, C. Escarmis, N. Sevilla, A. Moya, S. F. Elena, J. Quer, I. S. Novella, J. J. Holland, Basic concepts in RNA virus evolution. FASEB J. 10(1996).

3. L. D. Eckerle, M. M. Becker, R. A. Halpin, K. Li, E. Venter, X. Lu, S. Scherbakova, R. L. Graham, R. S. Baric, T. B. Stockwell, D. J. Spiro, M. R. Denison, Infidelity of SARS-CoV Nsp14-exonuclease mutant virus replication is revealed by complete genome sequencing. PLoS Pathog. 6, e1000896 (2010).

4. D. Muth, V. M. Corman, H. Roth, T. Binger, R. Dijkman, L. T. Gottula, F. Gloza-Rausch, A. Balboni, M. Battilani, D. Rihtaric, I. Toplak, R. S. Ameneiros, A. Pfeifer, V. Thiel, J. F. Drexler, M. A. Muller, C. Drosten, Attenuation of replication by a 29 nucleotide deletion in SARS-coronavirus acquired during the early stages of human-to-human transmission. Sci. Rep. 8, 15177 (2018).

5. A. S. Fauci, H. C. Lane, R. R. Redfield, Covid-19 - Navigating the Uncharted. N. Engl. J. Med. 382 (2020), pp. 1268–1269.

6. Y.-F. Tu, C.-S. Chien, A. A. Yarmishyn, Y.-Y. Lin, Y.-H. Luo, Y.-T. Lin, W.-Y. Lai, D.-M. Yang, S.-J. Chou, Y.-P. Yang, M.-L. Wang, S.-H. Chiou, A Review of SARS-CoV-2 and the Ongoing Clinical Trials. Int. J. Mol. Sci. 21 (2020), doi:10.3390/ijms21072657.

7. S. Belouzard, J. K. Millet, B. N. Licitra, G. R. Whittaker, Mechanisms of coronavirus cell entry mediated by the viral spike protein. Viruses. 4, 1011–1033 (2012).

8. M. Hoffmann, H. Kleine-Weber, S. Schroeder, N. Kruger, T. Herrler, S. Erichsen, T. S. Schiergens, G. Herrler, N.-H. Wu, A. Nitsche, M. A. Muller, C. Drosten, S. Pohlmann, SARS-CoV-2 Cell Entry Depends on ACE2 and TMPRSS2 and Is Blocked by a Clinically Proven Protease Inhibitor. Cell. 181, 271–280.e8 (2020).

9. S. A. Jeffers, S. M. Tusell, L. Gillim-Ross, E. M. Hemmila, J. E. Achenbach, G. J. Babcock, W. D. J. Thomas, L. B. Thackray, M. D. Young, R. J. Mason, D. M. Ambrosino, D. E. Wentworth, J. C. Demartini, K. V Holmes, CD209L (L-SIGN) is a receptor for severe acute respiratory syndrome coronavirus. Proc. Natl. Acad. Sci. U. S. A. 101, 15748–15753 (2004).

10. E. de Wit, N. van Doremalen, D. Falzarano, V. J. Munster, SARS and MERS: recent insights into emerging coronaviruses. Nat. Rev. Microbiol. 14, 523–534 (2016).

11. K. G. Andersen, A. Rambaut, W. I. Lipkin, E. C. Holmes, R. F. Garry, The proximal origin of SARS-CoV-2. Nat. Med. 26 (2020), pp. 450–452.

12. T. Heald-Sargent, T. Gallagher, Ready, set, fuse! The coronavirus spike protein and acquisition of fusion competence. Viruses. 4, 557–580 (2012).

13. T. Magoc, S. L. Salzberg, FLASH: fast length adjustment of short reads to improve genome assemblies. Bioinformatics. 27, 2957–2963 (2011).

14. R. C. Edgar, MUSCLE: a multiple sequence alignment method with reduced time and space complexity. BMC.Bioinformatics. 5, 113 (2004).

15. R. C. Team, R: A language and environment for statistical computing. (2016).

16. H. Pages, P. Aboyoun, R. Gentleman, S. DebRoy, ‘Biostrings:String objects representing biological sequences, and matching algorithms. R package 2.38.4’ (2012).

17. E. Paradis, J. Claude, K. Strimmer, APE: Analyses of Phylogenetics and Evolution in R language. Bioinformatics. 20, 289–290 (2004).

18. Z. Song, Y. Xu, L. Bao, L. Zhang, P. Yu, Y. Qu, H. Zhu, W. Zhao, Y. Han, C. Qin, From SARS to MERS, Thrusting Coronaviruses into the Spotlight. Viruses. 11 (2019), doi:10.3390/v11010059.

19. A. Caballero, D. Tabernero, M. Buti, F. Rodriguez-Frias, Hepatitis B virus: The challenge of an ancient virus with multiple faces and a remarkable replication strategy. Antiviral Res. 158, 34–44 (2018).

20. A. Kramvis, E.-G. Kostaki, A. Hatzakis, D. Paraskevis, Immunomodulatory Function of HBeAg Related to Short-Sighted Evolution, Transmissibility, and Clinical Manifestation of Hepatitis B Virus. Front. Microbiol. 9, 2521 (2018).

21. A. T. Borchers, C. Chang, M. E. Gershwin, L. J. Gershwin, Respiratory syncytial virus--a comprehensive review. Clin. Rev. Allergy Immunol. 45, 331–379 (2013).

22. D. A. Hendricks, K. McIntosh, J. L. Patterson, Further characterization of the soluble form of the G glycoprotein of respiratory syncytial virus. J. Virol. 62, 2228–2233 (1988).

23. F. P. Polack, P. M. Irusta, S. J. Hoffman, M. P. Schiatti, G. A. Melendi, M. F. Delgado, F. R. Laham, B. Thumar, R. M. Hendry, J. A. Melero, R. A. Karron, P. L. Collins, S. R. Kleeberger, The cysteine-rich region of respiratory syncytial virus attachment protein inhibits innate immunity elicited by the virus and endotoxin. Proc. Natl. Acad. Sci. U. S. A. 102, 8996–9001 (2005).

24. S. A. Hughes, H. Wedemeyer, P. M. Harrison, Hepatitis delta virus. Lancet (London, England). 378, 73–85 (2011).

25. A. Turner, in The Protective Arm of the Renin Angiotensin System (RAS) Functional Aspects and Therapeutic Implications, R. A. S. dos Thomas Unger, Ulrike M. Steckelings, Santos, Ed. (Elsevier Inc., 2015), pp. 185–189.

26. L. Matsushita, K; Ding, N; Kou, M; Hu, X; Chen, M; Gao, Y; Honda, Y; Dowdy, D; Mok, Y; Ishigami, J; Appel, The relationship of COVID-19 severity with cardiovascular disease and its traditional risk factors: A systematic review and meta-analysis. BMJ medRxiv Prepr. (2020), doi:https://doi.org/10.1101/2020.04.05.20054155.

27. S.-Y. Lau, P. Wang, B. W.-Y. Mok, A. J. Zhang, H. Chu, A. C.-Y. Lee, S. Deng, P. Chen, K.-H. Chan, W. Song, Z. Chen, K. K.-W. To, J. F.-W. Chan, K.-Y. Yuen, H. Chen, Attenuated SARS-CoV-2 variants with deletions at the S1/S2 junction. Emerg. Microbes Infect. 9, 837–842 (2020).

28. S. Xia, M. Liu, C. Wang, W. Xu, Q. Lan, S. Feng, F. Qi, L. Bao, L. Du, S. Liu, C. Qin, F. Sun, Z. Shi, Y. Zhu, S. Jiang, L. Lu, Inhibition of SARS-CoV-2 (previously 2019-nCoV) infection by a highly potent pan-coronavirus fusion inhibitor targeting its spike protein that harbors a high capacity to mediate membrane fusion. Cell Res. 30, 343–355 (2020).

