## Supplementary material for "Naturally occurring SARS-CoV-2 gene deletions close to the spike S1/S2 cleavage site in the viral quasispecies of COVID19 patients"

**SUPPLEMENTARY TABLES S1-S13**

**Table S1.** Amplicons positions related to reference GenBank sequence from Wuhan-Hu-1, reference MN908947.1

| Amplicon ID | Primer FW position | Primer RV position | 5' trim position | 3' trim position | Bp Length | Bp Overlap | Gene S 5' trim pos. | In frame 5' trim | Aa pos. |
|---|---|---|---|---|---|---|---|---|---|
| **N01--016** | 21517 | 21964 | 21546 | 21936 | 391 | --------- | -16 | 17 | 1 |
| **N02-0370** | 21907 | 22251 | 21932 | 22232 | 301 | 5 | 370 | 0 | 124 |
| **N03-0634** | 22170 | 22635 | 22196 | 22613 | 418 | 37 | 634 | 0 | 212 |
| **N04-0867** | 22407 | 22888 | 22429 | 22861 | 433 | 185 | 867 | 1 | 290 |
| **N05-1230** | 22768 | 23138 | 22792 | 23118 | 327 | 70 | 1230 | 1 | 411 |
| **N06-1536** | 23069 | 23426 | 23098 | 23406 | 309 | 21 | 1536 | 1 | 513 |
| **N07-1810** | 23347 | 23713 | 23372 | 23688 | 317 | 35 | 1810 | 0 | 604 |
| **N08-2103** | 23636 | 24074 | 23665 | 24051 | 387 | 24 | 2103 | 0 | 701 |
| **N09-2447** | 23987 | 24299 | 24009 | 24272 | 264 | 43 | 2447 | 2 | 817 |
| **N10-2610** | 24147 | 24616 | 24172 | 24590 | 419 | 101 | 2610 | 1 | 871 |
| **N11-2968** | 24507 | 24882 | 24530 | 24859 | 330 | 61 | 2968 | 0 | 990 |
| **N12-3250** | 24792 | 25166 | 24812 | 25142 | 331 | 48 | 3250 | 0 | 1084 |
| **N13-3418** | 24954 | 25418 | 24980 | 25392 | 413 | 163 | 3418 | 0 | 1140 |

**Table S2.** Quality scores of each MiSeq run named 55 to 58.

| Run | Amplicons | % PhiX | Cluster Density | % > Q30 | | | Pass Filter Reads | % Assigned |
|---|---|---|---|---|---|---|---|---|
| | | | | R1 | R3 | Total | | |
| **55** | 78 | 3,84 | 866 | 96,24 | 88,20 | 92,23 | 20.813.664 | 93,59 |
| **56** | 44 | 10,80 | 1276 | 94,89 | 90,75 | 90,75 | 29.050.702 | 90,99 |
| **57** | 62 | 15,02 | 936 | 95,64 | 86,41 | 90,98 | 22.170.213 | 88,48 |
| **58** | 66 | 14,03 | 1092 | 96,15 | 90,53 | 93,28 | 25.557.754 | 89,25 |
| **Total** | **250** | | | | | | **97.592.333** | |

**Table S3.** Experiment design. Runs assigned to each sample/amplicon

| | N01 | N02 | N03 | N04 | N05 | N06 | N07 | N08 | N09 | N10 | N11 | N12 | N13 | Clinics | Ct |
|---|---|---|---|---|---|---|---|---|---|---|---|---|---|---|---|
| **P01** | 55 | 55 | 55 | 57 | 55 | 55 | 55 | 58 | 55 | 55 | 55 | 55 | 57 | Mild | 19,00 |
| **P02** | 55 | 55 | 55 | 55 | 55 | 55 | 55 | 58 | 55 | 55 | 55 | 55 | 57 | Mild | 25,00 |
| **P03** | 55 | 55 | 55 | 55 | 55 | 55 | 55 | 58 | 55 | 55 | 55 | 55 | 57 | Mild | 16,40 |
| **P04** | 55 | 55 | 55 | 55 | 55 | 55 | 55 | 58 | 55 | 55 | 55 | 55 | 57 | Mild | 23,10 |
| **P05** | 55 | 55 | 55 | 55 | 55 | 55 | 55 | 58 | 55 | 55 | 55 | 55 | 57 | Mild | 25,98 |
| **P06** | 56 | 56 | 56 | 55 | 56 | 56 | 56 | 58 | 56 | 56 | 56 | 56 | 57 | Mild | 21,45 |
| **P07** | 57 | 57 | 57 | 57 | 57 | 57 | 57 | 58 | 57 | 57 | 57 | 57 | 57 | Mild | 25,94 |
| **P08** | 58 | 58 | 58 | 58 | 58 | 58 | 58 | 58 | 58 | 58 | 58 | 58 | 58 | Severe | NA |
| **P09** | 57 | 57 | 57 | 57 | 57 | 57 | 57 | 58 | 57 | 57 | 57 | 57 | 57 | Severe | 25,36 |
| **P10** | 56 | 56 | 56 | 57 | 56 | 56 | 56 | 58 | 56 | 56 | 56 | 56 | 57 | Severe | 21,23 |
| **P11** | 56 | 56 | 56 | 57 | 56 | 56 | 56 | 58 | 56 | 56 | 56 | 56 | 57 | Severe | 36,01 |
| **P12** | 58 | 58 | 58 | 58 | 58 | 58 | 58 | 58 | 58 | 58 | 58 | 58 | 58 | Severe | 31,04 |
| **P13** | 56 | 56 | 56 | 57 | 56 | 56 | 56 | 58 | 56 | 56 | 56 | 56 | 57 | Severe | 22,94 |
| **P14** | 57 | 57 | 57 | 57 | 57 | 57 | 57 | 58 | 57 | 57 | 57 | 57 | 57 | Mild | 23,71 |
| **P15** | 57 | 57 | 57 | 57 | 57 | 57 | 57 | 58 | 57 | 57 | 57 | 57 | 57 | Mild | 27,32 |
| **P16** | 58 | 58 | 58 | 58 | 58 | 58 | 58 | 58 | 58 | 58 | 58 | 58 | 58 | Severe | 31,35 |
| **P17** | 58 | 58 | 58 | 58 | 58 | 58 | 58 | 58 | 58 | 58 | 58 | 58 | 58 | Severe | 30,77 |
| **P18** | 55 | 55 | 55 | 57 | 55 | 55 | 55 | 58 | 55 | 55 | 55 | 55 | 57 | Mild | 15,50 |

**Table S4.** Final coverage, globally and by amplicons

| Coverage distribution, globally by amplicon | | | | | | |
|---|---|---|---|---|---|---|
| | **Min.** | **1st Qu.** | **Median** | **Mean** | **3rd Qu.** | **Max.** |
| **Globally** | 81202 | 139568 | 171478 | 208319 | 239172 | 597558 |
| **Coverage distribution, by amplicon** | | | | | | |
| **Amplicon** | **Min.** | **1st Qu.** | **Median** | **Mean** | **3rd Qu.** | **Max.** |
| **N01--016** | 108858 | 119879.2 | 139770.0 | 153125.4 | 179658.5 | 252666 |
| **N02-0370** | 163033 | 216452.8 | 270480.0 | 306943.6 | 342410.0 | 571306 |
| **N03-0634** | 111481 | 123999.2 | 140982.5 | 156211.4 | 167189.2 | 260079 |
| **N04-0867** | 133191 | 155911.8 | 161195.0 | 163940.7 | 168463.2 | 218955 |
| **N05-1230** | 159432 | 188796.5 | 221486.5 | 251823.3 | 312959.0 | 413965 |
| **N06-1536** | 126525 | 150571.2 | 182907.0 | 208838.9 | 230352.0 | 417160 |
| **N07-1810** | 81202 | 143585.2 | 179502.0 | 212005.9 | 247503.8 | 417674 |
| **N08-2101** | 109696 | 138632.2 | 145879.5 | 147679.3 | 156752.8 | 188798 |
| **N09-2447** | 88196 | 113809.2 | 165799.5 | 225772.8 | 281115.0 | 498361 |
| **N10-2610** | 97760 | 125432.2 | 176356.0 | 188779.4 | 232313.5 | 318908 |
| **N11-2968** | 128489 | 156074.2 | 200439.5 | 239287.4 | 267656.2 | 451180 |
| **N12-3250** | 170246 | 208136.0 | 259068.5 | 316778.6 | 345288.8 | 597558 |
| **N13-3418** | 93123 | 121680.2 | 132628.5 | 136960.3 | 149987.2 | 193516 |

**Table S5.** Total number of reads obtained by amplicon and per patient for InDel Bioinformatics analysis (see methods).

| | N01 | N02 | N03 | N04 | N05 | N06 | N07 | N08 | N09 | N10 | N11 | N12 | N13 | Total |
|---|---|---|---|---|---|---|---|---|---|---|---|---|---|---|
| **P01** | 126.140 | 214.410 | 123.583 | 155.414 | 172.903 | 173.278 | 162.753 | 188.798 | 130.031 | 130.134 | 169.759 | 207.326 | 123.284 | 2.077.813 |
| **P02** | 120.693 | 209.682 | 111.481 | 147.559 | 187.538 | 150.059 | 119.226 | 147.181 | 131.564 | 123.865 | 152.125 | 210.566 | 112.682 | 1.924.221 |
| **P03** | 132.797 | 280.219 | 122.298 | 172.784 | 192.572 | 137.806 | 88.629 | 167.853 | 101.547 | 97.760 | 128.489 | 186.463 | 97.426 | 1.906.643 |
| **P04** | 108.858 | 194.547 | 115.885 | 157.405 | 170.929 | 149.787 | 148.533 | 172.330 | 111.096 | 120.486 | 146.648 | 205.361 | 122.269 | 1.924.134 |
| **P05** | 119.151 | 200.164 | 125.248 | 164.460 | 159.432 | 140.610 | 141.936 | 123.580 | 96.938 | 106.203 | 135.932 | 185.940 | 117.774 | 1.817.368 |
| **P06** | 226.446 | 549.232 | 239.167 | 147.446 | 316.716 | 299.024 | 377.883 | 151.756 | 446.100 | 318.908 | 446.727 | 564.720 | 134.826 | 4.218.951 |
| **P07** | 187.731 | 238.043 | 150.720 | 187.152 | 242.302 | 172.026 | 150.994 | 137.992 | 121.949 | 167.580 | 186.481 | 233.956 | 132.828 | 2.309.754 |
| **P08** | 119.608 | 333.287 | 141.006 | 158.030 | 413.965 | 213.511 | 236.058 | 143.523 | 267.795 | 178.286 | 256.917 | 306.387 | 166.308 | 2.934.681 |
| **P09** | 155.441 | 249.554 | 147.531 | 167.822 | 213.635 | 192.536 | 192.954 | 131.492 | 172.699 | 174.426 | 214.398 | 270.683 | 146.733 | 2.429.904 |
| **P10** | 145.296 | 397.733 | 172.679 | 133.191 | 229.338 | 238.762 | 320.827 | 153.266 | 423.469 | 296.991 | 382.021 | 567.476 | 132.429 | 3.593.478 |
| **P11** | 227.142 | 511.453 | 236.148 | 163.172 | 357.145 | 417.160 | 382.413 | 163.666 | 424.874 | 261.499 | 400.440 | 551.456 | 131.133 | 4.227.701 |
| **P12** | 109.992 | 345.451 | 139.064 | 186.064 | 335.130 | 230.354 | 251.319 | 140.904 | 285.555 | 203.133 | 271.236 | 355.196 | 190.092 | 3.043.490 |
| **P13** | 252.666 | 571.306 | 260.079 | 159.218 | 373.205 | 338.914 | 417.674 | 140.553 | 498.361 | 315.226 | 451.180 | 597.558 | 133.612 | 4.509.552 |
| **P14** | 139.365 | 222.581 | 122.497 | 138.553 | 204.863 | 167.440 | 166.050 | 149.703 | 158.900 | 154.192 | 179.171 | 247.454 | 121.484 | 2.172.253 |
| **P15** | 192.348 | 163.033 | 184.162 | 218.955 | 168.990 | 126.525 | 133.830 | 144.578 | 88.196 | 229.561 | 167.922 | 215.279 | 193.516 | 2.226.895 |
| **P16** | 111.786 | 293.193 | 148.252 | 167.114 | 290.408 | 228.854 | 223.566 | 133.441 | 267.793 | 233.231 | 249.013 | 310.380 | 164.694 | 2.821.725 |
| **P17** | 140.623 | 290.356 | 140.959 | 168.677 | 301.688 | 230.346 | 220.260 | 157.915 | 243.100 | 185.798 | 239.174 | 315.567 | 151.072 | 2.785.535 |
| **P18** | 140.175 | 260.741 | 131.046 | 157.917 | 202.060 | 152.108 | 81.202 | 109.696 | 93.943 | 100.751 | 129.541 | 170.246 | 93.123 | 1.822.549 |
| **Total** | 2.756.258 | 5.524.985 | 2.811.805 | 2.950.933 | 4.532.819 | 3.759.100 | 3.816.107 | 2.658.227 | 4.063.910 | 3.398.030 | 4.307.174 | 5.702.014 | 2.465.285 | 48.746.647 |

**Table S6.** Percent frequency of master haplotype by amplicon and sample

| | N01 | N02 | N03 | N04 | N05 | N06 | N07 | N08 | N09 | N10 | N11 | N12 | N13 |
|---|---|---|---|---|---|---|---|---|---|---|---|---|---|
| **P01** | 98.9 | 97.7 | 98.5 | 97.5 | 97.5 | 98.8 | 95.4 | 97.1 | 98.5 | 97.8 | 98.2 | 100.0 | 97.8 |
| **P02** | 99.0 | 99.2 | 98.7 | 98.6 | 99.2 | 98.6 | 98.5 | 99.2 | 99.8 | 99.7 | 99.9 | 100.0 | 99.7 |
| **P03** | 96.3 | 95.6 | 95.7 | 94.0 | 96.2 | 93.8 | 95.3 | 99.2 | 95.5 | 93.7 | 92.9 | 93.1 | 93.4 |
| **P04** | 98.4 | 98.2 | 97.9 | 97.7 | 98.4 | 98.3 | 97.3 | 99.2 | 99.6 | 99.2 | 100.0 | 99.5 | 99.0 |
| **P05** | 99.8 | 99.2 | 99.8 | 99.8 | 99.4 | 99.4 | 98.0 | 95.1 | 99.6 | 99.9 | 100.0 | 99.8 | 99.9 |
| **P06** | 100.0 | 99.8 | 99.9 | 99.5 | 99.6 | 99.8 | 98.0 | 98.8 | 99.7 | 99.8 | 99.9 | 99.9 | 99.9 |
| **P07** | 99.1 | 98.7 | 99.1 | 97.9 | 98.4 | 97.2 | 95.4 | 94.7 | 95.6 | 92.9 | 91.8 | 93.3 | 92.2 |
| **P08** | 91.6 | 94.7 | 93.8 | 92.2 | 93.3 | 85.0 | 79.3 | 89.5 | 91.2 | 71.9 | 90.3 | 86.0 | 83.8 |
| **P09** | 96.4 | 97.5 | 96.6 | 96.9 | 96.1 | 97.2 | 92.7 | 97.0 | 97.8 | 98.2 | 98.3 | 98.1 | 97.3 |
| **P10** | 99.4 | 99.2 | 99.2 | 98.6 | 98.9 | 98.8 | 98.9 | 98.3 | 99.3 | 98.9 | 99.7 | 98.9 | 98.5 |
| **P11** | 98.5 | 99.0 | 98.6 | 98.3 | 98.9 | 97.3 | 82.7 | 96.2 | 99.7 | 99.6 | 98.2 | 88.3 | 97.1 |
| **P12** | 98.8 | 98.9 | 99.7 | 98.4 | 98.9 | 97.3 | 98.3 | 96.3 | 96.3 | 99.4 | 100.0 | 99.3 | 98.6 |
| **P13** | 99.3 | 99.7 | 99.0 | 99.3 | 99.4 | 99.2 | 99.7 | 98.4 | 100.0 | 99.8 | 99.9 | 99.7 | 99.9 |
| **P14** | 94.1 | 97.9 | 91.4 | 90.9 | 96.8 | 91.1 | 92.5 | 95.2 | 95.9 | 91.4 | 96.4 | 96.3 | 96.4 |
| **P15** | 96.6 | 97.8 | 97.9 | 96.4 | 92.0 | 93.9 | 93.2 | 92.8 | 94.8 | 98.5 | 98.6 | 99.2 | 99.2 |
| **P16** | 99.6 | 97.6 | 99.4 | 99.1 | 98.7 | 96.8 | 96.4 | 65.1 | 99.6 | 99.8 | 99.1 | 99.4 | 98.2 |
| **P17** | 86.5 | 81.9 | 86.6 | 88.6 | 87.8 | 84.7 | 84.6 | 88.3 | 90.9 | 86.5 | 84.6 | 80.9 | 73.1 |
| **P18** | 99.2 | 97.6 | 97.6 | 97.4 | 97.7 | 97.2 | 98.1 | 97.4 | 98.0 | 97.4 | 96.7 | 95.8 | 95.7 |

**Table S7.** Number of haplotypes by amplicon and sample

| | N01 | N02 | N03 | N04 | N05 | N06 | N07 | N08 | N09 | N10 | N11 | N12 | N13 |
|---|---|---|---|---|---|---|---|---|---|---|---|---|---|
| **P01** | 5 | 7 | 6 | 11 | 10 | 5 | 15 | 13 | 8 | 10 | 8 | 1 | 9 |
| **P02** | 7 | 6 | 9 | 10 | 6 | 10 | 8 | 5 | 2 | 3 | 2 | 1 | 3 |
| **P03** | 24 | 26 | 26 | 37 | 23 | 38 | 28 | 5 | 27 | 34 | 42 | 40 | 35 |
| **P04** | 6 | 12 | 10 | 12 | 7 | 11 | 15 | 6 | 4 | 6 | 1 | 3 | 6 |
| **P05** | 2 | 6 | 2 | 2 | 4 | 6 | 9 | 28 | 3 | 2 | 1 | 2 | 2 |
| **P06** | 1 | 3 | 2 | 5 | 3 | 3 | 9 | 7 | 2 | 3 | 2 | 2 | 2 |
| **P07** | 6 | 10 | 7 | 14 | 11 | 19 | 28 | 31 | 26 | 39 | 50 | 39 | 39 |
| **P08** | 21 | 31 | 31 | 35 | 37 | 41 | 31 | 21 | 27 | 29 | 51 | 38 | 42 |
| **P09** | 10 | 10 | 9 | 10 | 11 | 16 | 20 | 11 | 7 | 7 | 6 | 7 | 12 |
| **P10** | 5 | 6 | 6 | 9 | 8 | 8 | 9 | 10 | 6 | 8 | 3 | 8 | 10 |
| **P11** | 10 | 5 | 6 | 7 | 8 | 16 | 11 | 11 | 3 | 3 | 5 | 3 | 8 |
| **P12** | 8 | 8 | 3 | 11 | 8 | 19 | 8 | 8 | 7 | 5 | 1 | 6 | 7 |
| **P13** | 2 | 3 | 5 | 5 | 5 | 7 | 3 | 9 | 1 | 2 | 2 | 3 | 2 |
| **P14** | 16 | 12 | 12 | 18 | 14 | 24 | 26 | 16 | 13 | 25 | 16 | 17 | 18 |
| **P15** | 9 | 8 | 7 | 8 | 10 | 16 | 10 | 14 | 8 | 5 | 5 | 4 | 5 |
| **P16** | 4 | 10 | 4 | 6 | 9 | 15 | 6 | 6 | 4 | 2 | 5 | 2 | 6 |
| **P17** | 45 | 44 | 44 | 56 | 51 | 55 | 37 | 40 | 45 | 59 | 61 | 57 | 43 |
| **P18** | 6 | 16 | 16 | 16 | 16 | 19 | 10 | 15 | 14 | 15 | 23 | 25 | 24 |

**Table S8.** Gaps incidence by amplicon

| Amplicon | Npats | Npos | Nhpl | Nreads | Pctg |
|---|---|---|---|---|---|
| **N01--016** | 5 | 20 | 11 | 7563 | 0,3009 |
| **N02-0370** | 8 | 30 | 25 | 15143 | 0,3616 |
| **N03-0634** | 7 | 20 | 13 | 7111 | 0,2747 |
| **N04-0867** | 7 | 34 | 13 | 6296 | 0,2219 |
| **N05-1230** | 7 | 52 | 10 | 5888 | 0,1574 |
| **N06-1536** | 3 | 12 | 4 | 1410 | 0,0456 |
| **N07-1810** | 15 | 89 | 49 | 83485 | 1,9950 |
| **N08-2101** | 18 | 20 | 39 | 66057 | 2,6559 |
| **N09-2447** | 9 | 26 | 10 | 4817 | 0,1841 |
| **N10-2610** | 3 | 4 | 3 | 1444 | 0,0536 |
| **N11-2968** | 2 | 3 | 2 | 1400 | 0,0351 |
| **N12-3250** | 1 | 1 | 1 | 15201 | 0,2756 |
| **N13-3418** | 2 | 2 | 2 | 8745 | 0,2947 |

**Table S9.** Gaps incidence by sample and amplicon

| Pat.ID | Ampl.ID | Hpl | Reads | Deletions Reads | Deletions % | Pat.ID | Ampl.ID | Hpl | Reads | Deletions Reads | Deletions % |
|---|---|---|---|---|---|---|---|---|---|---|---|
| **P01** | N01--016 | 5 | 126140 | 837 | 0,66 | **P07** | N10-2610 | 39 | 167580 | 254 | 0,15 |
| **P01** | N02-0370 | 7 | 214410 | 3084 | 1,44 | **P08** | N03-0634 | 31 | 141006 | 833 | 0,59 |
| **P01** | N04-0867 | 11 | 155414 | 338 | 0,22 | **P08** | N04-0867 | 35 | 158030 | 871 | 0,55 |
| **P01** | N05-1230 | 10 | 172903 | 245 | 0,14 | **P08** | N07-1810 | 31 | 236058 | 23541 | 9,97 |
| **P01** | N06-1536 | 5 | 173278 | 521 | 0,30 | **P08** | N08-2101 | 21 | 143523 | 298 | 0,21 |
| **P01** | N07-1810 | 15 | 162753 | 3663 | 2,25 | **P08** | N12-3250 | 38 | 306387 | 15201 | 4,96 |
| **P01** | N08-2101 | 13 | 188798 | 2481 | 1,31 | **P08** | N13-3418 | 42 | 166308 | 8158 | 4,91 |
| **P01** | N09-2447 | 8 | 130031 | 536 | 0,41 | **P09** | N01--016 | 10 | 155441 | 3088 | 1,99 |
| **P01** | N10-2610 | 10 | 130134 | 671 | 0,52 | **P09** | N02-0370 | 10 | 249554 | 3954 | 1,58 |
| **P01** | N11-2968 | 8 | 169759 | 873 | 0,51 | **P09** | N03-0634 | 9 | 147531 | 1864 | 1,26 |
| **P02** | N02-0370 | 6 | 209682 | 642 | 0,31 | **P09** | N04-0867 | 10 | 167822 | 1619 | 0,96 |
| **P02** | N03-0634 | 9 | 111481 | 345 | 0,31 | **P09** | N05-1230 | 11 | 213635 | 1672 | 0,78 |
| **P02** | N04-0867 | 10 | 147559 | 227 | 0,15 | **P09** | N07-1810 | 20 | 192954 | 4392 | 2,28 |
| **P02** | N05-1230 | 6 | 187538 | 262 | 0,14 | **P09** | N08-2101 | 11 | 131492 | 1360 | 1,03 |
| **P02** | N06-1536 | 10 | 150059 | 581 | 0,39 | **P09** | N09-2447 | 7 | 172699 | 737 | 0,43 |
| **P02** | N07-1810 | 8 | 119226 | 876 | 0,73 | **P09** | N10-2610 | 7 | 174426 | 519 | 0,30 |
| **P02** | N08-2101 | 5 | 147181 | 516 | 0,35 | **P09** | N13-3418 | 12 | 146733 | 587 | 0,40 |
| **P03** | N07-1810 | 28 | 88629 | 223 | 0,25 | **P10** | N07-1810 | 9 | 320827 | 408 | 0,13 |
| **P03** | N08-2101 | 5 | 167853 | 869 | 0,52 | **P10** | N08-2101 | 10 | 153266 | 442 | 0,29 |
| **P03** | N09-2447 | 27 | 101547 | 165 | 0,16 | **P11** | N07-1810 | 11 | 382413 | 20109 | 5,26 |
| **P04** | N01--016 | 6 | 108858 | 665 | 0,61 | **P11** | N08-2101 | 11 | 163666 | 932 | 0,57 |
| **P04** | N02-0370 | 12 | 194547 | 2898 | 1,49 | **P12** | N08-2101 | 8 | 140904 | 361 | 0,26 |
| **P04** | N03-0634 | 10 | 115885 | 1537 | 1,33 | **P13** | N08-2101 | 9 | 140553 | 665 | 0,47 |
| **P04** | N04-0867 | 12 | 157405 | 2564 | 1,63 | **P14** | N01--016 | 16 | 139365 | 2789 | 2,00 |
| **P04** | N05-1230 | 7 | 170929 | 1776 | 1,04 | **P14** | N02-0370 | 12 | 222581 | 1145 | 0,51 |
| **P04** | N07-1810 | 15 | 148533 | 2003 | 1,35 | **P14** | N03-0634 | 12 | 122497 | 441 | 0,36 |
| **P04** | N08-2101 | 6 | 172330 | 991 | 0,58 | **P14** | N04-0867 | 18 | 138553 | 448 | 0,32 |
| **P04** | N09-2447 | 4 | 111096 | 137 | 0,12 | **P14** | N07-1810 | 26 | 166050 | 730 | 0,44 |
| **P05** | N01--016 | 2 | 119151 | 184 | 0,15 | **P14** | N08-2101 | 16 | 149703 | 2969 | 1,98 |
| **P05** | N02-0370 | 6 | 200164 | 1382 | 0,69 | **P14** | N09-2447 | 13 | 158900 | 500 | 0,31 |
| **P05** | N05-1230 | 4 | 159432 | 340 | 0,21 | **P15** | N03-0634 | 7 | 184162 | 1767 | 0,96 |
| **P05** | N07-1810 | 9 | 141936 | 2031 | 1,43 | **P15** | N07-1810 | 10 | 133830 | 3016 | 2,25 |
| **P05** | N08-2101 | 28 | 123580 | 506 | 0,41 | **P15** | N08-2101 | 14 | 144578 | 5407 | 3,74 |
| **P05** | N09-2447 | 3 | 96938 | 264 | 0,27 | **P15** | N09-2447 | 8 | 88196 | 1041 | 1,18 |
| **P06** | N02-0370 | 3 | 549232 | 1329 | 0,24 | **P16** | N08-2101 | 6 | 133441 | 45743 | 34,28 |
| **P06** | N03-0634 | 2 | 239167 | 324 | 0,14 | **P17** | N02-0370 | 44 | 290356 | 709 | 0,24 |
| **P06** | N04-0867 | 5 | 147446 | 229 | 0,16 | **P17** | N05-1230 | 51 | 301688 | 848 | 0,28 |
| **P06** | N05-1230 | 3 | 316716 | 745 | 0,24 | **P17** | N06-1536 | 55 | 230346 | 308 | 0,13 |
| **P06** | N07-1810 | 9 | 377883 | 5893 | 1,56 | **P17** | N07-1810 | 37 | 220260 | 15796 | 7,17 |
| **P06** | N08-2101 | 7 | 151756 | 1197 | 0,79 | **P17** | N08-2101 | 40 | 157915 | 323 | 0,20 |
| **P06** | N09-2447 | 2 | 446100 | 1318 | 0,30 | **P18** | N07-1810 | 10 | 81202 | 531 | 0,65 |
| **P06** | N11-2968 | 2 | 446727 | 527 | 0,12 | **P18** | N08-2101 | 15 | 109696 | 491 | 0,45 |
| **P07** | N08-2101 | 31 | 137992 | 506 | 0,37 | **P18** | N09-2447 | 14 | 93943 | 119 | 0,13 |

**Table S10.** Premature STOPs incidence

| ID | Ampl | Pos | Freq % | ID | Ampl | Pos | Freq % | ID | Ampl | Pos | Freq % |
|---|---|---|---|---|---|---|---|---|---|---|---|
| **P01** | A01-0001 | 109 | 0,48 | **P04** | A07-0604 | 672 | 0,21 | **P09** | A08-0701 | 825 | 0,43 |
| **P01** | A02-0124 | 212 | 0,65 | **P04** | A07-0604 | 648 | 0,13 | **P09** | A08-0701 | 821 | 0,30 |
| **P01** | A02-0124 | 155 | 0,78 | **P04** | A08-0701 | 826 | 0,13 | **P09** | A08-0701 | 731 | 0,30 |
| **P01** | A04-0290 | 400 | 0,22 | **P04** | A08-0701 | 731 | 0,28 | **P09** | A09-0817 | 825 | 0,43 |
| **P01** | A07-0604 | 684 | 0,18 | **P04** | A09-0817 | 838 | 0,12 | **P09** | A10-0871 | 996 | 0,30 |
| **P01** | A07-0604 | 678 | 0,56 | **P05** | A01-0001 | 110 | 0,15 | **P10** | A07-0604 | 678 | 0,13 |
| **P01** | A07-0604 | 682 | 0,33 | **P05** | A02-0124 | 155 | 0,23 | **P10** | A08-0701 | 731 | 0,29 |
| **P01** | A07-0604 | 660 | 0,29 | **P05** | A02-0124 | 144 | 0,13 | **P11** | A06-0513 | 580 | 0,17 |
| **P01** | A07-0604 | 683 | 0,28 | **P05** | A05-0411 | 434 | 0,21 | **P11** | A07-0604 | 674 | 5,07 |
| **P01** | A07-0604 | 672 | 0,24 | **P05** | A07-0604 | 680 | 0,39 | **P11** | A07-0604 | 674 | 0,18 |
| **P01** | A08-0701 | 821 | 0,23 | **P05** | A07-0604 | 678 | 0,61 | **P11** | A08-0701 | 731 | 0,57 |
| **P01** | A08-0701 | 826 | 0,23 | **P05** | A07-0604 | 669 | 0,27 | **P12** | A08-0701 | 731 | 0,26 |
| **P01** | A08-0701 | 731 | 0,31 | **P05** | A07-0604 | 672 | 0,16 | **P13** | A08-0701 | 731 | 0,47 |
| **P01** | A09-0817 | 867 | 0,20 | **P05** | A08-0701 | 731 | 0,27 | **P14** | A01-0001 | 110 | 0,29 |
| **P01** | A09-0817 | 835 | 0,22 | **P06** | A02-0124 | 211 | 0,12 | **P14** | A02-0124 | 212 | 0,51 |
| **P01** | A11-0990 | 1013 | 0,51 | **P06** | A02-0124 | 155 | 0,12 | **P14** | A03-0212 | 271 | 0,19 |
| **P02** | A02-0124 | 203 | 0,18 | **P06** | A05-0411 | 434 | 0,24 | **P14** | A04-0290 | 387 | 0,32 |
| **P02** | A02-0124 | 151 | 0,13 | **P06** | A07-0604 | 680 | 0,26 | **P14** | A07-0604 | 644 | 0,69 |
| **P02** | A03-0212 | 244 | 0,14 | **P06** | A07-0604 | 678 | 0,68 | **P14** | A07-0604 | 613 | 0,48 |
| **P02** | A04-0290 | 363 | 0,15 | **P06** | A07-0604 | 682 | 0,12 | **P14** | A07-0604 | 677 | 0,46 |
| **P02** | A05-0411 | 434 | 0,14 | **P06** | A07-0604 | 669 | 0,34 | **P14** | A08-0701 | 821 | 0,39 |
| **P02** | A06-0513 | 598 | 0,23 | **P06** | A08-0701 | 731 | 0,32 | **P14** | A08-0701 | 826 | 1,14 |
| **P02** | A06-0513 | 546 | 0,15 | **P06** | A11-0990 | 1060 | 0,12 | **P14** | A08-0701 | 731 | 0,31 |
| **P02** | A07-0604 | 680 | 0,21 | **P07** | A07-0604 | 678 | 0,18 | **P14** | A09-0817 | 853 | 0,31 |
| **P02** | A07-0604 | 678 | 0,34 | **P07** | A08-0701 | 826 | 0,14 | **P14** | A09-0817 | 867 | 0,31 |
| **P02** | A07-0604 | 669 | 0,18 | **P07** | A08-0701 | 731 | 0,23 | **P14** | A10-0871 | 935 | 0,78 |
| **P02** | A08-0701 | 731 | 0,35 | **P07** | A10-0871 | 922 | 0,15 | **P14** | A10-0871 | 992 | 0,48 |
| **P03** | A07-0604 | 678 | 0,25 | **P08** | A04-0290 | 387 | 0,55 | **P14** | A10-0871 | 949 | 0,29 |
| **P03** | A08-0701 | 731 | 0,26 | **P08** | A07-0604 | 669 | 6,64 | **P14** | A11-0990 | 992 | 0,45 |
| **P04** | A01-0001 | 110 | 0,15 | **P08** | A07-0604 | 674 | 3,18 | **P15** | A01-0001 | 13 | 0,78 |
| **P04** | A02-0124 | 212 | 0,14 | **P08** | A07-0604 | 669 | 0,15 | **P15** | A03-0212 | 249 | 0,96 |
| **P04** | A02-0124 | 212 | 0,24 | **P08** | A08-0701 | 731 | 0,21 | **P15** | A07-0604 | 676 | 1,18 |
| **P04** | A02-0124 | 211 | 0,14 | **P08** | A12-1084 | 1177 | 4,96 | **P15** | A08-0701 | 767 | 2,20 |
| **P04** | A02-0124 | 209 | 0,14 | **P08** | A13-1140 | 1177 | 4,91 | **P15** | A08-0701 | 731 | 0,28 |
| **P04** | A02-0124 | 151 | 0,17 | **P09** | A01-0001 | 110 | 1,10 | **P16** | A02-0124 | 183 | 0,95 |
| **P04** | A02-0124 | 152 | 0,23 | **P09** | A02-0124 | 210 | 0,13 | **P16** | A08-0701 | 767 | 34,13 |
| **P04** | A02-0124 | 155 | 0,13 | **P09** | A02-0124 | 154 | 0,23 | **P16** | A08-0701 | 731 | 0,14 |
| **P04** | A04-0290 | 386 | 0,21 | **P09** | A02-0124 | 155 | 0,59 | **P17** | A01-0001 | 23 | 0,21 |
| **P04** | A04-0290 | 387 | 0,30 | **P09** | A03-0212 | 248 | 0,38 | **P17** | A02-0124 | 212 | 0,24 |
| **P04** | A04-0290 | 342 | 0,20 | **P09** | A03-0212 | 244 | 0,88 | **P17** | A05-0411 | 510 | 0,28 |
| **P04** | A04-0290 | 363 | 0,14 | **P09** | A07-0604 | 628 | 0,69 | **P17** | A06-0513 | 546 | 0,13 |
| **P04** | A05-0411 | 433 | 0,15 | **P09** | A07-0604 | 680 | 0,36 | **P17** | A08-0701 | 731 | 0,20 |
| **P04** | A05-0411 | 502 | 0,18 | **P09** | A07-0604 | 679 | 0,17 | **P17** | A13-1140 | 1214 | 6,63 |
| **P04** | A05-0411 | 430 | 0,70 | **P09** | A07-0604 | 671 | 0,26 | **P18** | A07-0604 | 680 | 0,23 |
| **P04** | A07-0604 | 680 | 0,22 | **P09** | A07-0604 | 681 | 0,55 | **P18** | A07-0604 | 678 | 0,43 |
| **P04** | A07-0604 | 682 | 0,18 | **P09** | A07-0604 | 672 | 0,18 | **P18** | A08-0701 | 731 | 0,31 |
| **P04** | A07-0604 | 669 | 0,33 | **P09** | A07-0604 | 650 | 0,27 | | | | |

**Table S11A.** List of deletions found in amplicon N07 at the nucleotide (nt) level. wt=wild type; S=stop; Lost+S = lose of reading frame and appearance of a stop codon; rRF = recover Reading frame. Underlined in yellow haplotypes that do not lose TATA-box-like, and in light blue haplotype with deletion upstream of TATA sequence.

| Patient MILD/SEVERE | Nucleotide alignments |
|---|---|
| MN908947.3 | ***5'1974 CTCATATGAGTGTGACATACCCATTGGTGCAGGTATATGCGCTAGTTATCAGACTCAGACTAATTCTCCTCGGCGG 2049 3'*** |
| P01 | *CTCATATGAGTGTGACATACCCATTGGTGCAGG--TATGCGCTAGTTATCAGACTCAGACTAATTCTCCTCGGCGG* |
| P09 | *CTCATATGAGTGTGACATACCCATTGGTGCAGG--TATGCGCTAGTTATCAGACTCAGACTAATTCTCCTCGGCGG* |
| P05 | *CTCATATGAGTGTGACATACCCATTGGTGCAGG--TATGCGCTAGTTATCAGACTCAGACTAATTCTCCTCGGCGG* |
| P04 | *CTCATATGAGTGTGACATACCCATTGGTGCAGG--TATGCGCTAGTTATCAGACTCAGACTAATTCTCCTCGGCGG* |
| P01 | *CTCATATGAGTGTGACATACCCATT~~~~~~~~~-TATGCGCTAGTTATCAGACTCAGACTAATTCTCCTCGGCGG* |
| P01 | *CTCATATGAGTGTGACATA~~~~~~~~~~~~~~~-TATGCGCTAGTTATCAGACTCAGACTAATTCTCCTCGGCGG* |
| P04 | *CTCATATGAGTGTGACATA~~~~~~~~~~~~~~~-TATGCGCTAGTTATCAGACTCAGACTAATTCTCCTCGGCGG* |
| P06 | *CTCATATGAGTGTGACATA~~~~~~~~~~~~~~~-TATGCGCTAGTTATCAGACTCAGACTAATTCTCCTCGGCGG* |
| P09 | *CTCATATGAGTGTGACATA~~~~~~~~~~~~~~~~~~~~~~~~-GTTATCAGACTCAGACTAATTCTCCTCGGCGG* |
| P02 | *CTCATATGAGTGTGACATACCCA--~~~~~~~~~~~~~~~~~~--TTATCAGACTCAGACTAATTCTCCTCGGCGG* |
| P06 | CTCATATGAGTGTGACATACCCATTGGTGCAGG-~~~~~~~~~--TAATCAGACTCAGACTAATTCTCCTCGGCGG |
| P06 | CTCATATGAGTGTGACATACCCA--~~~~~~~~~~~~~~~~~~--TTATCAGACTCAGACTAATTCTCCTCGGCGG |
| P08 | CTCATATGAGTGTGACA--~~~~~~~~~~~~~~~~~~~~~~~~~~~TATCAGACTCAGACTAATTCTCCTCGGCGG |
| P08 | CTCATATGAGTGTGACA--~~~~~~~~~~~~~~~~~~~~~~~~~~~TATCAGACTCAGACTAATTCTCCTCGGCGG |
| P08 | CTCATATGAGTGTGACA--~~~~~~~~~~~~~~~~~~~~~~~~~~~TATCAGACTCAGACTAATTCTCCTCGGCGG |
| P01 | CTCATATGAGTGTGACATACCCATTGGTGCAGG-~~~~~~~~~~~~TATCAGACTCAGACTAATTCTCCTCGGCGG |
| P01 | CTCATATGAGTGTGACATACCCATTGGTGCAGGTA--~~~~~~~~~-ATCAGACTCAGACTAATTCTCCTCGGCGG |
| P01 | CTCATATGAGTGTGACATA~~~~~~~~~~~~~~~~~~~~~~~~~~~-ATCAGACTCAGACTAATTCTCCTCGGCGG |
| P02 | CTCATATGAGTGTGACATA~~~~~~~~~~~~~~~~~~~~~~~~~~~-ATCAGACTCAGACTAATTCTCCTCGGCGG |
| P02 | CTCATATGAGTGTGACAT-~~~~~~~~~~~~~~~~~~~~~~~~~~~-ATCAGACTCAGACTAATTCTCCTCGGCGG |
| P03 | CTCATATGAGTGTGACATA~~~~~~~~~~~~~~~~~~~~~~~~~~~-ATCAGACTCAGACTAATTCTCCTCGGCGG |
| P04 | CTCATATGAGTGTGACATACCCATTG--~~~~~~~~~~~~~~~~~~-ATCAGACTCAGACTAATTCTCCTCGGCGG |
| P04 | CTCATATGAGTGTGACATACCCATT~~~~~~~~~~~~~~~~~~~~~-ATCAGACTCAGACTAATTCTCCTCGGCGG |
| P04 | CTCATATGAGTGTGACAT-~~~~~~~~~~~~~~~~~~~~~~~~~~~-ATCAGACTCAGACTAATTCTCCTCGGCGG |
| P05 | CTCATATGAGTGTGACATACCCATT~~~~~~~~~~~~~~~~~~~~~-ATCAGACTCAGACTAATTCTCCTCGGCGG |
| P05 | CTCATATGAGTGTGACATA~~~~~~~~~~~~~~~~~~~~~~~~~~~-ATCAGACTCAGACTAATTCTCCTCGGCGG |
| P05 | CTCATATGAGTGTGACAT-~~~~~~~~~~~~~~~~~~~~~~~~~~~-ATCAGACTCAGACTAATTCTCCTCGGCGG |
| P06 | CTCATATGAGTGTGACATA~~~~~~~~~~~~~~~~~~~~~~~~~~~-ATCAGACTCAGACTAATTCTCCTCGGCGG |
| P07 | CTCATATGAGTGTGACATA~~~~~~~~~~~~~~~~~~~~~~~~~~~-ATCAGACTCAGACTAATTCTCCTCGGCGG |
| P09 | CTCATATGAGTGTGACATACCCATT~~~~~~~~~~~~~~~~~~~~~-ATCAGACTCAGACTAATTCTCCTCGGCGG |
| P10 | CTCATATGAGTGTGACATA~~~~~~~~~~~~~~~~~~~~~~~~~~~-ATCAGACTCAGACTAATTCTCCTCGGCGG |
| P14 | CTCATATGAGTGTGACATACCCATTGGTGCAGGTA--~~~~~~~~~-ATCAGACTCAGACTAATTCTCCTCGGCGG |
| P15 | CTCATATGAGTGTGACATACCCATTGGTGCAGGTA--~~~~~~~~~-ATCAGACTCAGACTAATTCTCCTCGGCGG |
| P18 | CTCATATGAGTGTGACATA~~~~~~~~~~~~~~~~~~~~~~~~~~~-ATCAGACTCAGACTAATTCTCCTCGGCGG |
| P18 | CTCATATGAGTGTGACATACCCATT~~~~~~~~~~~~~~~~~~~~~-ATCAGACTCAGACTAATTCTCCTCGGCGG |
| P06 | CTCATATGAGTGTGACATA~~~~~~~~~~~~~~~~~~~~~~~~~~~--TCAGACTCAGACTAATTCTCCTCGGCGG |
| P09 | CTCATATGAGTGTGACATACCCATT~~~~~~~~~~~~~~~~~~~~~--TCAGACTCAGACTAATTCTCCTCGGCGG |
| P15 | CTCATATGAGTGTG--~~~~~~~~~~~~~~~~~~~~~~~~~~~~~~--TCAGACTCAGACTAATTCTCCTCGGCGG |
| P01 | CTCATA-~~~~~~~~~~~~~~~~~~GGTGCAGGTATATGCGCTAGTTATCAGACTCAGACTAATTCTCCTCGGCGG |
| P08 | CTCATATGAGTGTGACATACCCATTGGTGCAGG**TATA**TGCGCTAGTTA--AGACTCAGACTAATTCTCCTCGGCGG |
| P11 | CTCATATGAGTGTGACATACCCATTGGTGCAGG**TATA**TGCGCTAGTT---AGACTCAGACTAATTCTCCTCGGCGG |
| P11 | CTCATATGAGTGTGACATACCCATTGGTGCAGG**TATA**TGCGCTAGTT---AGACTCAGACTAATTCTCCTCGGCGG |
| P04 | CTCATATGAGTGTGACATACCCATTGGTGCAGG**TATA**TGCGCTAGTT--~~~~~~~~~~~~-ATTCTCCTCGGCGG |

**Table S11B.** List of deletions found in amplicon N07 at the amino acid (aa) level. wt=wild type; S=stop; Lost+S = lose of reading frame and appearance of a stop codon; rRF = recover Reading frame. Underlined in yellow haplotypes that do not lose TATA-box-like, and in light blue haplotype with deletion upstream of TATA sequence.

| Patient MILD/SEVERE | Amino acid alignments | |
|---|---|---|
| **MN908947.3** | **5'654 EHVNNSYECDIPIGAGICASYQTQTNSPRRARSVASQSIIAYTMSLGAENSVAYS 708 3'** | |
| **P01** | EHVNNSYECDIPIGAGMR* | S |
| **P09** | EHVNNSYECDIPIGAGMR* | Lost+S |
| **P05** | EHVNNSYECDIPIGAGMR* | Lost+S |
| **P04** | EHVNNSYECDIPIGAGMR* | Lost+S |
| **P01** | EHVNNSYECDIPIYALVIRLRLILLGGHVV* | Lost+S |
| **P01** | EHVNNSYECDIYALVIRLRLILLGGHVV* | Lost+S |
| **P04** | EHVNNSYECDIYALVIRLRLILLGGHVV* | Lost+S |
| **P06** | EHVNNSYECDIYALVIRLRLILLGGHVV* | Lost+S |
| **P09** | EHVNNSYECDIVIRLRLILLGGHVV* | Lost+S |
| **P02** | EHVNNSYECDIPIIRLRLILLGGHVV* | Lost+S |
| **P06** | EHVNNSYECDIPIGAG---**N**QTQTNSPRRARSVASQSIIAYTMSLGAENSVAYS | rRF |
| **P06** | EHVNNSYECDIPIIRLRLILLGGHVV* | Lost+S |
| **P08** | EHVNNSYECDISDSD* | Lost+S |
| **P08** | EHVNNSYECDISDSD* | Lost+S |
| **P08** | EHVNNSYECDISDSD* | Lost+S |
| **P01** | EHVNNSYECDIPIGAGIRLRLILLGGHVV* | Lost+S |
| **P01** | EHVNNSYECDIPIGAG---------NQTQTNSPRRARSVASQSIIAYTMSLGAENSVAYS | rRF |
| **P01** | EHVNNSYECDIIRLRLILLGGHVV* | Lost+S |
| **P02** | EHVNNSYECDIIRLRLILLGGHVV* | Lost+S |
| **P02** | EHVNNSYECDISDSD* | Lost+S |
| **P03** | EHVNNSYECDIIRLRLILLGGHVV* | Lost+S |
| **P04** | EHVNNSYECDIPI-------DQTQTNSPRRARSVASQSIIAYTMSLGAENSVAYS | rRF |
| **P04** | EHVNNSYECDIPIIRLRLILLGGHVV* | Lost+S |
| **P04** | EHVNNSYECDISDSD* | Lost+S |
| **P05** | EHVNNSYECDIPIIRLRLILLGGHVV* | Lost+S |
| **P05** | EHVNNSYECDIIRLRLILLGGHVV* | Lost+S |
| **P05** | EHVNNSYECDISDSD* | Lost+S |
| **P06** | EHVNNSYECDIIRLRLILLGGHVV* | Lost+S |
| **P07** | EHVNNSYECDIIRLRLILLGGHVV* | Lost+S |
| **P09** | EHVNNSYECDIPIIRLRLILLGGHVV* | Lost+S |
| **P10** | EHVNNSYECDIIRLRLILLGGHVV* | Lost+S |
| **P14** | EHVNNSYECDIPIGAG----**N**QTQTNSPRRARSVASQSIIAYTMSLGAENSVAYS | rRF |
| **P15** | EHVNNSYECDIPIGAG----**N**QTQTNSPRRARSVASQSIIAYTMSLGAENSVAYS | rRF |
| **P18** | EHVNNSYECDIIRLRLILLGGHVV* | Lost+S |
| **P18** | EHVNNSYECDIPIIRLRLILLGGHVV* | Lost+S |
| **P06** | EHVNNSYECDISDSD* | Lost+S |
| **P09** | EHVNNSYECDIPISDSD* | Lost+S |
| **P15** | EHVNNSYECVRLRLILLGGHVV* | Lost+S |
| **P01** | EHVNNS* | S |
| **P08** | EHVNNSYECDIPIGAGICAS* | S |
| **P11** | EHVNNSYECDIPIGAGICAS* | S |
| **P11** | EHVNNSYECDIPIGAGICAS* | S |
| **P04** | EHVNNSYECDIPIGAGICASY----SPRRARSVASQSIIAYTMSLGAENSVAYS | rRF |

**Table S12.** Fisher-test comparing number of deletions in mild versus severe COVID-19 patients

| | With deletions | No deletions |
|---|---|---|
| **Severe** | 4 | 4 |
| **Mild** | 10 | 0 |

Fisher test:
odds-ratio: 95 confidence interval 0.0 - 0.9605
p-value = 0.02288

**Table S13A.** External oligos used at the RT-PCR to amplify the whole spike region (*Reference sequence MN908947.3Wuhan-Hu-1*)

| Fragment | 5'END | Sequence | Product Size |
|---|---|---|---|
| **EXT.1** | 20776 | ATAATGATGAATGTCGCAAAATATAC | 3314 |
| | 24089 | CACCAAGGCAATCACCATAT | |
| **EXT.2** | 22771 | AGGTGATGAAGTCAGACAAAT | 3591 |
| | 26361 | CAATCGAAGCGCAGTAAGGAT | |

**Table S13B.** Internal oligos used during the second amplification round. (*Reference sequence MN908947.3Wuhan-Hu-1*)

| Fragment | 5'END | Sequence | Product Size |
|---|---|---|---|
| **INT.1** | 21517 | AGAGTTGTTATTTCTAGTGATGTTCTTGT | 448 |
| | 21964 | TTGAAATTCACAGACTTTAATAACAACA | |
| **INT.2** | 21907 | GTCCCTACTTATTGTTAATAACGCT | 345 |
| | 22251 | GGCAAATCTACCAATGGTT | |
| **INT.3** | 22170 | TATATTCTAAGCACACGCCTATTAAT | 466 |
| | 22635 | ATTCTCTTCCTGTTCCAAGCAT | |
| **INT.4** | 22407 | ATGGAACCATTACAGATGCTGT | 482 |
| | 22888 | ATCAAGATTGTTAGAATTCCAAGCTAT | |
| **INT.5** | 22768 | TAGAGGTGATGAAGTCAGACAAAT | 371 |
| | 23138 | CACAAACAGTTGCTGGTGCA | |
| **INT.6** | 23069 | GTTGGTTACCAACCATACAGAGTAGTAGT | 358 |
| | 23426 | CAGGGACTTCTGTGCAGTTA | |
| **INT.7** | 23347 | CAGTGTTATAACACCAGGAACAAAT | 367 |
| | 23713 | ATTTGTGGGTATGGCAATAGAGTTA | |
| **INT.8** | 23636 | GTTGCTTACTCTAATAACTCTATTGCCAT | 439 |
| | 24074 | CATATTGTTTGATGAAGCCAGCA | |
| **INT.9** | 23987 | CCATCAAAACCAAGCAAGAGGT | 313 |
| | 24299 | GTGTAACTCCAATACCATTAAACCTAT | |
| **INT.10** | 24147 | CACCTTTGCTCACAGATGAAATGAT | 470 |
| | 24616 | GATTTCTGCAGCTCTAATTAATTGTT | |
| **INT.11** | 24507 | CACGTCTTGACAAAGTTGAGGCT | 376 |
| | 24882 | CTTTGTGTTACAAACCAGTGTGT | |
| **INT.12** | 24792 | CTGCTCCTGCCATTTGTCAT | 375 |
| | 25166 | CTTGGAGATCGATGAGAGATTCAT | |
| **INT.13** | 24954 | GAATTGTCAACAACACAGTTTATGAT | 465 |
| | 25418 | GTGAAGATTCTCATAAACAAATCCAT | |

**SUPPLEMENTARY FIGURES S1-S14.**

Bar plot of deletions for the 18 patients and by amplicons are provided as supplementary figures S1 to S14 for nucleotides. The x axis provides the multiple alignment (MA) nucleotide positions and the amplitude of the deletions by subregions, and the y axis shows the frequency of the deletion (percentage on the right) and the number of reads (on the left). As no insertions were observed the MA positions correspond to S gene positions. Discontinued lines indicates S1/S2 (left) and S2' (right) cleavage sites. (P01, patient 1; N01, nucleotide amplicon 1; 016 indicates the in-frame nucleotide

**Figure S1.** Bar plot of deletions in amplicon N01 in the 18 patients (P01-P18) at the nucleotide level.

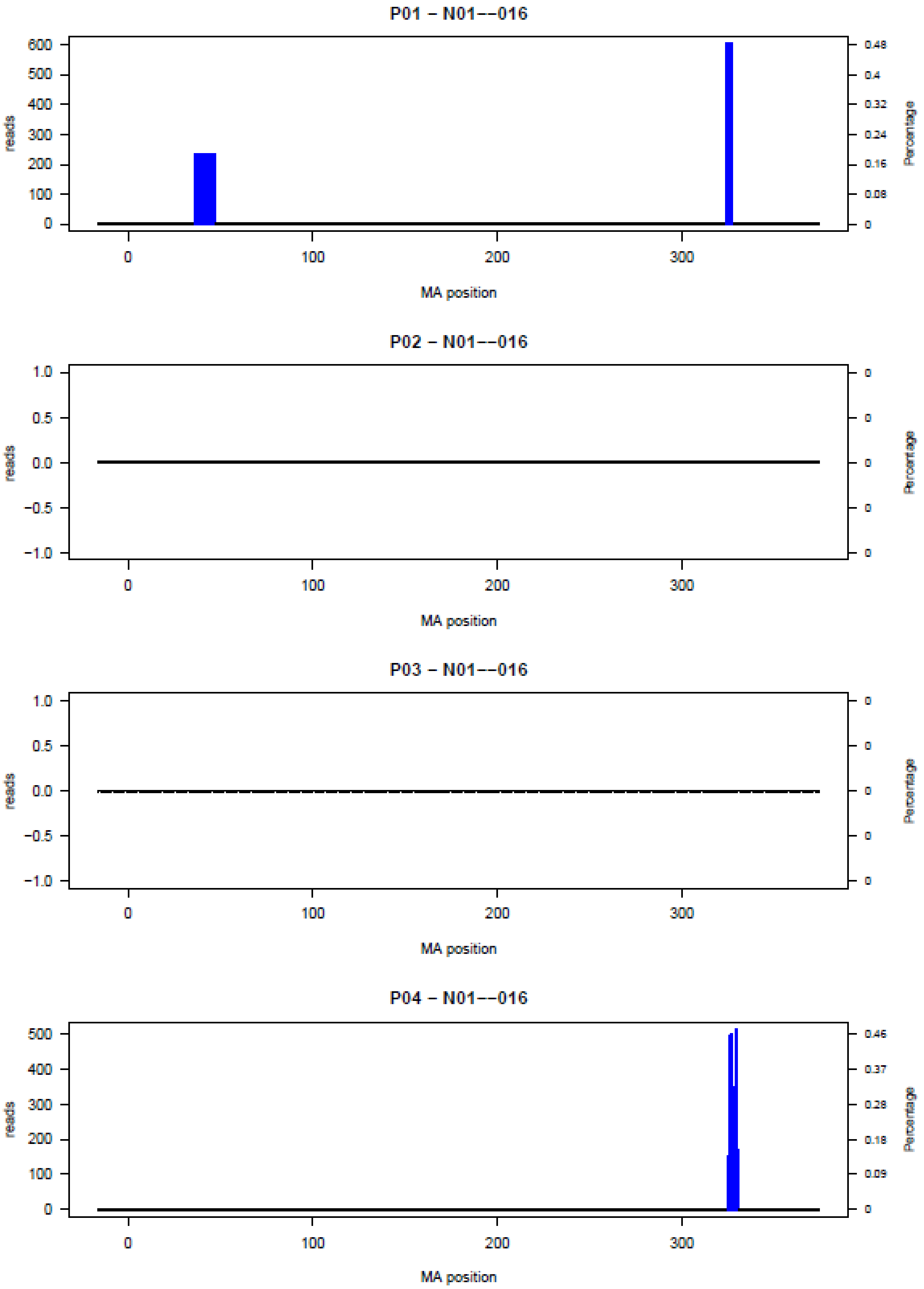

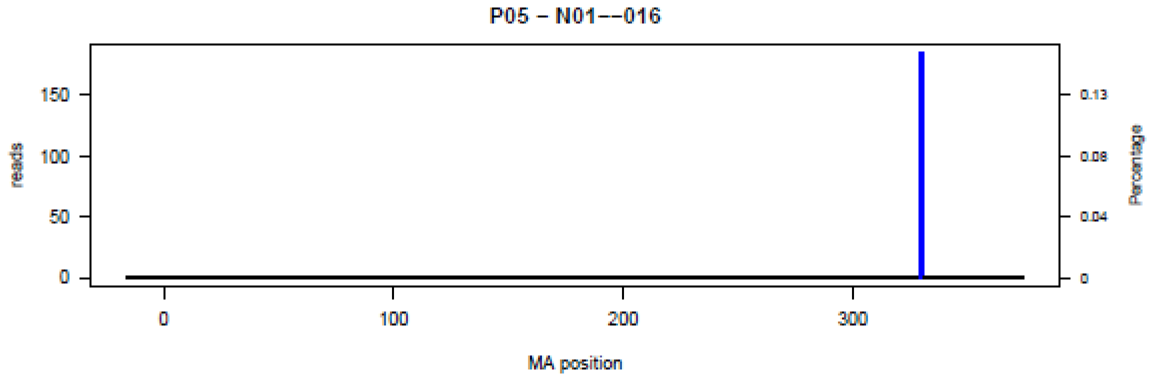
P05 – N01––016
reads
Percentage
MA position

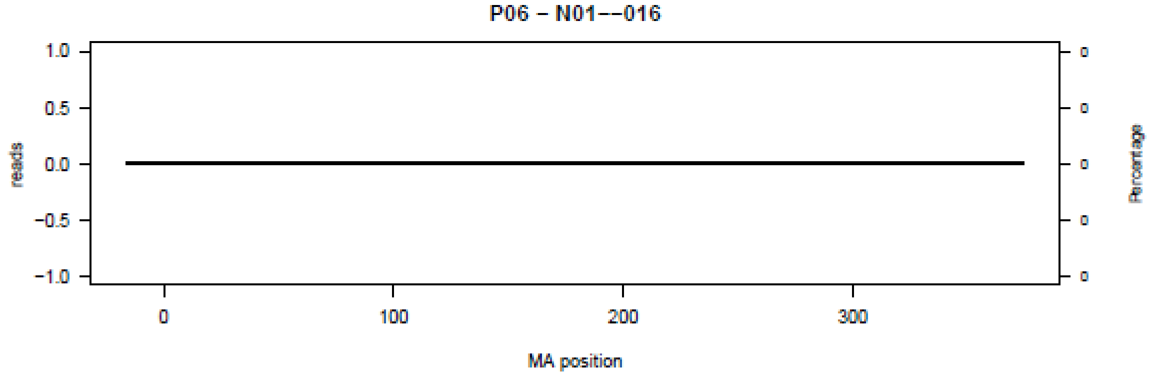
P06 – N01––016
reads
Percentage
MA position

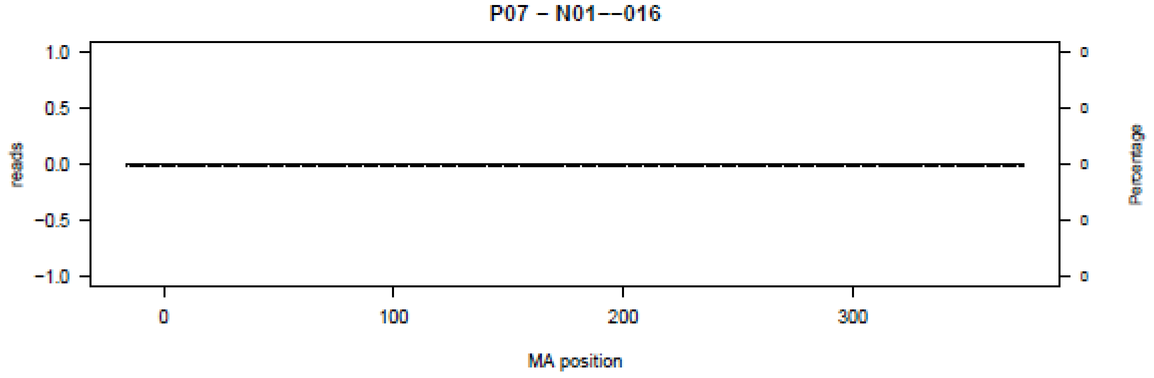
P07 – N01––016
reads
Percentage
MA position

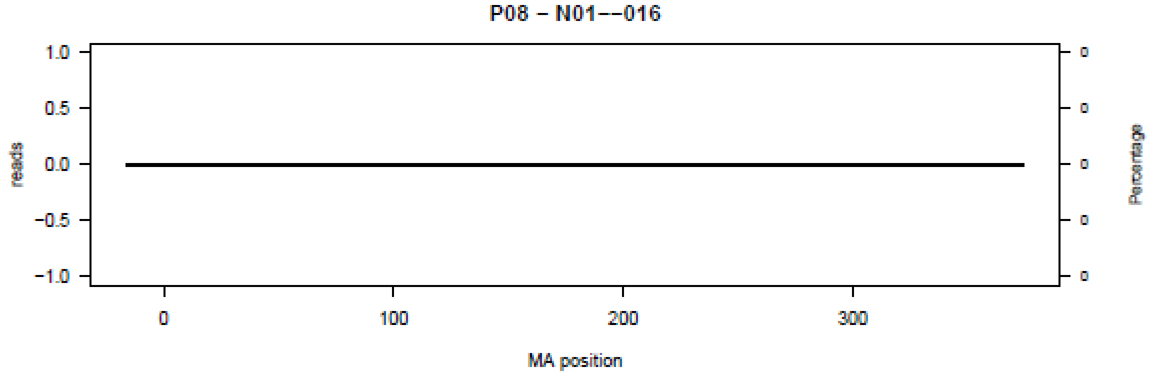
P08 – N01––016
reads
Percentage
MA position

P09 – N01--016

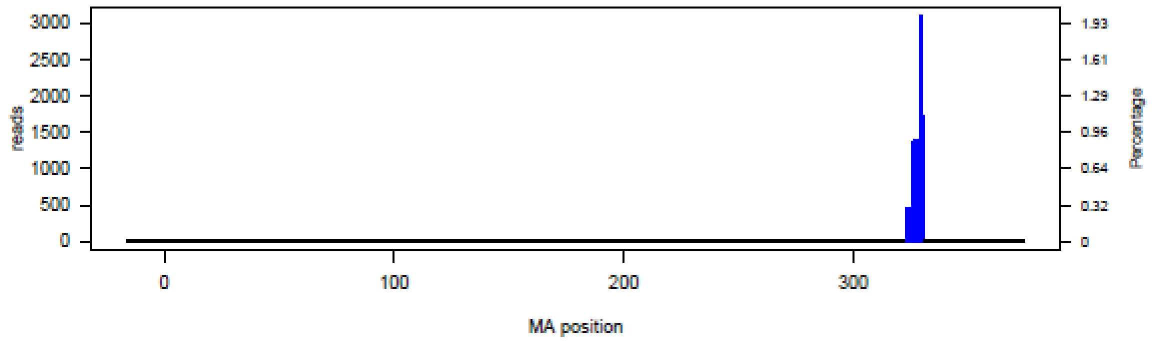


P10 – N01--016

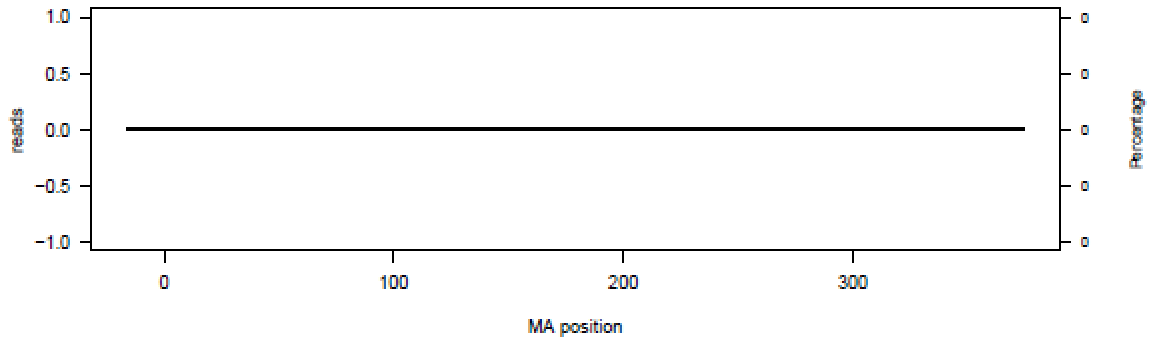


P11 – N01--016

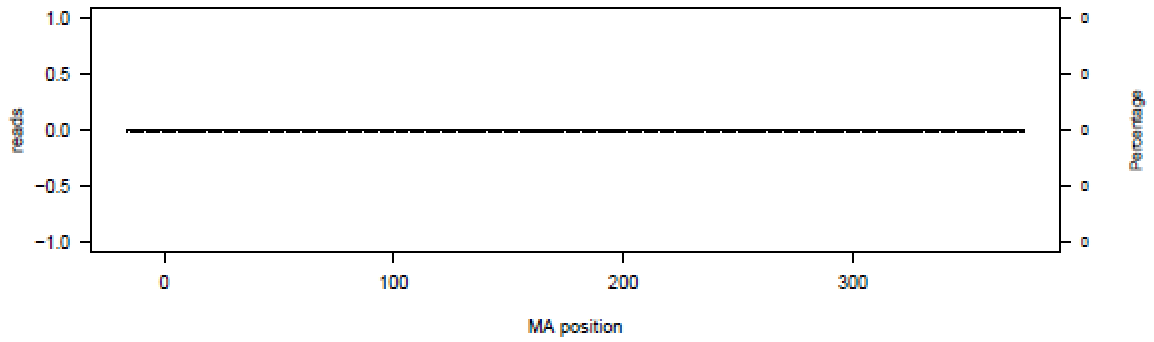


P12 – N01--016

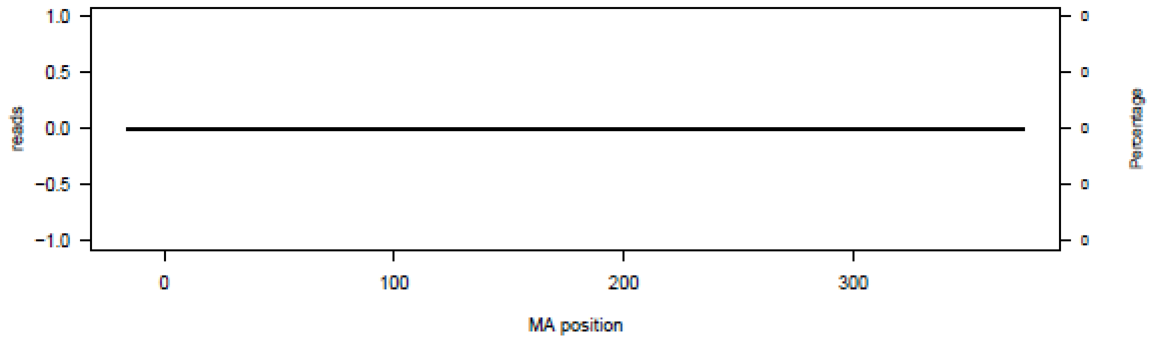

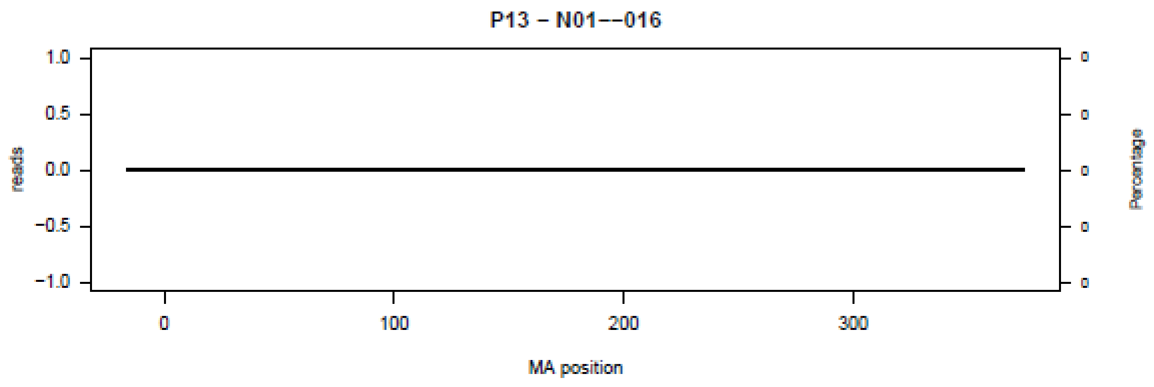
P13 – N01--016
reads
MA position
Percentage

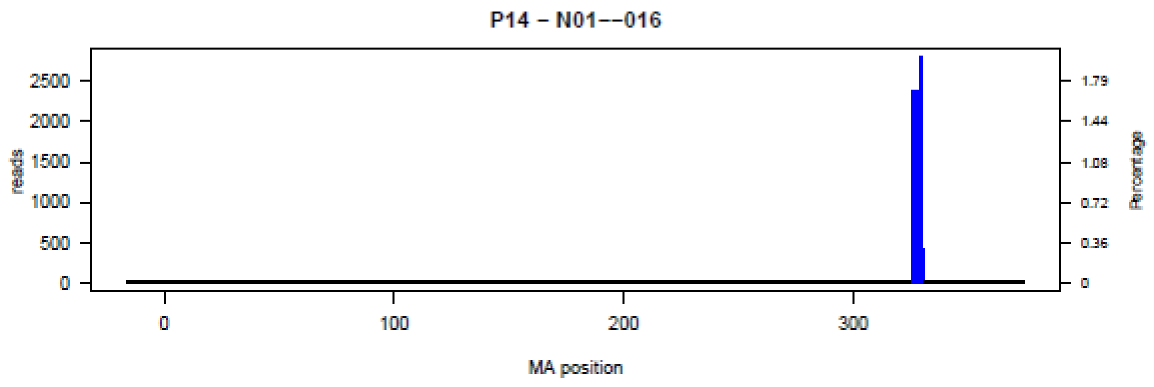
P14 – N01--016
reads
MA position
Percentage

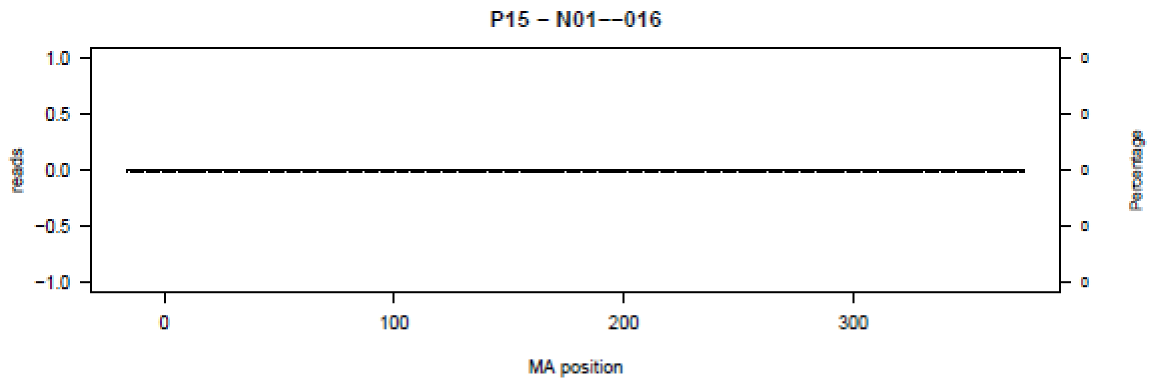
P15 – N01--016
reads
MA position
Percentage

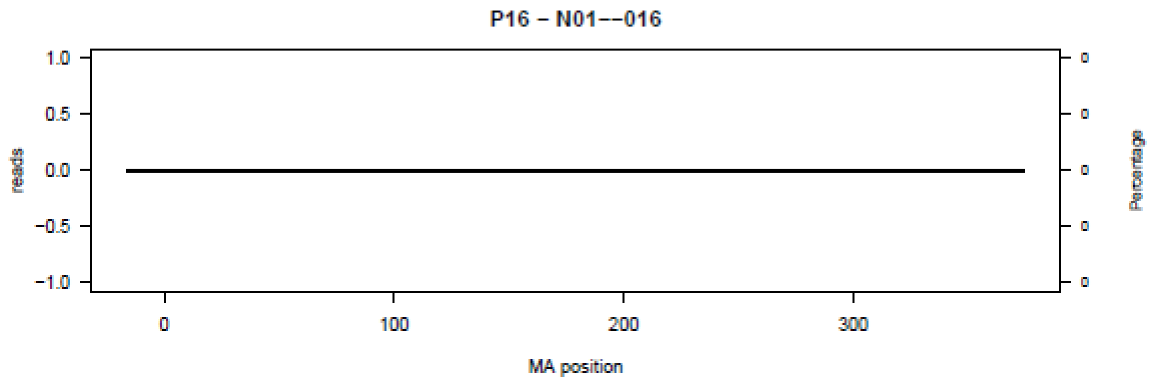
P16 – N01--016
reads
MA position
Percentage

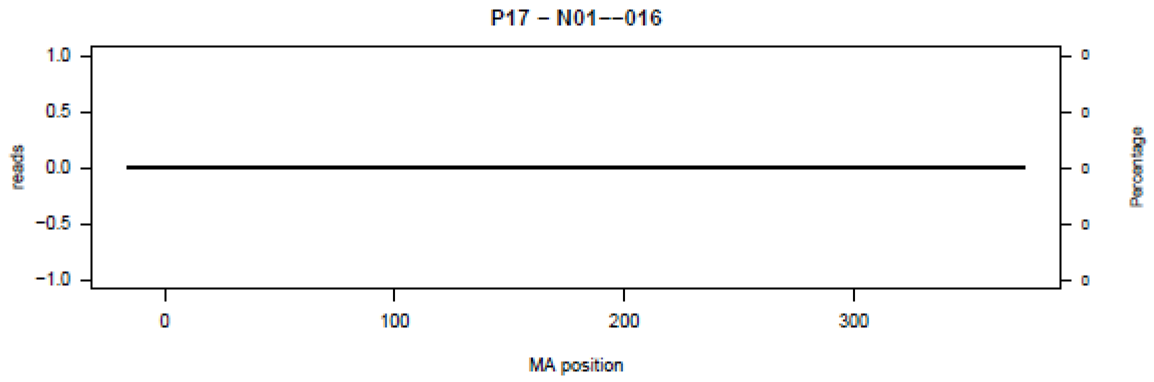
P17 – N01--016
1.0
0.5
0.0
-0.5
-1.0
reads
0
100
200
300
MA position
Percentage

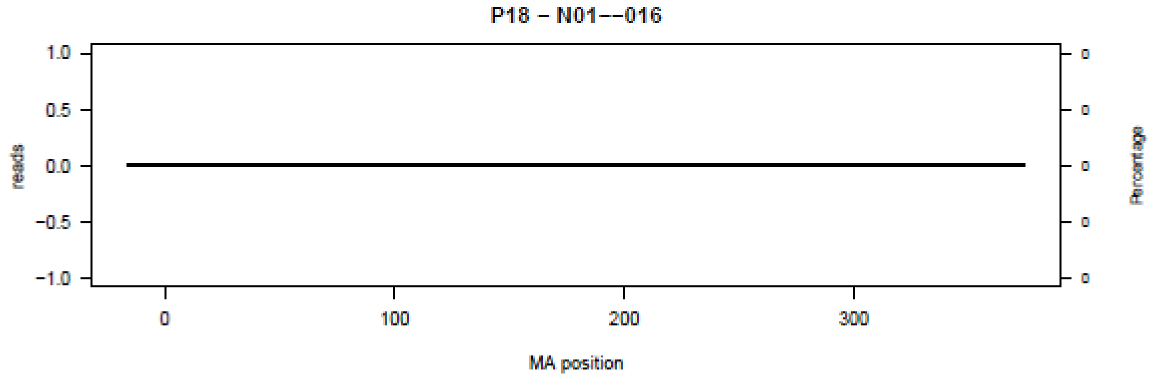
P18 – N01--016
1.0
0.5
0.0
-0.5
-1.0
reads
0
100
200
300
MA position
Percentage

**Figure S2.** Bar plot of deletions in amplicon N02 in the 18 patients (P01-P18) at the nucleotide level.

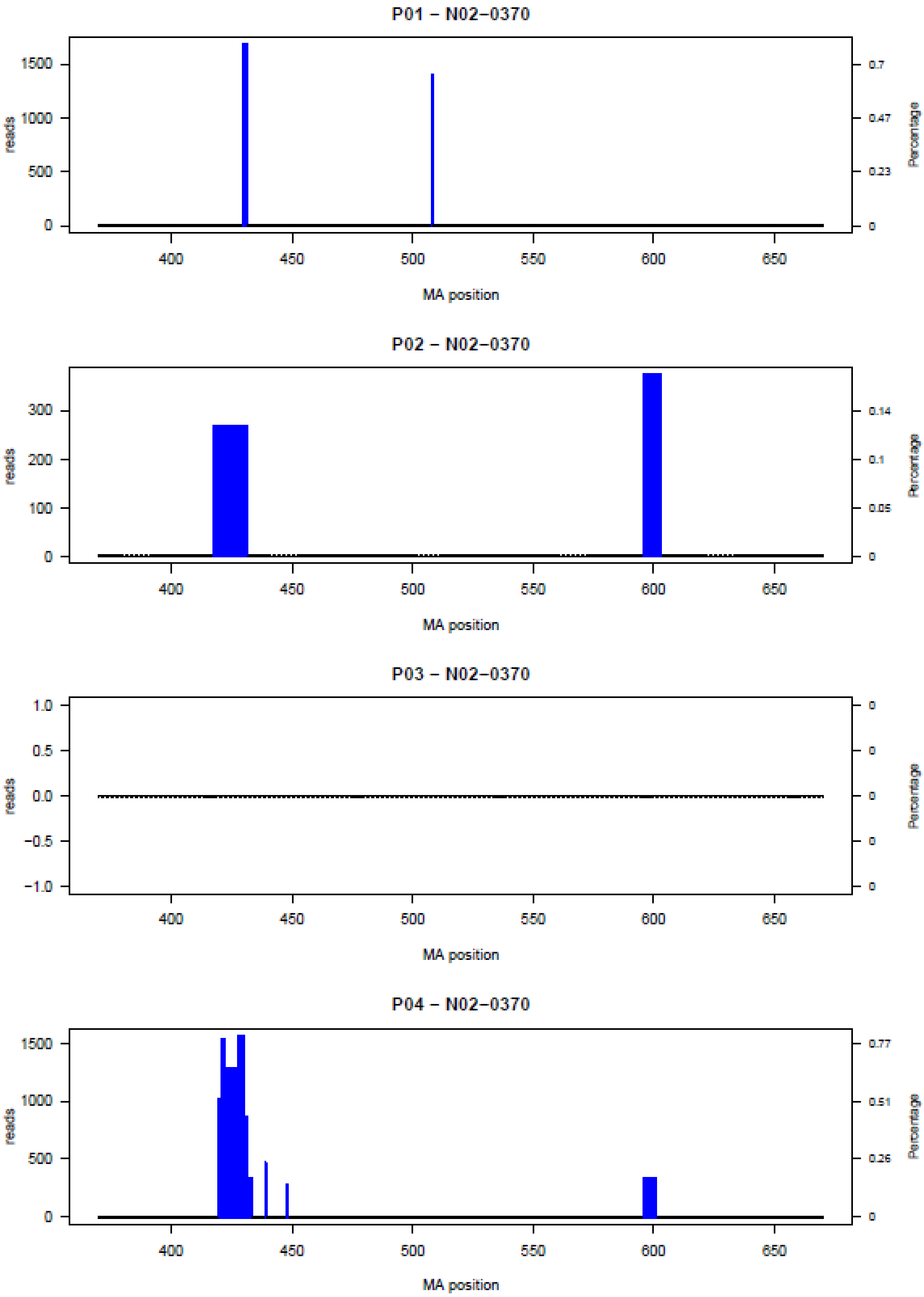

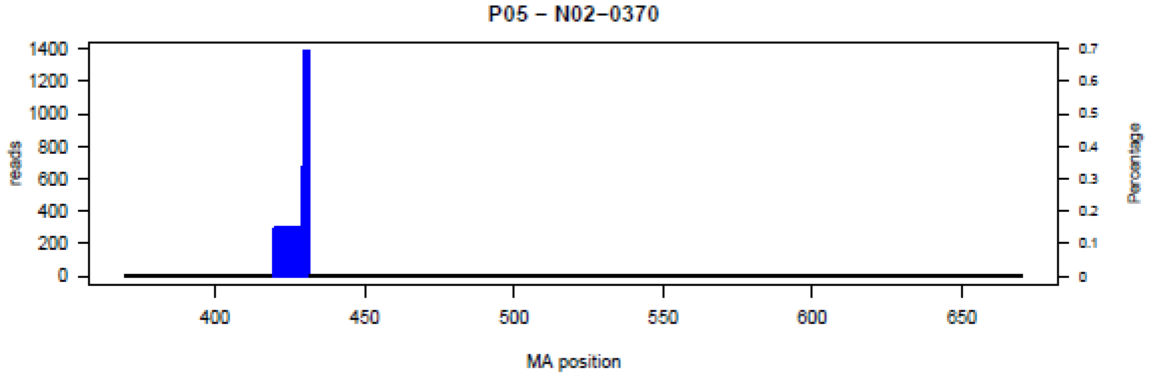
P05 – N02–0370
reads
Percentage
MA position

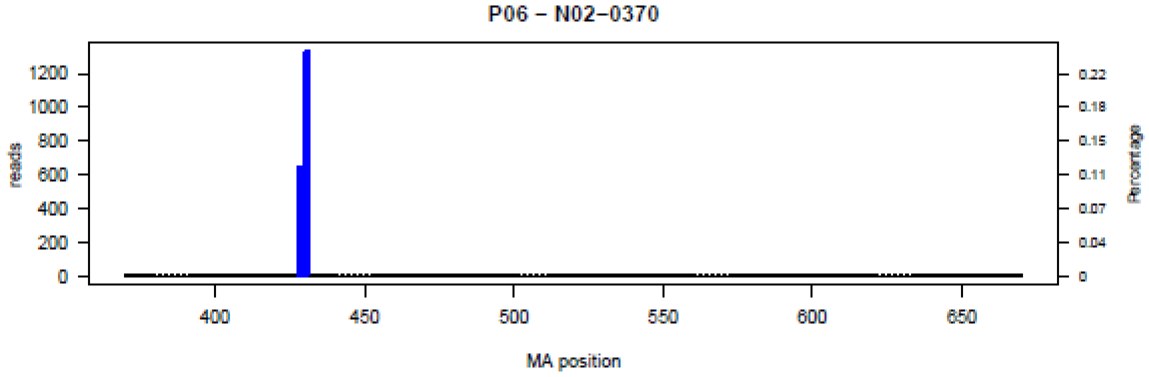
P06 – N02–0370
reads
Percentage
MA position

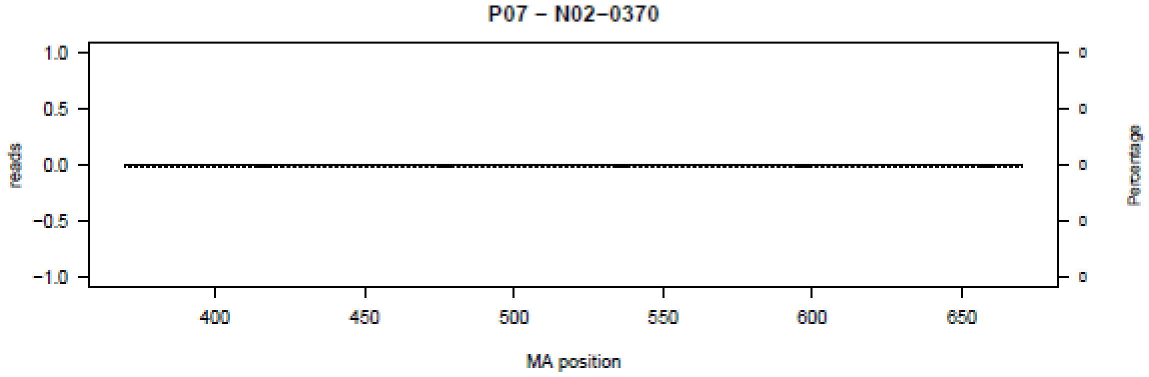
P07 – N02–0370
reads
Percentage
MA position

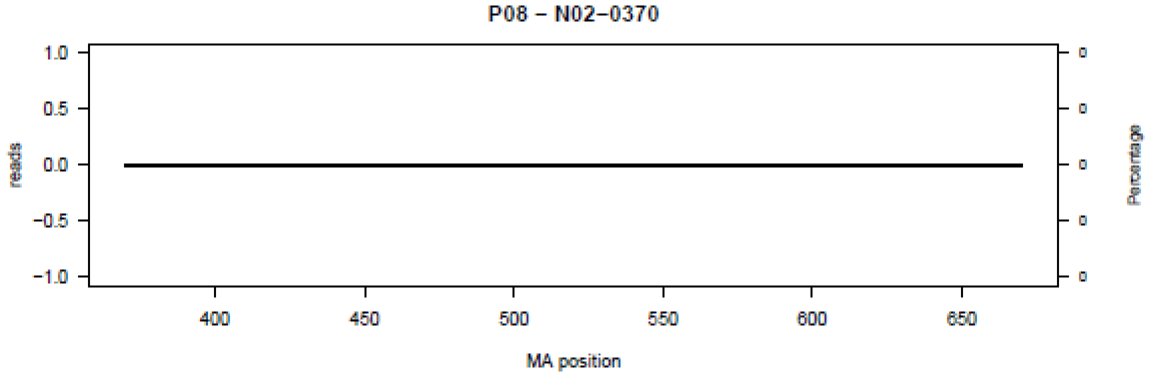
P08 – N02–0370
reads
Percentage
MA position

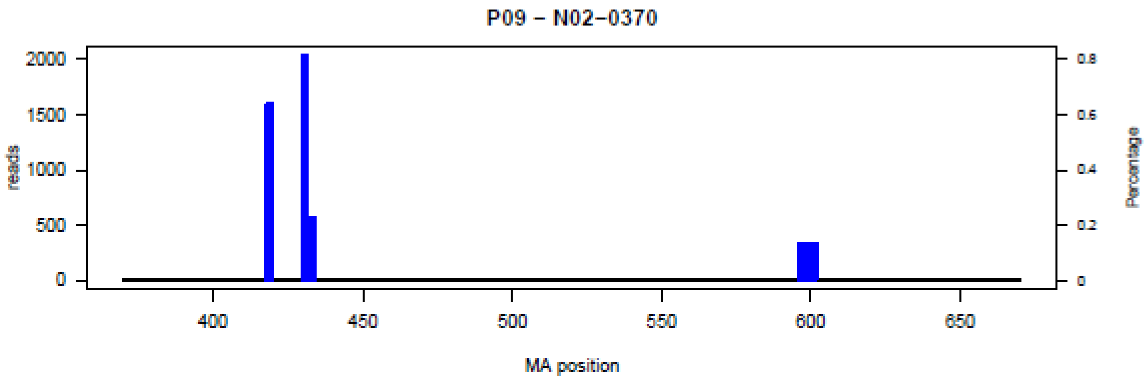
P09 – N02–0370
reads
2000
1500
1000
500
0
Percentage
0.8
0.6
0.4
0.2
0
400
450
500
550
600
650
MA position

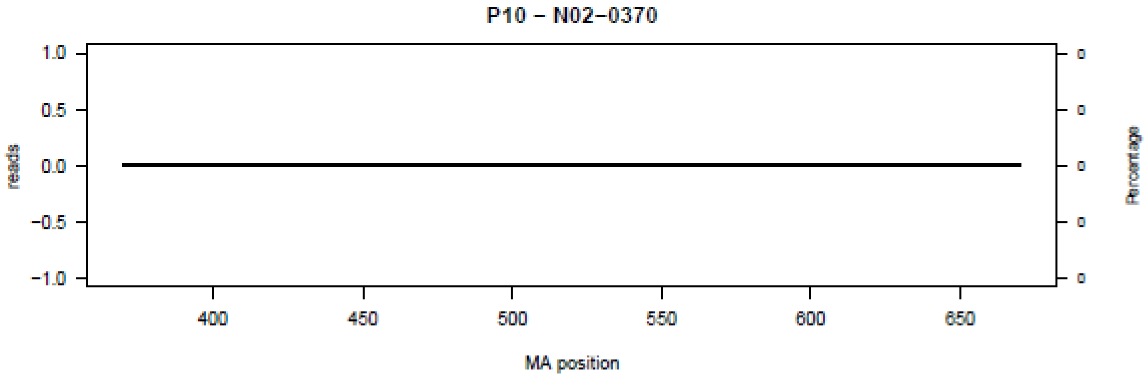
P10 – N02–0370
reads
1.0
0.5
0.0
−0.5
−1.0
Percentage
0
0
0
0
0
400
450
500
550
600
650
MA position

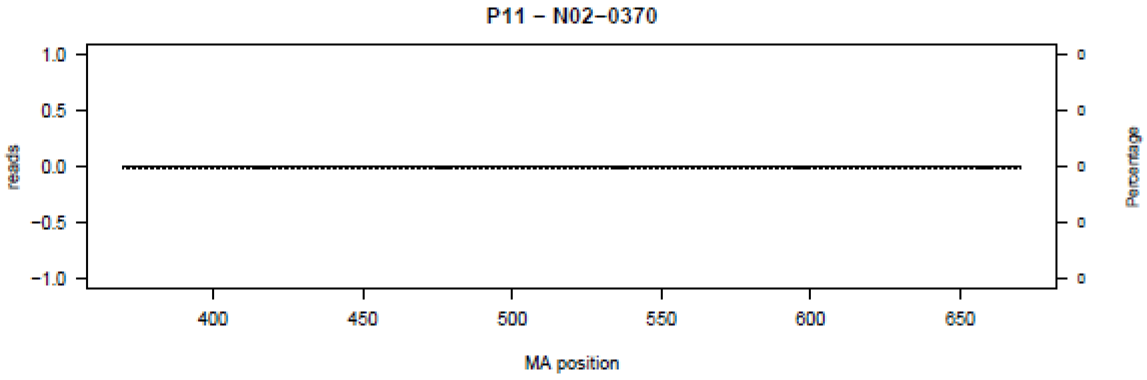
P11 – N02–0370
reads
1.0
0.5
0.0
−0.5
−1.0
Percentage
0
0
0
0
0
400
450
500
550
600
650
MA position

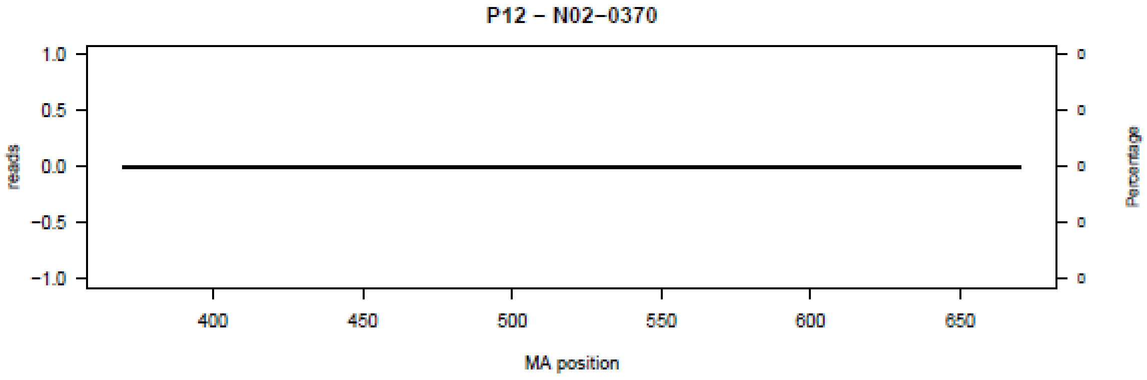
P12 – N02–0370
reads
1.0
0.5
0.0
−0.5
−1.0
Percentage
0
0
0
0
0
400
450
500
550
600
650
MA position

**P13 – N02–0370**

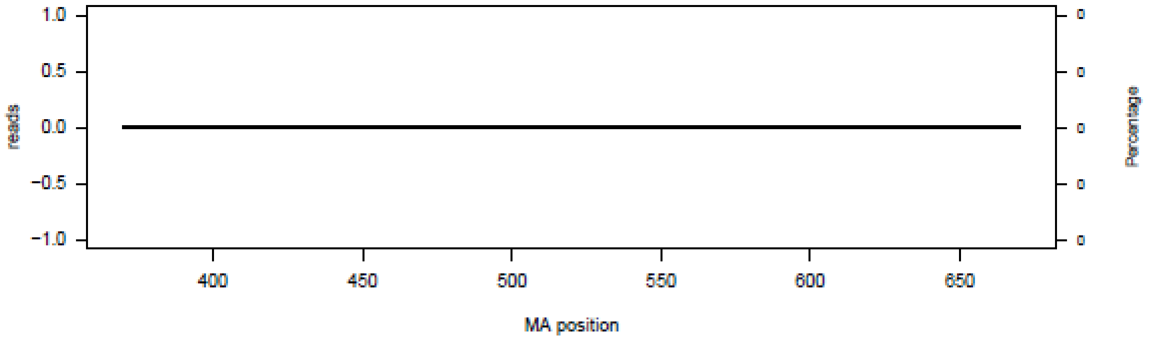


**P14 – N02–0370**

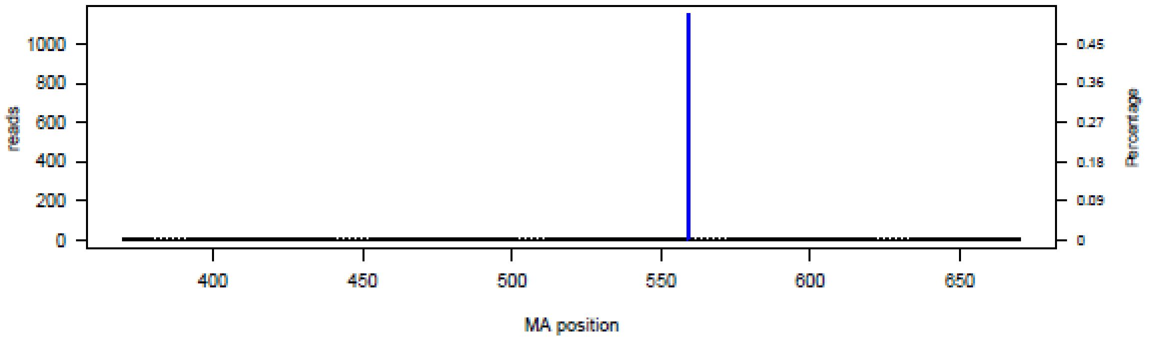


**P15 – N02–0370**

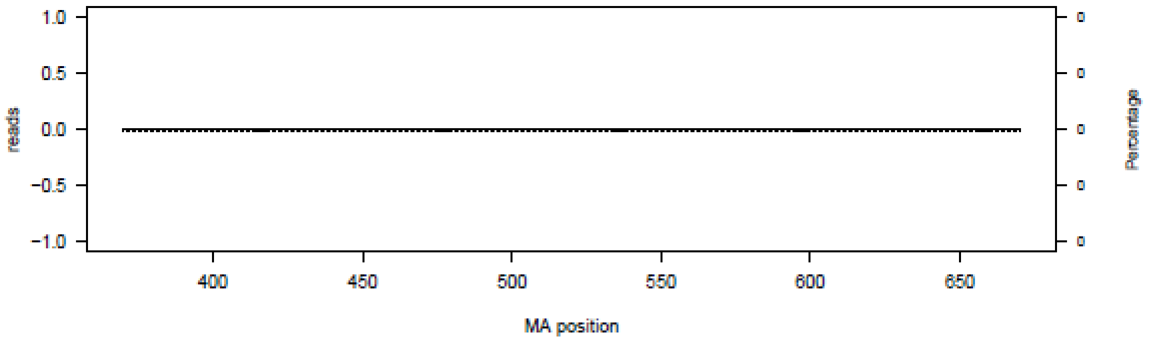


**P16 – N02–0370**

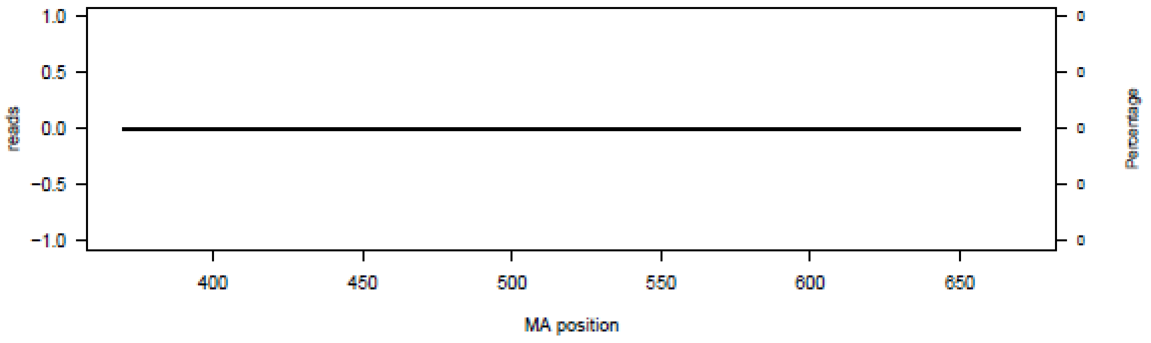

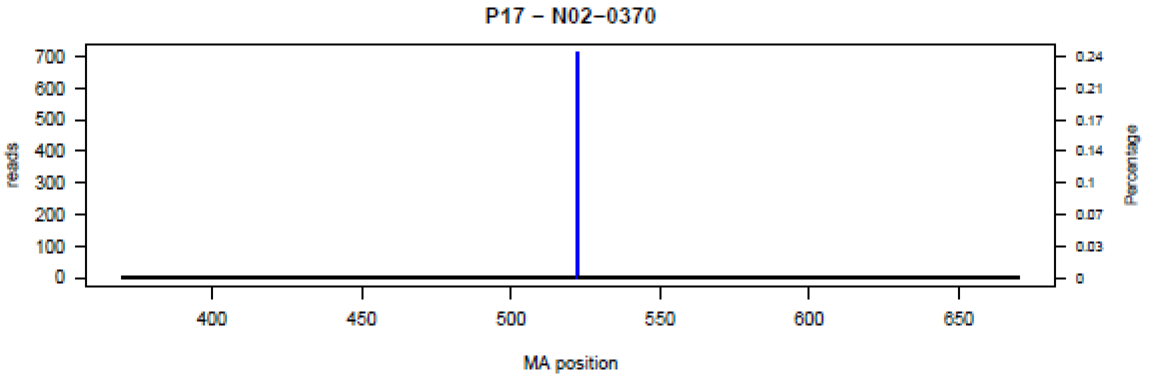

P17 – N02–0370
reads
Percentage
MA position


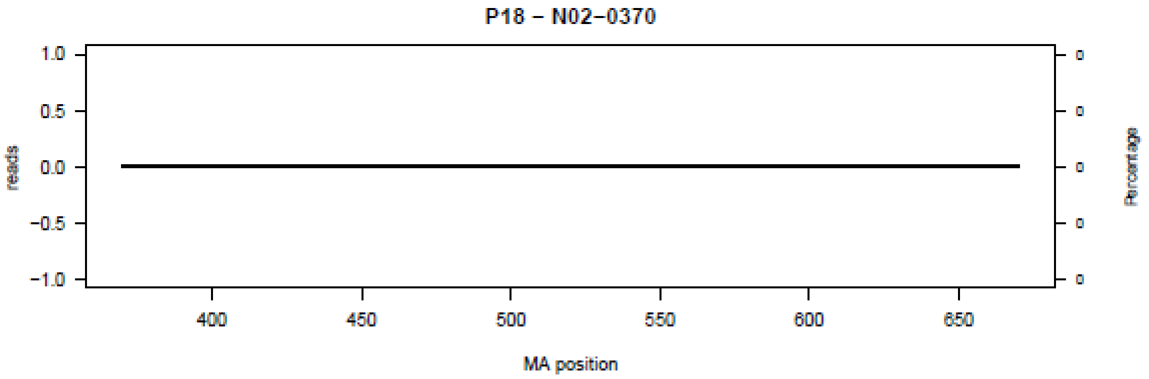

P18 – N02–0370
reads
Percentage
MA position

**Figure S3.** Bar plot of deletions in amplicon N03 in the 18 patients (P01-P18) at the nucleotide level.

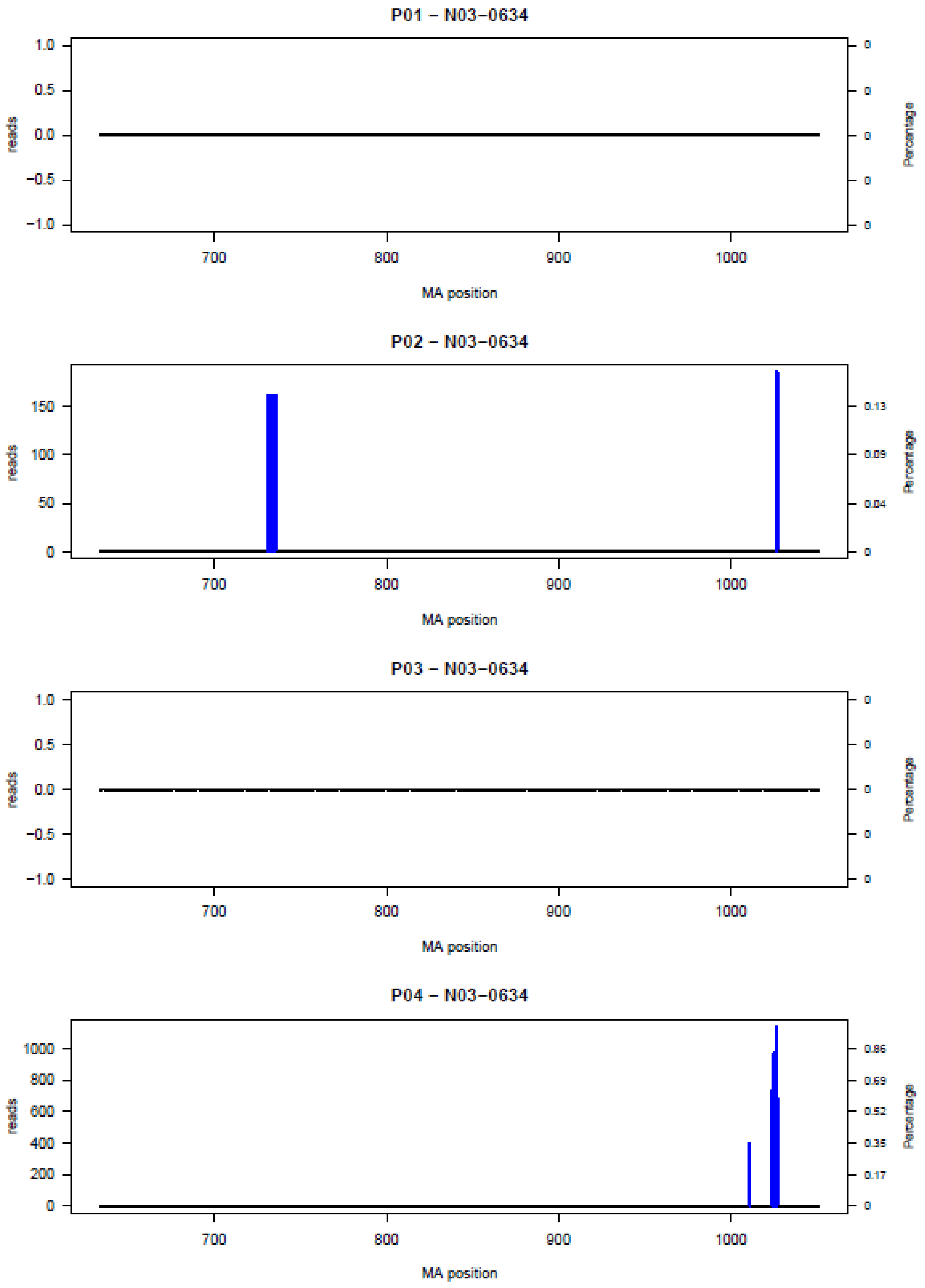

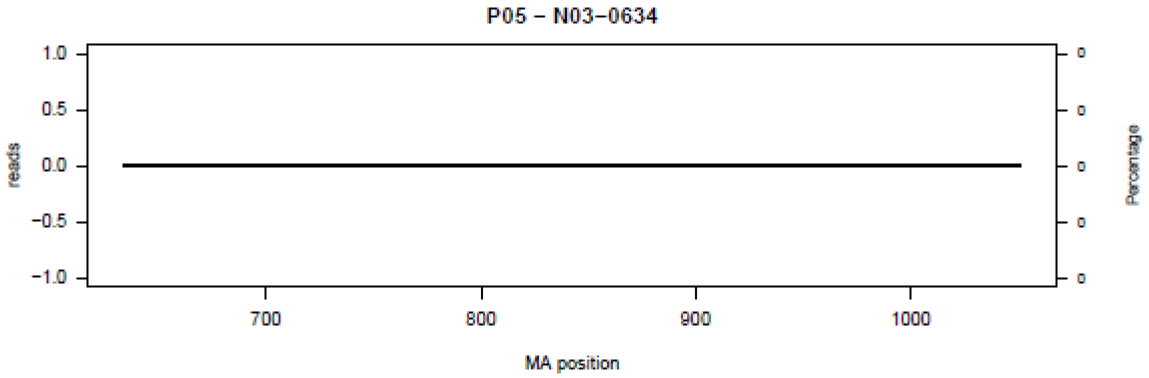
P05 – N03–0634
reads
1.0
0.5
0.0
−0.5
−1.0
0
Percentage
700
800
900
1000
MA position

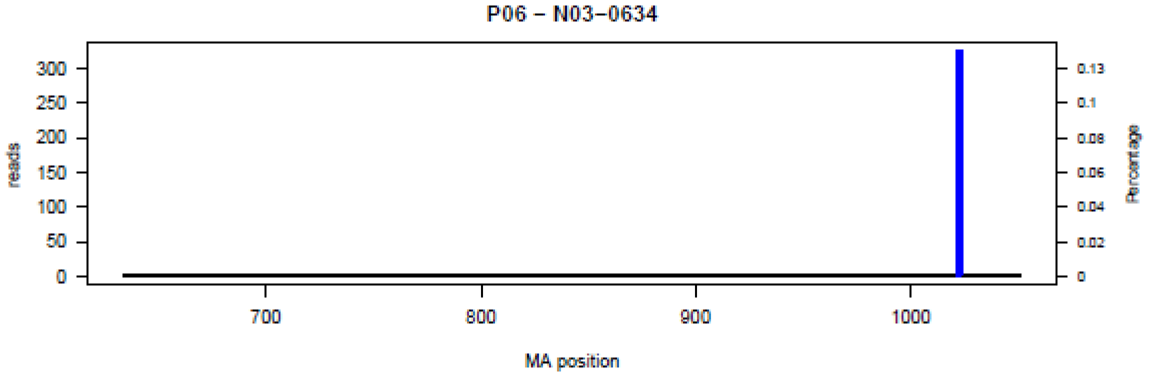
P06 – N03–0634
reads
300
250
200
150
100
50
0
0.13
0.1
0.08
0.06
0.04
0.02
0
Percentage
700
800
900
1000
MA position

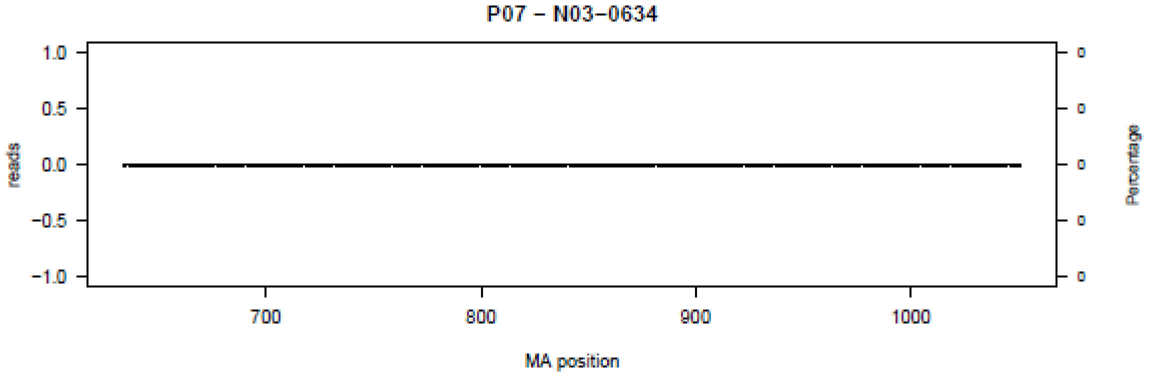
P07 – N03–0634
reads
1.0
0.5
0.0
−0.5
−1.0
0
Percentage
700
800
900
1000
MA position

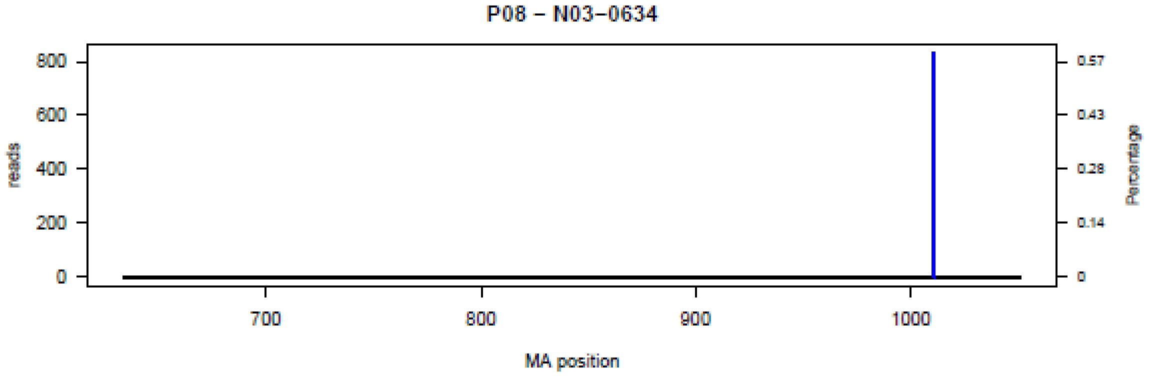
P08 – N03–0634
reads
800
600
400
200
0
0.57
0.43
0.28
0.14
0
Percentage
700
800
900
1000
MA position

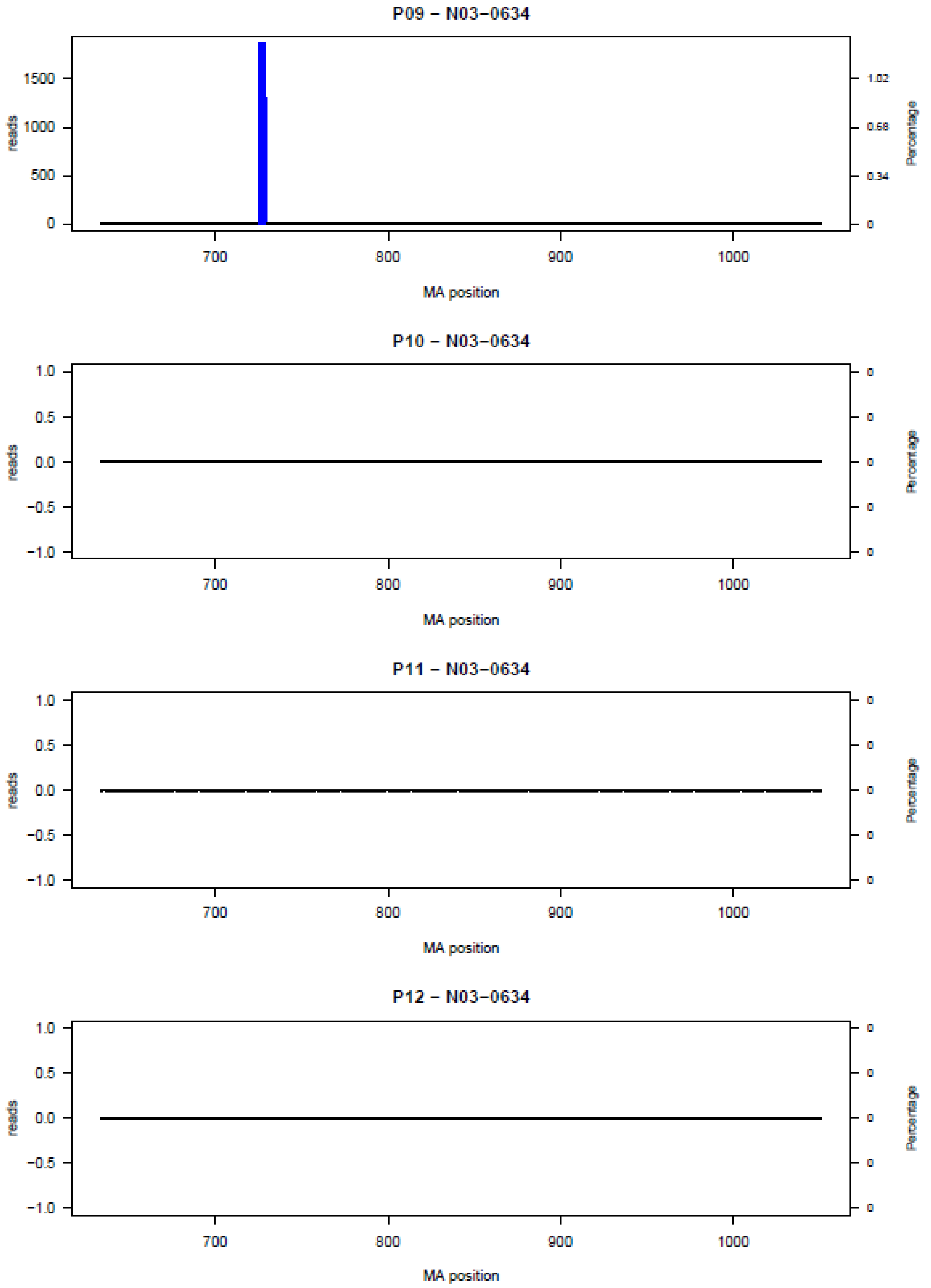
P09 – N03–0634
reads
Percentage
1500
1000
500
0
1.02
0.68
0.34
0
700
800
900
1000
MA position
P10 – N03–0634
1.0
0.5
0.0
−0.5
−1.0
reads
Percentage
MA position
P11 – N03–0634
reads
Percentage
MA position
P12 – N03–0634
reads
Percentage
MA position

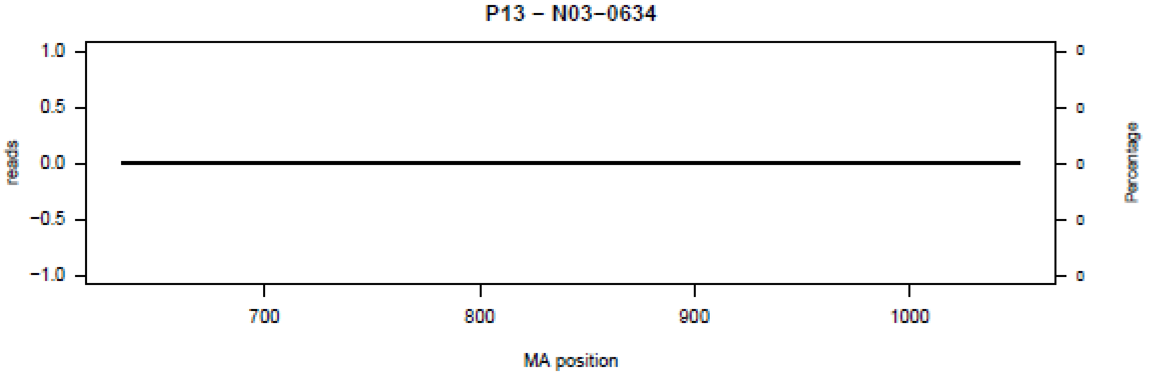
P13 – N03–0634
1.0
0.5
0.0
−0.5
−1.0
reads
0
0
0
0
0
Percentage
700
800
900
1000
MA position

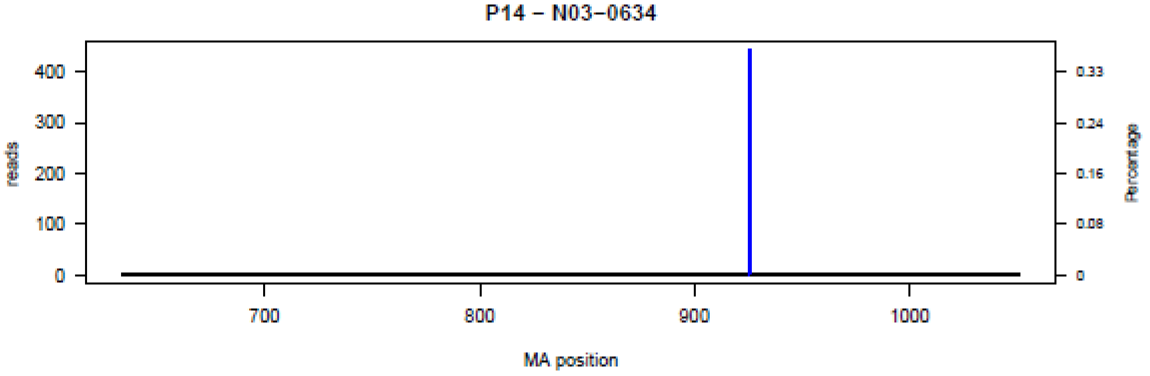
P14 – N03–0634
400
300
200
100
0
reads
0.33
0.24
0.16
0.08
0
Percentage
700
800
900
1000
MA position

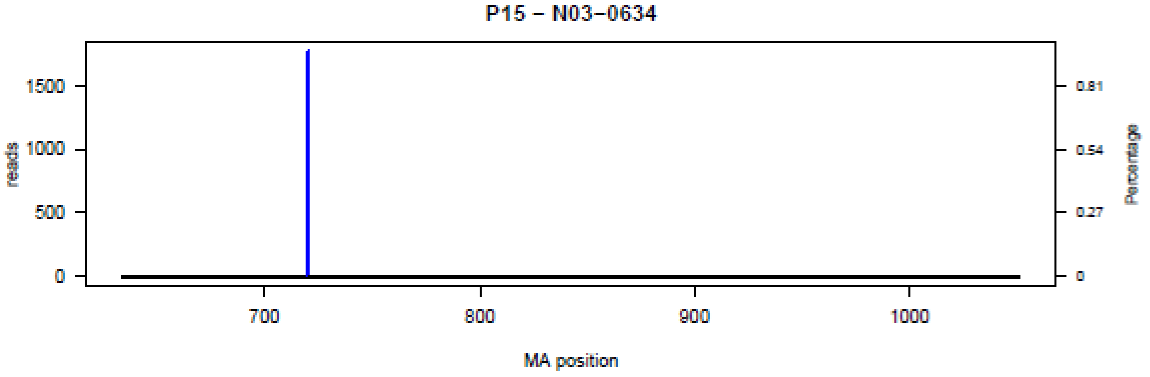
P15 – N03–0634
1500
1000
500
0
reads
0.81
0.54
0.27
0
Percentage
700
800
900
1000
MA position

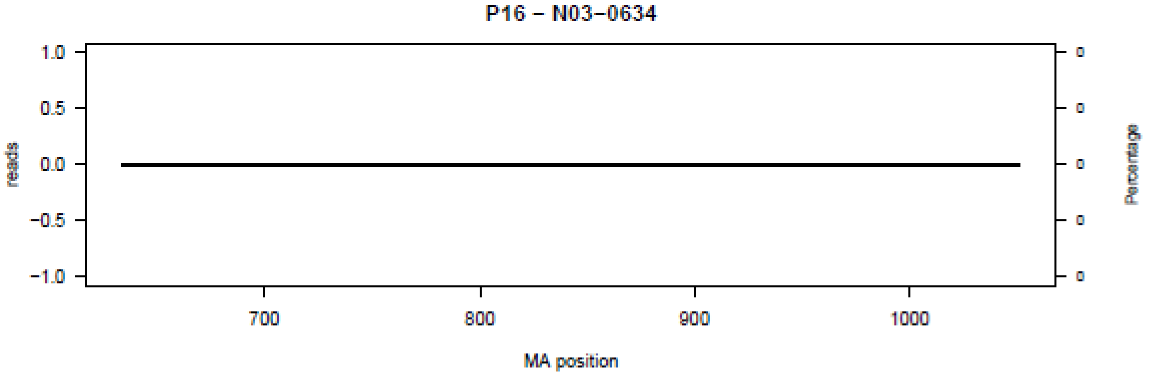
P16 – N03–0634
1.0
0.5
0.0
−0.5
−1.0
reads
0
0
0
0
0
Percentage
700
800
900
1000
MA position

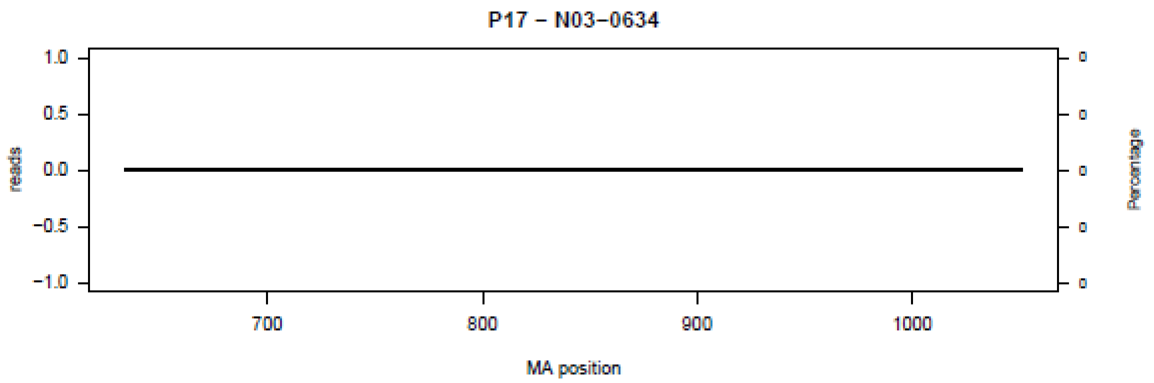
P17 – N03−0634
reads
Percentage
MA position

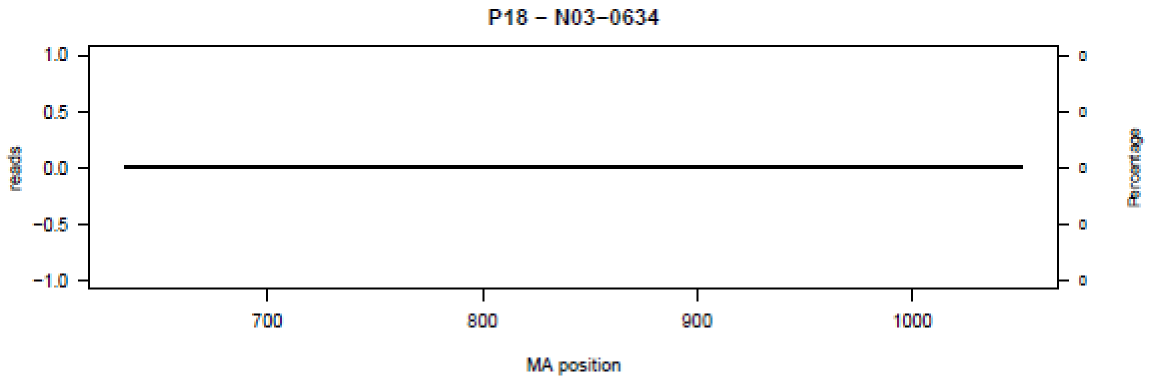
P18 – N03−0634
reads
Percentage
MA position

**Figure S4.** Bar plot of deletions in amplicon N04 in the 18 patients (P01-P18) at the nucleotide level.

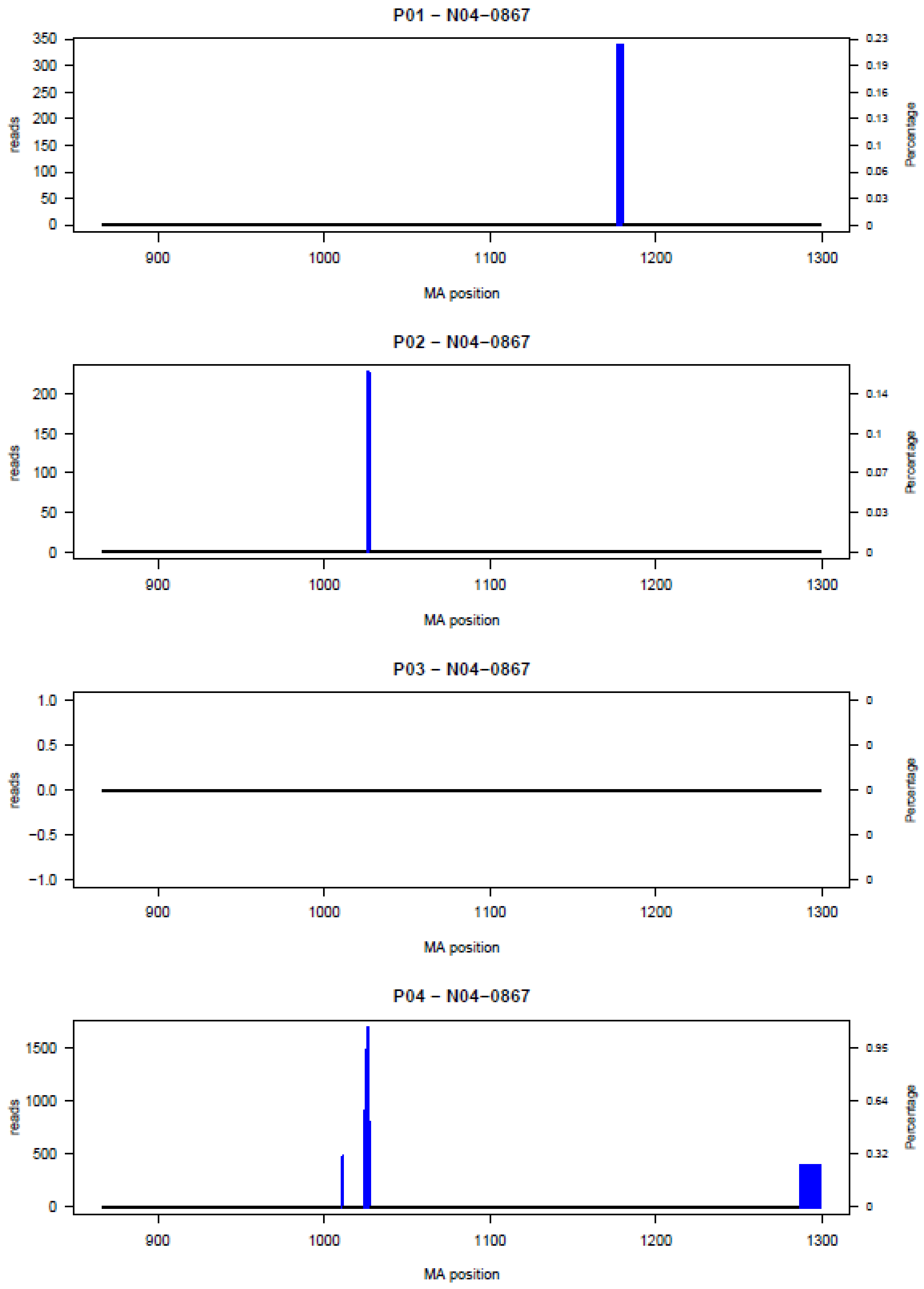

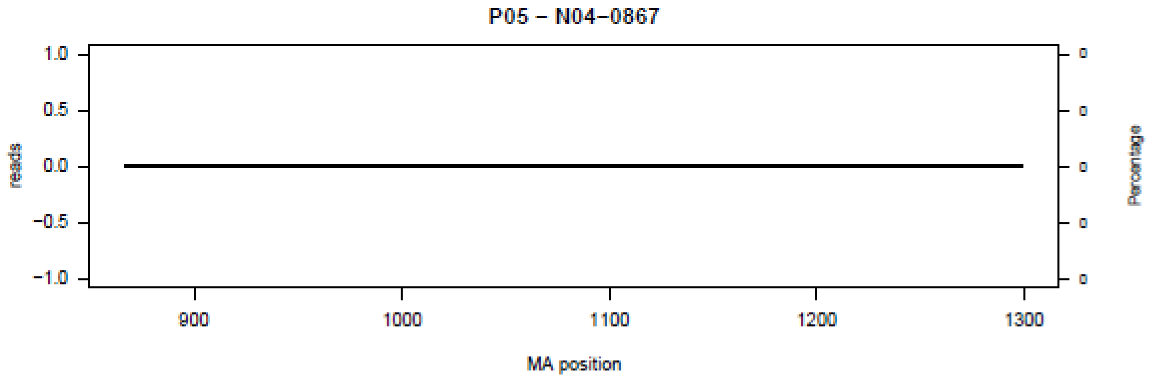
P05 – N04–0867
reads
1.0
0.5
0.0
−0.5
−1.0
900
1000
1100
1200
1300
MA position
Percentage
0
0
0
0
0

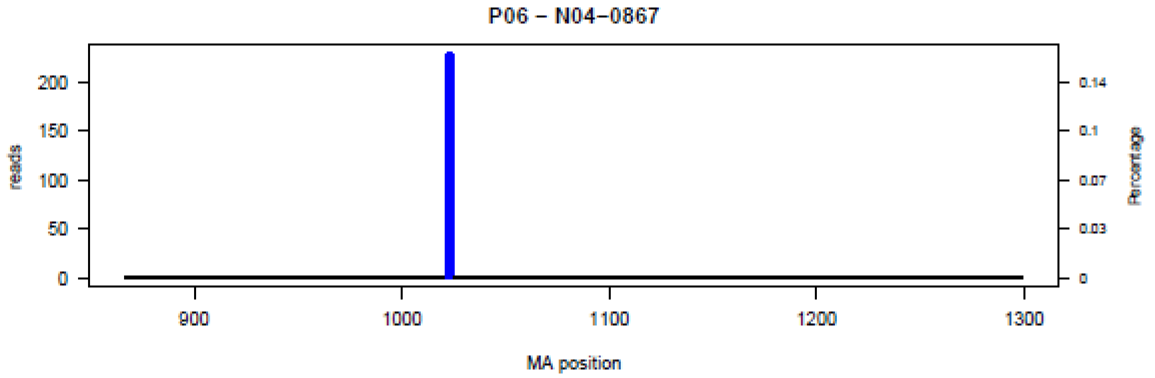
P06 – N04–0867
reads
200
150
100
50
0
900
1000
1100
1200
1300
MA position
Percentage
0.14
0.1
0.07
0.03
0

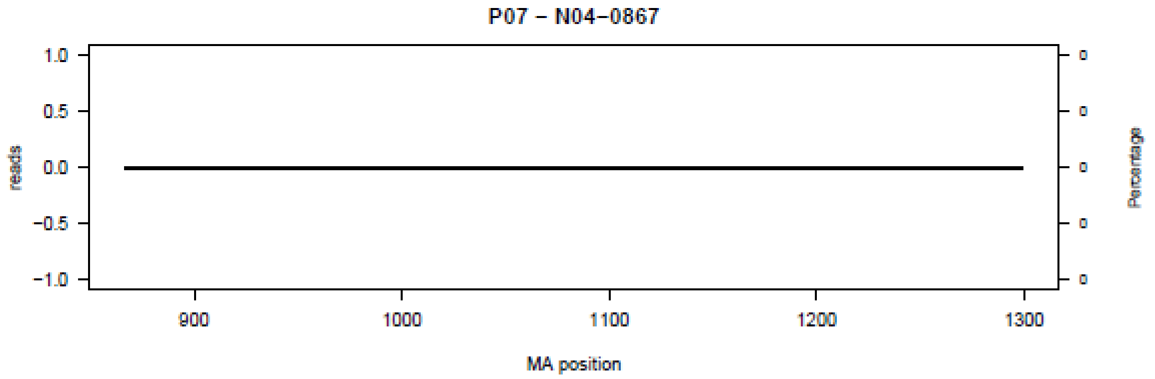
P07 – N04–0867
reads
1.0
0.5
0.0
−0.5
−1.0
900
1000
1100
1200
1300
MA position
Percentage
0
0
0
0
0

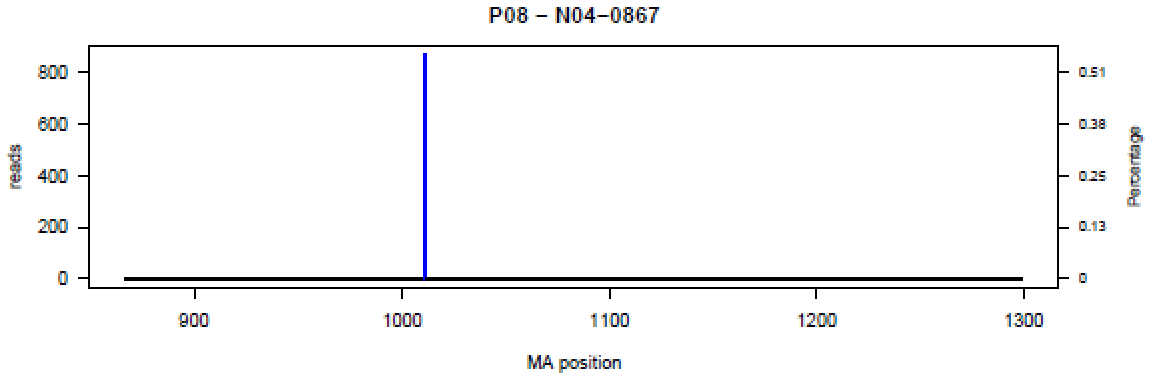
P08 – N04–0867
reads
800
600
400
200
0
900
1000
1100
1200
1300
MA position
Percentage
0.51
0.38
0.25
0.13
0

P09 – N04–0867

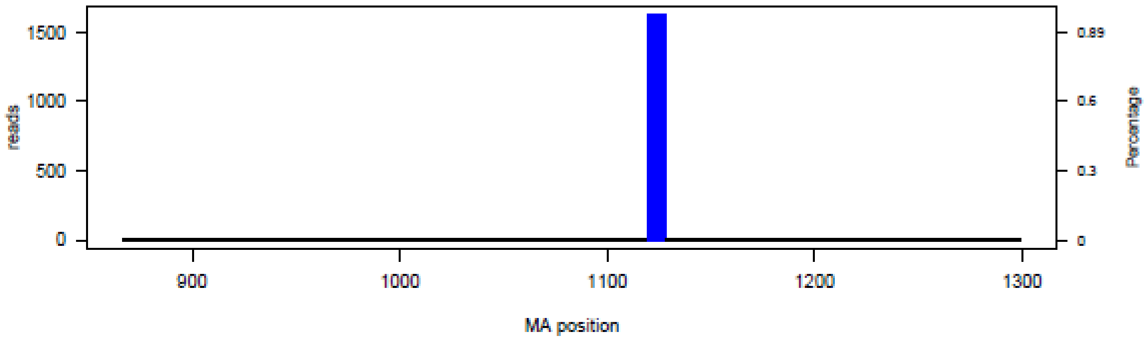


P10 – N04–0867

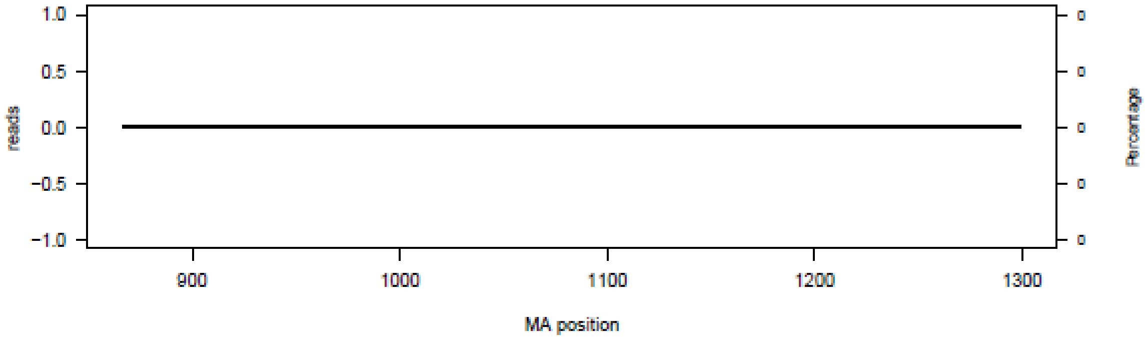


P11 – N04–0867

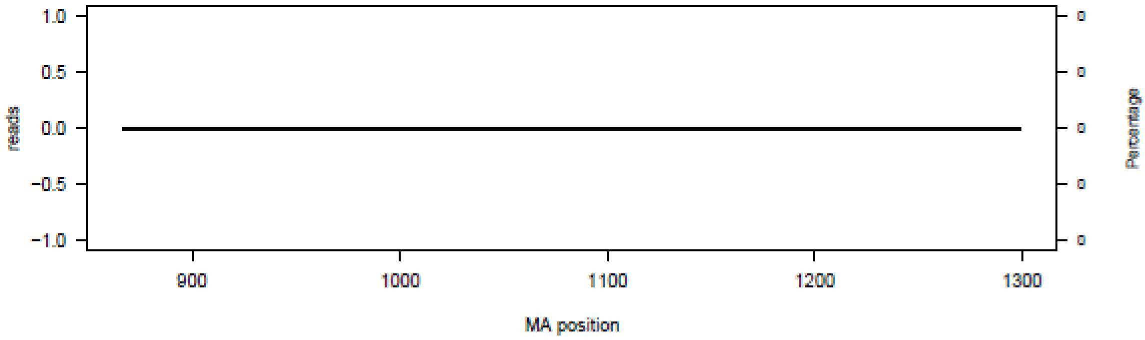


P12 – N04–0867

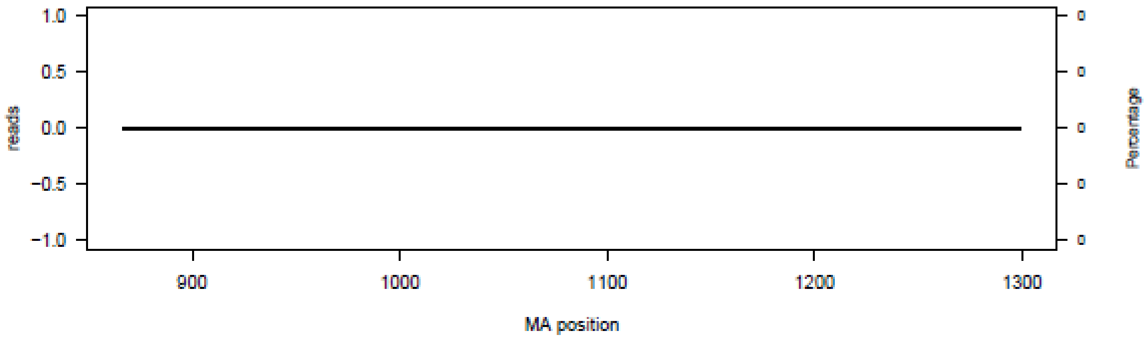

**P13 – N04–0867**

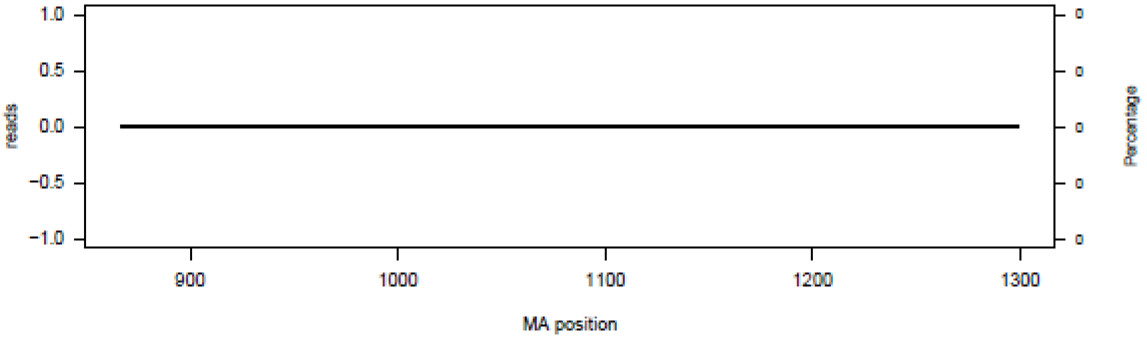


**P14 – N04–0867**

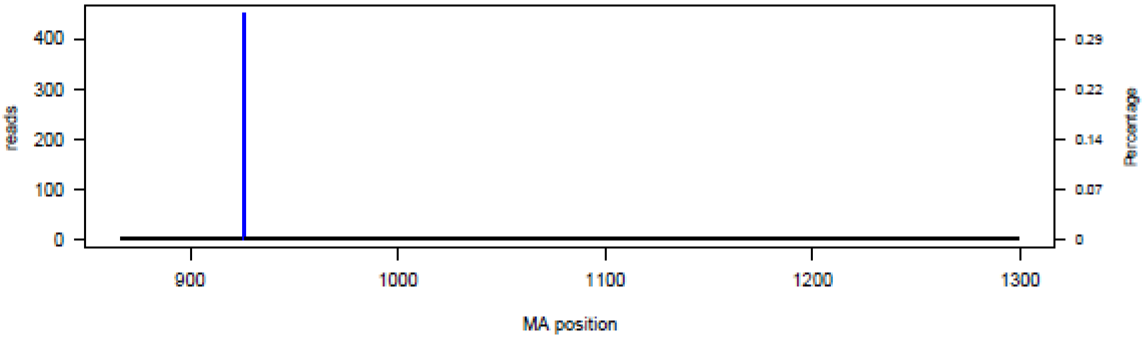


**P15 – N04–0867**

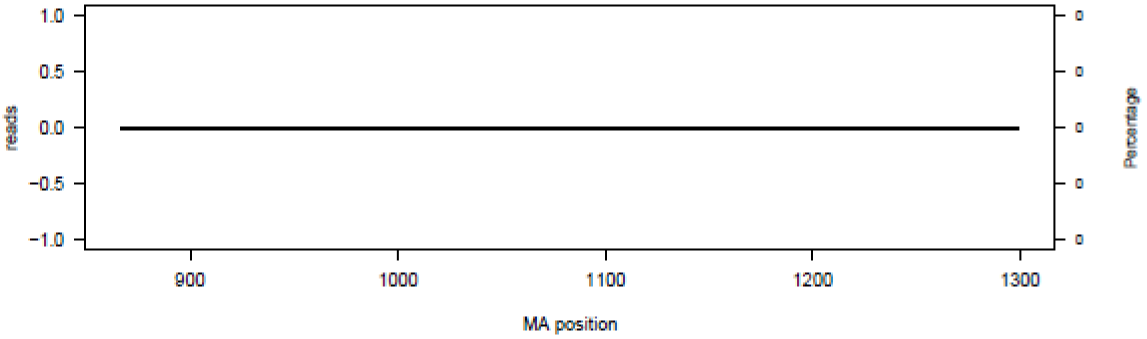


**P16 – N04–0867**

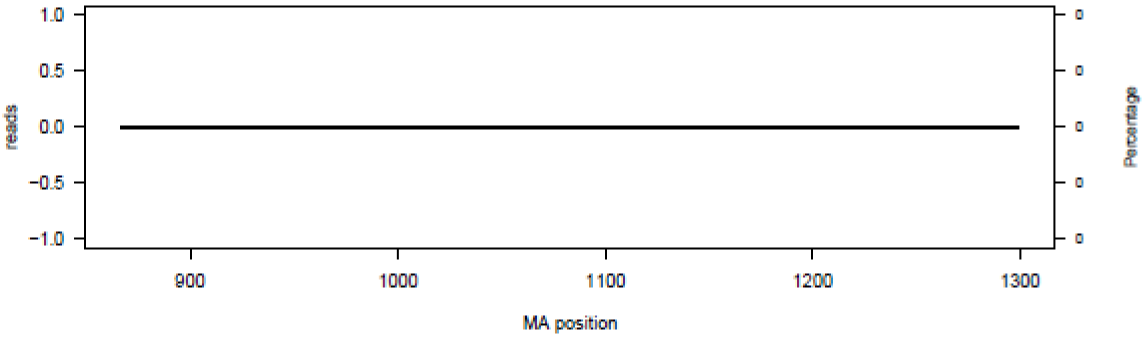

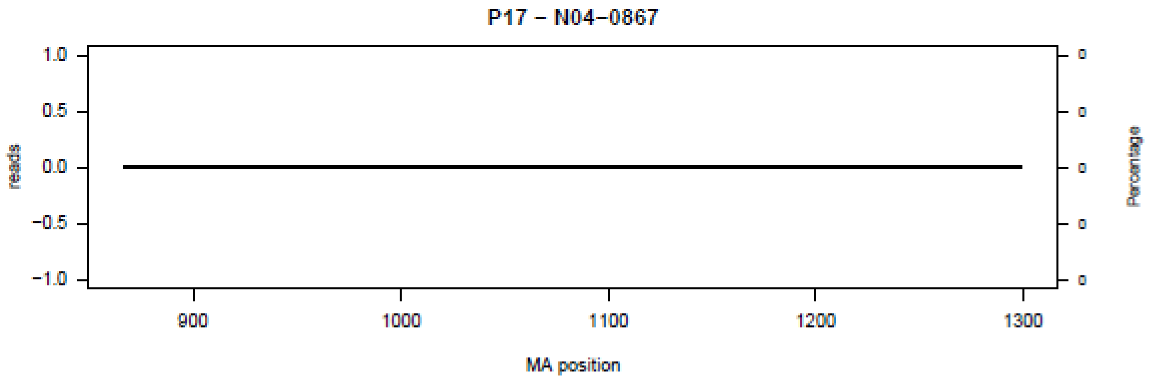
P17 – N04–0867
reads
Percentage
MA position
1.0
0.5
0.0
−0.5
−1.0
900
1000
1100
1200
1300
0
0
0
0
0

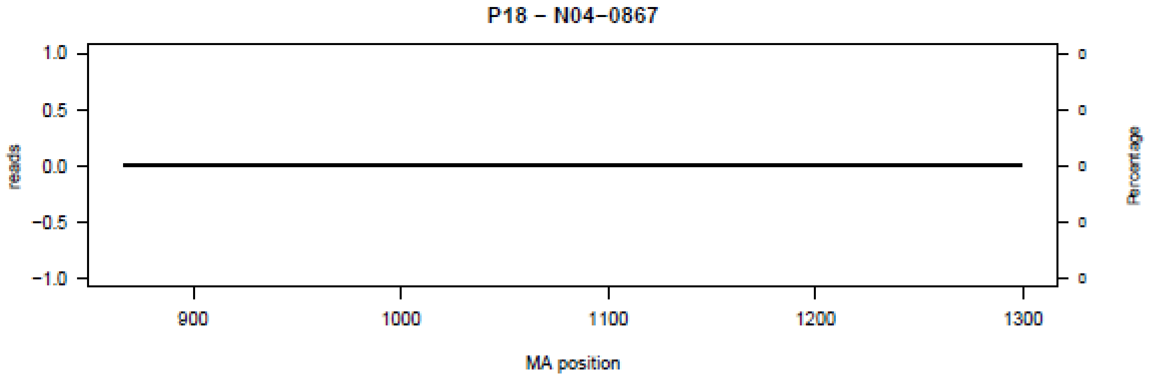
P18 – N04–0867
reads
Percentage
MA position
1.0
0.5
0.0
−0.5
−1.0
900
1000
1100
1200
1300
0
0
0
0
0

**Figure S5.** Bar plot of deletions in amplicon N05 in the 18 patients (P01-P18) at the nucleotide level.

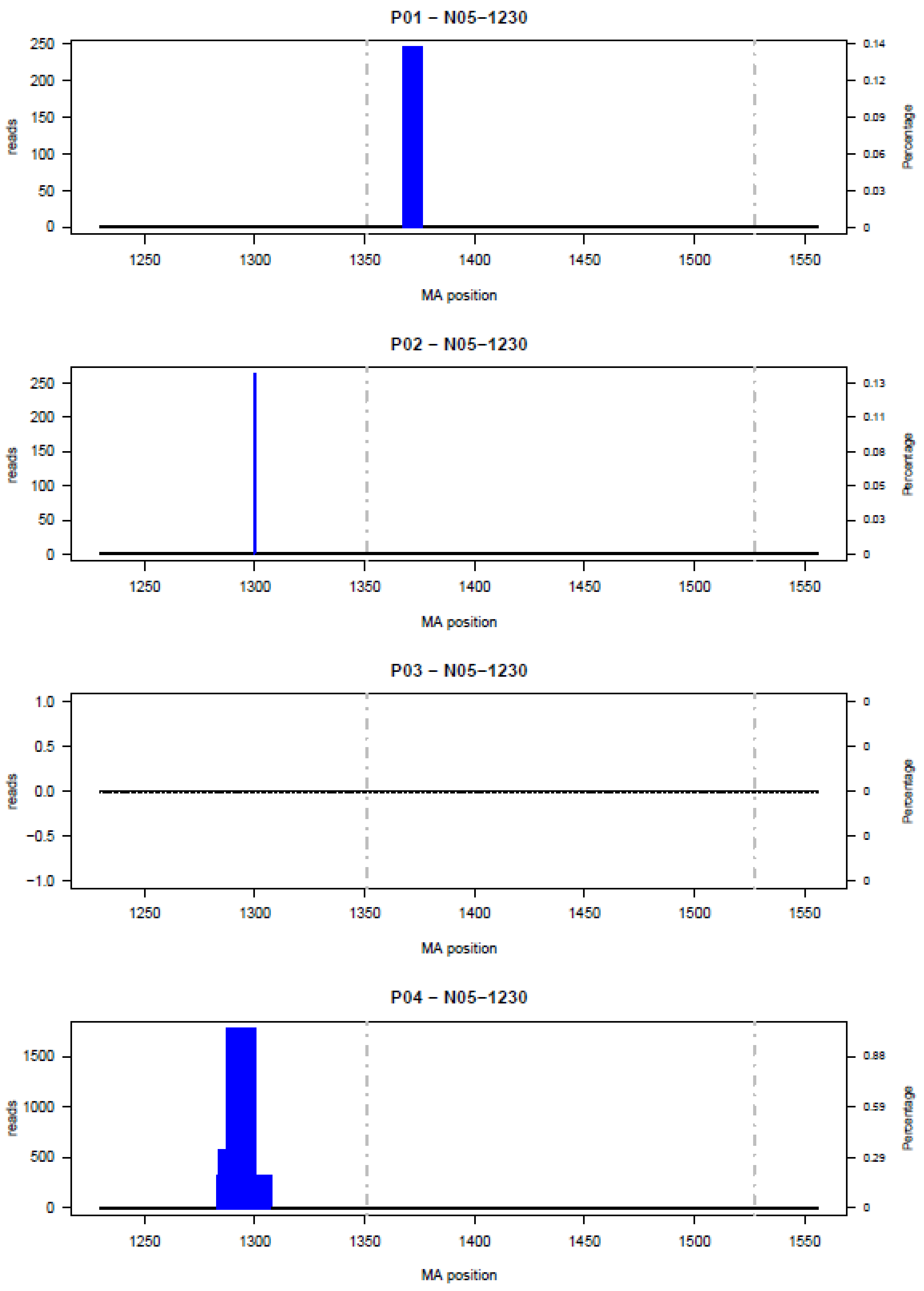

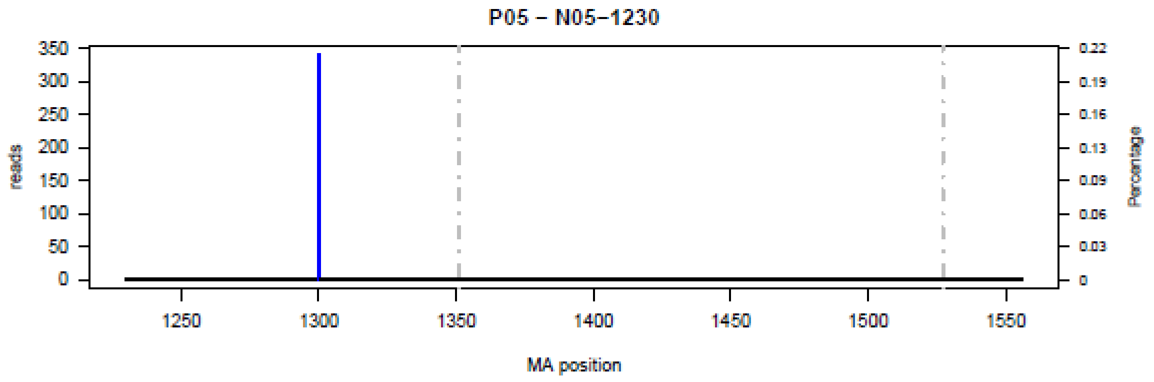
P05 – N05–1230
reads
Percentage
MA position

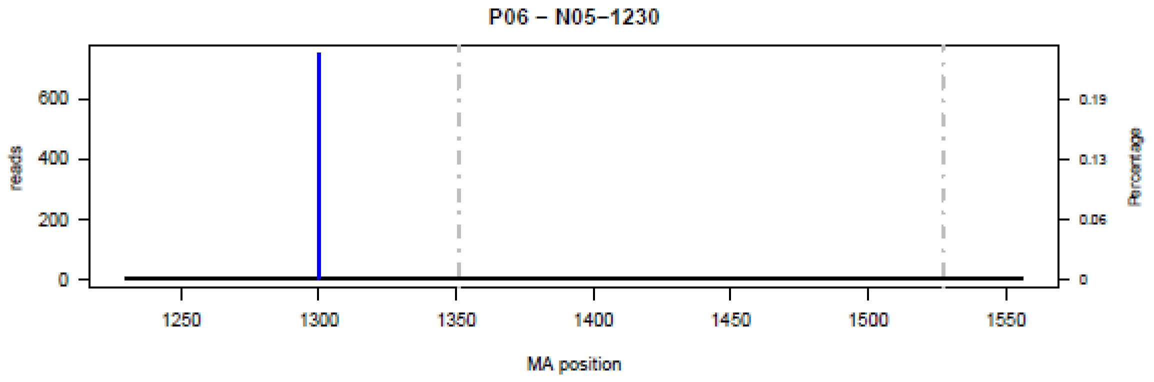
P06 – N05–1230
reads
Percentage
MA position

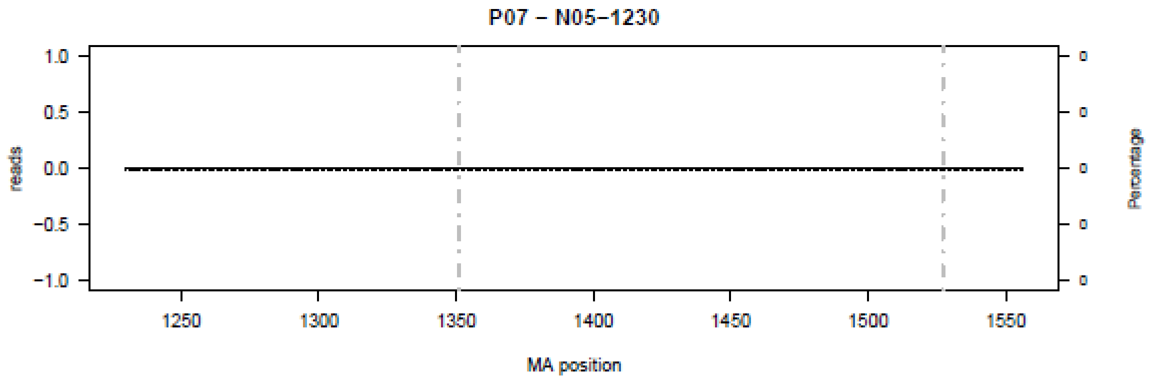
P07 – N05–1230
reads
Percentage
MA position

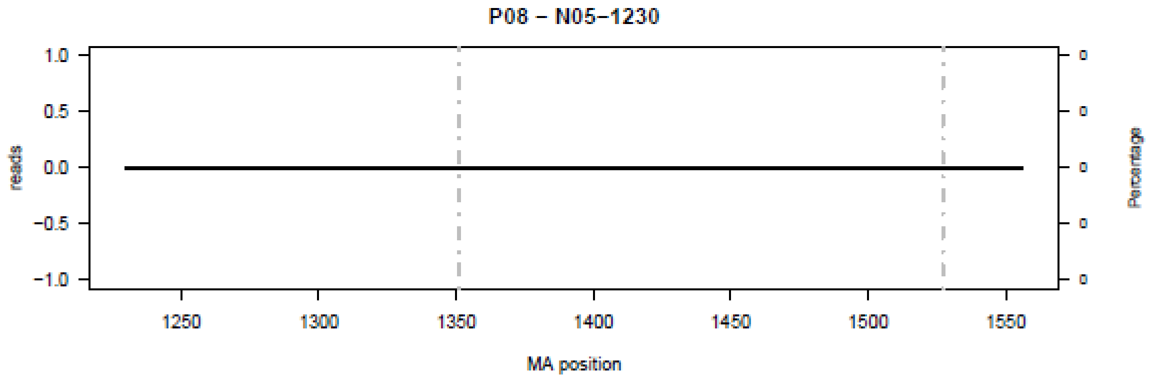
P08 – N05–1230
reads
Percentage
MA position

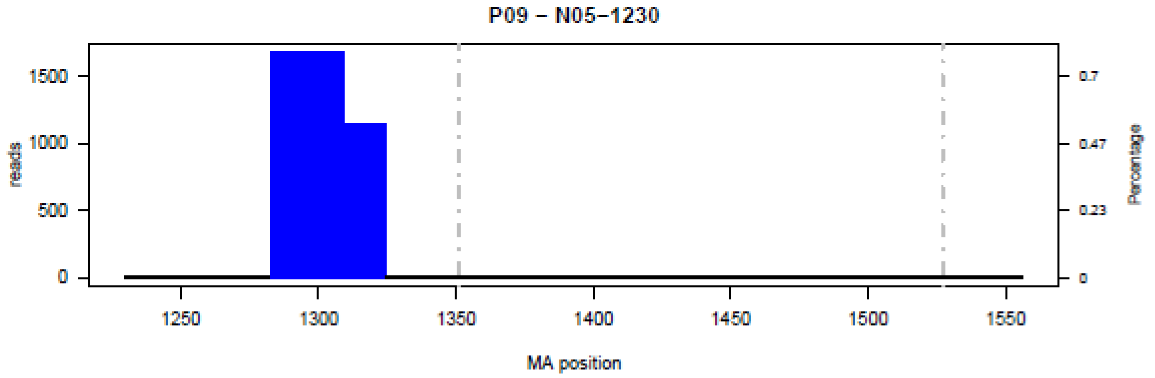
P09 – N05–1230
reads
Percentage
MA position
1500
1000
500
0
0.7
0.47
0.23
0
1250
1300
1350
1400
1450
1500
1550

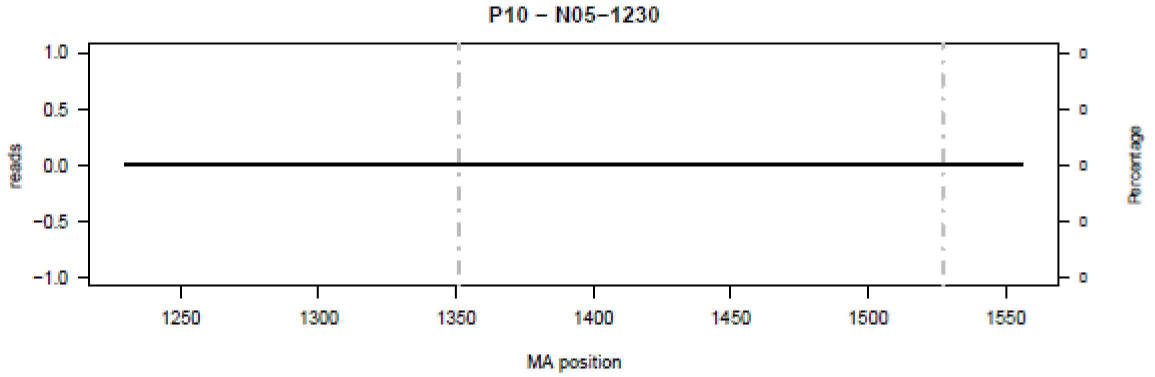
P10 – N05–1230
reads
Percentage
MA position

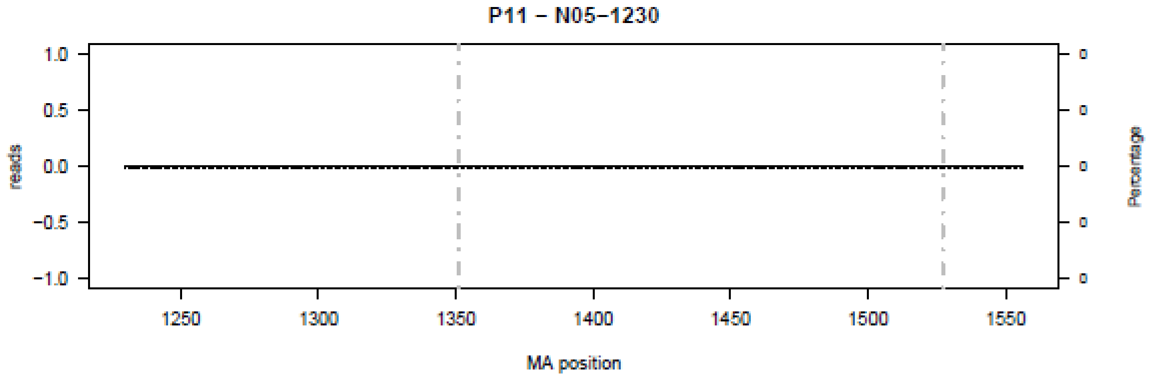
P11 – N05–1230
reads
Percentage
MA position

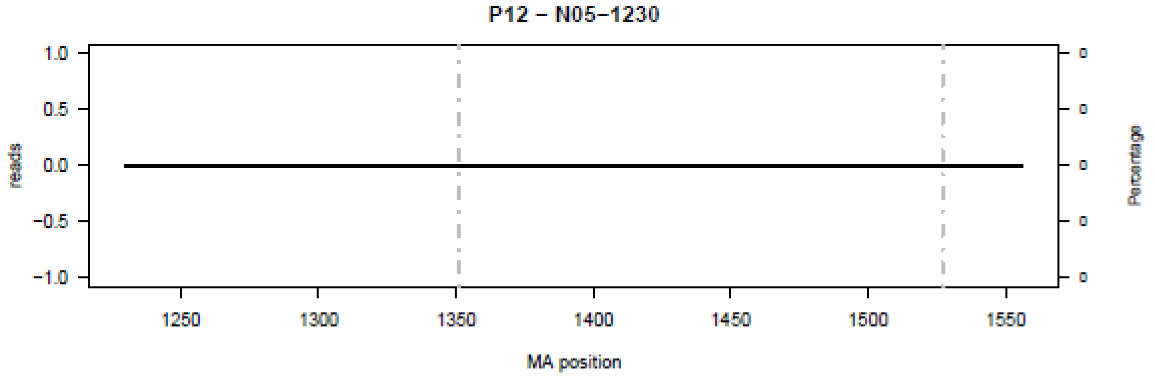
P12 – N05–1230
reads
Percentage
MA position

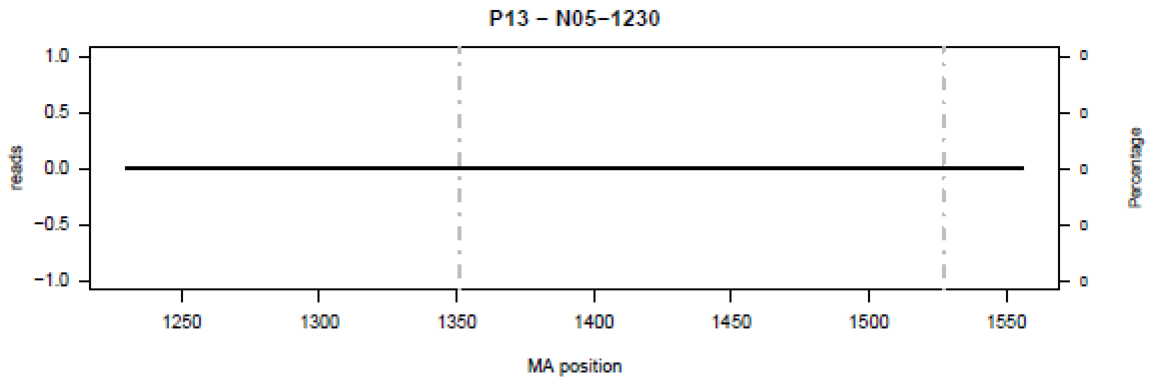
P13 – N05−1230
reads
Percentage
MA position

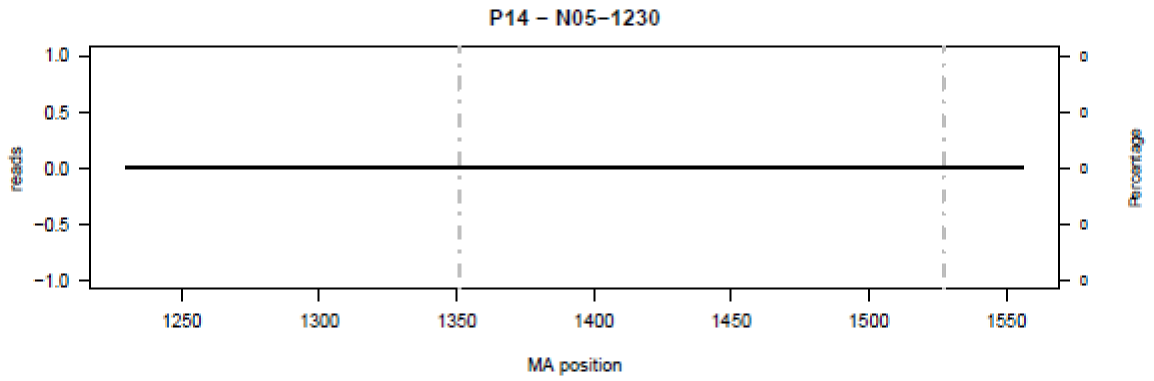
P14 – N05−1230
reads
Percentage
MA position

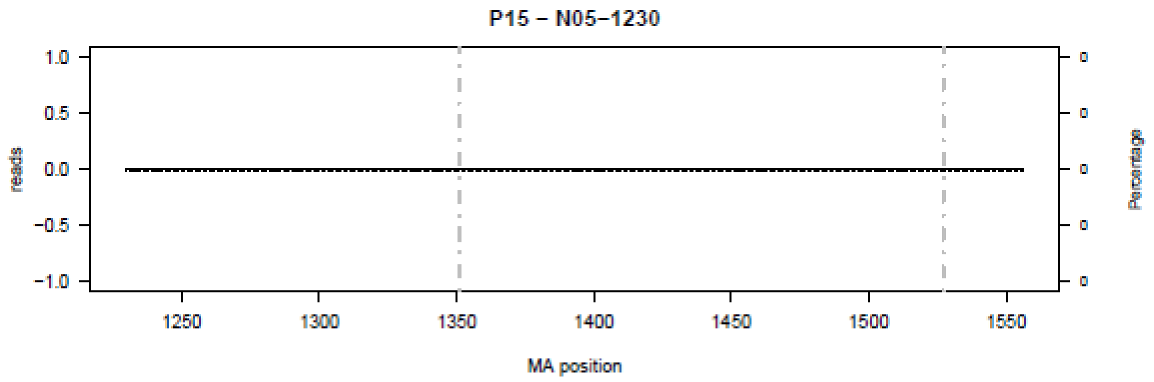
P15 – N05−1230
reads
Percentage
MA position

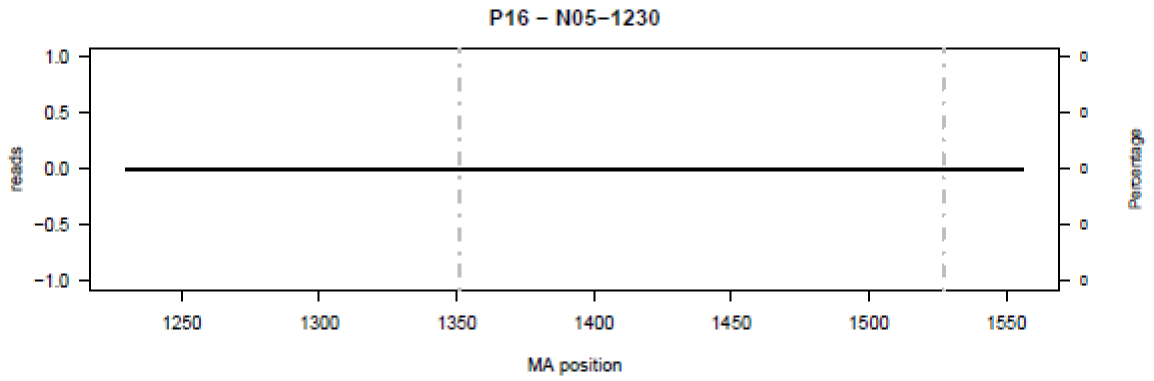
P16 – N05−1230
reads
Percentage
MA position

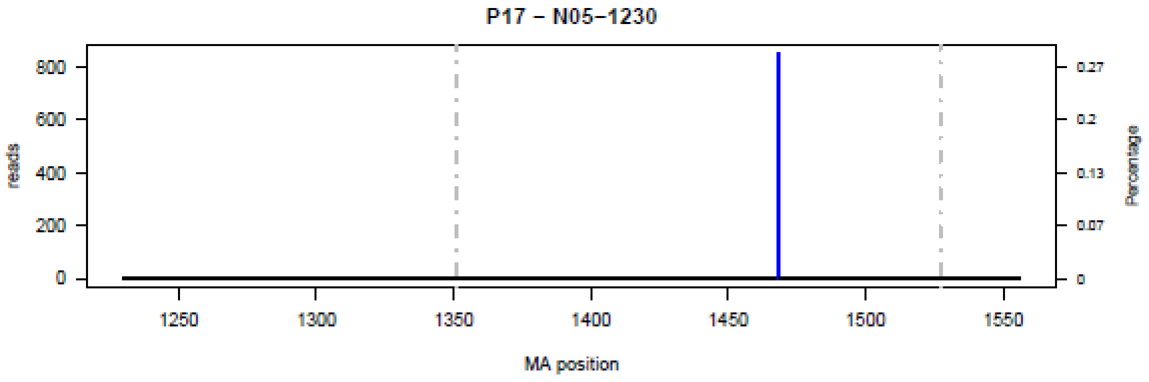

P17 − N05−1230
reads
800
600
400
200
0
0.27
0.2
0.13
0.07
0
Percentage
1250
1300
1350
1400
1450
1500
1550
MA position


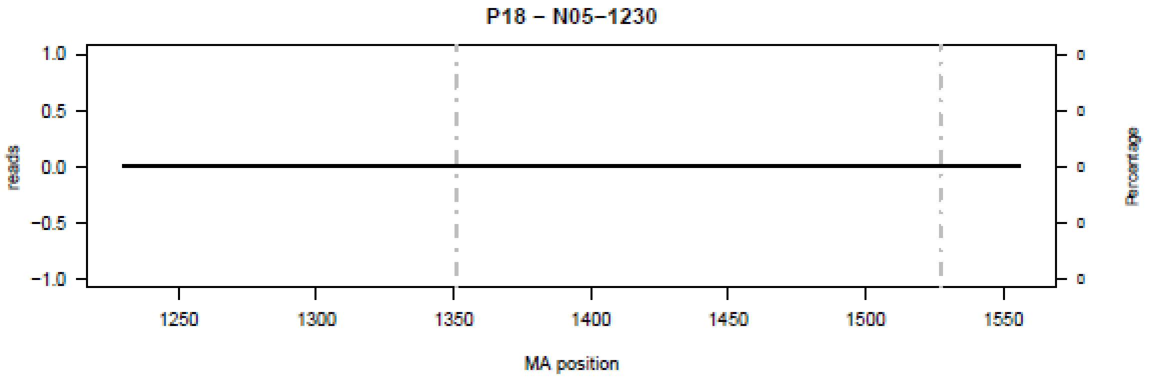

P18 − N05−1230
reads
1.0
0.5
0.0
−0.5
−1.0
Percentage
1250
1300
1350
1400
1450
1500
1550
MA position

**Figure S6.** Bar plot of deletions in amplicon N06 in the 18 patients (P01-P18) at the nucleotide level.

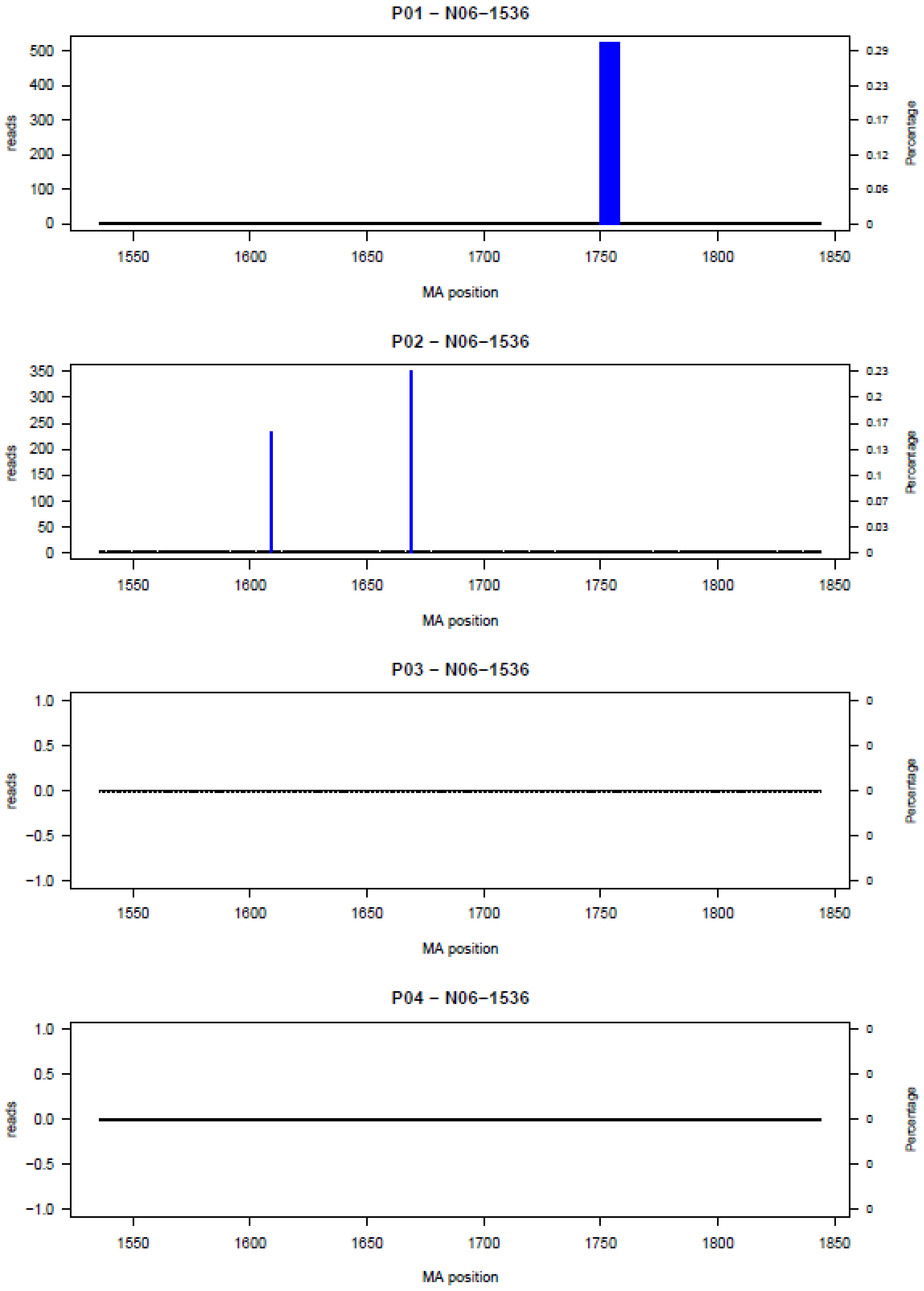

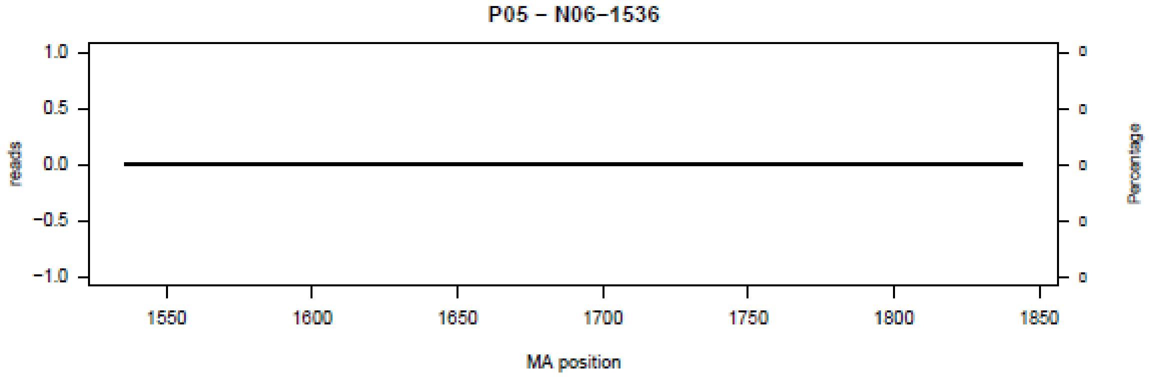

P05 – N06–1536
reads
Percentage
MA position


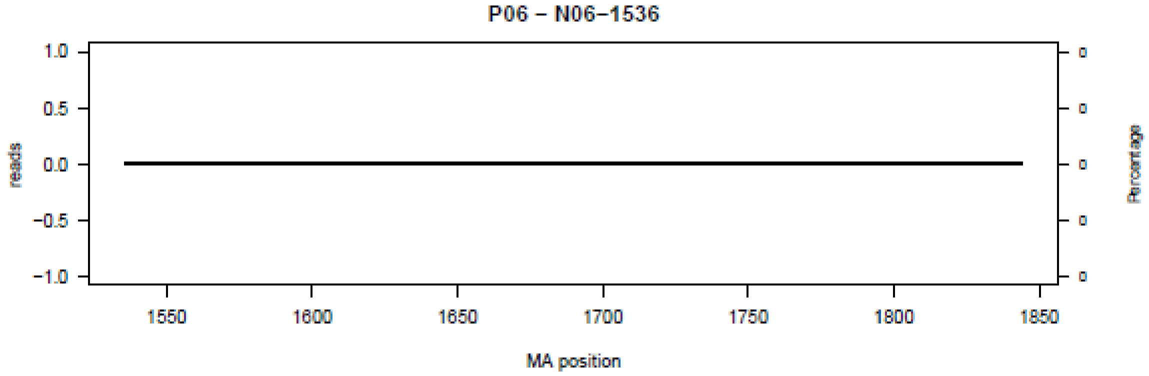

P06 – N06–1536
reads
Percentage
MA position


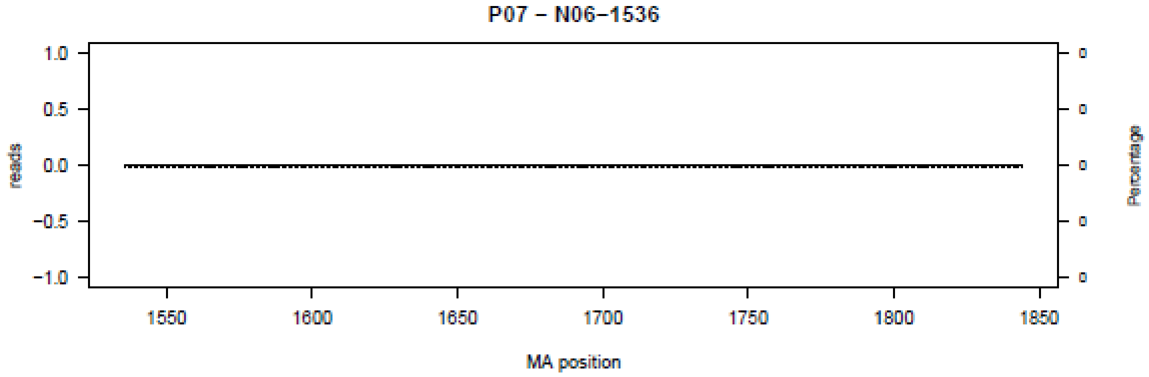

P07 – N06–1536
reads
Percentage
MA position


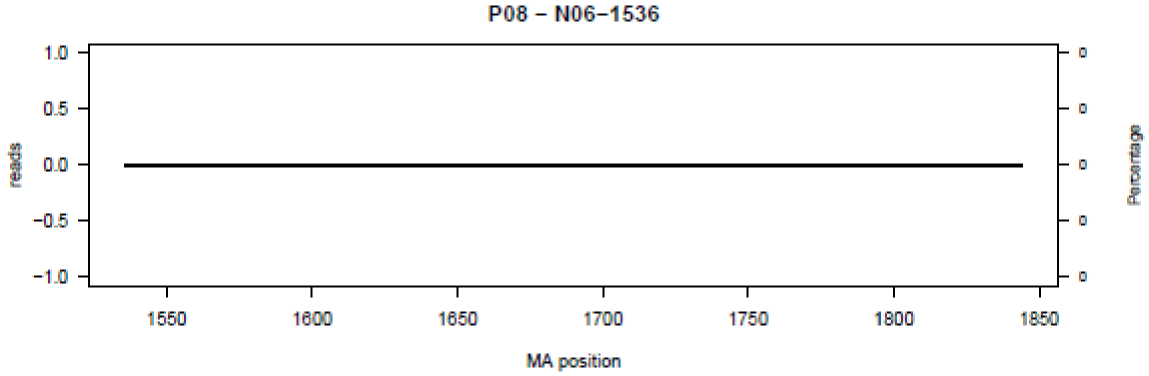

P08 – N06–1536
reads
Percentage
MA position

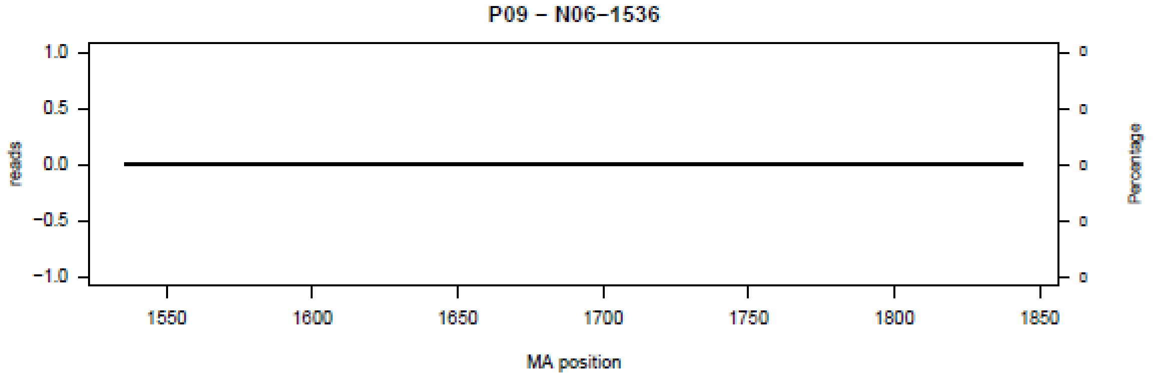
P09 – N06–1536
reads
Percentage
MA position

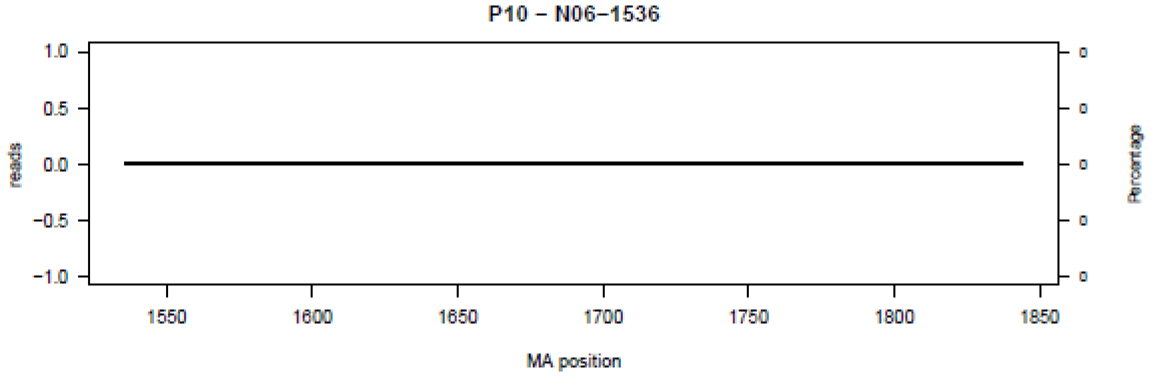
P10 – N06–1536
reads
Percentage
MA position

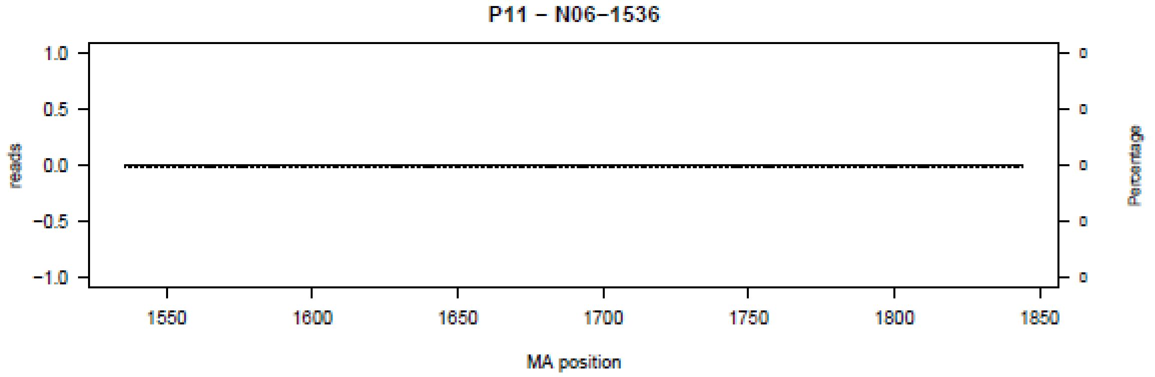
P11 – N06–1536
reads
Percentage
MA position

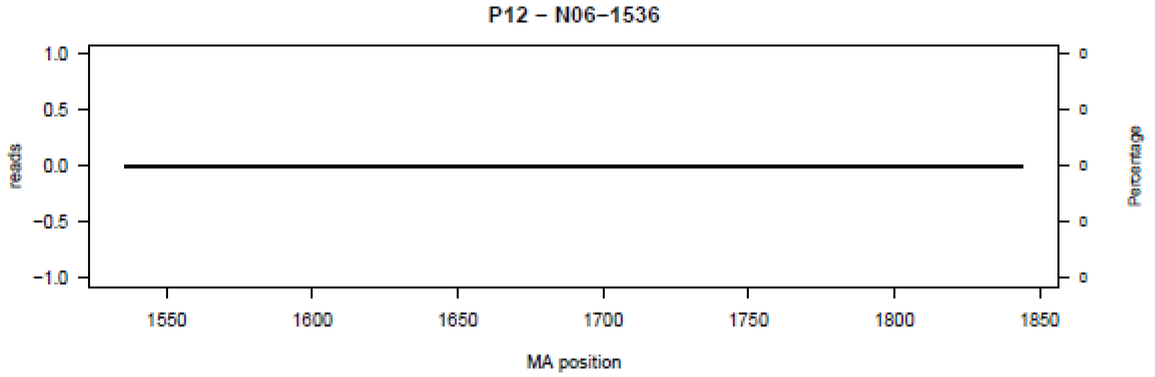
P12 – N06–1536
reads
Percentage
MA position

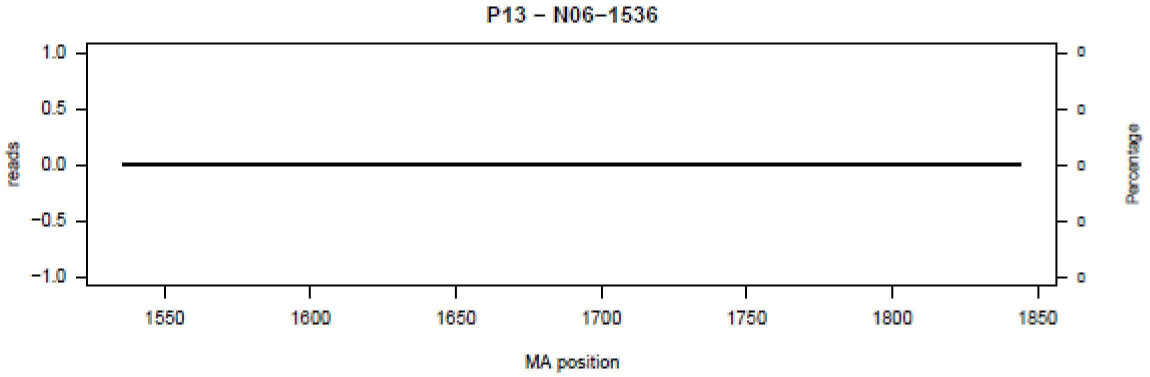
P13 – N06–1536
reads
Percentage
MA position

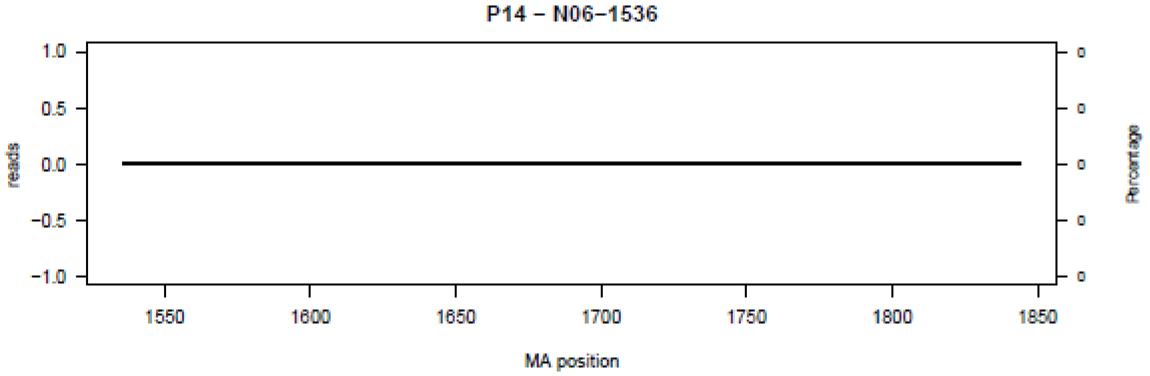
P14 – N06–1536
reads
Percentage
MA position

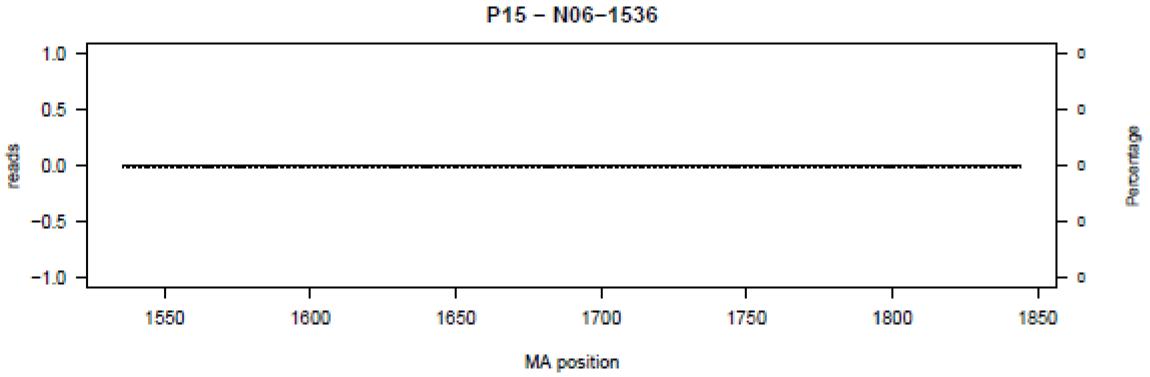
P15 – N06–1536
reads
Percentage
MA position

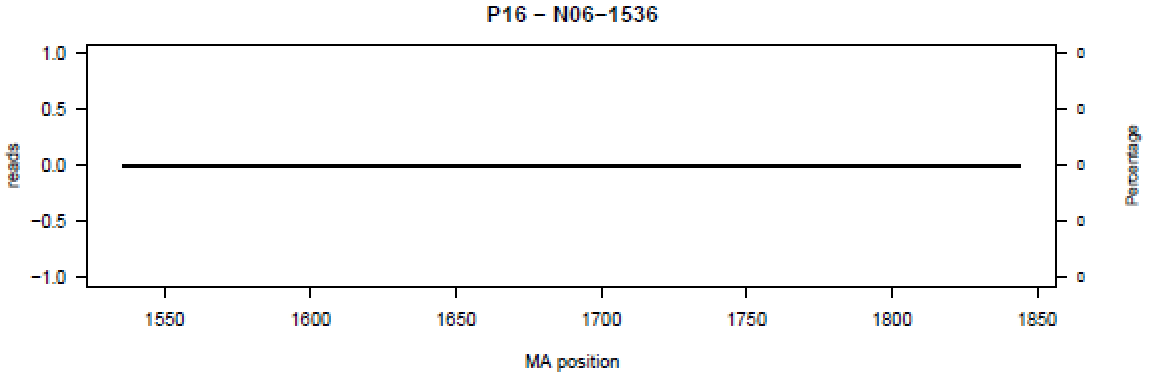
P16 – N06–1536
reads
Percentage
MA position

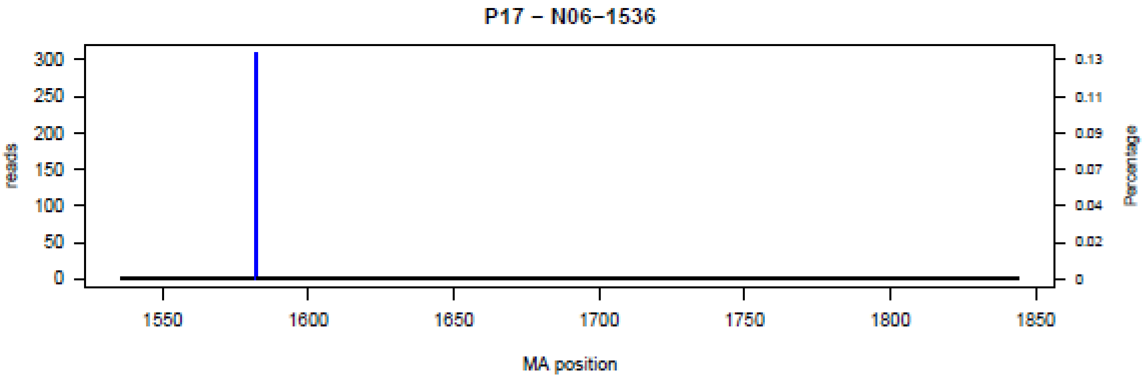
P17 – N06–1536
reads
Percentage
MA position
300
250
200
150
100
50
0
0.13
0.11
0.09
0.07
0.04
0.02
0
1550
1600
1650
1700
1750
1800
1850

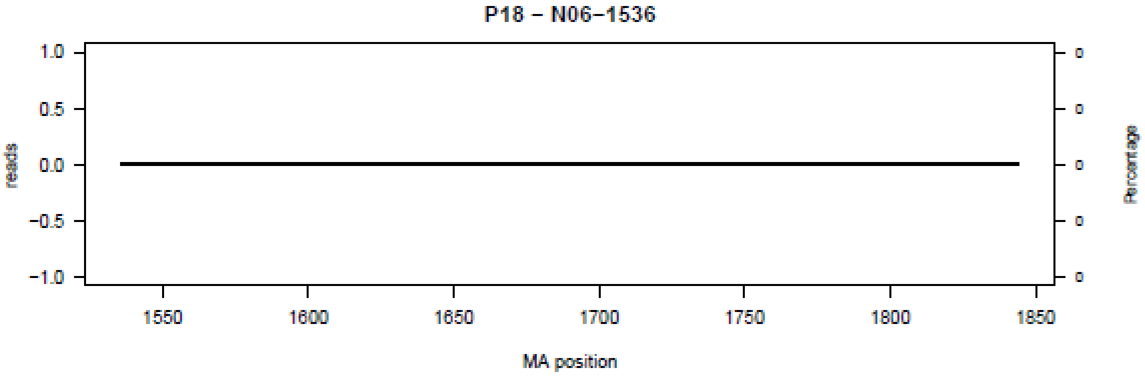
P18 – N06–1536
reads
Percentage
MA position
1.0
0.5
0.0
−0.5
−1.0
0
1550
1600
1650
1700
1750
1800
1850

**Figure S7.** Bar plot of deletions in amplicon N07 in the 18 patients (P01-P18) at the nucleotide level.

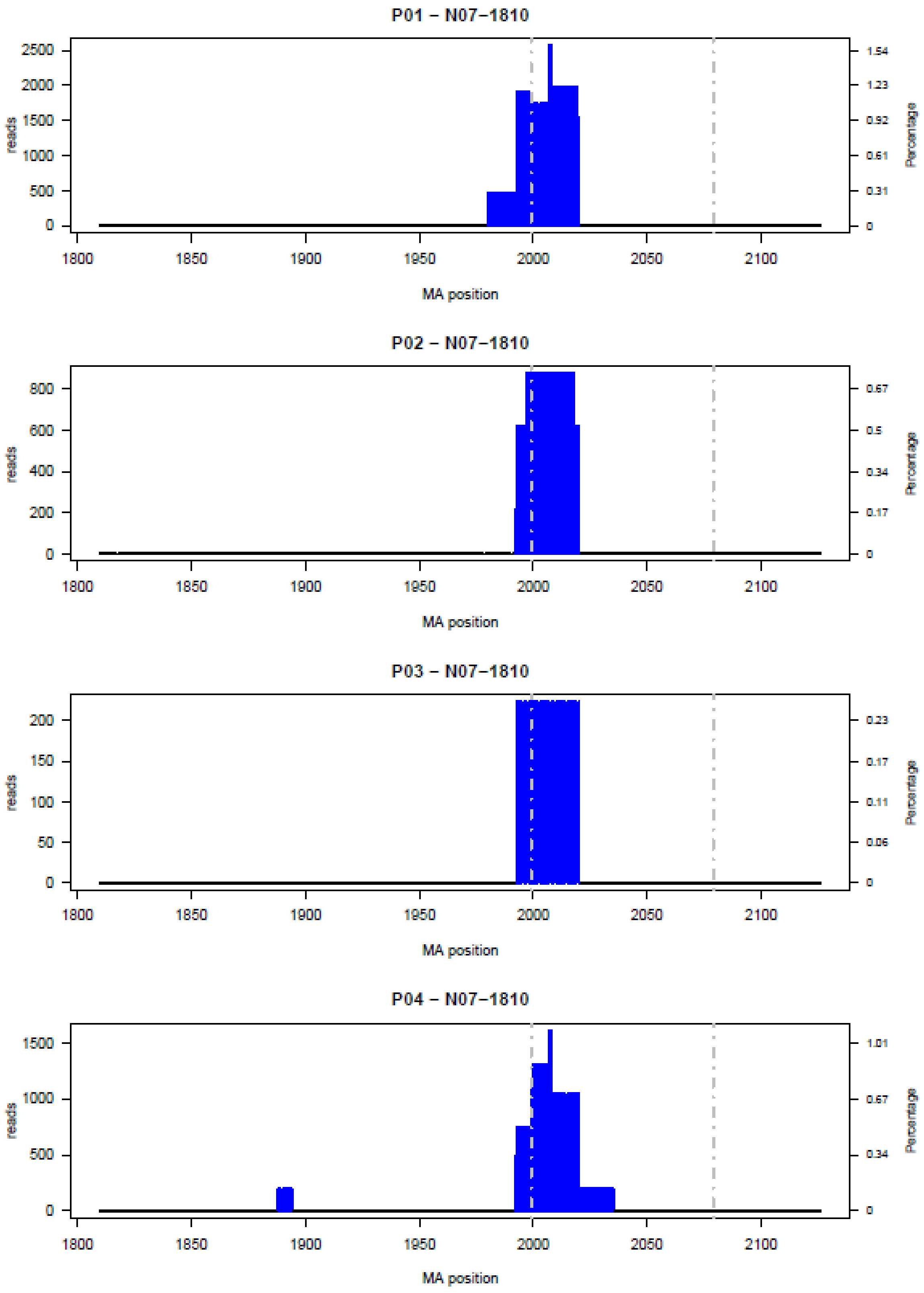

**P05 – N07–1810**

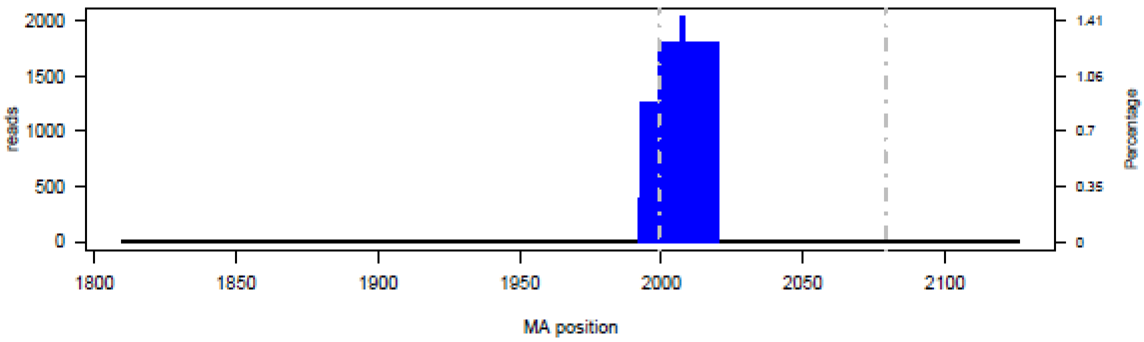


**P06 – N07–1810**

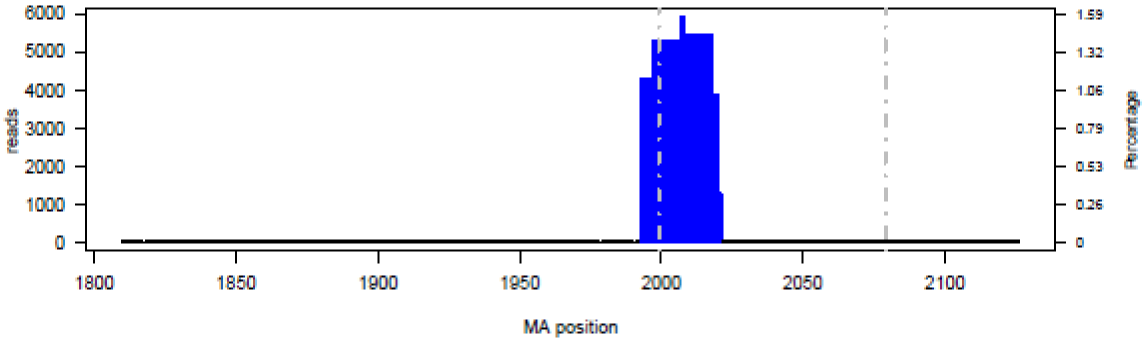


**P07 – N07–1810**

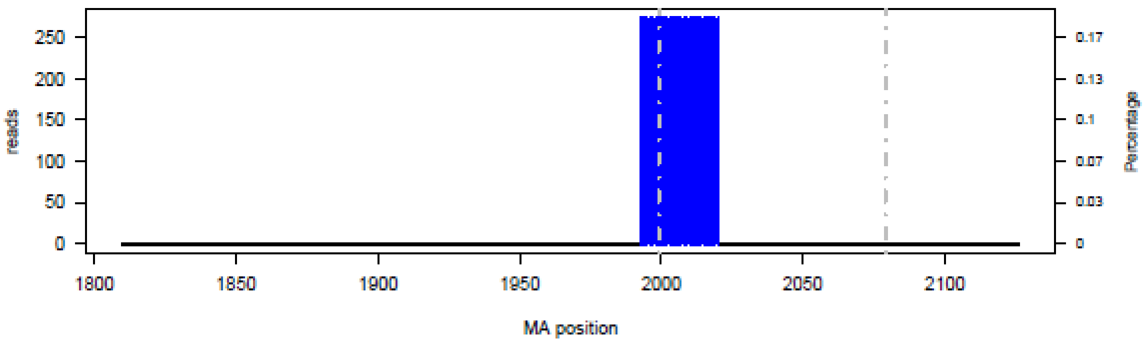


**P08 – N07–1810**

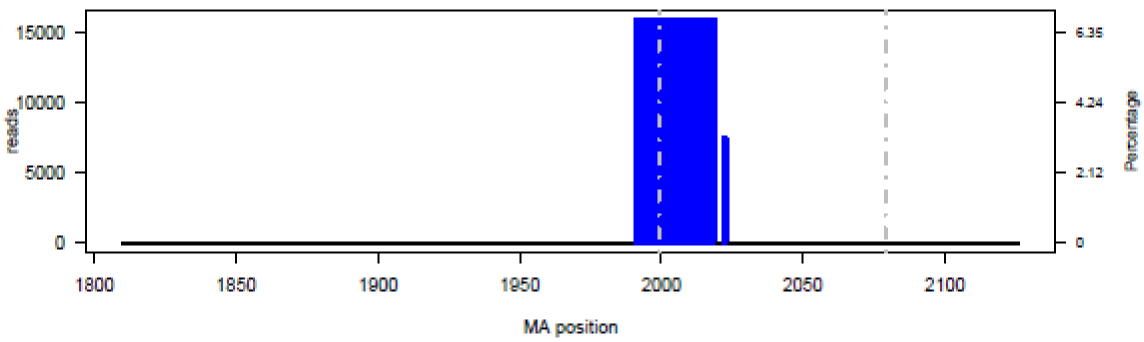

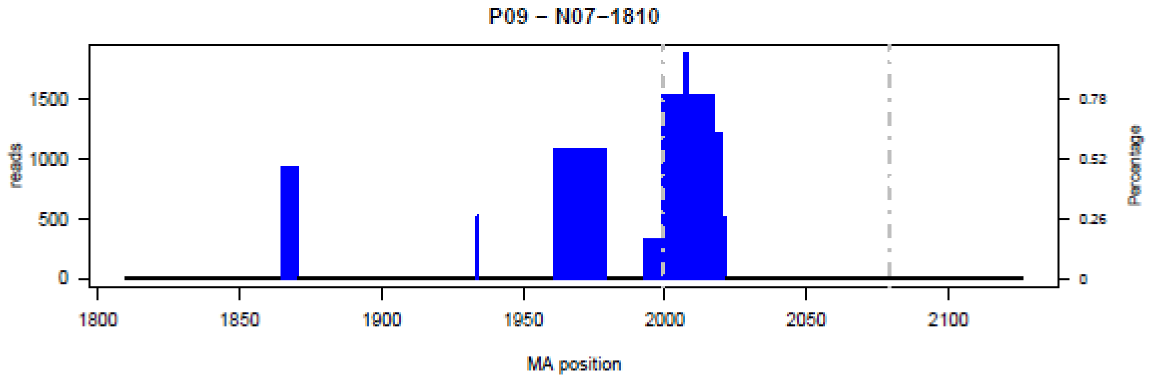
P09 – N07–1810
reads
Percentage
MA position
0
500
1000
1500
0
0.26
0.52
0.78
1800
1850
1900
1950
2000
2050
2100

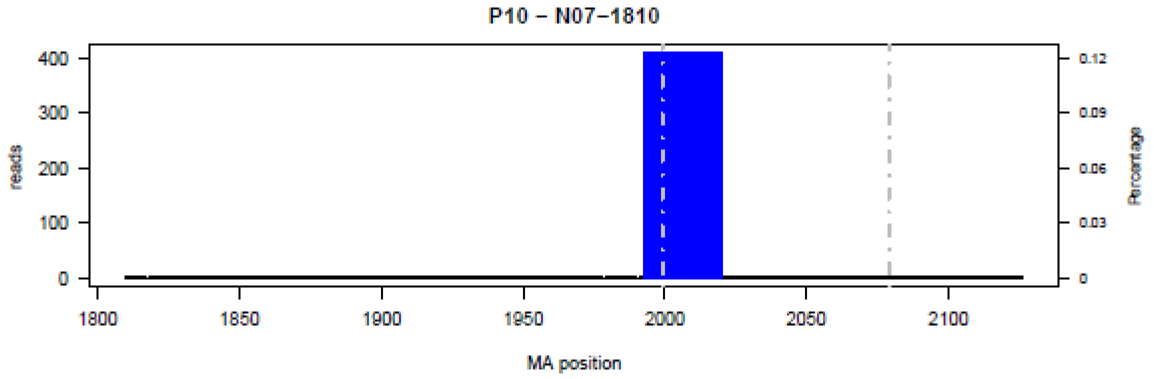
P10 – N07–1810
reads
Percentage
MA position
0
100
200
300
400
0
0.03
0.06
0.09
0.12
1800
1850
1900
1950
2000
2050
2100

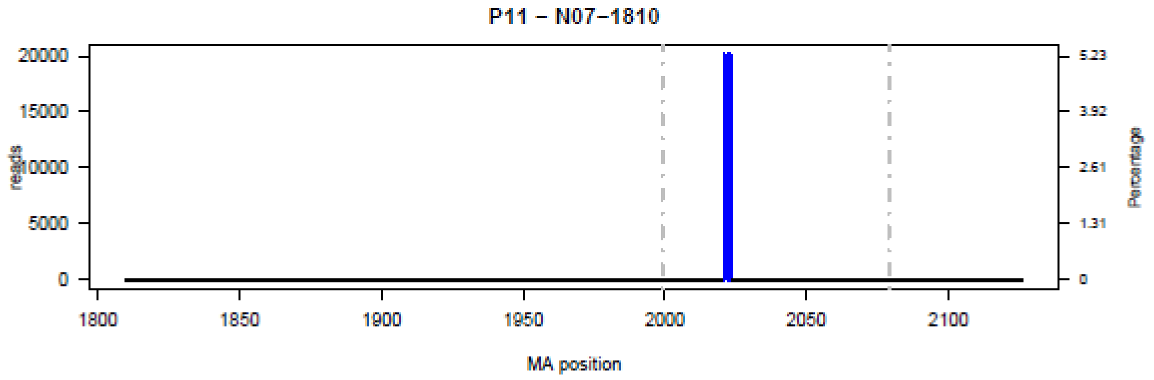
P11 – N07–1810
reads
Percentage
MA position
0
5000
10000
15000
20000
0
1.31
2.61
3.92
5.23
1800
1850
1900
1950
2000
2050
2100

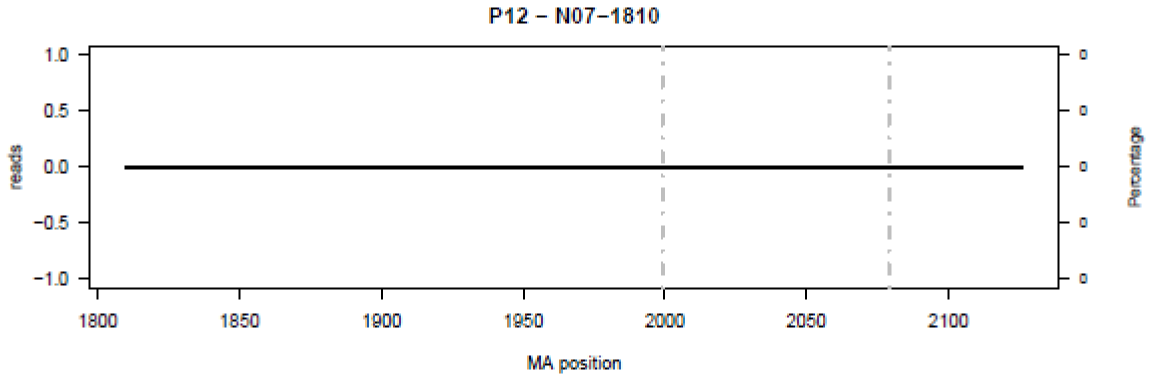
P12 – N07–1810
reads
Percentage
MA position
−1.0
−0.5
0.0
0.5
1.0
0
0
0
0
0
1800
1850
1900
1950
2000
2050
2100

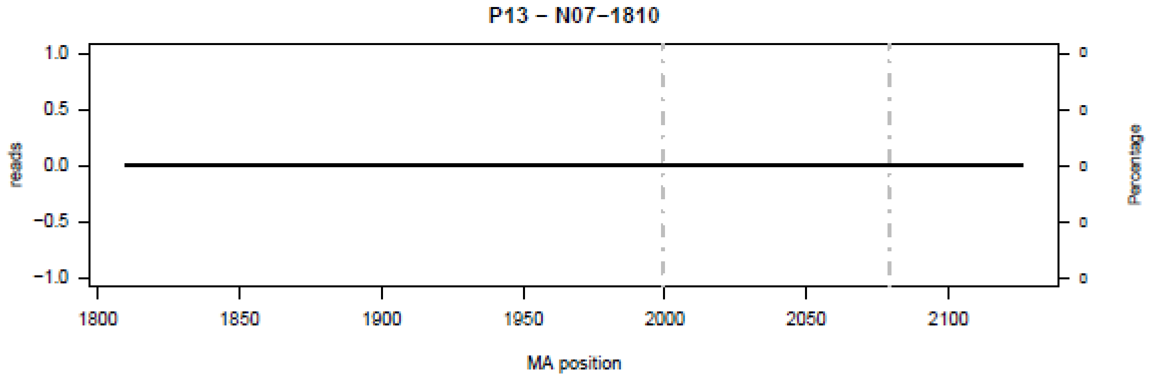

P13 – N07–1810
reads
Percentage
MA position


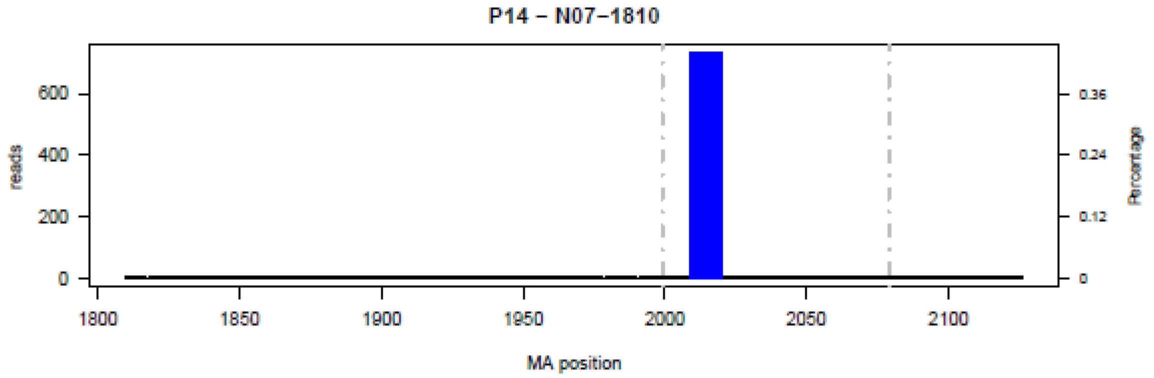

P14 – N07–1810
reads
Percentage
MA position


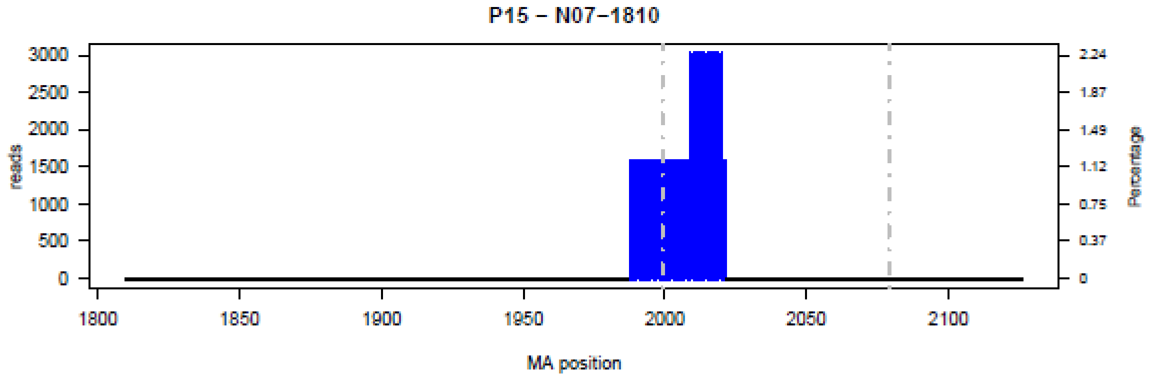

P15 – N07–1810
reads
Percentage
MA position


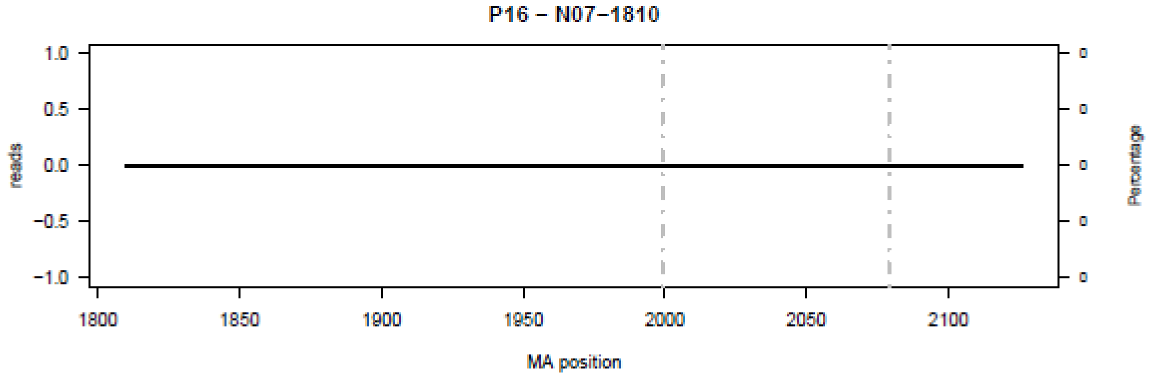

P16 – N07–1810
reads
Percentage
MA position

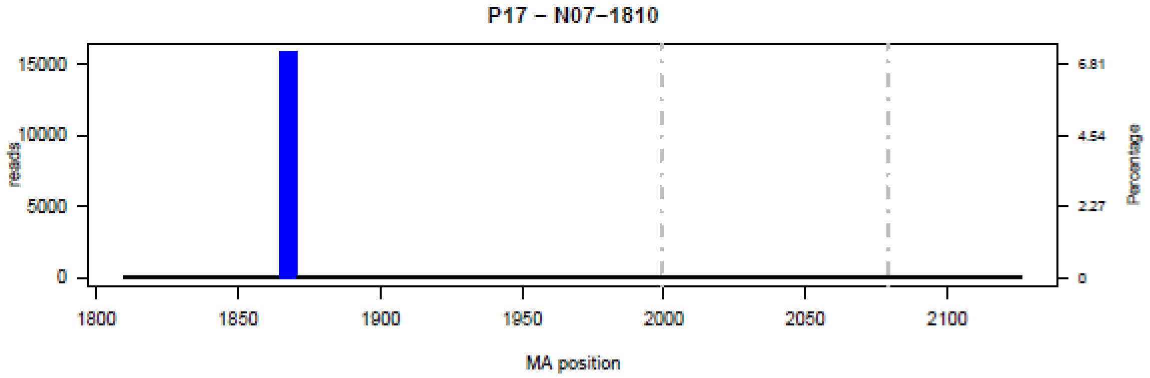
P17 – N07–1810
15000
10000
5000
0
reads
6.81
4.54
2.27
0
Percentage
1800
1850
1900
1950
2000
2050
2100
MA position

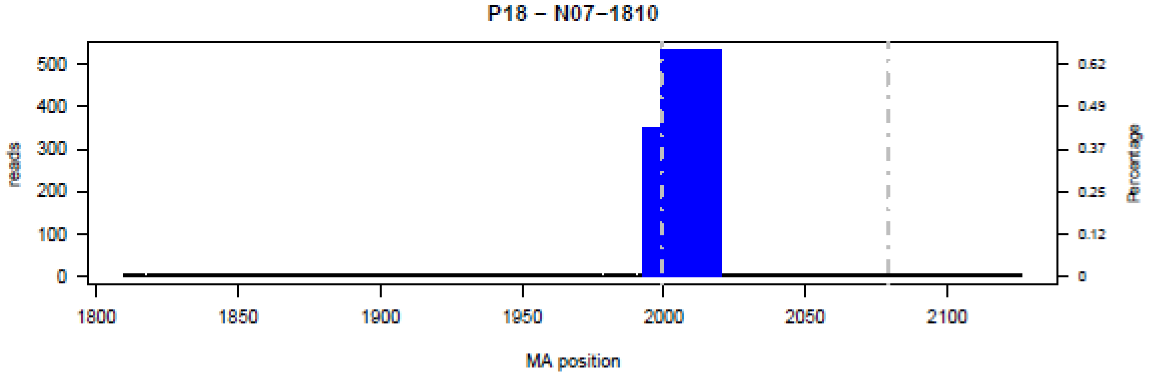
P18 – N07–1810
500
400
300
200
100
0
reads
0.62
0.49
0.37
0.25
0.12
0
Percentage
1800
1850
1900
1950
2000
2050
2100
MA position

**Figure S8.** Bar plot of deletions in amplicon N08 in the 18 patients (P01-P18) at the nucleotide level.

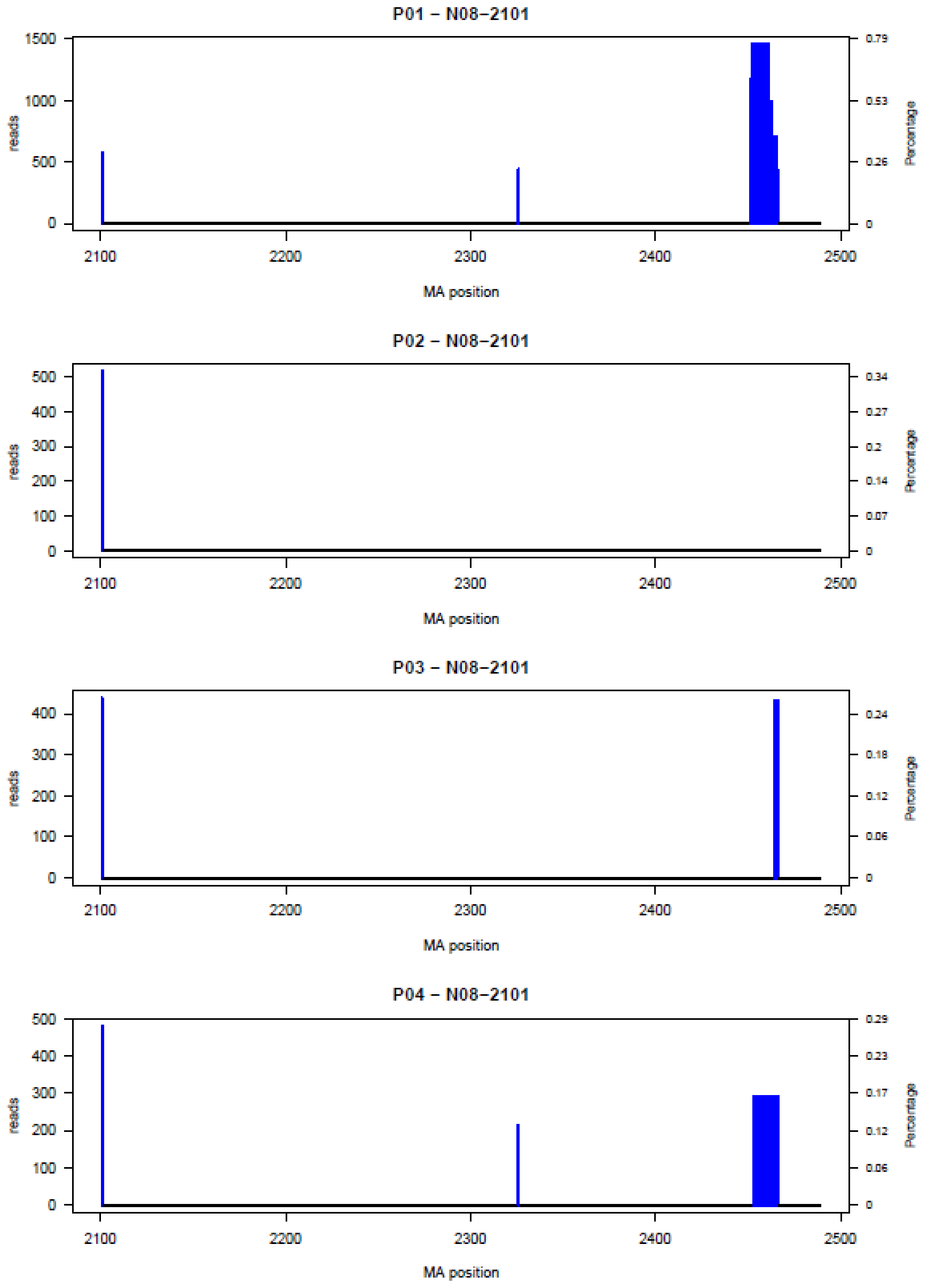

P05 – N08–2101

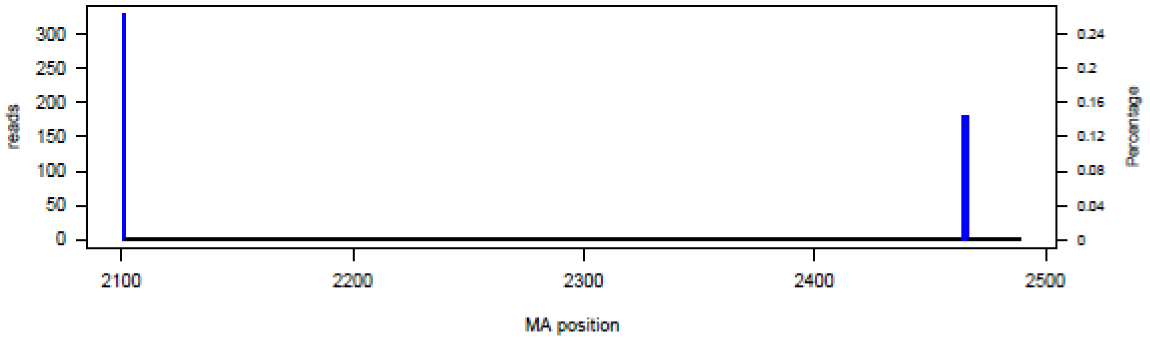


P06 – N08–2101

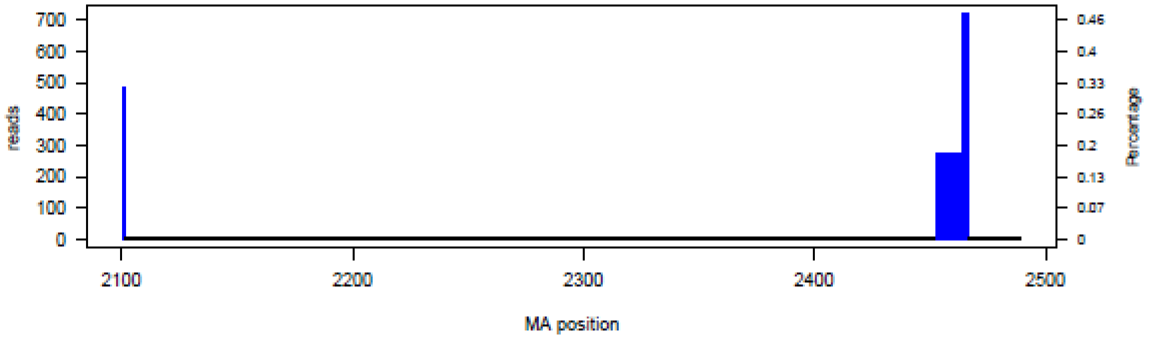


P07 – N08–2101

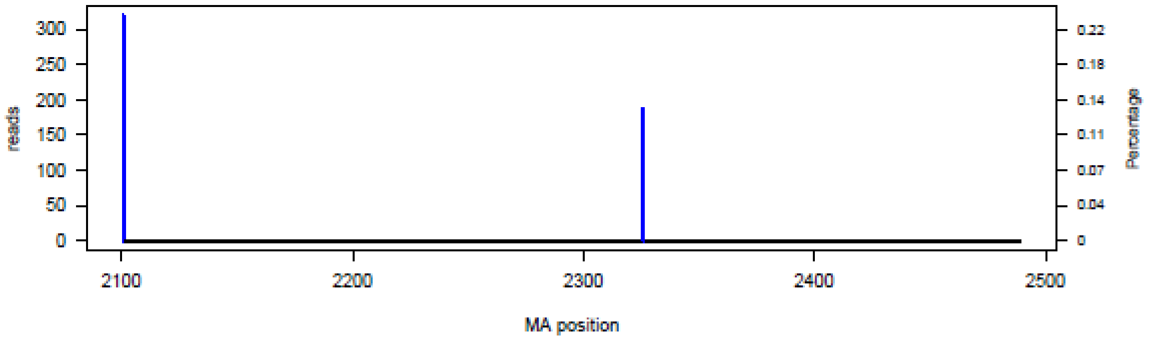


P08 – N08–2101

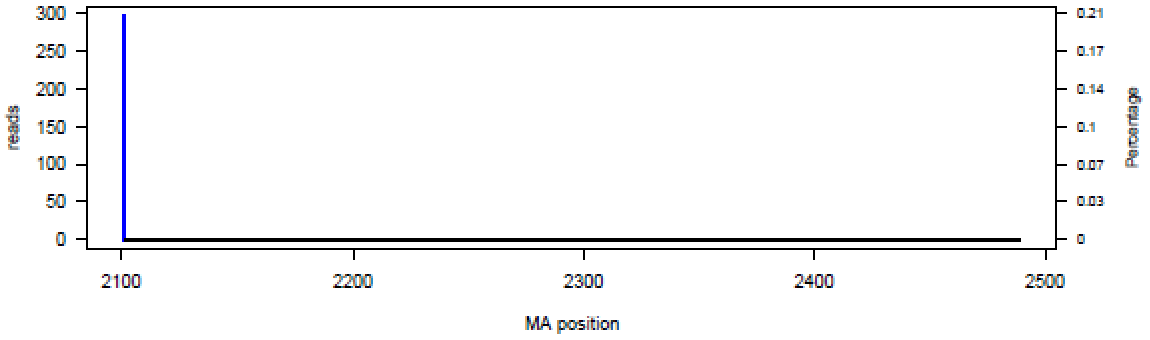

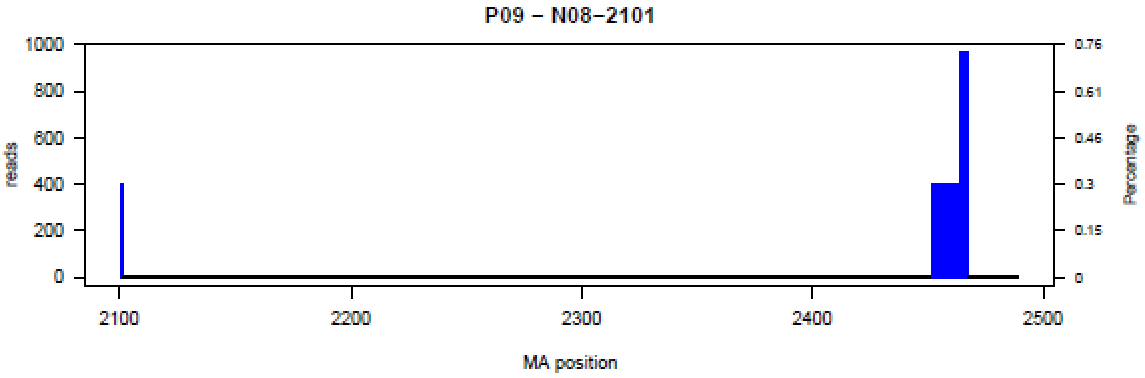

P09 – N08–2101
reads
Percentage
1000
800
600
400
200
0
0.76
0.61
0.46
0.3
0.15
0
2100
2200
2300
2400
2500
MA position


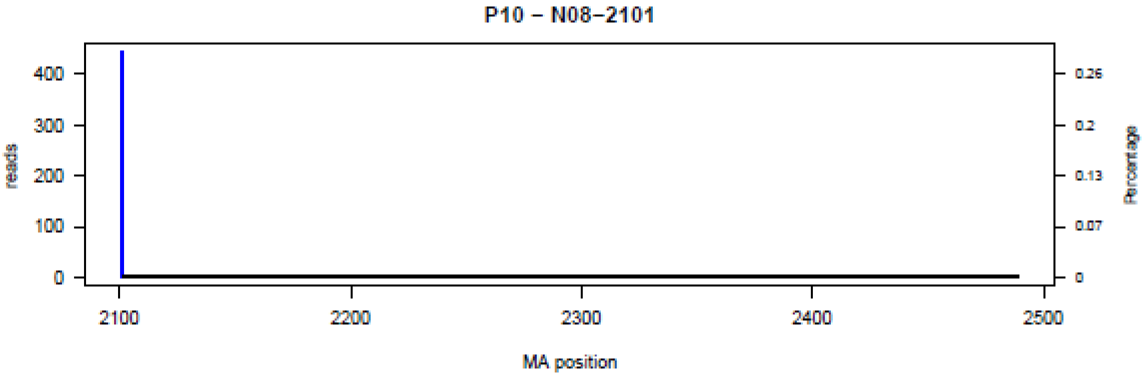

P10 – N08–2101
reads
Percentage
400
300
200
100
0
0.26
0.2
0.13
0.07
0
2100
2200
2300
2400
2500
MA position


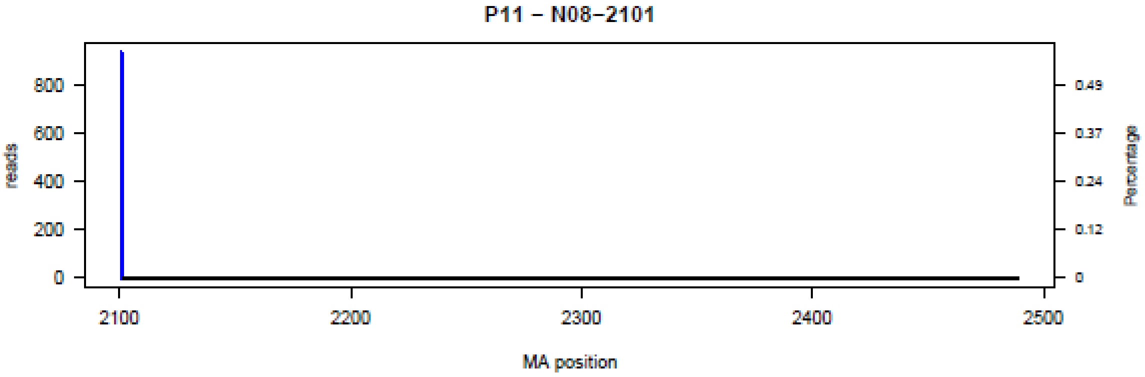

P11 – N08–2101
reads
Percentage
800
600
400
200
0
0.49
0.37
0.24
0.12
0
2100
2200
2300
2400
2500
MA position


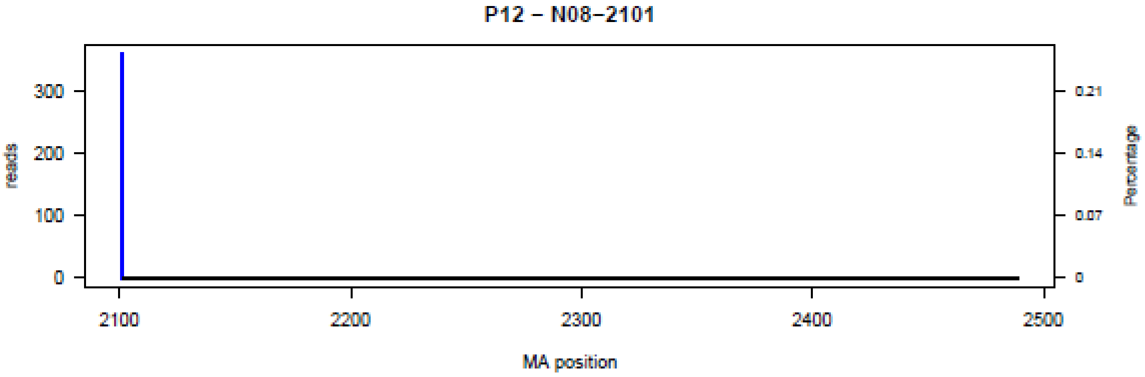

P12 – N08–2101
reads
Percentage
300
200
100
0
0.21
0.14
0.07
0
2100
2200
2300
2400
2500
MA position

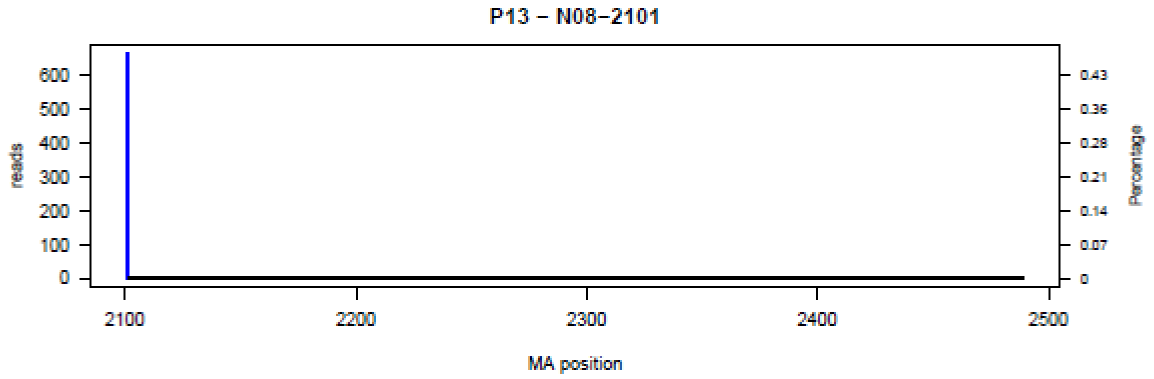
P13 – N08–2101
reads
Percentage
0
100
200
300
400
500
600
0
0.07
0.14
0.21
0.28
0.36
0.43
2100
2200
2300
2400
2500
MA position

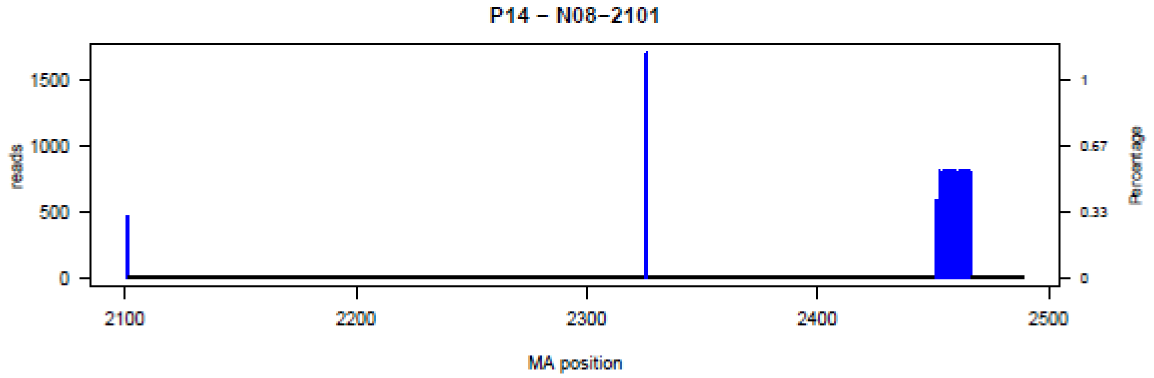
P14 – N08–2101
reads
Percentage
0
500
1000
1500
0
0.33
0.67
1
2100
2200
2300
2400
2500
MA position

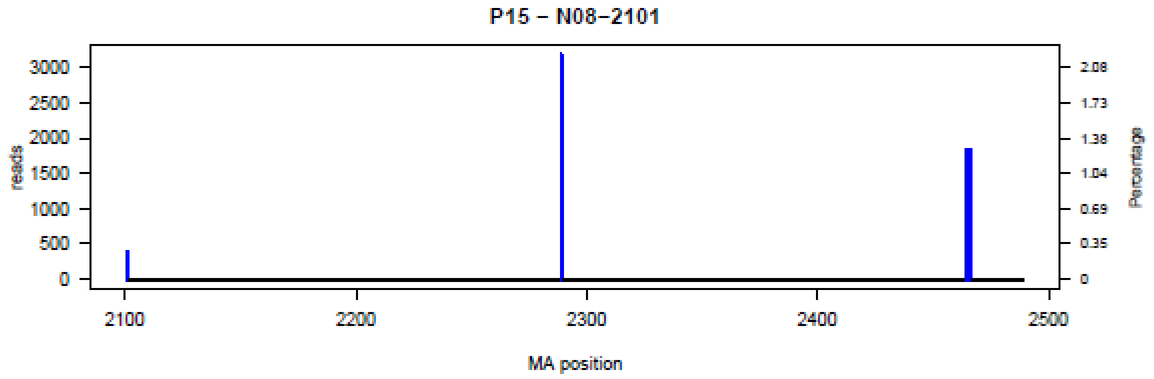
P15 – N08–2101
reads
Percentage
0
500
1000
1500
2000
2500
3000
0
0.35
0.69
1.04
1.38
1.73
2.08
2100
2200
2300
2400
2500
MA position

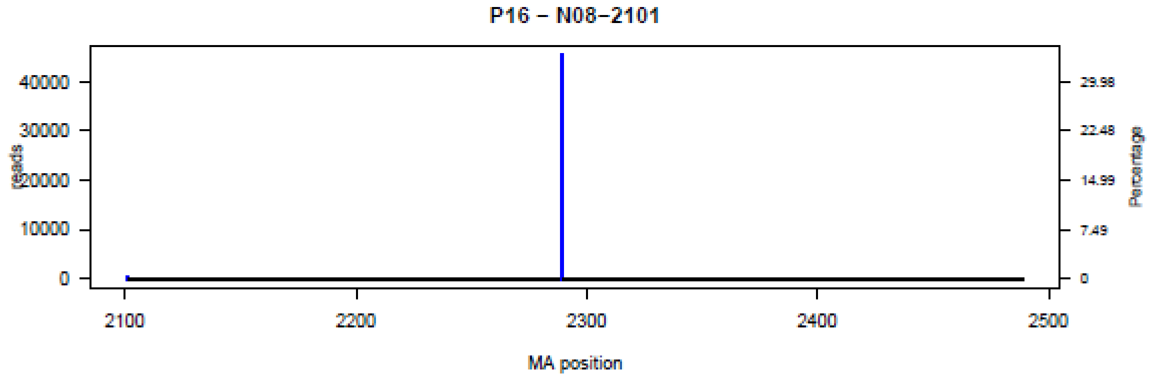
P16 – N08–2101
reads
Percentage
0
10000
20000
30000
40000
0
7.49
14.99
22.48
29.98
2100
2200
2300
2400
2500
MA position

**P17 – N08–2101**

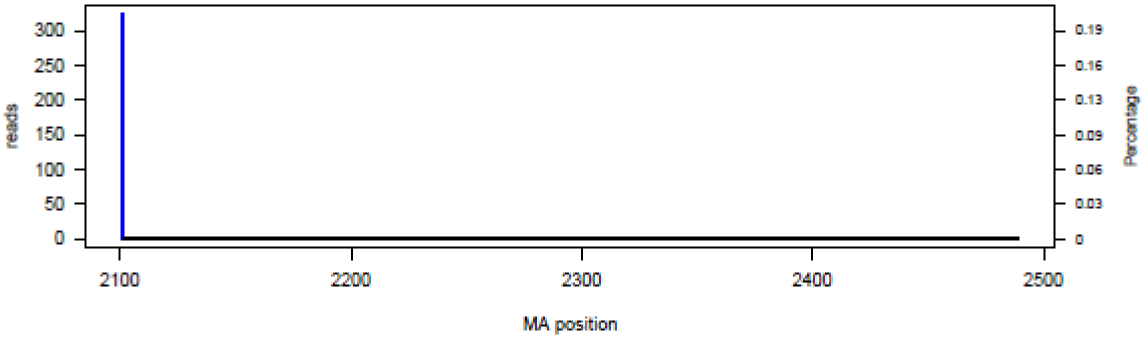


**P18 – N08–2101**

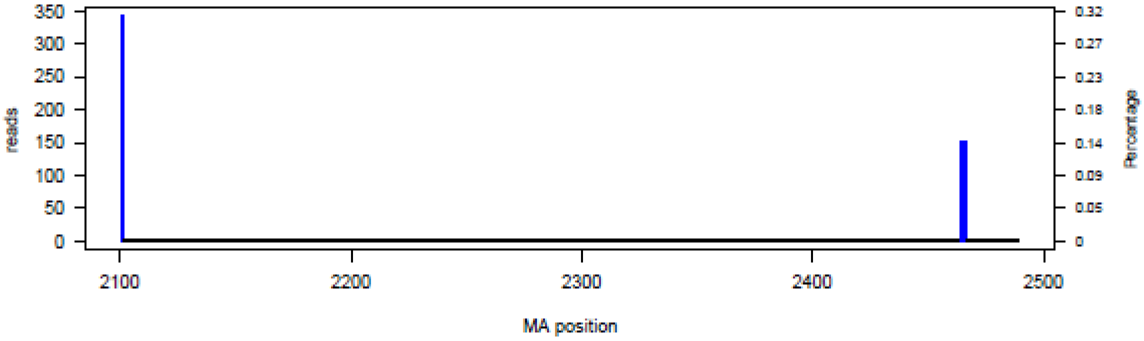

**Figure S9.** Bar plot of deletions in amplicon N09 in the 18 patients (P01-P18) at the nucleotide level.

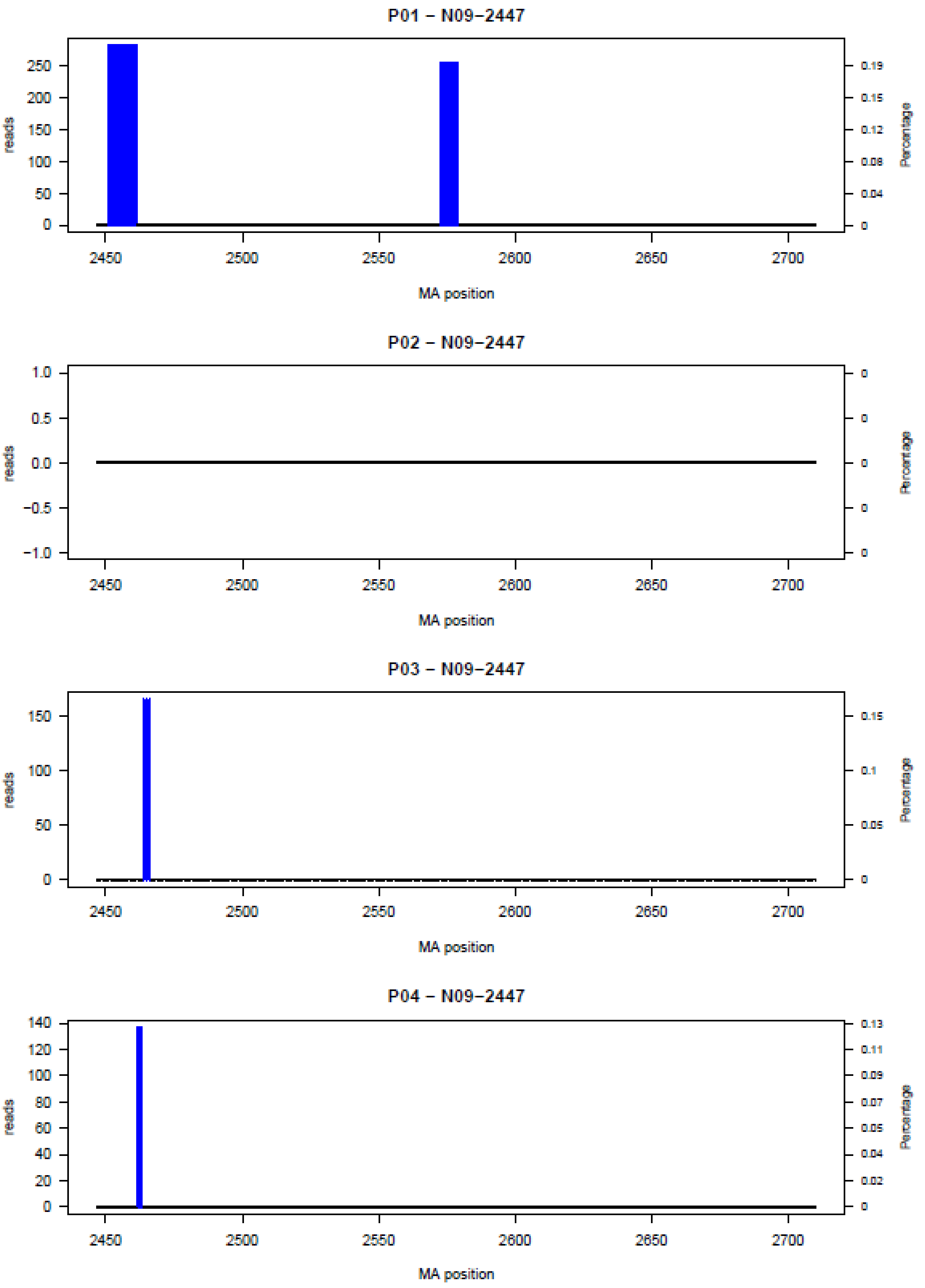

P05 − N09−2447

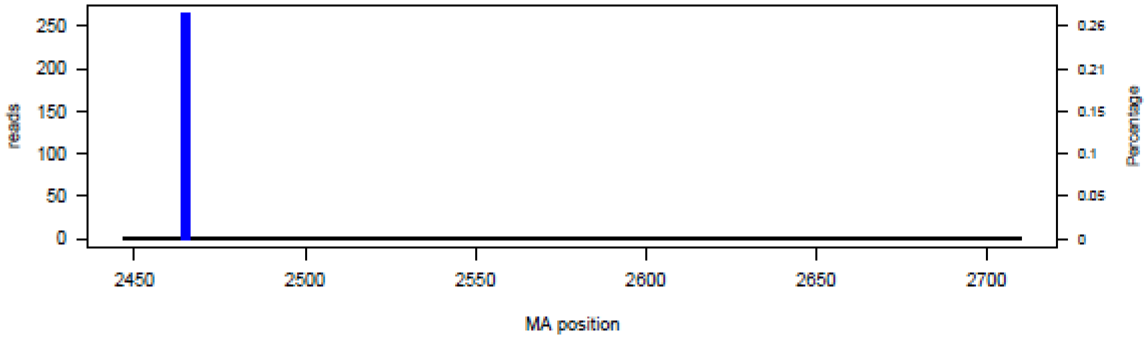


P06 − N09−2447

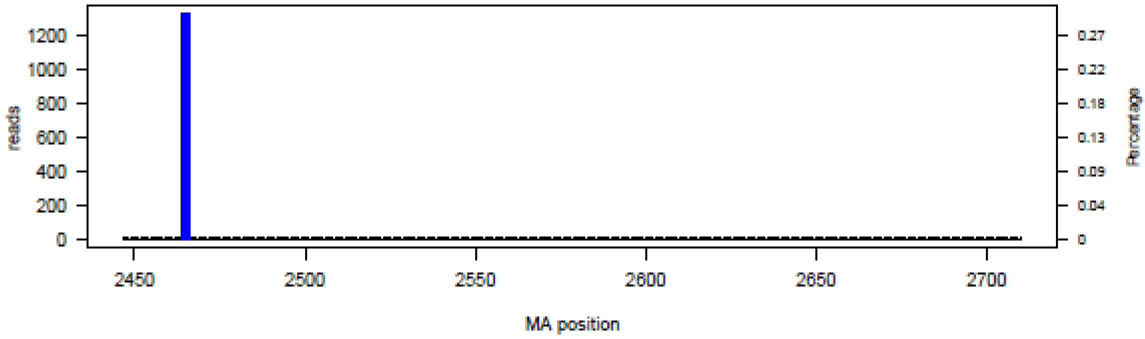


P07 − N09−2447

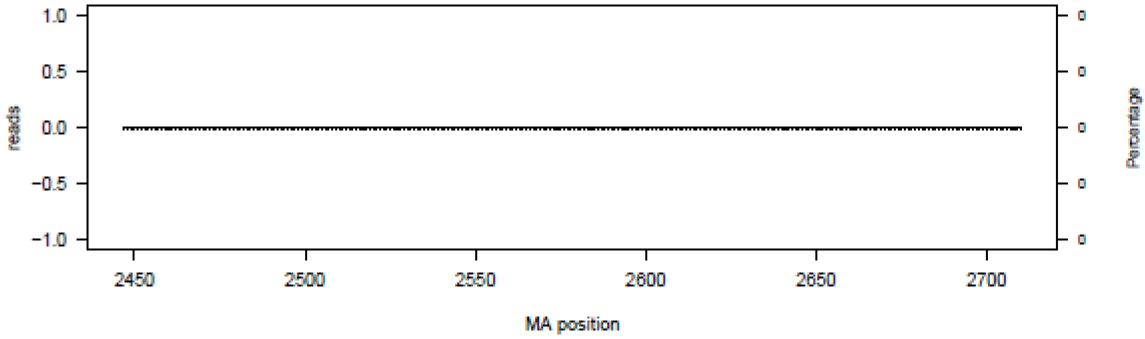


P08 − N09−2447

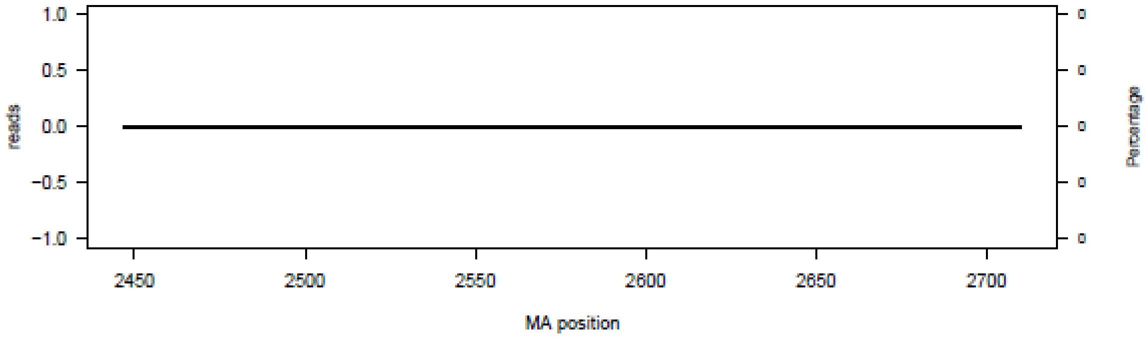

## P09 – N09–2447

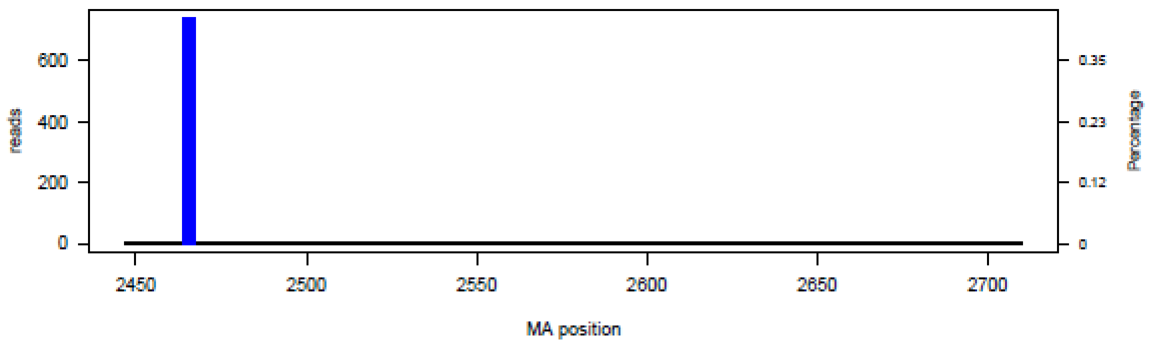


## P10 – N09–2447

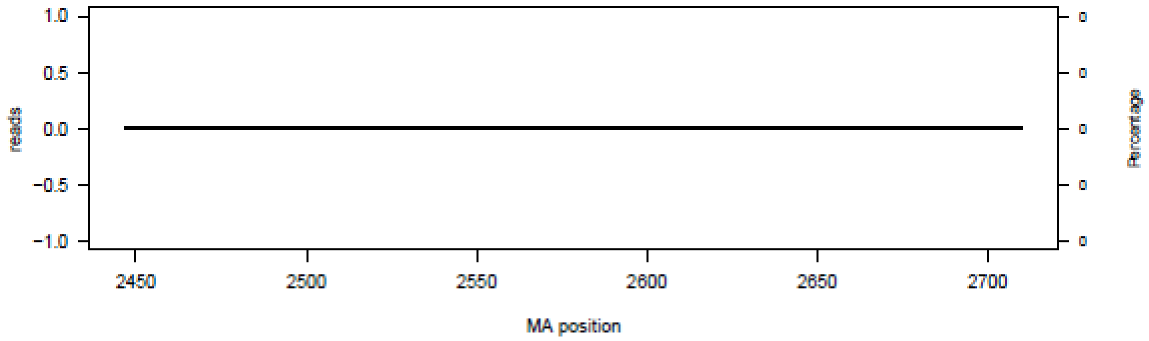


## P11 – N09–2447

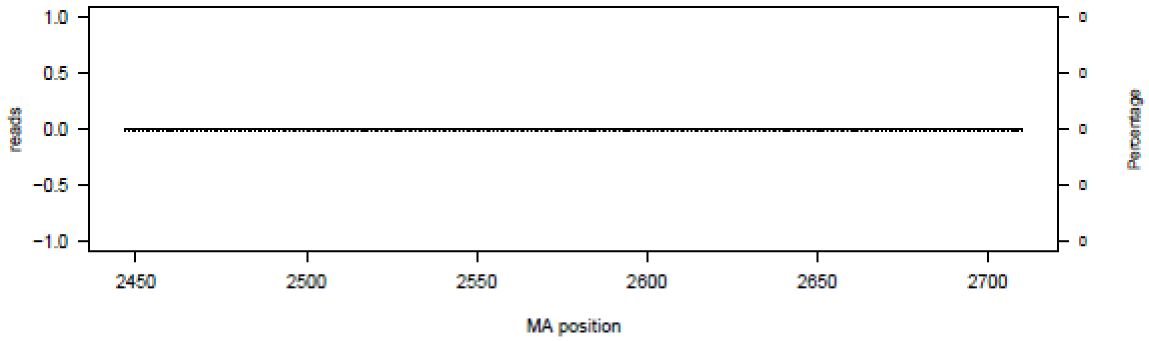


## P12 – N09–2447

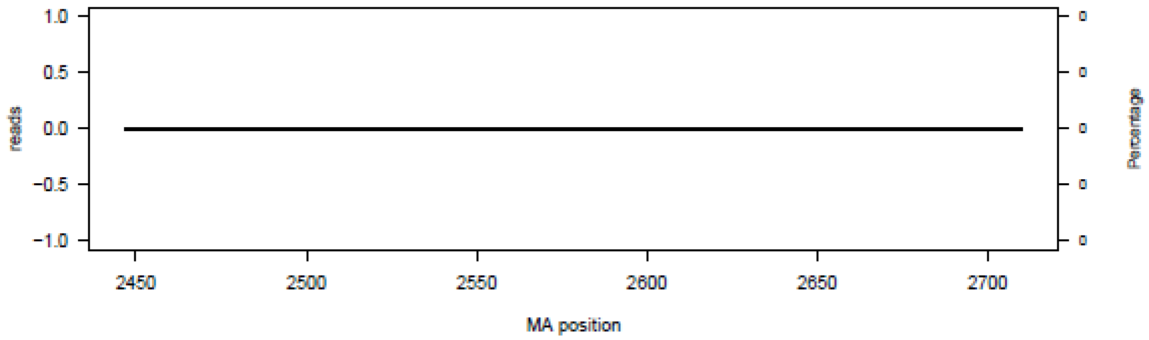

### P13 – N09−2447

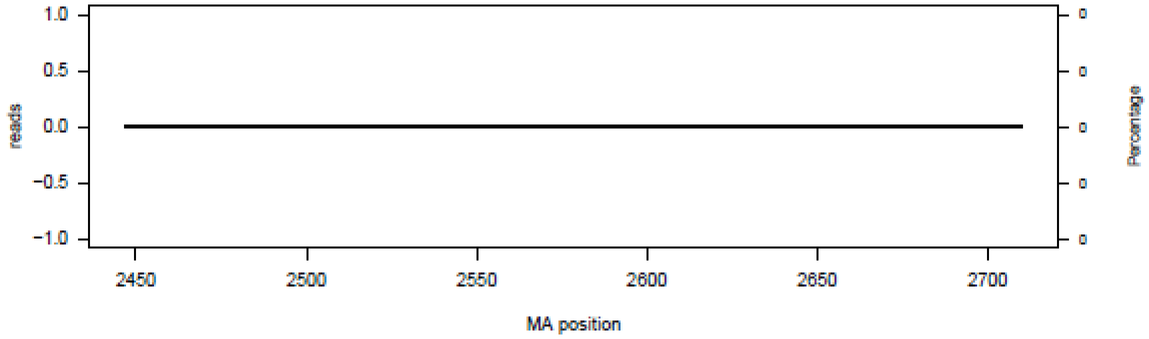


### P14 – N09−2447

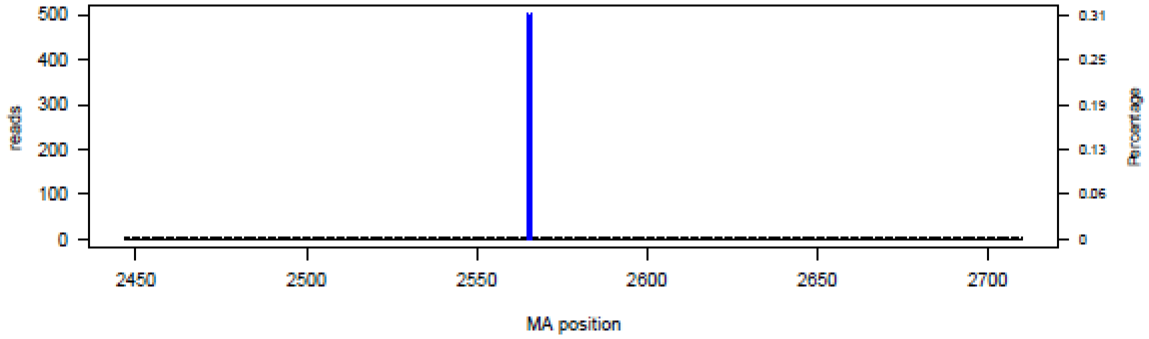


### P15 – N09−2447

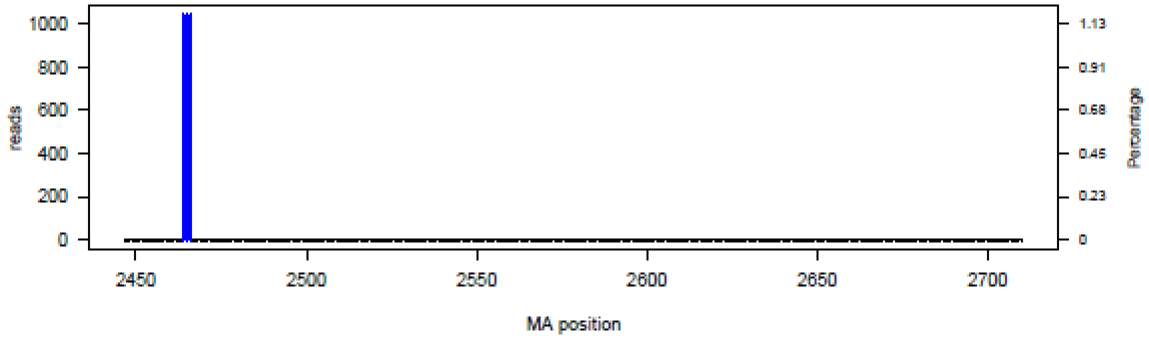


### P16 – N09−2447

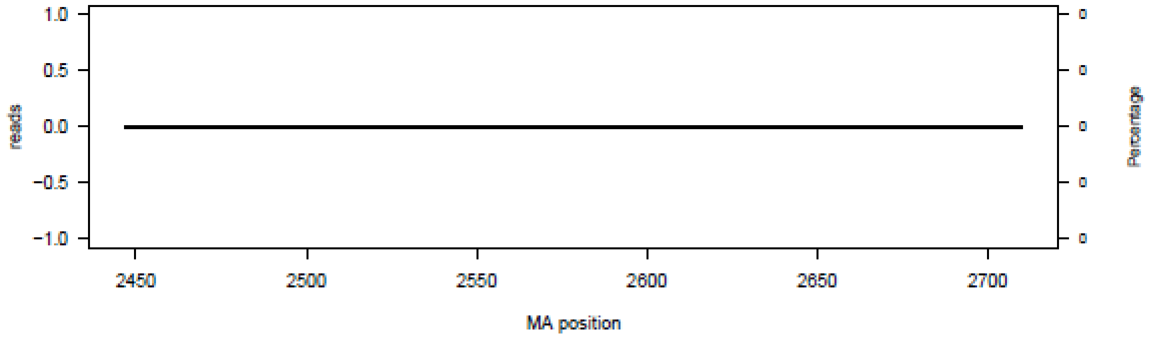

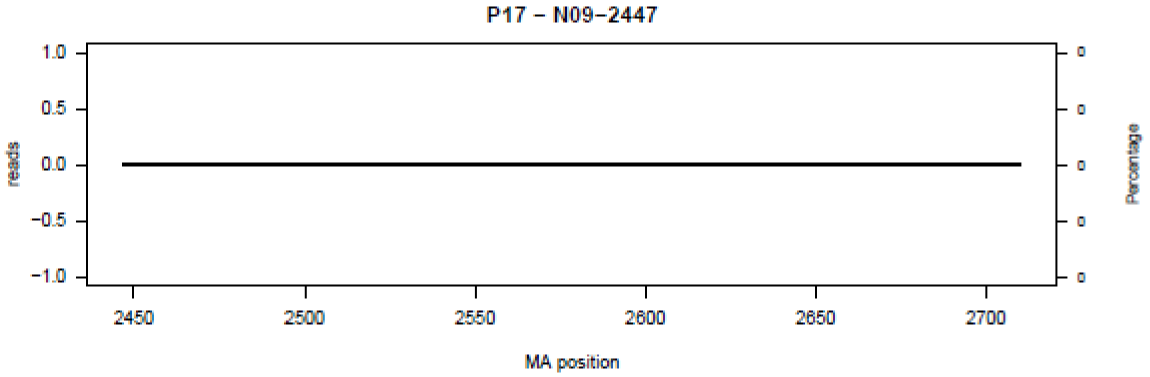
P17 − N09−2447
reads
Percentage
MA position
1.0
0.5
0.0
−0.5
−1.0
2450
2500
2550
2600
2650
2700

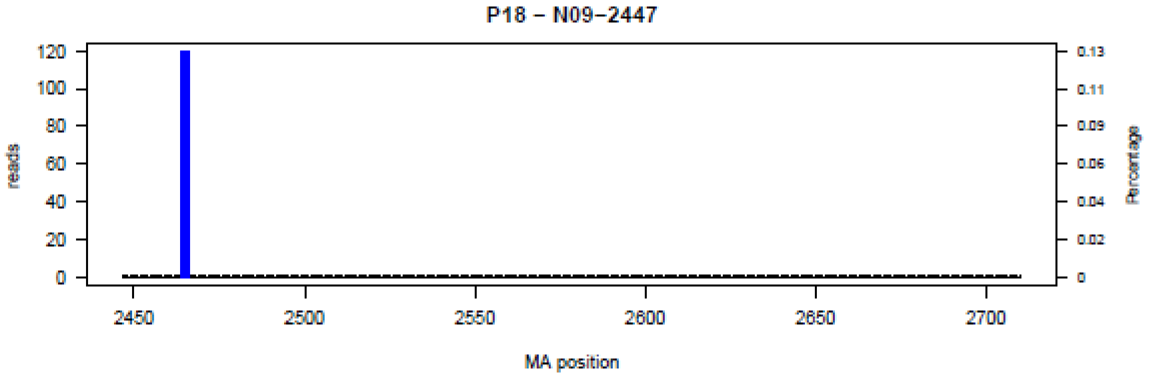
P18 − N09−2447
reads
Percentage
MA position
120
100
80
60
40
20
0
0.13
0.11
0.09
0.06
0.04
0.02
2450
2500
2550
2600
2650
2700

**Figure S10.** Bar plot of deletions in amplicon N10 in the 18 patients (P01-P18) at the nucleotide level.

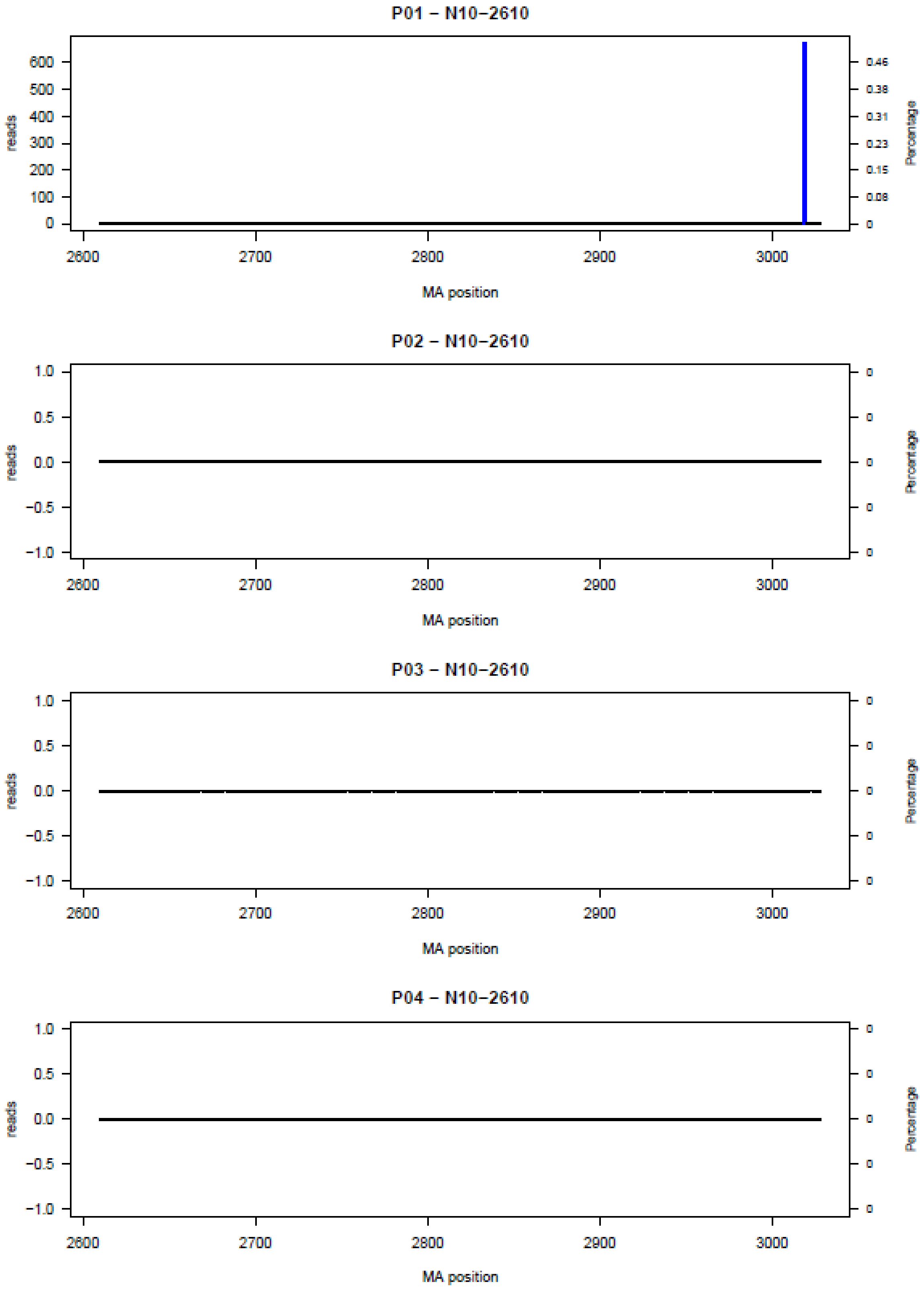

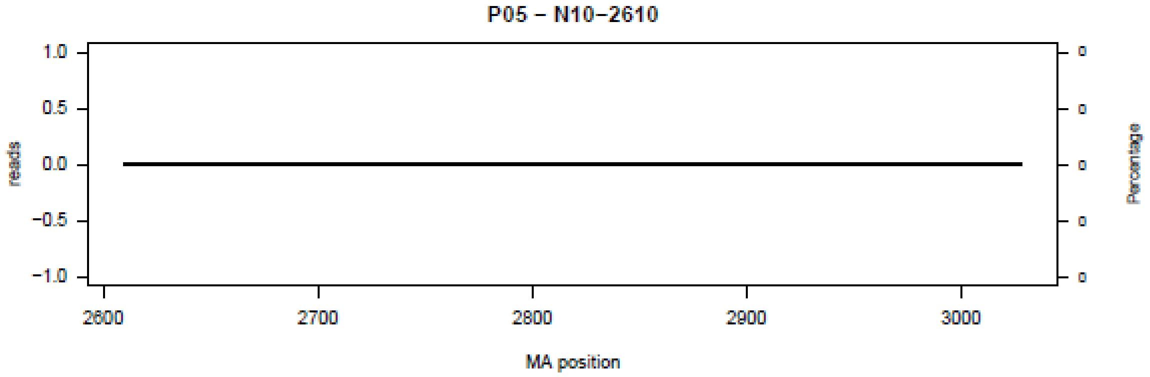
P05 – N10–2610
reads
Percentage
MA position

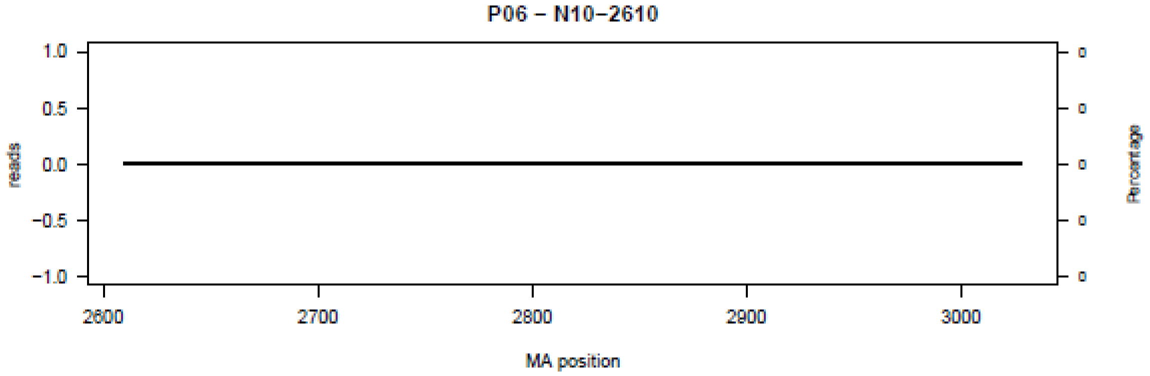
P06 – N10–2610
reads
Percentage
MA position

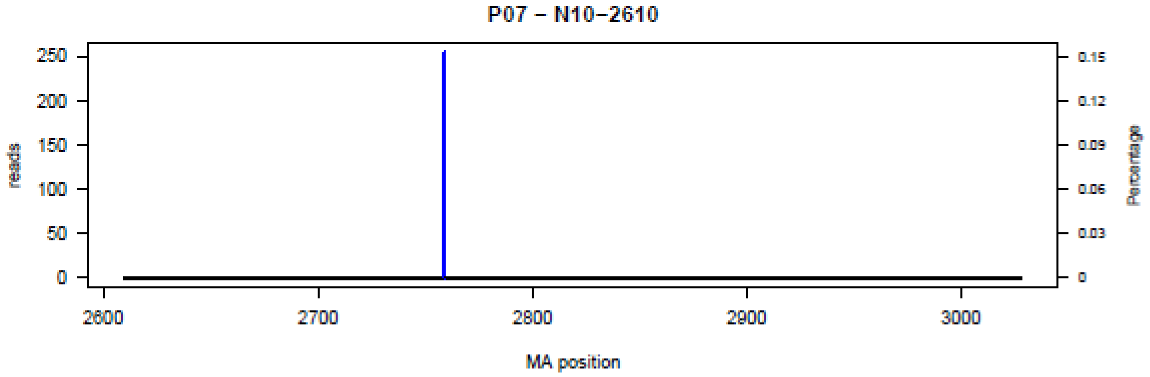
P07 – N10–2610
reads
Percentage
MA position

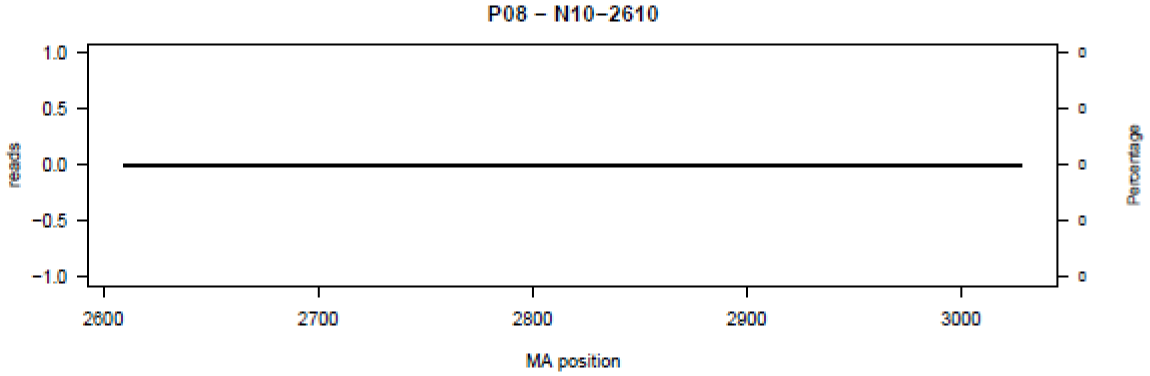
P08 – N10–2610
reads
Percentage
MA position

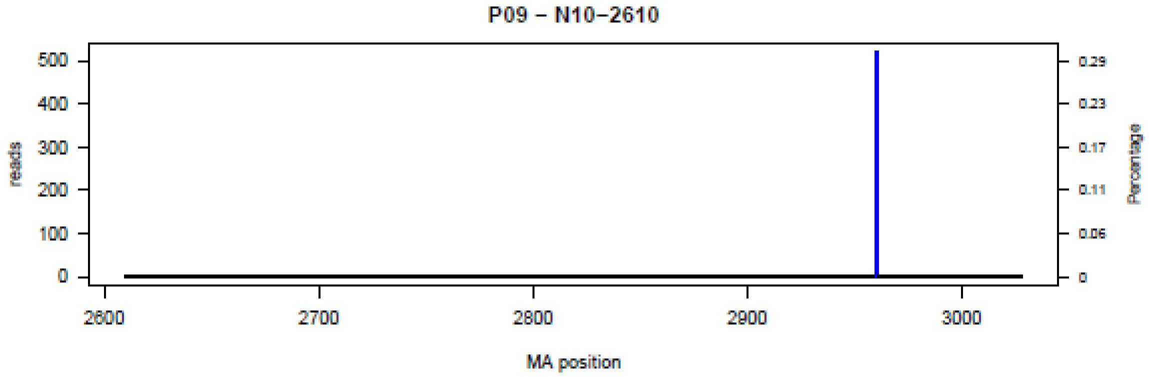
P09 – N10–2610
reads
Percentage
MA position
500
400
300
200
100
0
0.29
0.23
0.17
0.11
0.06
0
2600
2700
2800
2900
3000

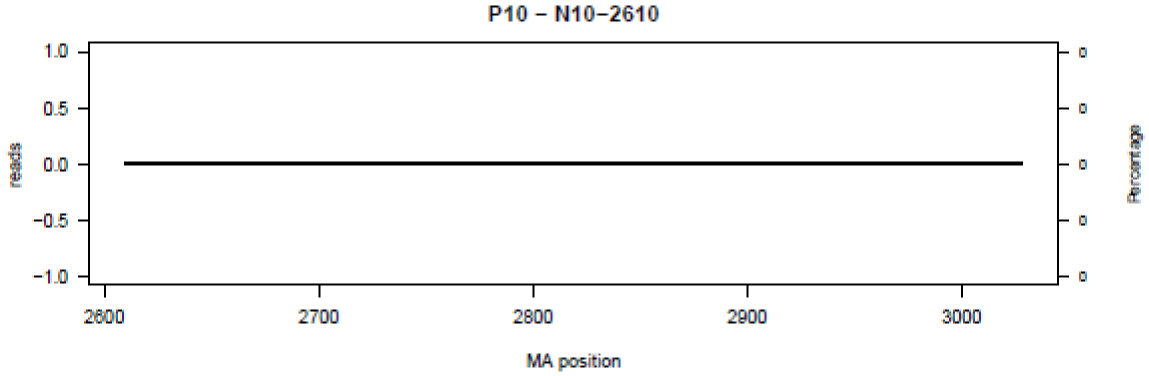
P10 – N10–2610
reads
Percentage
MA position
1.0
0.5
0.0
−0.5
−1.0
2600
2700
2800
2900
3000

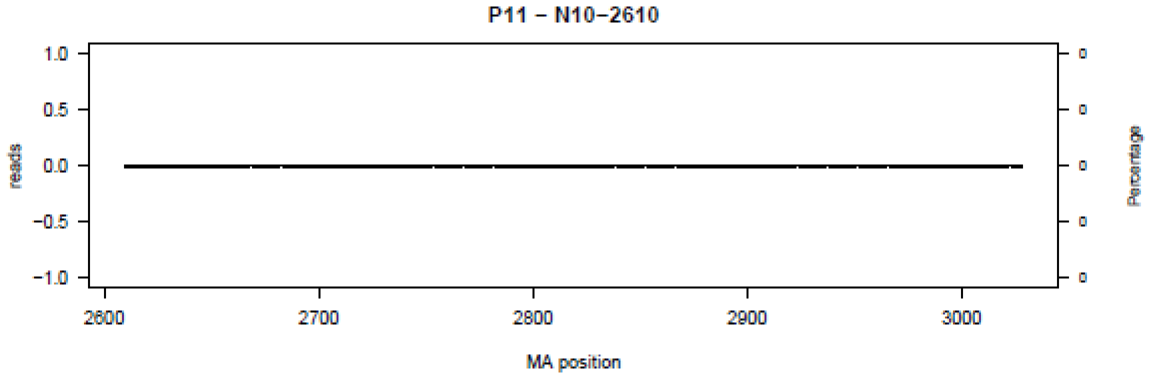
P11 – N10–2610
reads
Percentage
MA position
1.0
0.5
0.0
−0.5
−1.0
2600
2700
2800
2900
3000

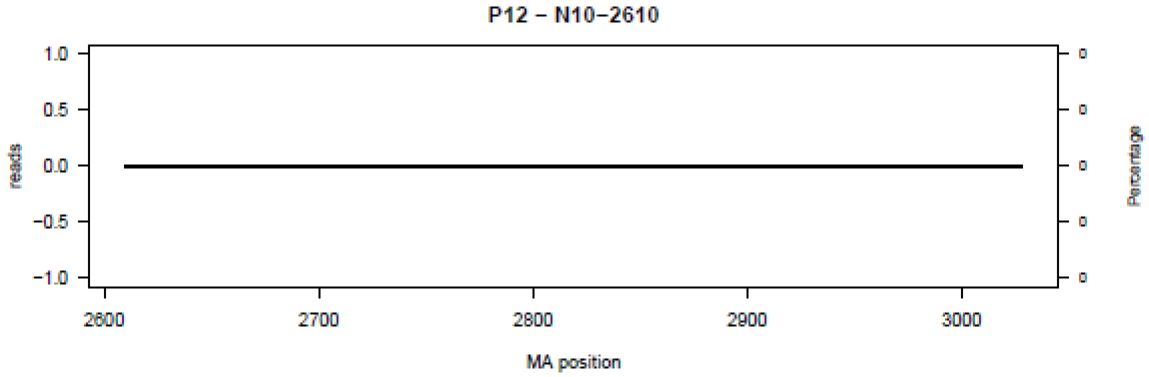
P12 – N10–2610
reads
Percentage
MA position
1.0
0.5
0.0
−0.5
−1.0
2600
2700
2800
2900
3000

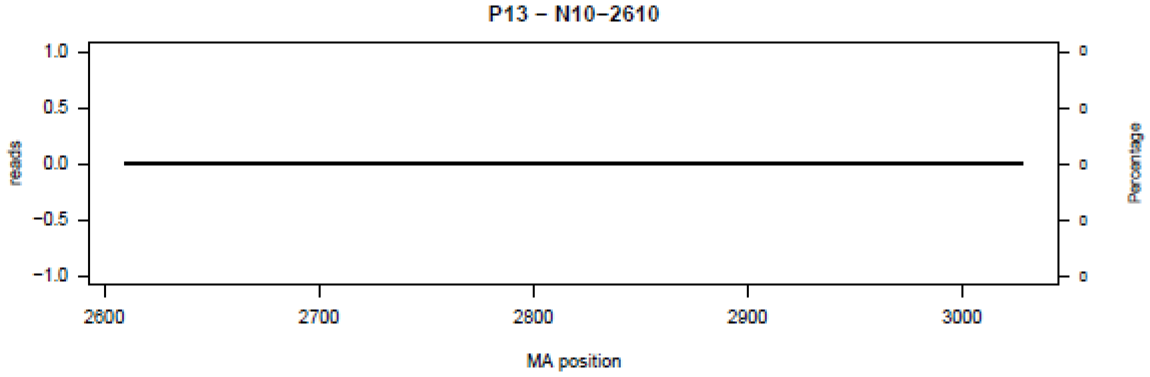
P13 – N10−2610
reads
Percentage
MA position
1.0
0.5
0.0
−0.5
−1.0
2600
2700
2800
2900
3000
0

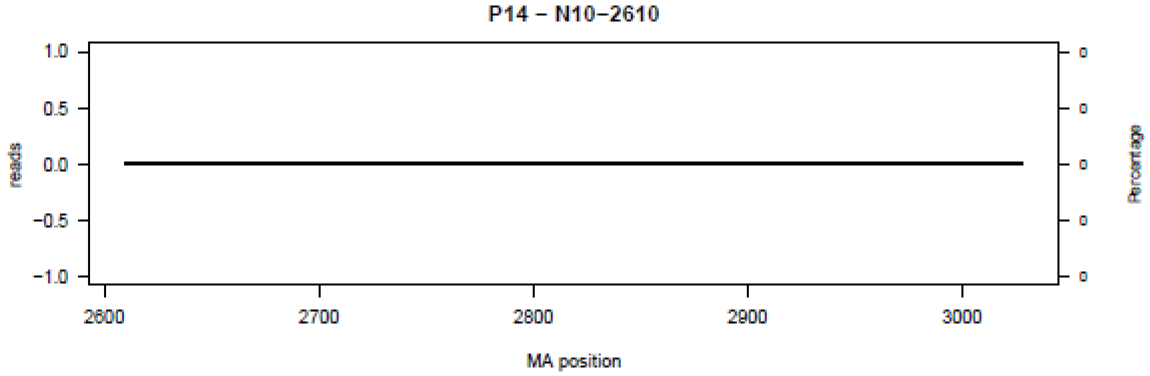
P14 – N10−2610
reads
Percentage
MA position
1.0
0.5
0.0
−0.5
−1.0
2600
2700
2800
2900
3000
0

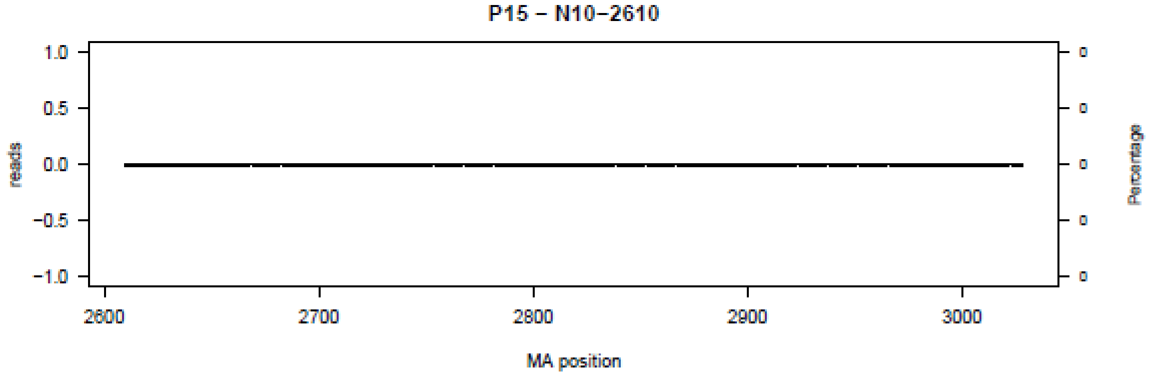
P15 – N10−2610
reads
Percentage
MA position
1.0
0.5
0.0
−0.5
−1.0
2600
2700
2800
2900
3000
0

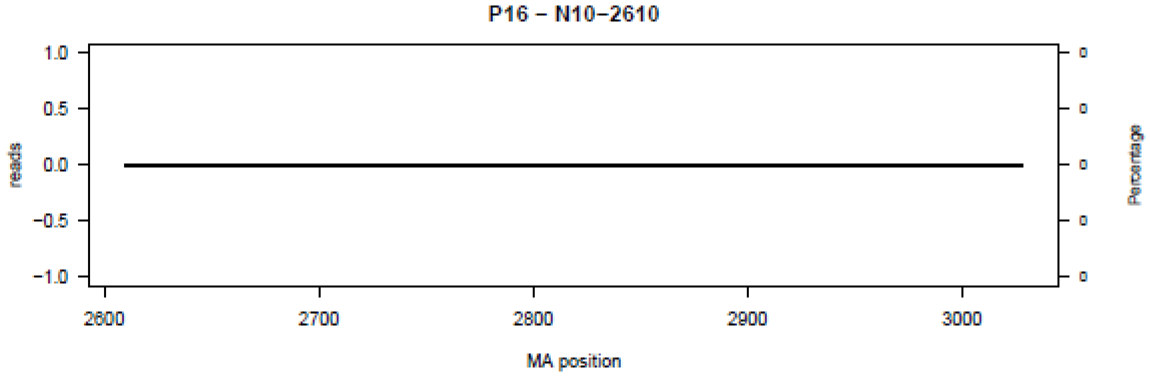
P16 – N10−2610
reads
Percentage
MA position
1.0
0.5
0.0
−0.5
−1.0
2600
2700
2800
2900
3000
0

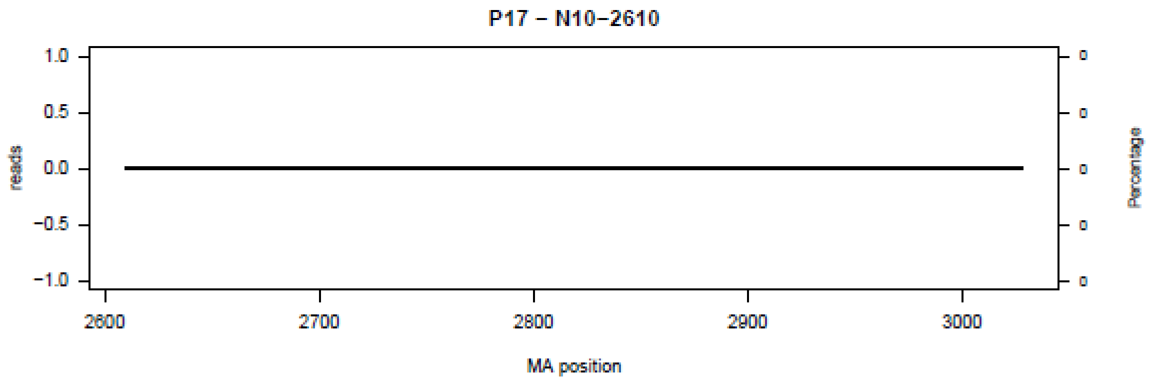
P17 – N10–2610
1.0
0.5
0.0
−0.5
−1.0
reads
0
0
0
0
0
Percentage
2600
2700
2800
2900
3000
MA position

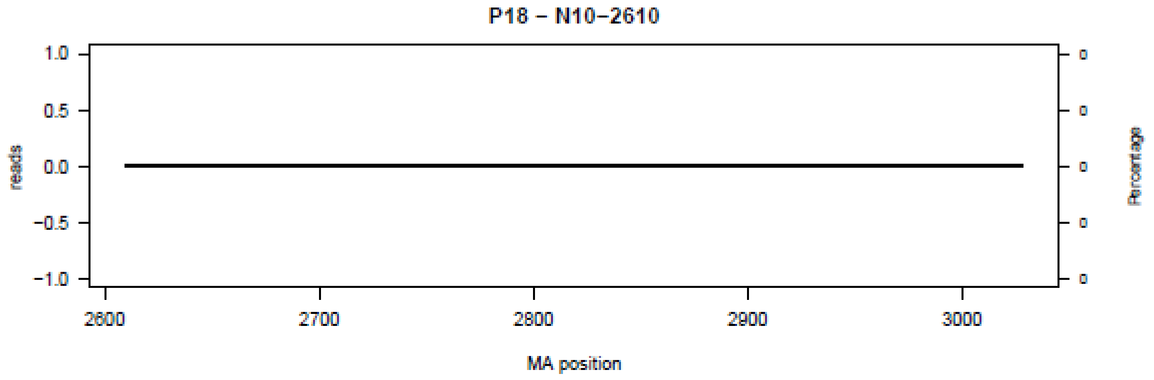
P18 – N10–2610
1.0
0.5
0.0
−0.5
−1.0
reads
0
0
0
0
0
Percentage
2600
2700
2800
2900
3000
MA position

**Figure S11.** Bar plot of deletions in amplicon N11 in the 18 patients (P01-P18) at the nucleotide level.

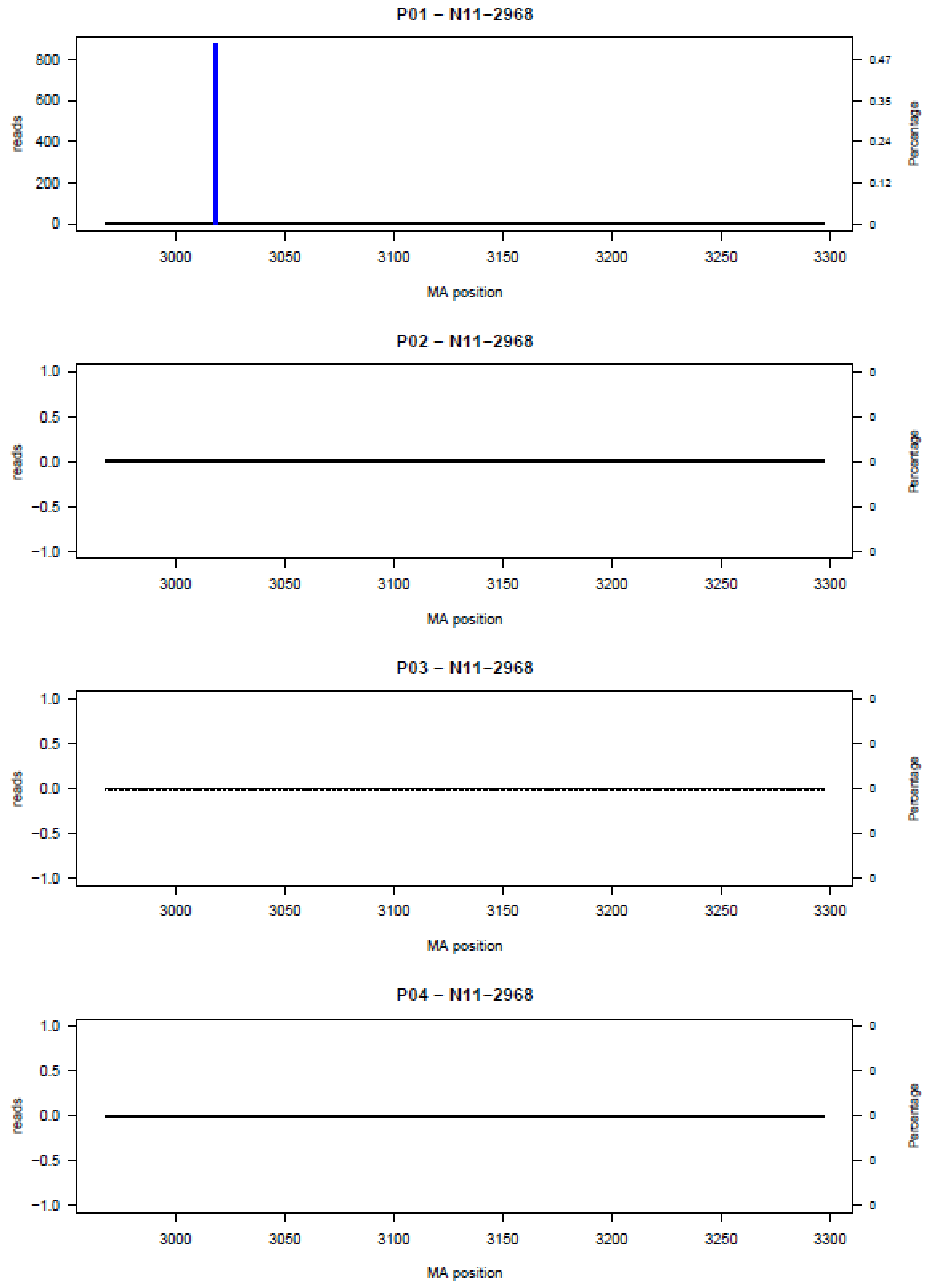

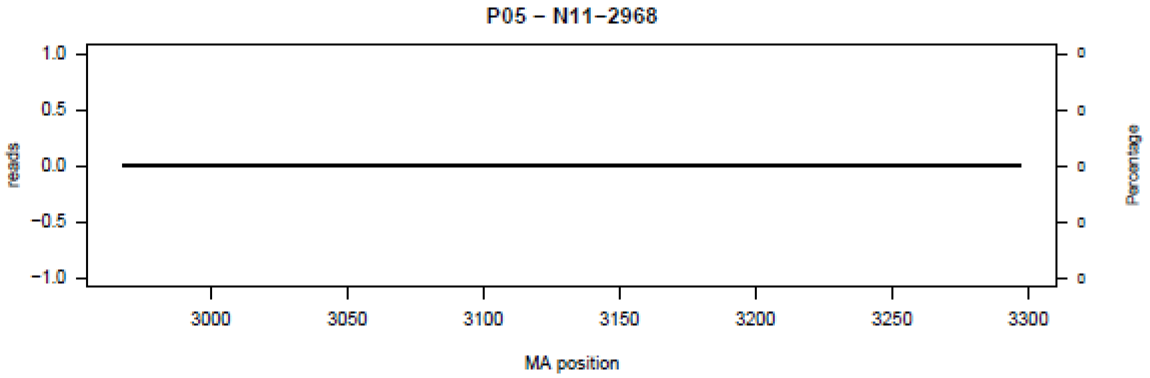
P05 – N11–2968
reads
Percentage
MA position
1.0
0.5
0.0
−0.5
−1.0
3000
3050
3100
3150
3200
3250
3300

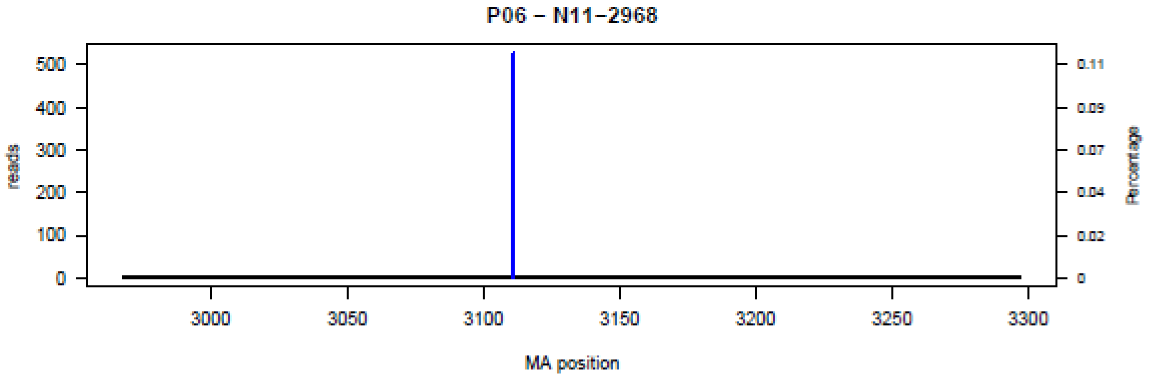
P06 – N11–2968
reads
Percentage
MA position
500
400
300
200
100
0
0.11
0.09
0.07
0.04
0.02
3000
3050
3100
3150
3200
3250
3300

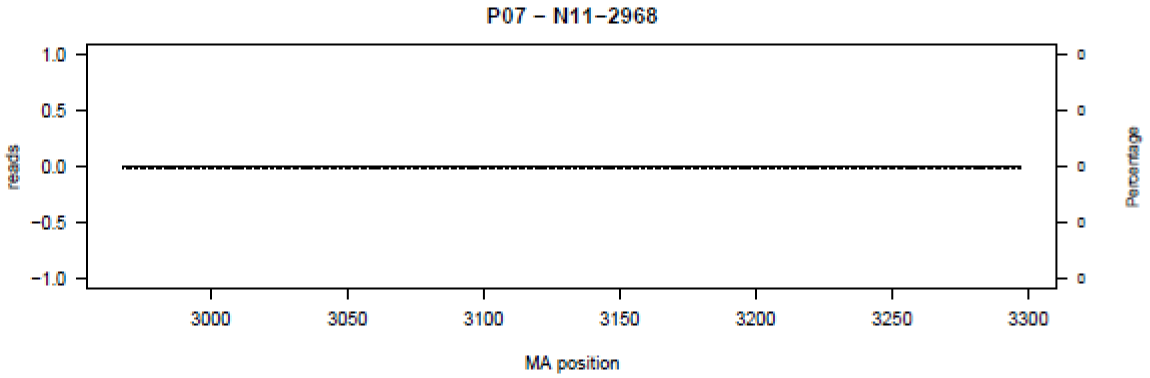
P07 – N11–2968
reads
Percentage
MA position
1.0
0.5
0.0
−0.5
−1.0
3000
3050
3100
3150
3200
3250
3300

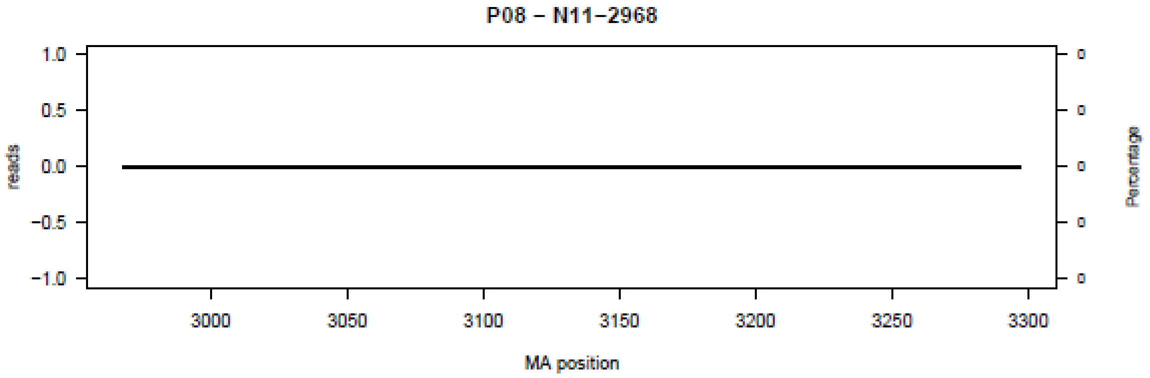
P08 – N11–2968
reads
Percentage
MA position
1.0
0.5
0.0
−0.5
−1.0
3000
3050
3100
3150
3200
3250
3300

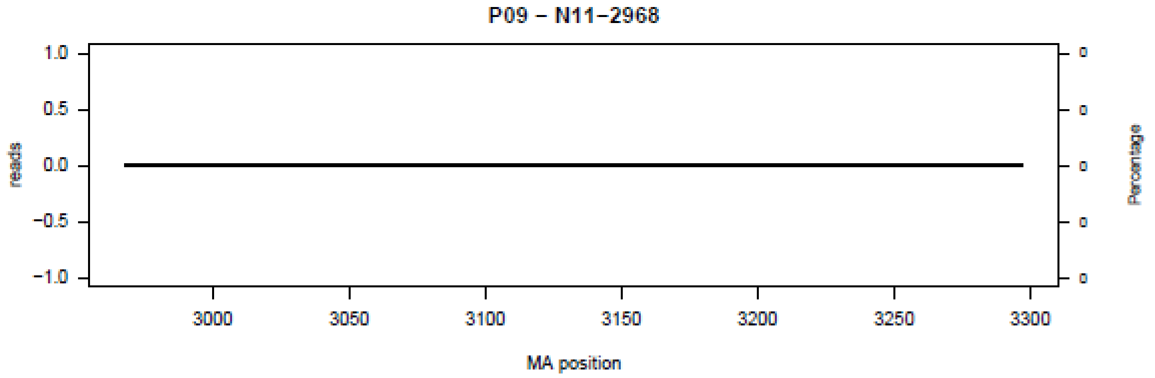
P09 – N11–2968
reads
Percentage
MA position
1.0
0.5
0.0
−0.5
−1.0
3000
3050
3100
3150
3200
3250
3300

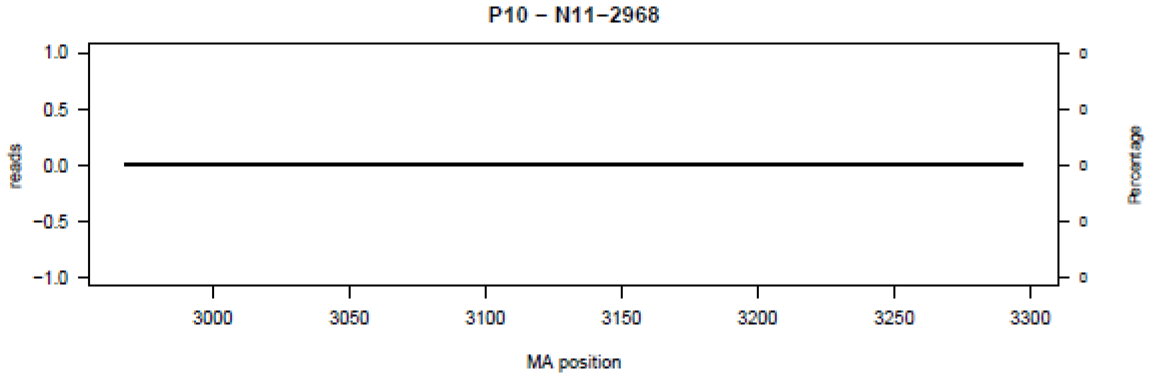
P10 – N11–2968
reads
Percentage
MA position
1.0
0.5
0.0
−0.5
−1.0
3000
3050
3100
3150
3200
3250
3300

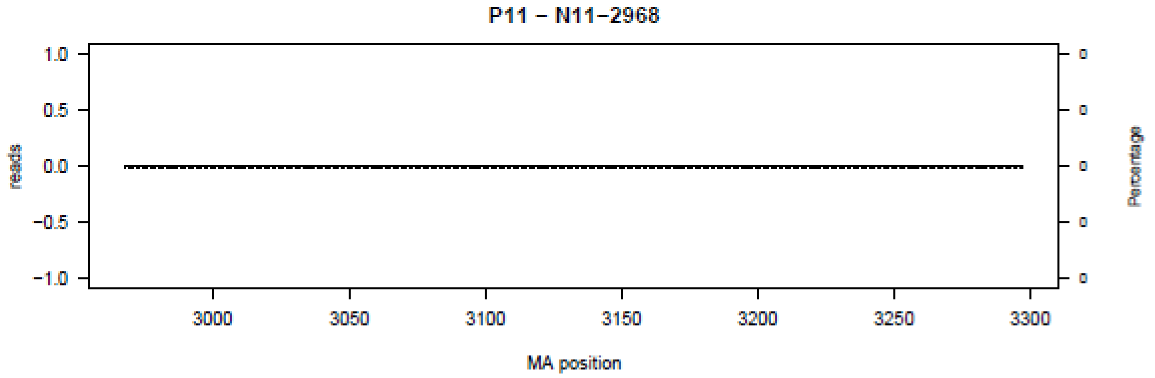
P11 – N11–2968
reads
Percentage
MA position
1.0
0.5
0.0
−0.5
−1.0
3000
3050
3100
3150
3200
3250
3300

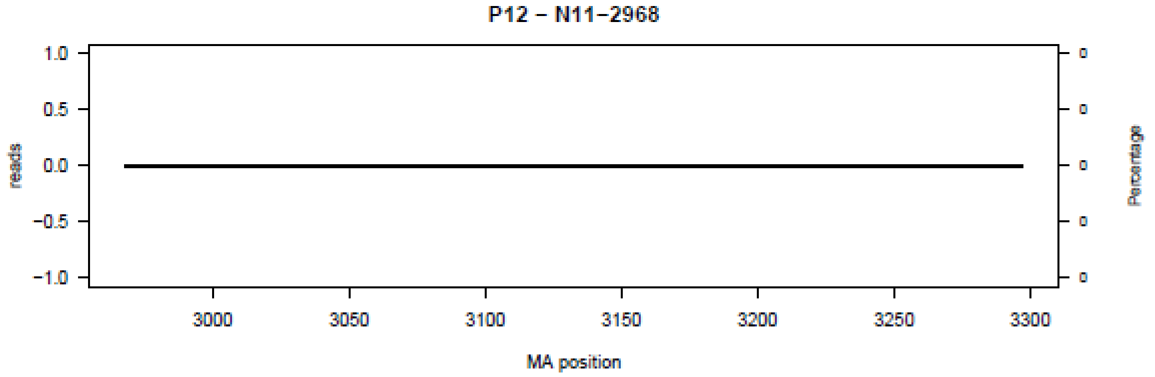
P12 – N11–2968
reads
Percentage
MA position
1.0
0.5
0.0
−0.5
−1.0
3000
3050
3100
3150
3200
3250
3300

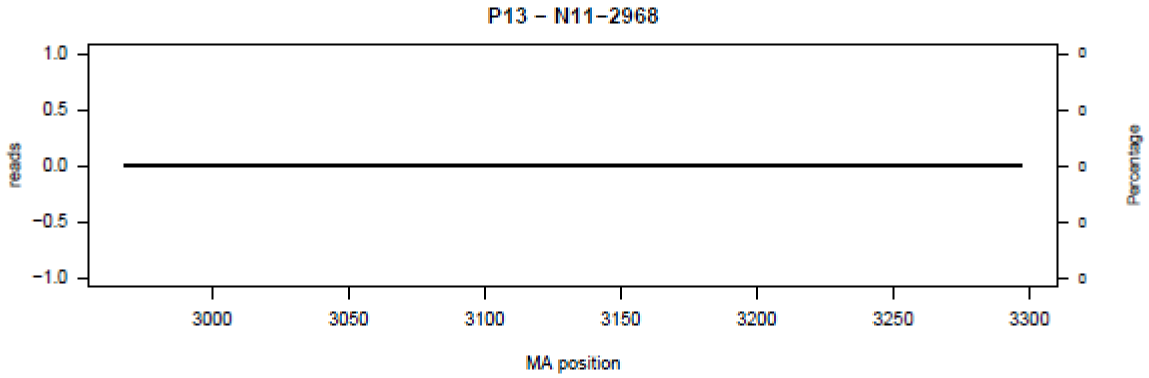
P13 – N11–2968
reads
Percentage
MA position

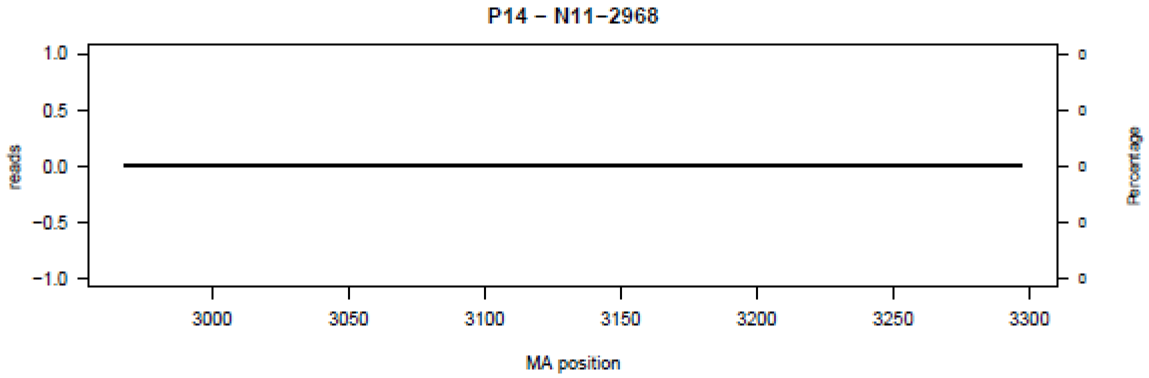
P14 – N11–2968
reads
Percentage
MA position

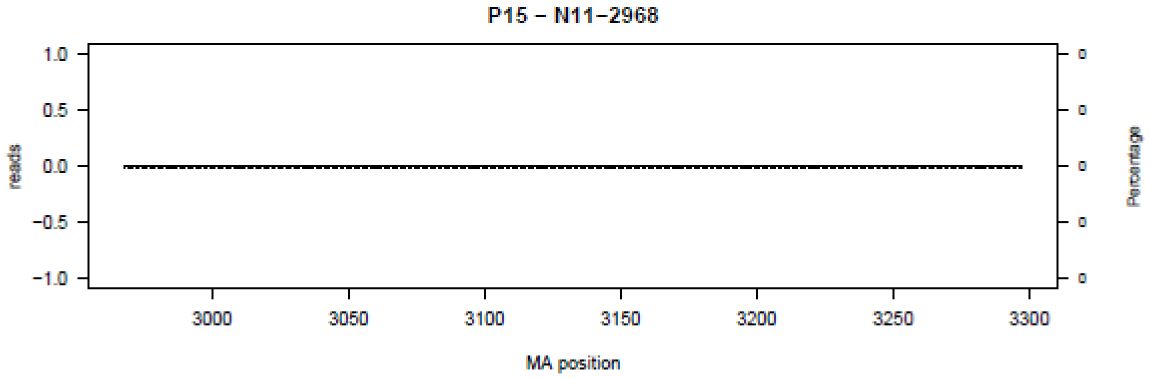
P15 – N11–2968
reads
Percentage
MA position

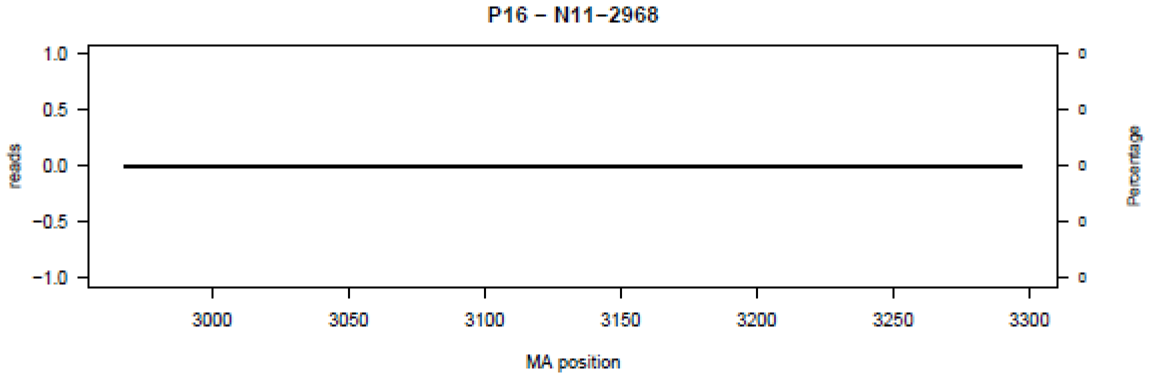
P16 – N11–2968
reads
Percentage
MA position

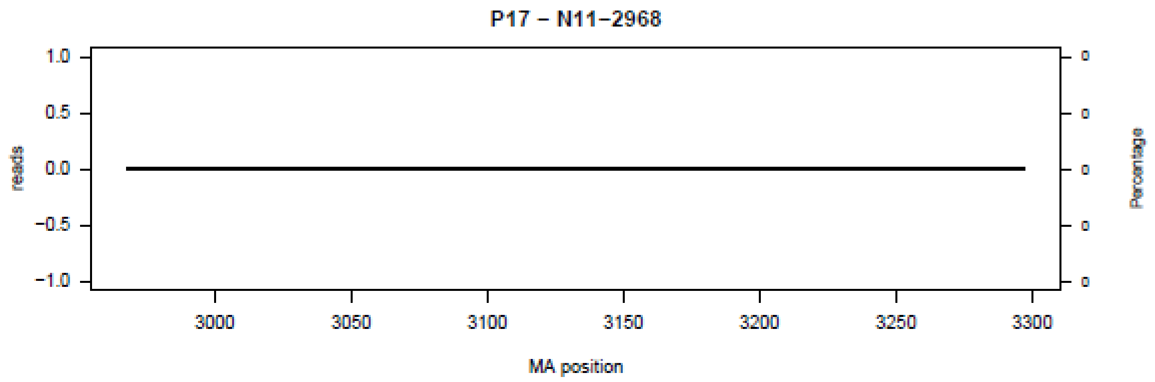
P17 – N11–2968
reads
Percentage
MA position
1.0
0.5
0.0
−0.5
−1.0
3000
3050
3100
3150
3200
3250
3300

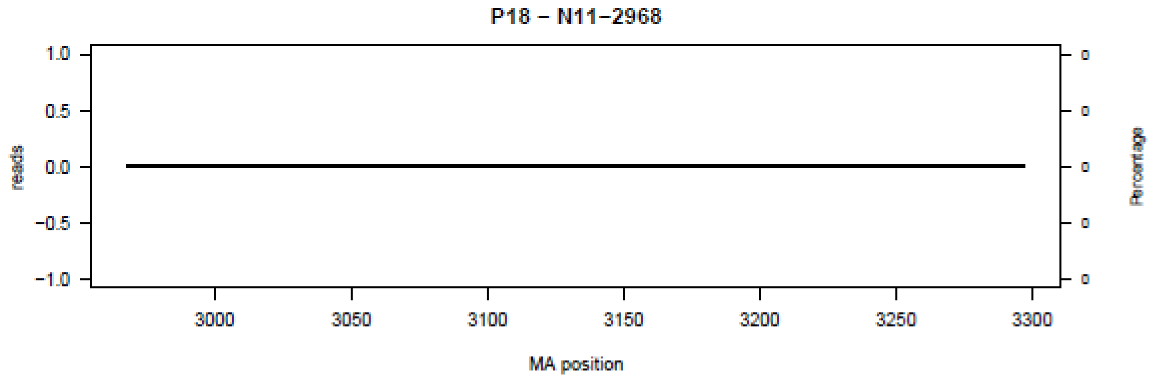
P18 – N11–2968
reads
Percentage
MA position
1.0
0.5
0.0
−0.5
−1.0
3000
3050
3100
3150
3200
3250
3300

**Figure S12.** Bar plot of deletions in amplicon N12 in the 18 patients (P01-P18) at the nucleotide level.

P01 – N12–3250

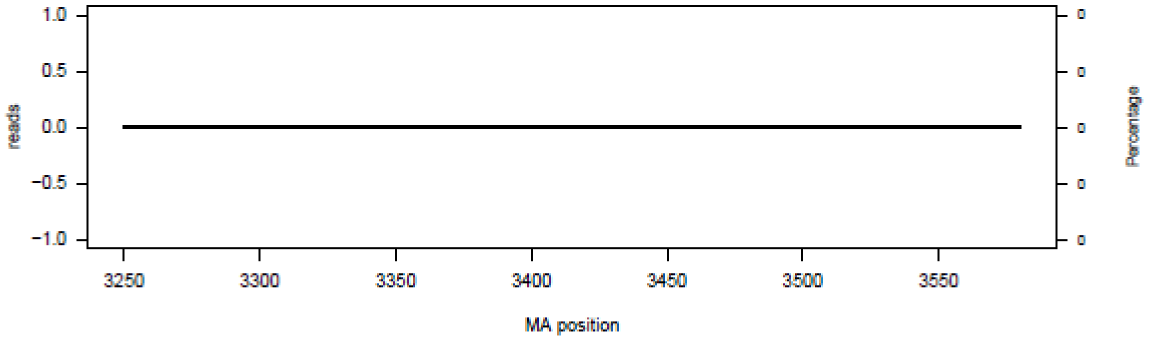


P02 – N12–3250

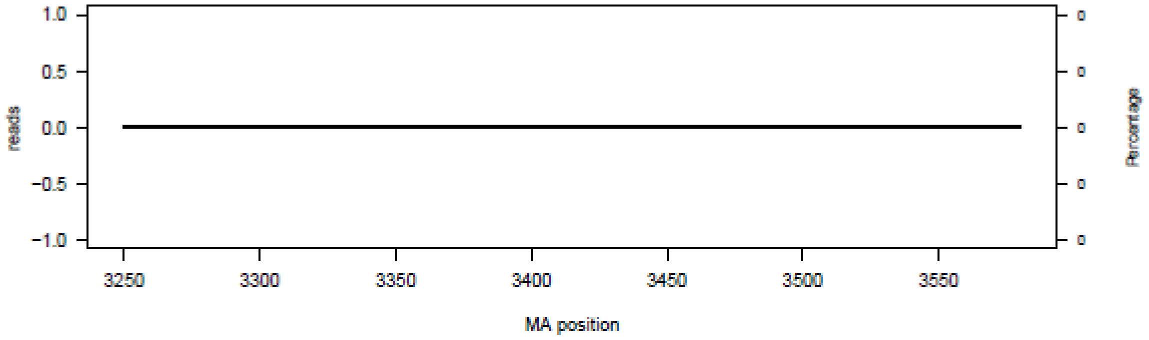


P03 – N12–3250

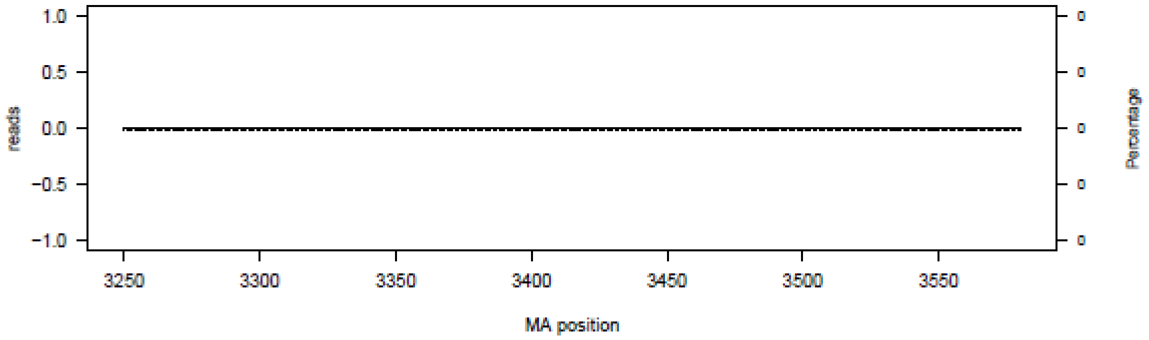


P04 – N12–3250

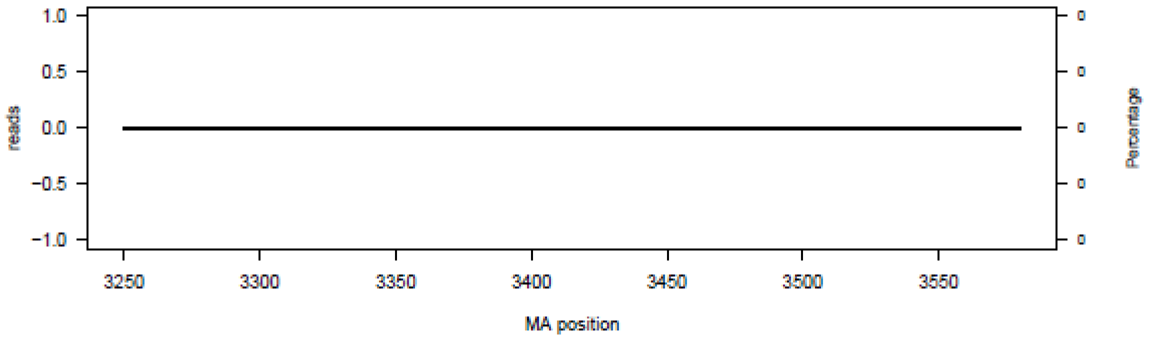

P05 – N12–3250

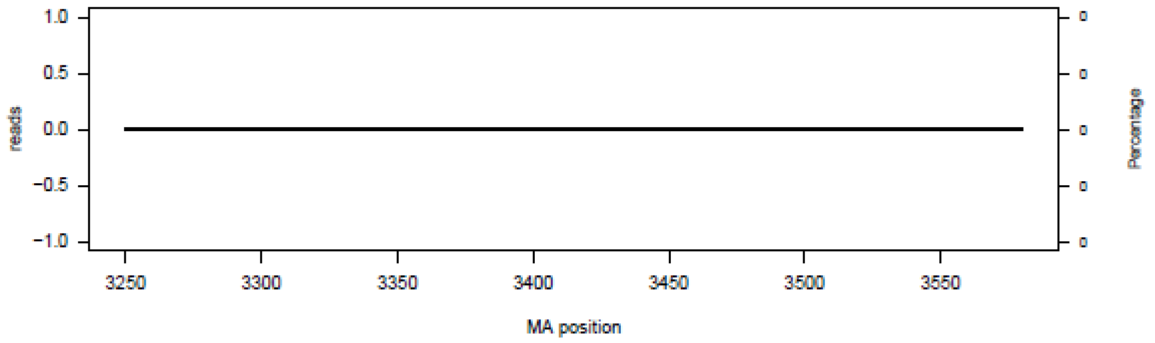


P06 – N12–3250

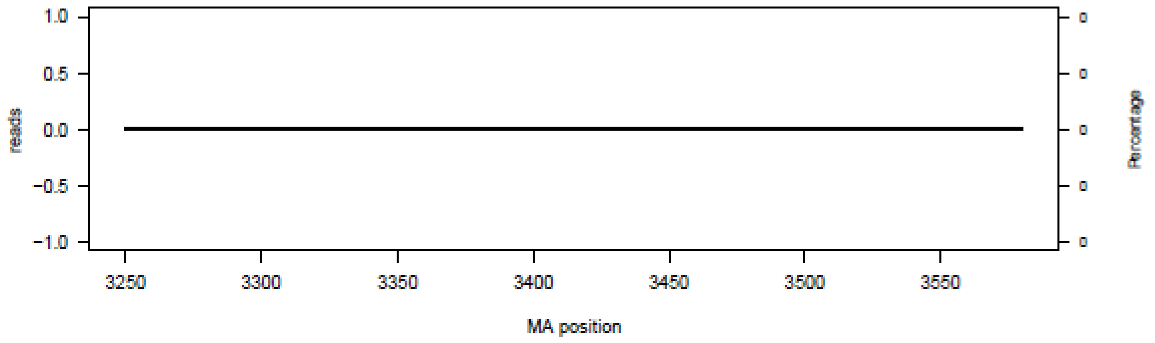


P07 – N12–3250

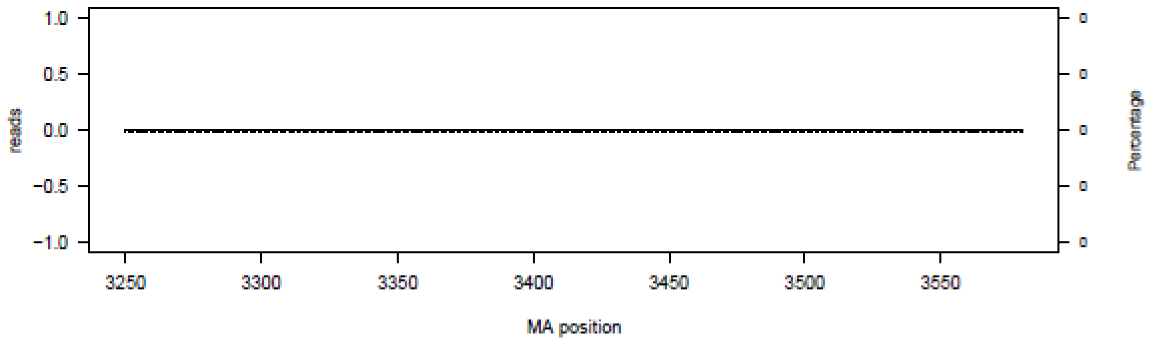


P08 – N12–3250

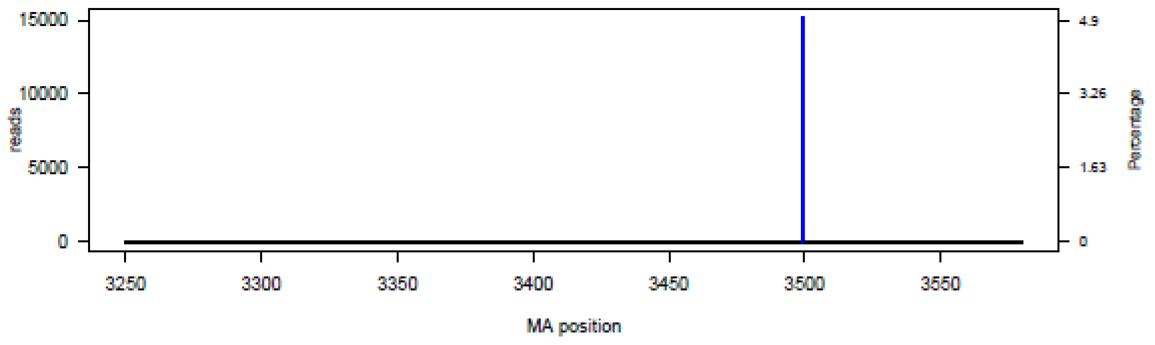

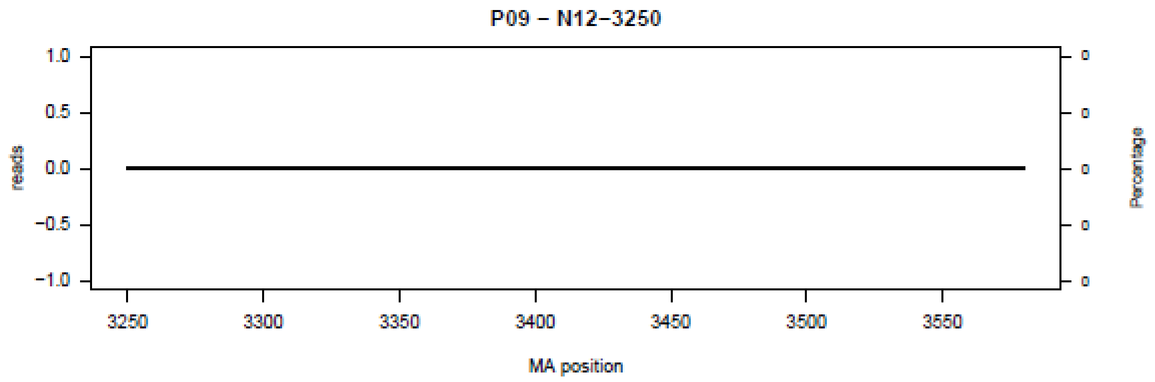
P09 – N12–3250
1.0
0.5
0.0
−0.5
−1.0
reads
3250
3300
3350
3400
3450
3500
3550
MA position
Percentage

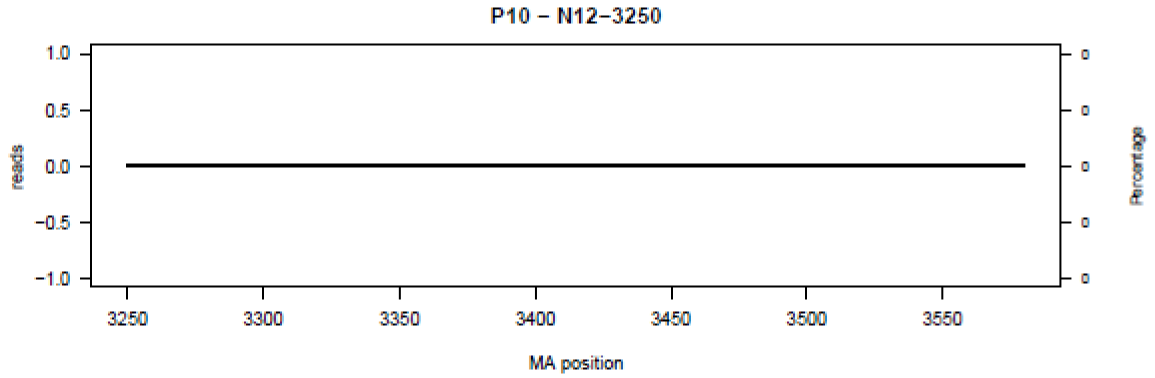
P10 – N12–3250
1.0
0.5
0.0
−0.5
−1.0
reads
3250
3300
3350
3400
3450
3500
3550
MA position
Percentage

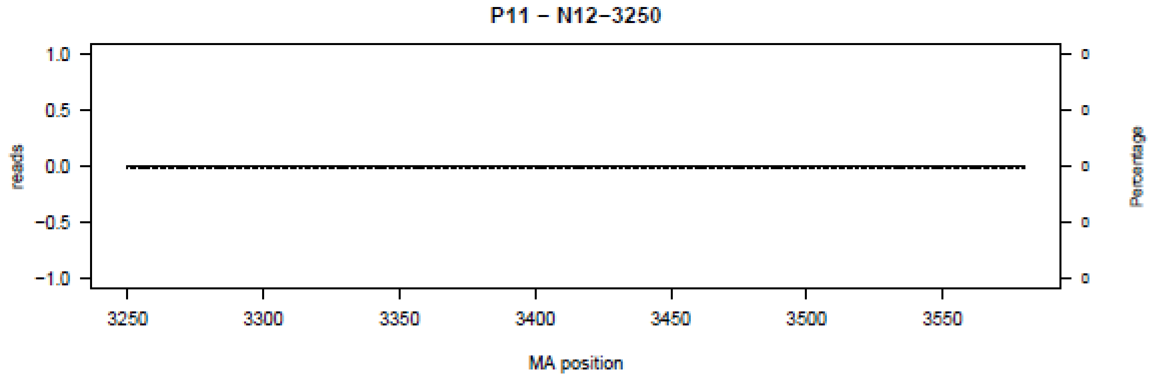
P11 – N12–3250
1.0
0.5
0.0
−0.5
−1.0
reads
3250
3300
3350
3400
3450
3500
3550
MA position
Percentage

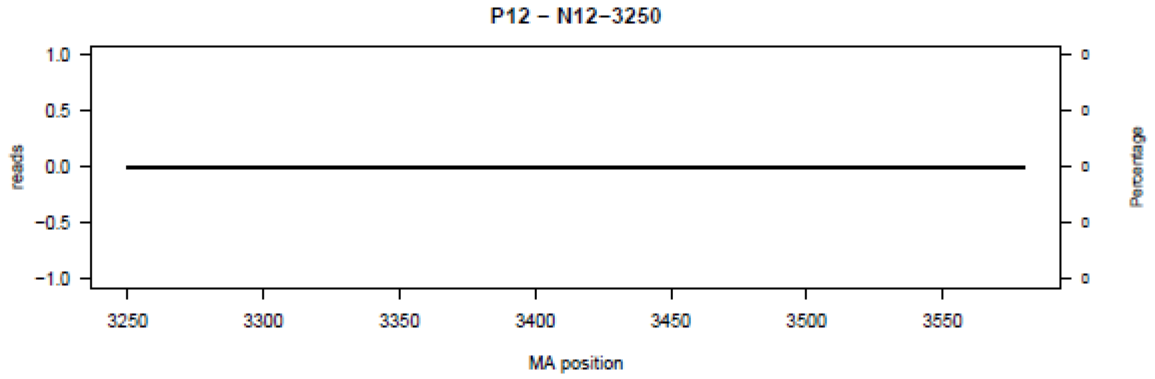
P12 – N12–3250
1.0
0.5
0.0
−0.5
−1.0
reads
3250
3300
3350
3400
3450
3500
3550
MA position
Percentage

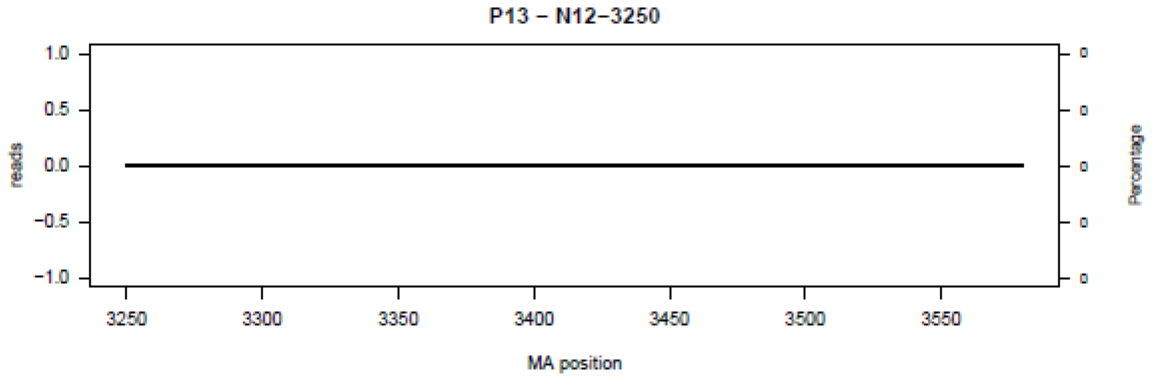
P13 – N12–3250
reads
Percentage
MA position
1.0
0.5
0.0
−0.5
−1.0
3250
3300
3350
3400
3450
3500
3550
0

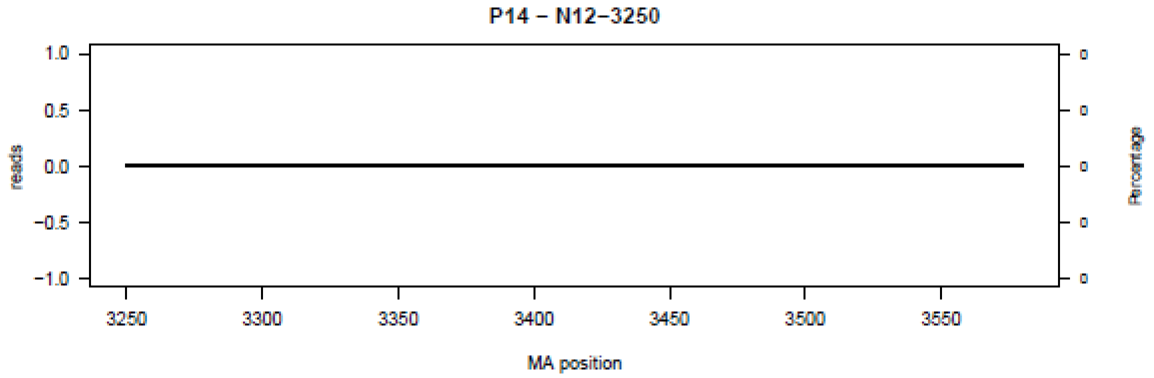
P14 – N12–3250
reads
Percentage
MA position
1.0
0.5
0.0
−0.5
−1.0
3250
3300
3350
3400
3450
3500
3550
0

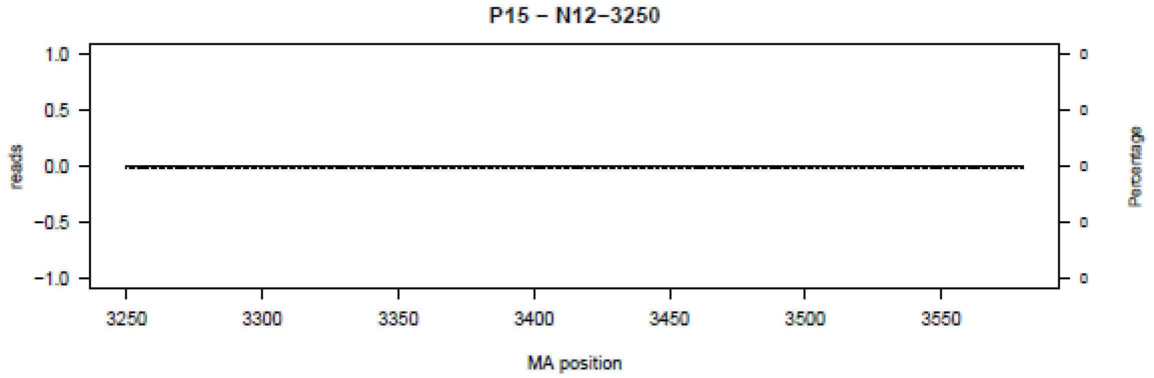
P15 – N12–3250
reads
Percentage
MA position
1.0
0.5
0.0
−0.5
−1.0
3250
3300
3350
3400
3450
3500
3550
0

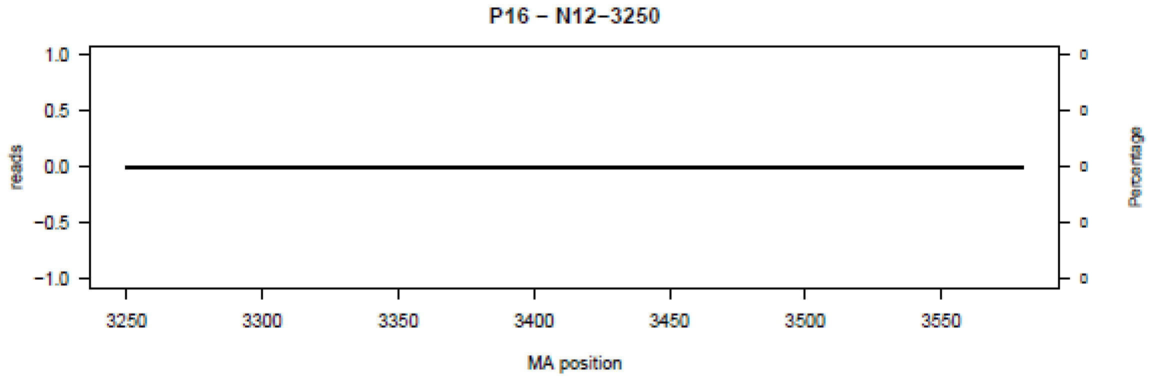
P16 – N12–3250
reads
Percentage
MA position
1.0
0.5
0.0
−0.5
−1.0
3250
3300
3350
3400
3450
3500
3550
0

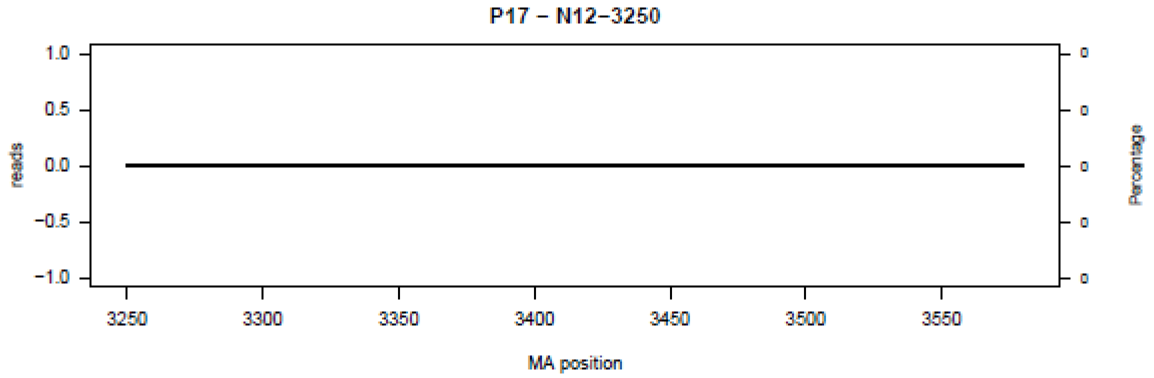
P17 – N12–3250
reads
Percentage
MA position
1.0
0.5
0.0
−0.5
−1.0
3250
3300
3350
3400
3450
3500
3550

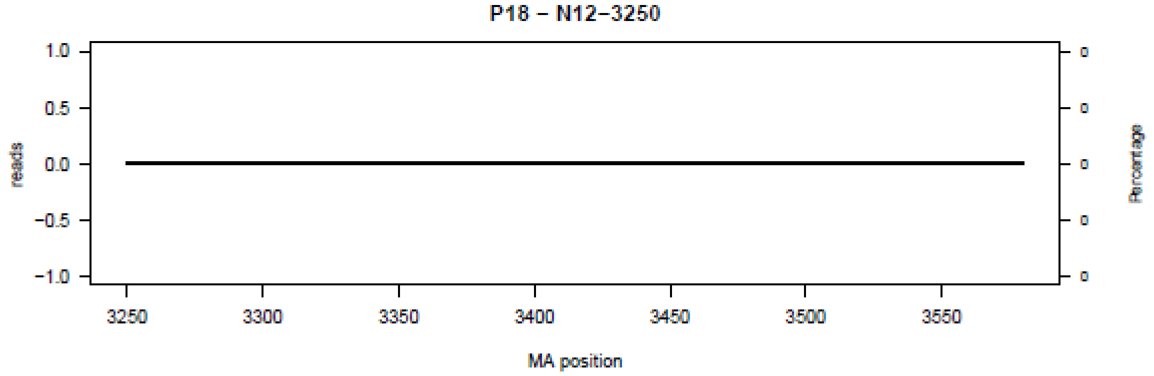
P18 – N12–3250
reads
Percentage
MA position
1.0
0.5
0.0
−0.5
−1.0
3250
3300
3350
3400
3450
3500
3550

**Figure S13.** Bar plot of deletions in amplicon N13 in the 18 patients (P01-P18) at the nucleotide level.

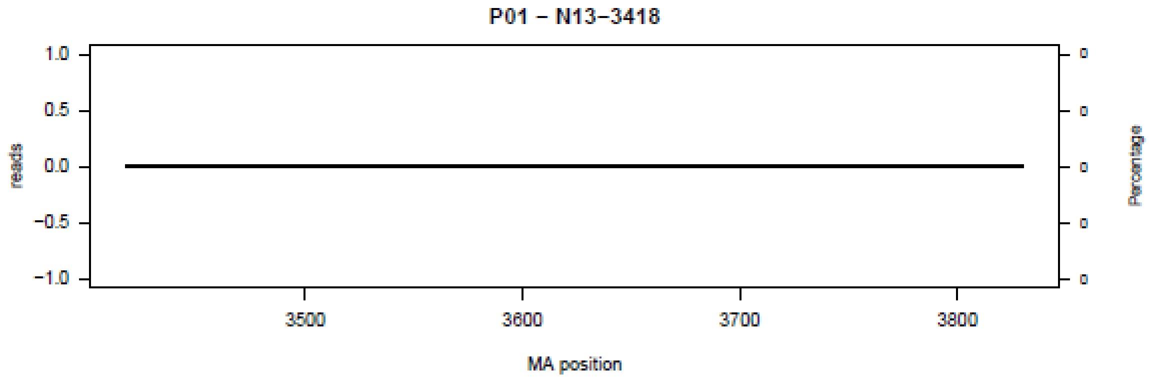


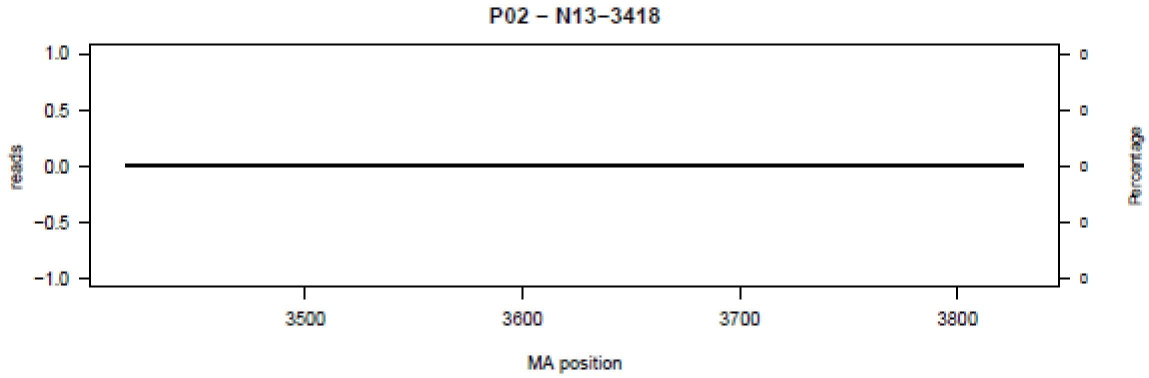


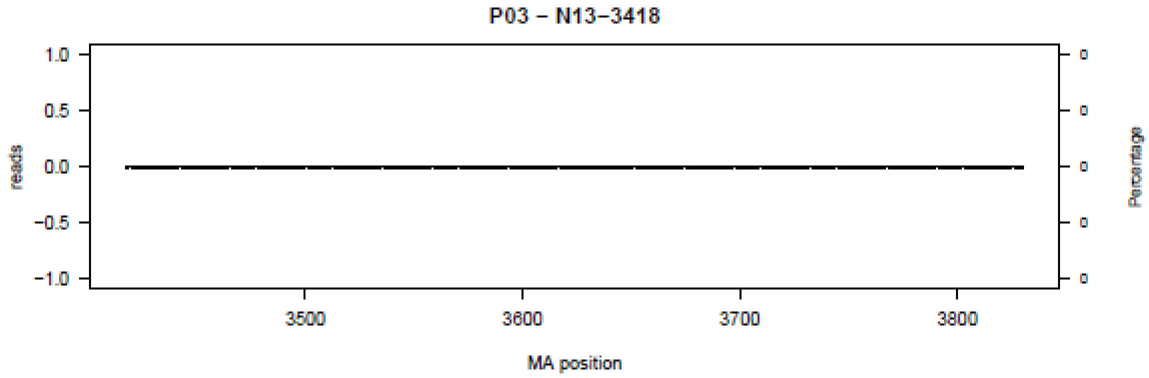


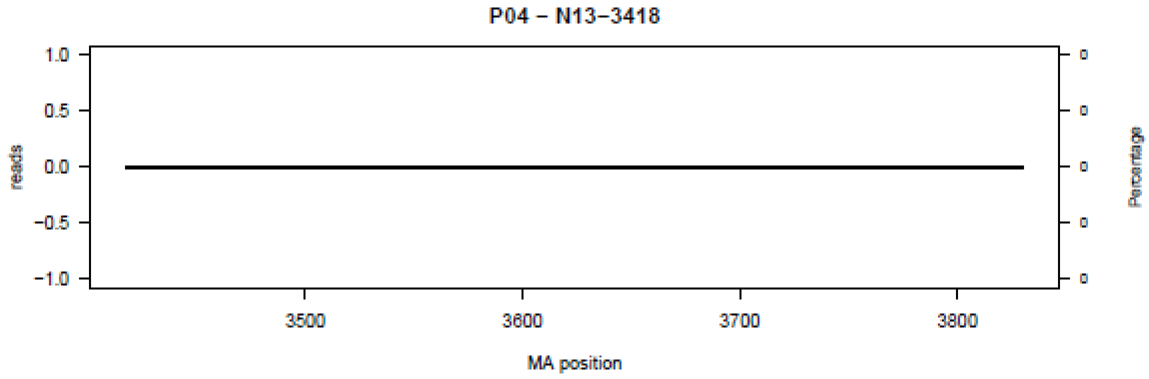

**P05 – N13–3418**

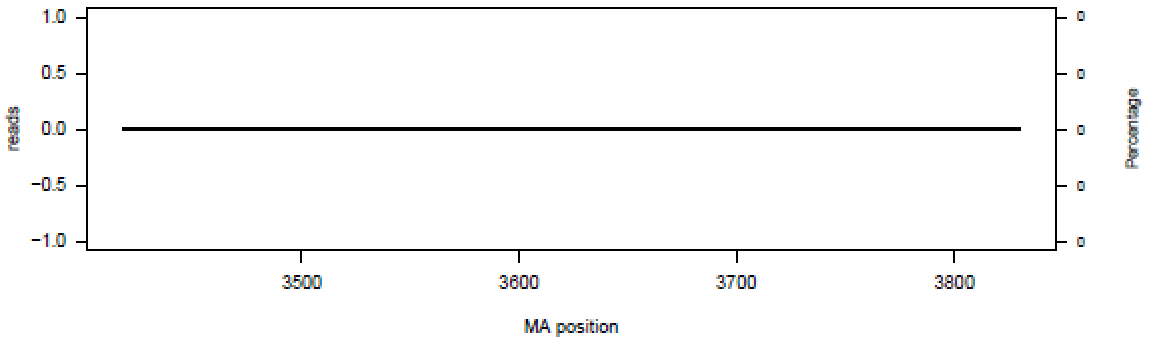


**P06 – N13–3418**

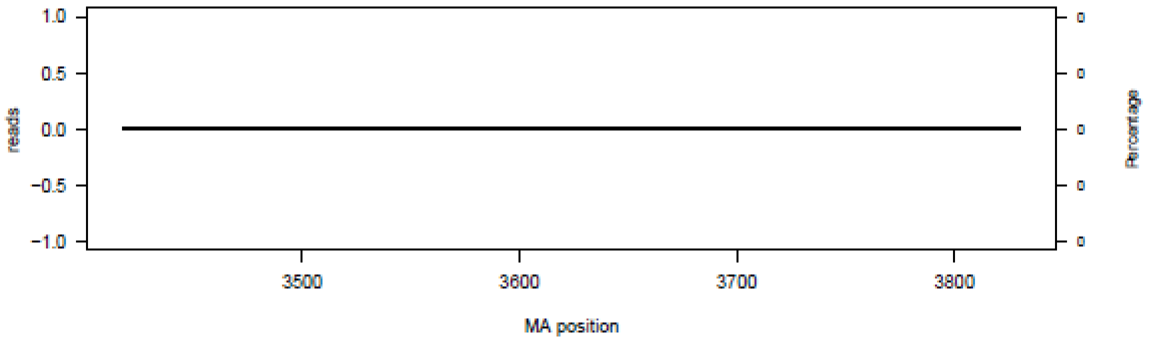


**P07 – N13–3418**

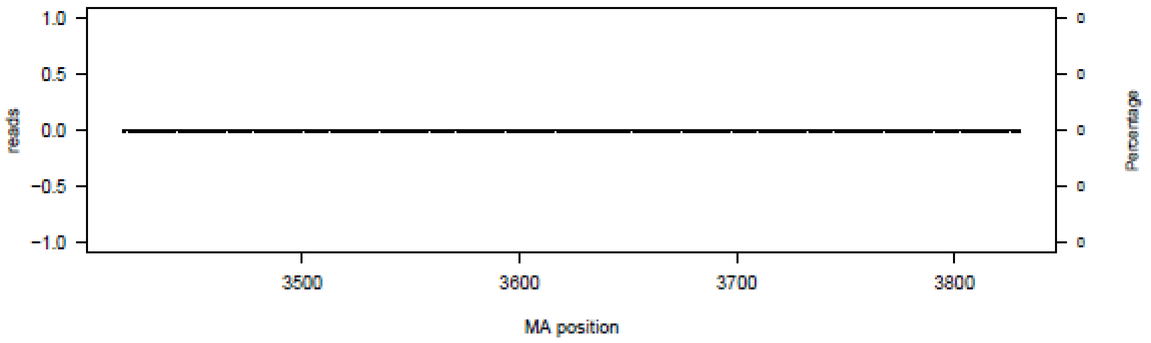


**P08 – N13–3418**

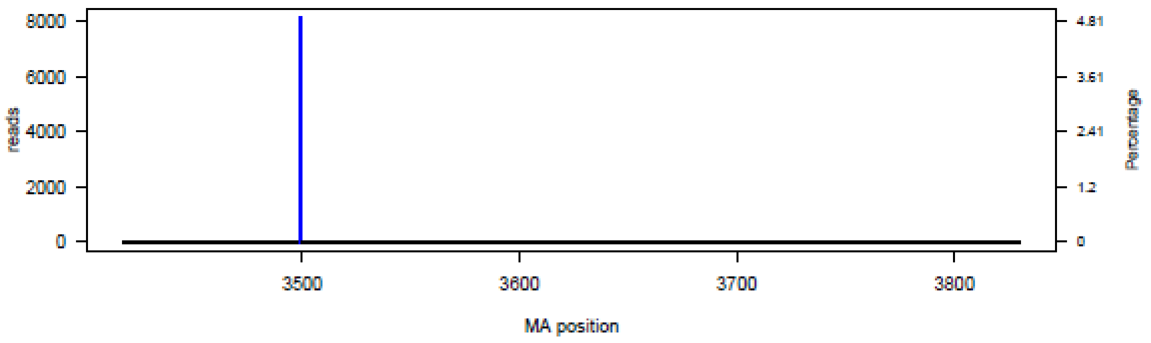

P09 – N13–3418

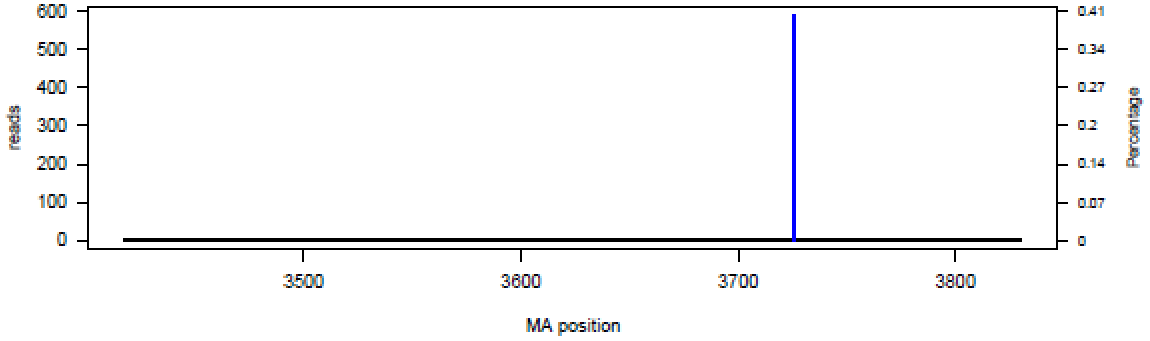


P10 – N13–3418

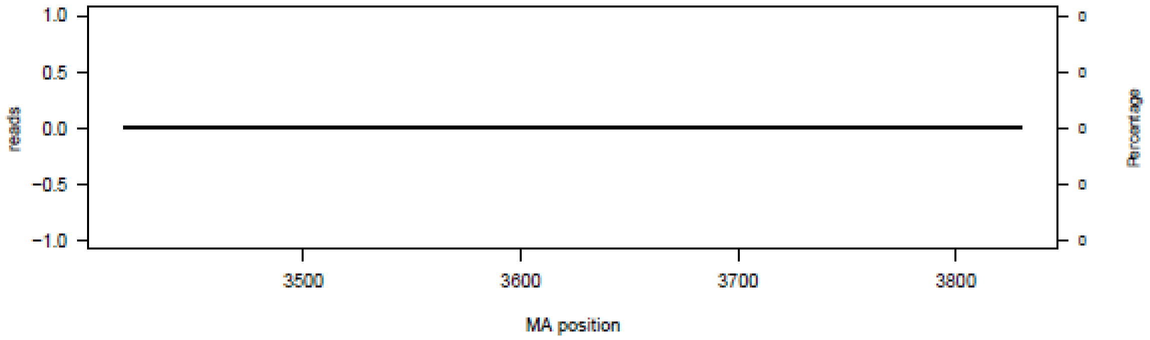


P11 – N13–3418

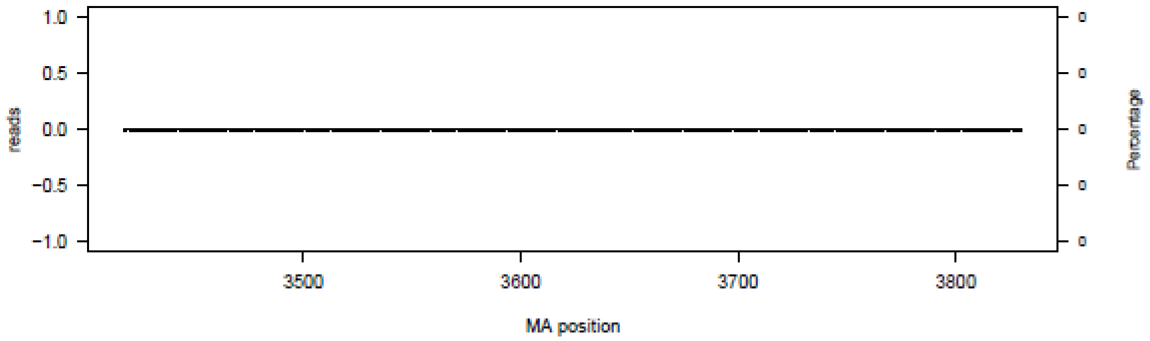


P12 – N13–3418

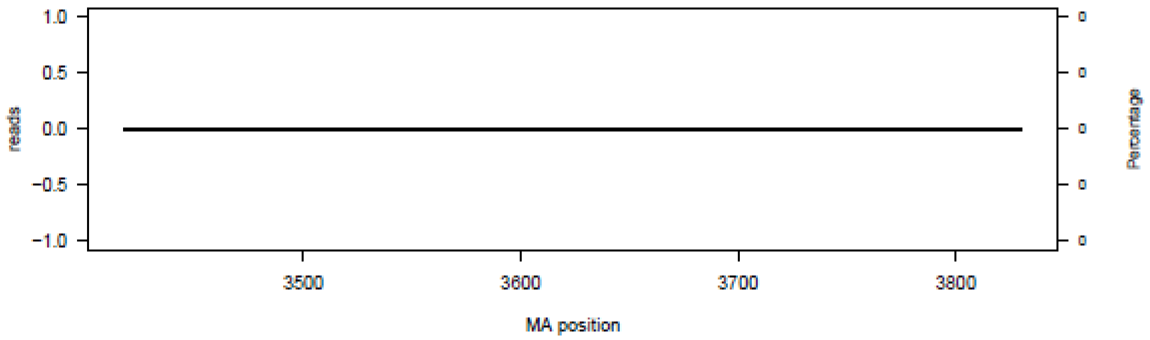

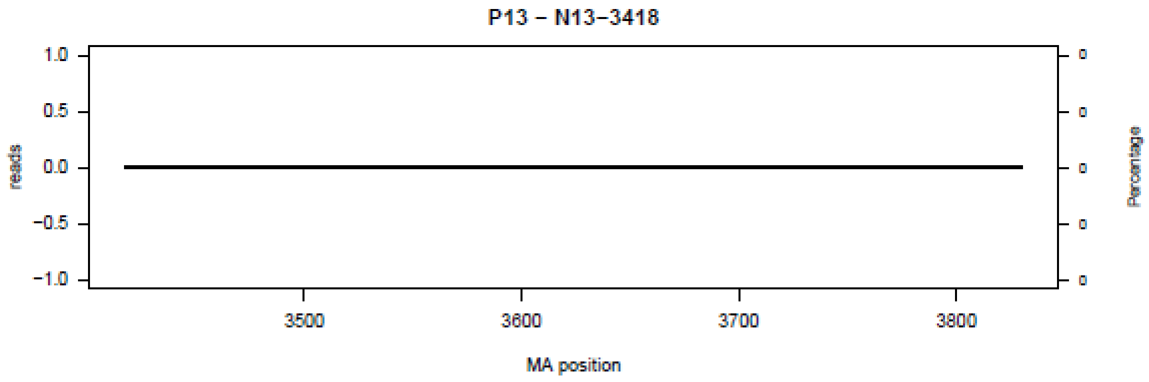
P13 – N13–3418
reads
Percentage
MA position
3500
3600
3700
3800

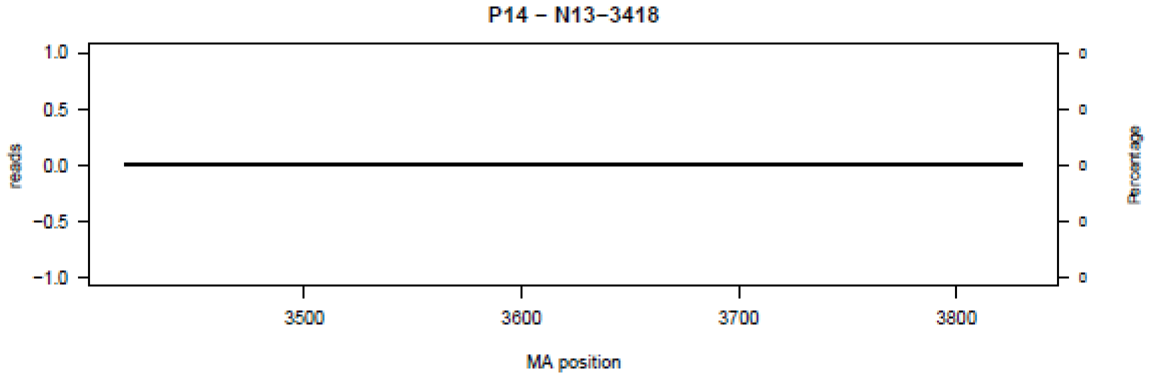
P14 – N13–3418
reads
Percentage
MA position
3500
3600
3700
3800

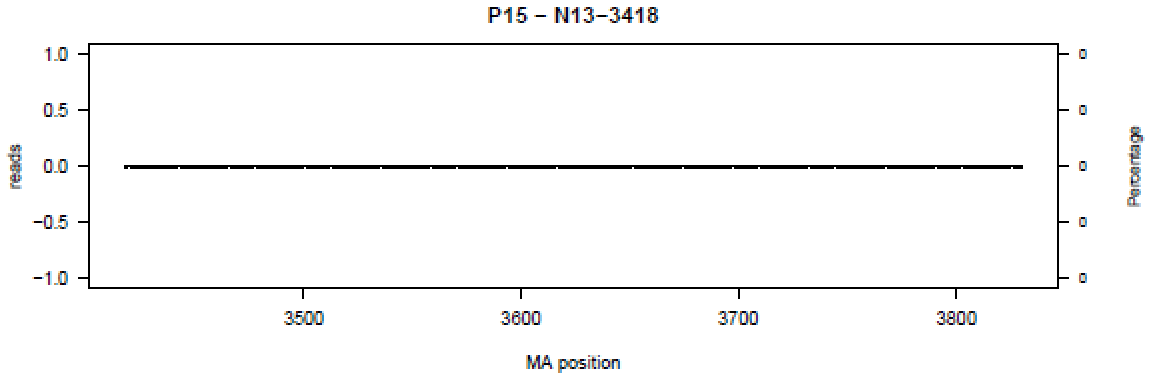
P15 – N13–3418
reads
Percentage
MA position
3500
3600
3700
3800

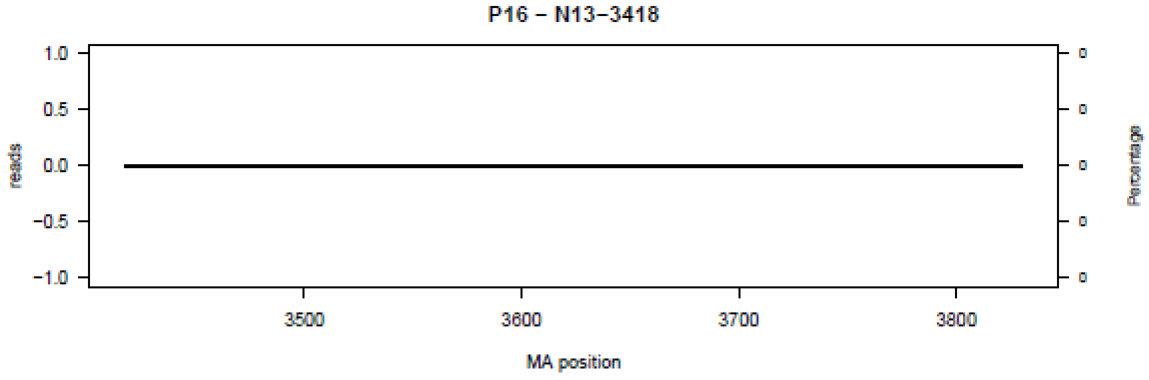
P16 – N13–3418
reads
Percentage
MA position
3500
3600
3700
3800

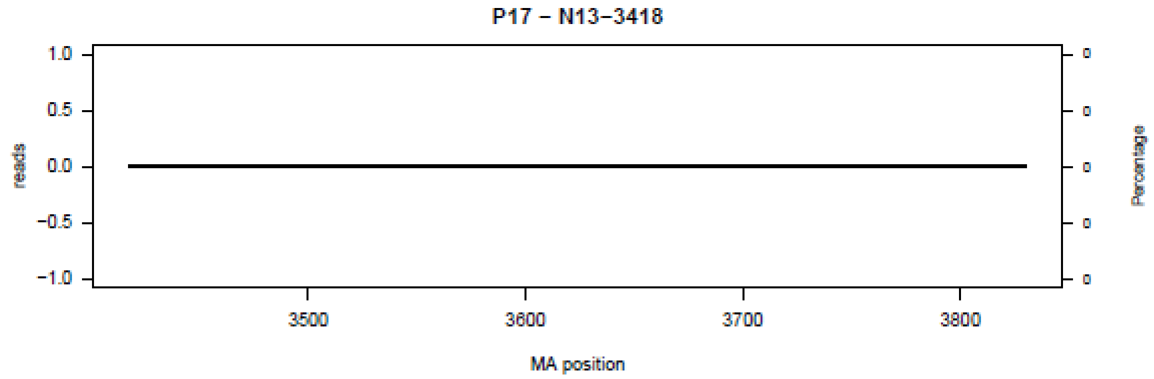
P17 – N13–3418
reads
1.0
0.5
0.0
−0.5
−1.0
3500
3600
3700
3800
MA position
Percentage
0
0
0
0
0

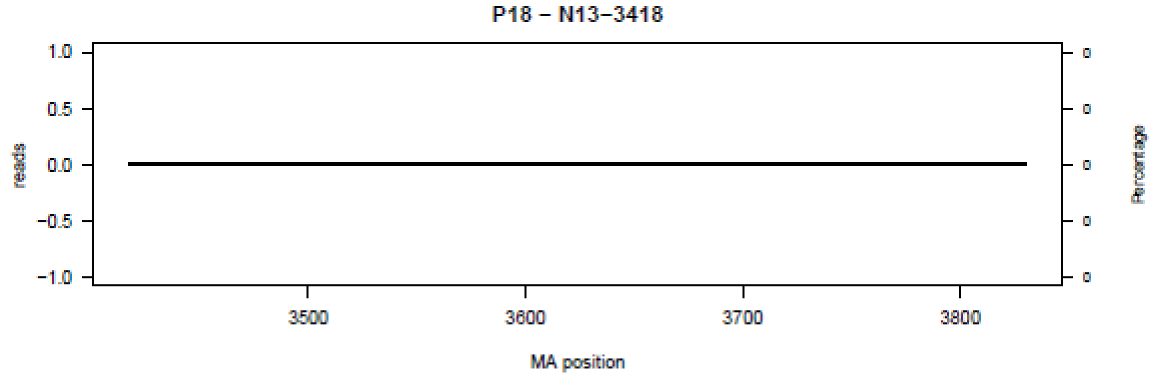
P18 – N13–3418
reads
1.0
0.5
0.0
−0.5
−1.0
3500
3600
3700
3800
MA position
Percentage
0
0
0
0
0

**Figure S14.** Bar plot summary of all deletions per nucleotide region.

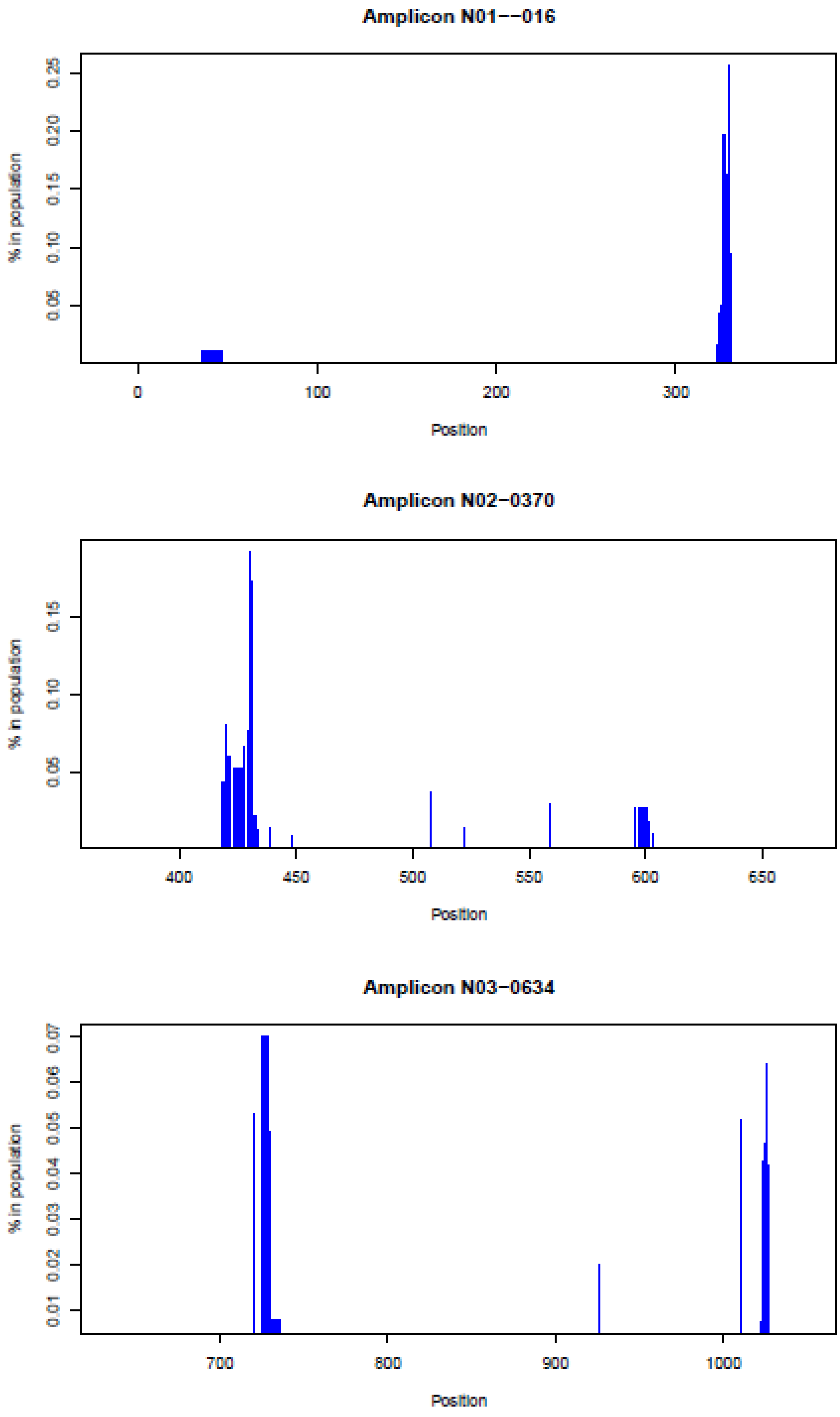

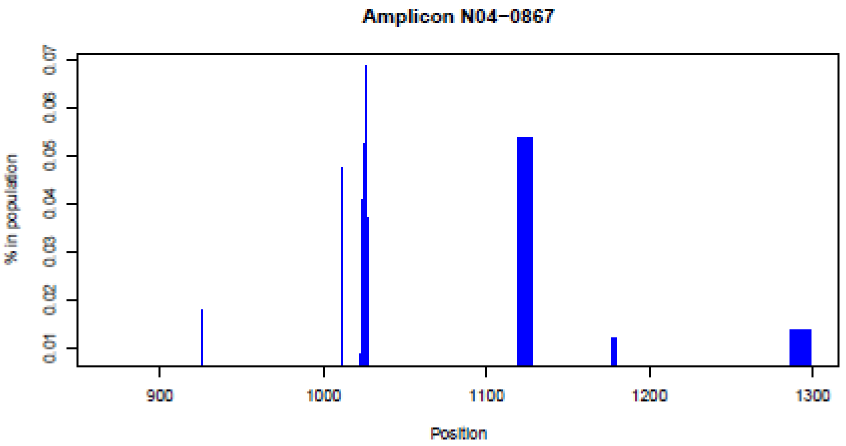
Amplicon N04−0867
% in population
0.01
0.02
0.03
0.04
0.05
0.06
0.07
900
1000
1100
1200
1300
Position

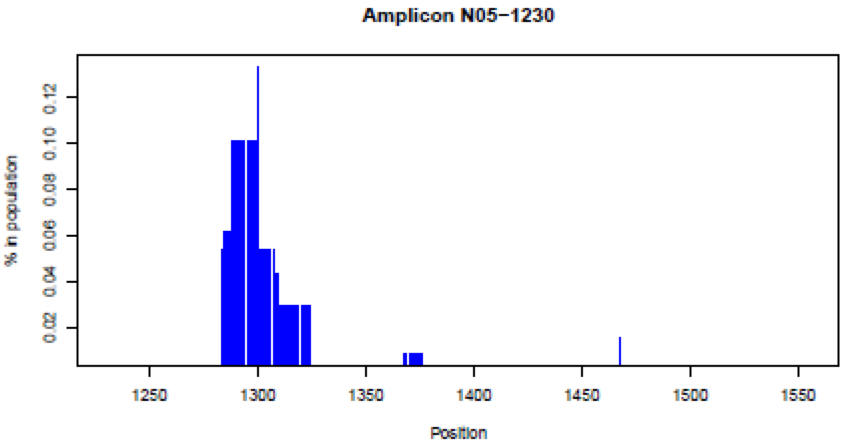
Amplicon N05−1230
% in population
0.02
0.04
0.06
0.08
0.10
0.12
1250
1300
1350
1400
1450
1500
1550
Position

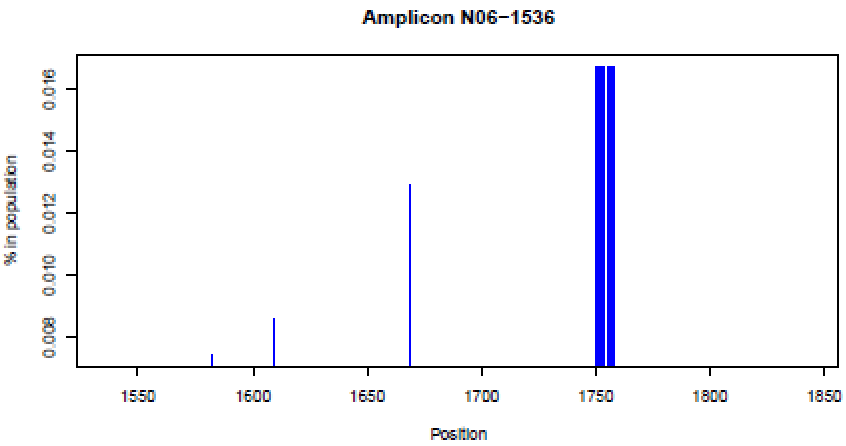
Amplicon N06−1536
% in population
0.008
0.010
0.012
0.014
0.016
1550
1600
1650
1700
1750
1800
1850
Position

Amplicon N07−1810

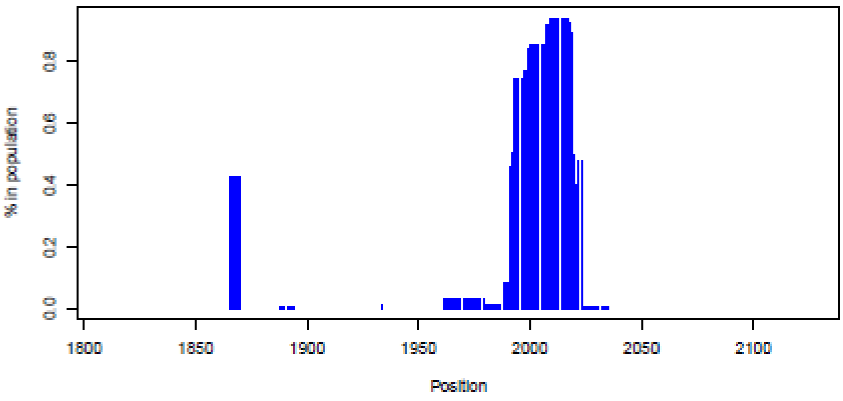


Amplicon N08−2101

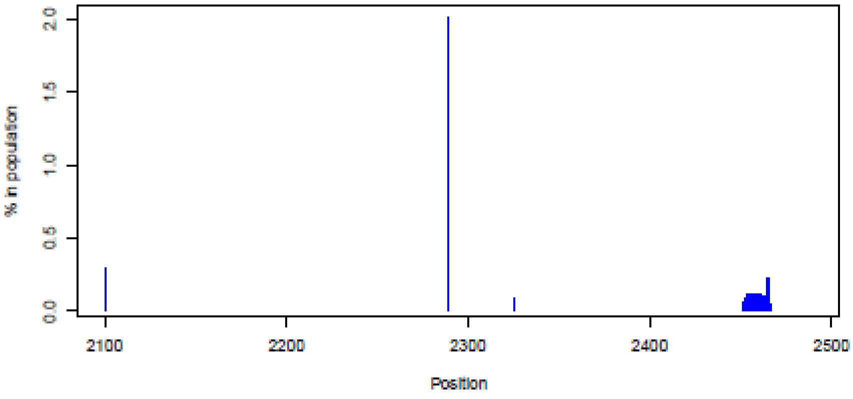


Amplicon N09−2447

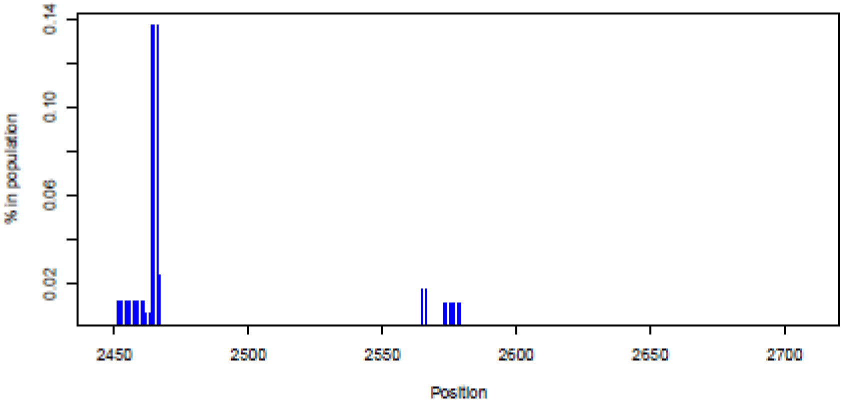

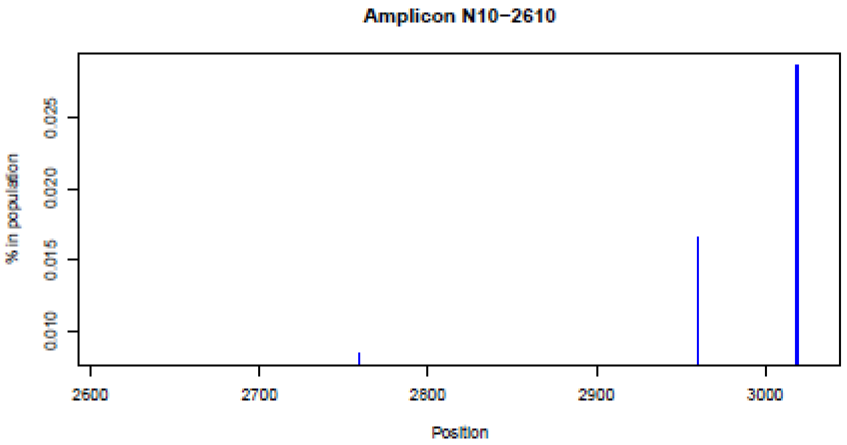
Amplicon N10−2610
% in population
0.010
0.015
0.020
0.025
2600
2700
2800
2900
3000
Position

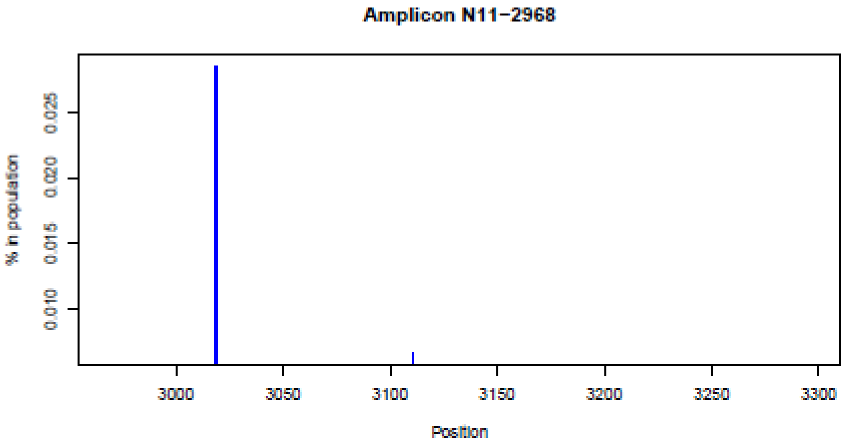
Amplicon N11−2968
% in population
0.010
0.015
0.020
0.025
3000
3050
3100
3150
3200
3250
3300
Position

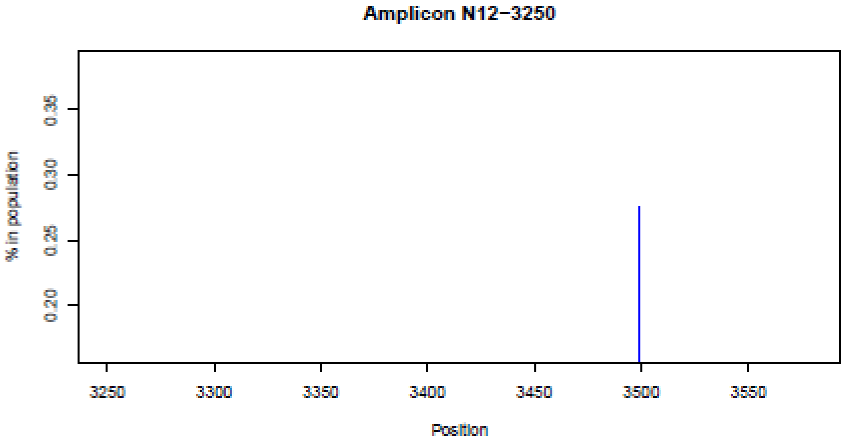
Amplicon N12−3250
% in population
0.20
0.25
0.30
0.35
3250
3300
3350
3400
3450
3500
3550
Position

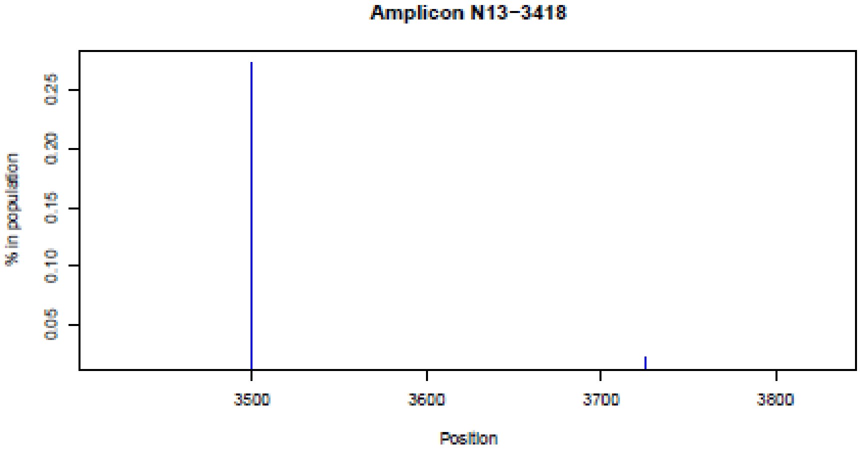
Amplicon N13−3418
% in population
0.05
0.10
0.15
0.20
0.25
3500
3600
3700
3800
Position

**SUPPLEMENTARY FIGURE S15**

**Figure S15.** Schematic amplicon design for the amplification of the whole RNA genomic spike gene, using MN908947.3 Wuhan-Hu-1 as a reference

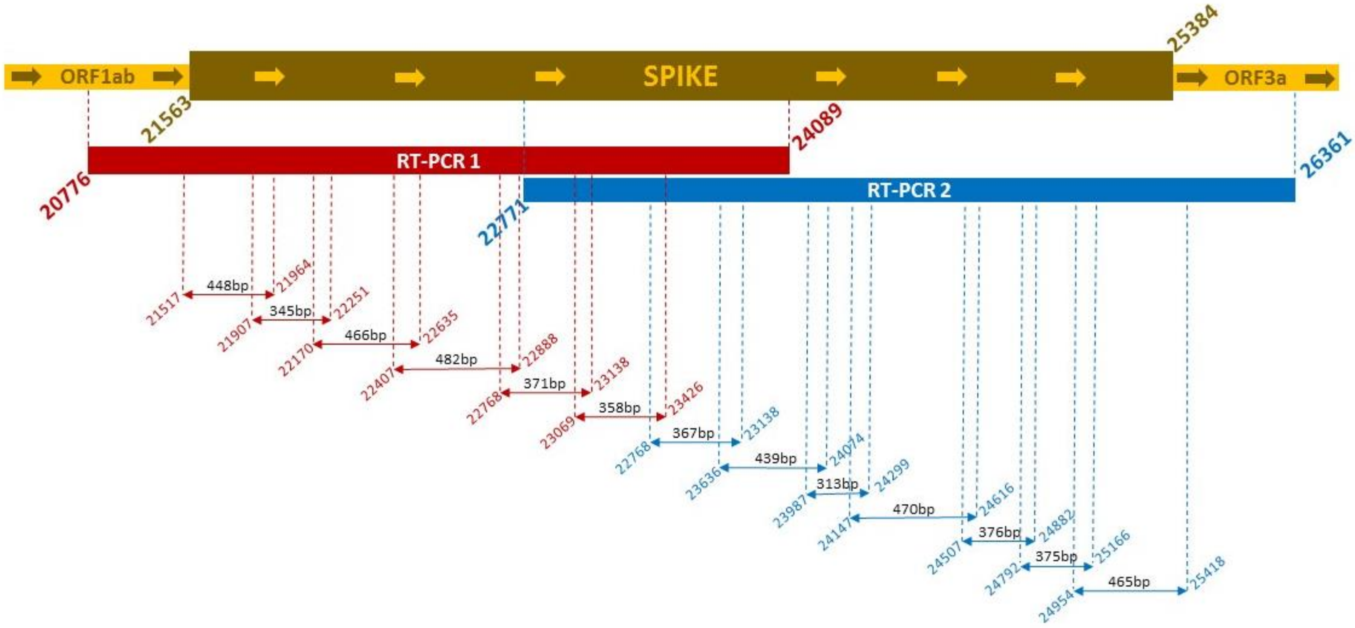
